# Genome-wide analysis of flavonoid biosynthetic genes in Musaceae (*Ensete*, *Musella*, and *Musa* species) reveals amplification of flavonoid 3’,5’-hydroxylase

**DOI:** 10.1101/2024.02.08.579575

**Authors:** Dongli Cui, Gui Xiong, Lyuhan Ye, Richard Gornall, Ziwei Wang, Pat Heslop-Harrison, Qing Liu

**Affiliations:** State Key Laboratory of Plant Diversity and Specialty Crops / Guangdong Provincial Key Laboratory of Applied Botany, South China Botanical Garden, Chinese Academy of Sciences, Guangzhou 510650, China; South China National Botanical Garden, Guangzhou 510650, China; University of Chinese Academy of Sciences, Beijing 100049, China; University of Leicester, Department of Genetics and Genome Biology, Institute for Environmental Futures, Leicester LE1 7RH, UK; Henry Fok School of Biology and Agriculture, Shaoguan University, Shaoguan 512005, China

**Keywords:** Anthocyanins, banana, diversification, F3’5’H, F3’H, flavonoids, genetics, genomics, ginger, monocotyledons, Musaceae

## Abstract

Flavonoids in Musaceae are involved in pigmentation and stress responses, including pathogen defense and UV protection, and are a component of the healthy human diet. Identification and analysis of the sequence and copy number of flavonoid biosynthetic genes are valuable for understanding the nature and diversity of flavonoid evolution in Musaceae species. In this study, we identified 71 to 80 flavonoid biosynthetic genes in chromosome-scale genome sequence assemblies of Musaceae, including those of *Ensete glaucum*, *Musella lasiocarpa*, *Musa beccarii*, *M. acuminata*, *M. balbisiana*, and *M. schizocarpa,* checking annotations with BLAST and determining the presence of conserved domains. The number of genes increased through segmental duplication and tandem duplication. Orthologues of both structural and regulatory genes in the flavonoid biosynthetic pathway are highly conserved across Musaceae. The flavonoid 3’,5’-hydroxylase gene *F3’5’Hs* was amplified in Musaceae and ginger compared with grasses (rice, Brachypodium, *Avena longiglumis,* and sorghum). One group of genes from this gene family amplified near the centromere of chromosome 2 in the *x* = 11 Musaceae species. Flavonoid biosynthetic genes displayed few consistent responses in the yellow and red bracts of *Musella lasiocarpa* when subjected to low temperatures. The expression levels of *MlDFR2/3* increased while *MlLAR* was reduced by half. These results establish the range of diversity in both sequence and copy number of flavonoid biosynthetic genes during evolution of Musaceae. The combination of allelic variants of genes, changes in their copy numbers, and variation in transcription factors with the modulation of expression under cold treatments and between bract-colours suggests the variation may be exploited in plant breeding programmes, particularly for improvement of stress-resistance in the banana crop.

## Introduction

The Musaceae comprises three genera: *Ensete* (7 spp.), *Musella* (1 sp.), and *Musa* (c. 83 spp.) (Cheesman 1947; Häkkinen 2013), distributed across southeastern Asia and Africa (Janssens *et al*., 2016). Cold resistance occurs especially in Asian *Musella lasiocarpa* and *Ensete glaucum*, which grow at 1500–2500 meters and 800–1100 meters above sea level, respectively (Li, 1978). *Musa* and *Ensete* are important crops (Christelová *et al*., 2011; Amah *et al*., 2020) that provide a source of carbohydrates to tropical communities. The worldwide production of bananas and plantains reached 179,263,140 tons in 2022 (https://www.fao.org/faostat/, accessed 30 January 2024). The basic chromosome number in *Ensete* and *Musella* is considered to be *x*=9, with all the species being diploid (Cheesman 1947; Li 1978; Li 1979; Zhang *et al*., 2018; Wang *et al*., 2022; Wang *et al*., 2023). *Musa* has *x*=9, 10, and 11 species, including diploids, with triploids and tetraploids present in some species and as hybrids (Wang *et al*., 2019). The wild species are diploids with a signature of ancient genome triplication, while most cultivated lineages of banana and plantains are sterile parthenocarpic triploids (Heslop-Harrison *et al*., 2007, Passos *et al*., 2012). Genetic diversity has been exploited in all three genera through selection for cultivation as food (banana and starch ensete) (Neumann and Hildebrand 2009; Borrell *et al*., 2019), potential medicines (Ranjha *et al*., 2022), and horticulture (Nugraha *et al*., 2023).

Flavonoids occur in all photosynthetic lineages, from cyanobacteria, haptophytes, ochrophytes, rhodophytes and chlorophytes to bryophytes and vascular plants (Goiris *et al*. 2014). These compounds are synthesized in the cytosol and transported into the vacuole, where they can function as bioactive molecules (Rausher 2006; Zhao 2015; Davies *et al*., 2020). Flavonoids are important to plants for the production of colours, especially flowers and fruits, in the context of pollination and seed dispersal (Giusti *et al*., 2023); pollen, spore and seed germination; and seedling development (Wang *et al*., 2020). Flavonoids protect plants from different abiotic and biotic stresses, including by acting as UV filters for young growth and playing roles in frost hardiness and drought resistance (Samanta *et al*., 2011; Ferreyra *et al*., 2021). They are important in defence against pathogens, pests, and herbivores (Mahajan *et al*., 2011; van der Kooi *et al*., 2019; Tiku *et al*., 2020). Attention has been given to the potential nutraceutical, pharmaceutical, medicinal and cosmetic applications and benefits of flavonoids due to their antioxidative, antiviral, antimicrobial, inflammatory, mutagenic, and anticarcinogenic properties (Someya *et al*., 2002; Panche *et al*., 2016). In particular, *Ensete ventricosum* has a wide range of therapeutic uses, and landraces are cultivated for use in traditional medicine (Nuraga *et al*., 2022), while various relatives of Zingiberales (including turmeric and ginger) are considered to have health benefits.

Flavonoids are the largest class of polyphenols, comprising 9000 kinds in the plant kingdom (Bate-Smith 1954; Williams *et al*., 2004). These compounds bear a common diphenylpropane (C6-C3-C6) backbone in which two aromatic rings are linked via a three-carbon chain (Saito *et al*., 2013) (Fig. 1; inset flavan nucleus). Flavonoids consist of a benzene ring (A) linked to a pyrone ring (C), which at the 2 or 3 position takes a phenyl ring (B) as a substitute (Fig. 1); based on the oxygenation pattern of the heterocyclic C ring, they can be sub-classified into six groups, namely flavones, flavonols, flavanones, flavanols, anthocyanidins, and isoflavones (Alseekh *et al*., 2020; Wen *et al*., 2020). Flavonoid biosynthesis has been studied extensively and the pathways to the various compounds are well-established (Shirley 1996; Ng and Smith 2016; Chu *et al*., 2017; Zhao *et al*., 2020; Manzoor *et al*., 2023). The genes controlling the individual processes have been characterized in detail (Fig. 1; Falcone Ferreyra *et al*., 2012; Peng *et al*., 2020). The biosynthetic pathway starts with phenylalanine via the general phenylpropanoid pathway (GPP) (Wei *et al*., 2023). Three major gene families, PAL, C4H, and 4CL, are involved in the production of p-coumaroyl-CoA via cinnamic acid (precursor to salicylic acid). The resulting structural genes can be divided into two groups (Guo *et al*., 2014). Early Biosynthetic Genes (EBGs), which include a group of hydroxylases and synthases and Late Biosynthetic Genes (LBGs), which include reductases (Lepiniec *et al*., 2006; Petroni and Tonelli 2011; Pucker *et al*., 2020). In the LBG group, glycosyl transferase genes (UFGT) add the finishing modifications to the molecules. The structural genes have been identified in many species, including Musaceae (Pucker et al., 2020; Rijzaani et al., 2022; Busche et al., 2023).

**Fig. 1.**
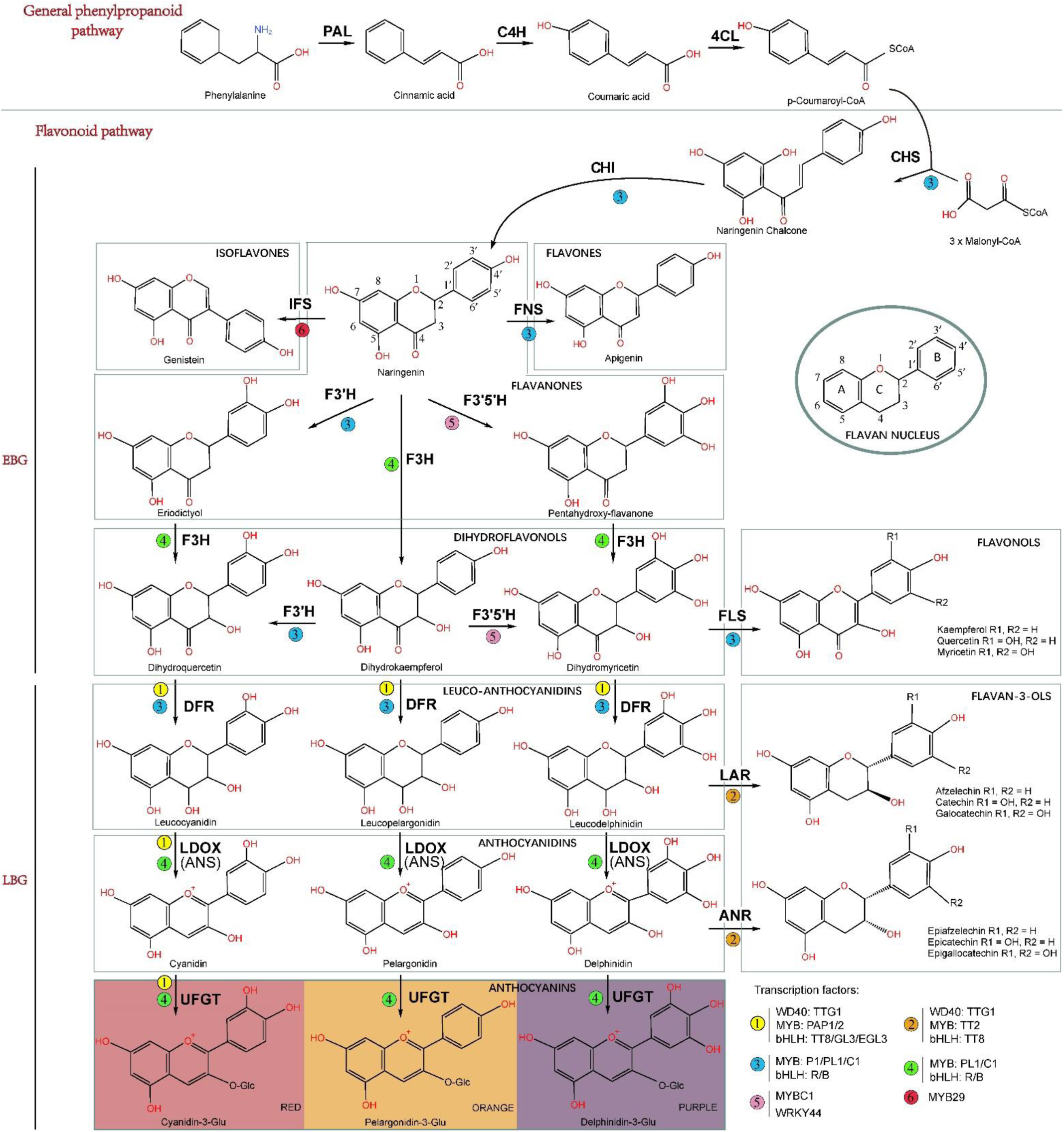
The part of the flavonoid biosynthetic pathway analysed here, with regulators of the genes, in plant cells (based on maize, Arabidopsis, *Brassica, Musa*, kiwifruit, and soybean). Structural genes encoding enzymes are: PAL, phenylalanine ammonia lyase; C4H, cinnamic acid 4-hydroxylase; 4CL, 4 coumarate CoA ligase; CHS, chalcone synthase; CHI, chalcone isomerase; FNS, flavone synthase; IFS, isoflavone synthase; F3H, flavanone 3-hydroxylase; F3’H, flavanone 3’-hydroxylase; F3’5’H, flavanone 3’5’- hydroxylase; DFR, dihydroflavonol reductase; FLS, flavonol synthase; ANS/LDOX, anthocyanidin synthase/leucoanthocyanidin dioxygenase; UGP/UFGT, UDP-flavonoid glucosyl transferase; ANR, anthocyanidin reductase; LAR, leucoanthocyanidin reductase. EBG: early biosynthetic genes; LBG: late biosynthetic genes. Regulatory transcription factors MYB (P1/PL1/C1 or PL1/C1) and bHLH (R/B) are from maize; WD40 (TTG1), MYB (TT2/PAP1/PAP2), and bHLH (TT8/GL3/EGL3) are from Arabidopsis; MYBC1 and WRKY44 are from kiwifruit; and MYB29 is from soybean

Three transcription factor (TF) families, R2R3-MYB, bHLH, and WD40, comprise the ternary MBW complexes which regulate different branches of the pathway by activating a subset of biosynthetic genes (Wilkins et al., 2009; Zheng *et al*., 2019; Pucker 2022). The expression of several genes has been studied; for example, the expression of the *Musa F3’5’H-1* gene is strongly affected by abiotic stress conditions and following application of phytohormones to banana plants (Negi *et al*., 2023). Transcriptome expression profiling, e.g. in *Musa* (Mingmanit *et al*., 2023; Liu *et al*. 2023) is increasingly used to complement studies of gene alleles and copy number variations.

Both structural and regulatory genes in the flavonoid pathway have served as model systems for understanding a variety of evolutionary processes, such as gene duplication, evolutionary rate variation among genes, and the relative importance of structural and regulatory genes in the evolution of ecologically important characters (Rausher 2006), including studies of *Musa acuminata* (Pandey *et al*., 2016), *Oryza sativa* (rice) (Xia *et al*., 2021), *Arabidopsis thaliana* (Arabidopsis), and *Brassica rapa* (Guo *et al*., 2014). Analysis of their evolution in both the sequence and copy number of these genes is valuable for phylogenetic reconstruction.

In the present study, we aimed to perform a comparative analysis of flavonoid biosynthetic genes in Musaceae based on recent publications of chromosome-scale genome assemblies and transcriptome data (Droc *et al*., 2022). Interspecific variations in gene copy number, structural organization, genomic location, and the nature and expression of key structural and regulatory genes were investigated, especially in the context of response to low temperatures.

## Materials and Methods

### Identification of putative flavonoid biosynthetic genes

The genome and inferred protein sequences of *Oryza sativa*, *Sorghum bicolor*, *Zingiber officinale*, *Ensete glaucum*, *Musella lasiocarpa*, *Musa beccarii*, *M. acuminata*, *M. balbisiana*, and *M. schizocarpa* were downloaded from the Banana Genome Hub (https://banana-genome-hub.southgreen.fr/content/download) (Droc *et al*., 2022); *Brachypodium distachyon* protein sequences were downloaded from NCBI (PRJNA32607; https://www.ncbi.nlm.nih.gov); and *Avena longiglumis* protein sequences (PRJNA956334) were downloaded from Liu *et al*. (2022) to build 11 local protein databases.

Hidden Markov Model (HMM) profiles of the characterized and conserved domains of flavonoid biosynthetic genes were retrieved from the protein family database (https://www.ebi.ac.uk/interpro/) (Mistry *et al*., 2021; Paysan-Lafosse *et al*., 2023) to search for these gene families in the Musaceae. The flavonoid biosynthetic protein sequences were identified based on the HMM profiles using HMMER software with default parameters (Finn *et al*., 2011) and a cut-off value of 0.01 (Xie *et al*., 2018). To ensure the accuracy of the identification of flavonoid biosynthetic genes, we further blasted the flavonoid protein sequences of Arabidopsis, rice and Brachypodium (Qian *et al*., 2011) against the *E. glaucum*, *Musella lasiocarpa*, *Musa beccarii*, *M. acuminata*, *M. balbisiana*, and *M. schizocarpa* genome databases. The conserved domains of all the candidate genes were examined using the Pfam and Simple Modular Architecture Research Tool (SMART, http://smart.embl-heidelberg.de/) programs. The sequence length, molecular weight and isoelectric point of the flavonoid biosynthetic proteins were predicted using tools from ExPasy (https://web.expasy.org/protparam/).

Artefacts arise from automated genome-wide annotation, with Bartas *et al*. (2018) noting an apparent triplication of the CHS domain in *M. balbisiana*. Here, we found one (*M. schizocarpa F3’H*; *M. balbisiana F3’5’H*, and *DFR*) or two (*M. schizocarpa F3’5’H*) annotated genes which could not be validated and Table 1 shows the revised gene numbers.

**Table 1.** Number of flavonoid genes in four species of *Musa* and one each of *Musella*, *Ensete*, *Zingiber*, *Avena*, *Brachypodium*, *Oryza*, Sorghum, and Arabidopsis.

| Gene family | *Musa acuminata*<br>sect. *Musa*<br>(Musaceae)<br>2n=2x=22 | *Musa balbisiana*<br>sect. *Musa*<br>(Musaceae)<br>2n=2x=22 | *Musa schizocarpa*<br>sect. *Musa*<br>(Musaceae)<br>2n=2x=22 | *Musa beccarii*<br>sect.<br>*Callimusa*<br>(Musaceae)<br>2n=2x=18 | *Musella lasiocarpa*<br>(Musaceae)<br>2n=2x=18 | *Ensete glaucum*<br>(Musaceae)<br>2n=2x=18 | *Zingiber officinale*<br>(Zingiberaceae)<br>2n=2x=22 | *Avena longiglumis*<br>(Poaceae)<br>2n=2x=14 | *Brachypodium distachyon*<br>(Poaceae)<br>2n=2x=10 | *Oryza sativa*<br>(Poaceae)<br>2n=2x=24 | *Sorghum bicolor*<br>(Poaceae)<br>2n=2x=20 | *Arabidopsis thaliana*<br>(Brassicaceae)<br>2n=2x=10 |
| --- | --- | --- | --- | --- | --- | --- | --- | --- | --- | --- | --- | --- |
| *Structural genes* |  |  |  |  |  |  |  |  |  |  |  |  |
| PAL<sup>a</sup> | 8 | 7 | 8 | 8 | 9 | 9 | 6 | 26 | 8 | 12 | 8 | 4 |
| C4H<sup>a</sup> | 4 | 4 | 3 | 4 | 4 | 4 | 3 | 5 | 2 | 1 | 2 | 1 |
| 4CL<sup>a</sup> | 18 | 16 | 18 | 18 | 18 | 16 | 24 | 24 | 14 | 16 | 17 | 4 |
| CHS<sup>b</sup> | 6 | 8 | 5 | 5 | 4 | 6 | 7 | 1 | 1 | 2 | 10 | 1 |
| CHI<sup>b</sup> | 2 | 2 | 2 | 2 | 2 | 2 | 2 | 2 | 2 | 3 | 2 | 1 |
| F3H<sup>b</sup> | 2 | 2 | (1)2 | 2 | 2 | 2 | 1 | 1 | 1 | 1 | 2 | 1 |
| F3'H<sup>b</sup> | 1 | 1 | 1 | 1 | 1 | 1 | 2 | 2 | 4 | 2 | 5 | 1 |
| F3'5'H<sup>b</sup> | 13 | (13)14 | (8)10 | 10 | 12 | 10 | 8 | 1 | 2 | 1 | 2 | 7 |
| FLS<sup>b</sup> | 2 | 5 | 3 | 4 | 5 | 4 | 3 | 3 | 1 | 1 | 2 | 6 |
| DFR<sup>c</sup> | 3 | (2)3 | 3 | 3 | 3 | 3 | 1 | 1 | 2 | 1 | 4 | 1 |
| LDOX<sup>c</sup> | 1 | 1 | 1 | 1 | 1 | 1 | 1 | 2 | 0 | 2 | 1 | 1 |
| LAR<sup>c</sup> | 1 | 1 | 1 | 1 | 1 | 1 | 0 | 1 | 1 | 1 | 0 | 0 |
| ANR<sup>c</sup> | 5 | 1 | 3 | 1 | 4 | 2 | 4 | 19 | 8 | 5 | 2 | 1 |
| UGT78D2<sup>c</sup> | 3 | 3 | 3 | 3 | 3 | 3 | 5 | 1 | 2 | 2 | 2 | 1 |
| UGT75C1<sup>c</sup> | 4 | 3 | 4 | 3 | 3 | 3 | 7 | 9 | 9 | 6 | 9 | 1 |
| UGT79B1<sup>c</sup> | 1 | (0)1 | 1 | 1 | 1 | 1 | 1 | 1 | 2 | 1 | 3 | 1 |
| *Regulatory genes (transcription factors)* |  |  |  |  |  |  |  |  |  |  |  |  |
| bHLH: TT8 (R1) | 2 | 2 | 2 | 2 | 2 | 2 | 1 | 1 | 3 | 1 | 1 | 1 |
| WD40: TTG1 (PAC1) | 2 | 2 | 2 | 2 | 2 | 2 | 3 | 1 | 1 | 1 | 1 | 1 |
| MYB: C1 | 2 | 3 | 2 | 3 | 3 | 2 | 5 | 2 | 1 | 1 | 1 | 0 |
| Total | 80 | 76 | 71 | 74 | 80 | 74 | 84 | 103 | 64 | 60 | 74 | 34 |

<sup>a</sup> Flavonoid precursor; <sup>b</sup> early-pathway; <sup>c</sup> late-pathway. The red number in parentheses is the original published annotated gene copy number, with the corrected annotation number next to it (c.f. Bartas *et al.*, 2018).

### *In-silico* analysis of promoter cis-acting elements

A 1,500-bp (base pair) upstream region from the initial codon of each candidate flavonoid biosynthetic gene was used to search cis-elements with PlantCARE (http://bioinformatics.psb.ugent.be/webtools/plantcare/ html/) (Lescot *et al*., 2002).

### Chromosomal localization and phylogenetic analysis

The physical locations of flavonoid biosynthetic genes were determined in *E. glaucum, Musella lasiocarpa, Musa beccarii, M. acuminata, M. balbisiana*, and *M. schizocarpa* genome databases and displayed in MapChart (Voorrips *et al*., 2022). Multiple sequence alignments with the respective protein families in Arabidopsis, *B. distachyon*, *A. longiglumis*, *O. sativa*, *S. bicolor*, *Z. officinale*, *E. glaucum*, *Musella lasiocarpa*, *Musa beccarii*, *M. acuminata*, *M. balbisiana* and *M. schizocarpa* were performed using ClustalW in MEGA 7.0 (Kumar *et al*., 2018). Phylogenetic trees were constructed using a Neighbor-Joining (NJ) method with 1,000 bootstrap replications.

### Analysis of collinearity and duplication

Gene duplication events and collinearity (conserved synteny) were analysed using the Multiple Collinearity Scan toolkit (MCScanX). Amino acid sequences were analysed using BLASTP with an e-value of 1.0 ×10^−5^. The collinear blocks were detected by submitting a whole-genome gff file (downloaded from Banana Genome Hub) and BLASTP results to MCScanX. The positions of gene pairs of flavonoid biosynthetic genes were mapped using Circos tool (Krzywinski *et al*., 2009).

### Evolutionary analysis of FBGs

Coding sequences (CDSs) of orthologous flavonoid flavonoid biosynthetic gene pairs between *E. glaucum*, *Musella lasiocarpa*, *Musa beccarii*, *M. acuminata*, *M. balbisiana*, and *M. schizocarpa* were aligned using ClustalW. The synonymous (Ks) and non-synonymous (Ka) substitution rates were calculated using the Ka/Ks_calculator 2.0 (Wang *et al*., 2010). The divergence time (T) was estimated using the equation T = Ks/2r, where “r” represents the synonymous substitution rate (4.5 per 10^9^ years) (Lescot *et al*., 2008).

### Plant materials and temperature stress treatments

Four-year-old *Musella lasiocarpa* var. *lasiocarpa* (yellow bracts) and *M. lasiocarpa* var. *rubribracteata* (red bracts) originated from Puge city (E 102.68°, N 23.45°), Liangshan Yi Autonomous Prefecture, Sichuan Province, China. The plants were subsequently grown in a growth cabinet at the South China Botanical Garden for 16 hours at 70% relative humidity under two temperature regimes: 15°C in the low-temperature-treated group and 25°C in the control group. After 48 hours of treatment, the bracts were collected. Three replicates were conducted for each species and two treatments. The specimens were frozen in liquid nitrogen for RNA extraction and kept at −80°C.

### RNA-seq-based gene expression analysis

RNA was extracted from the bracts of *Musella lasiocarpa* using an ENZA Plant RNA Kit (R6827, Omega, Guangzhou). The library was constructed using the MGIEasy RNA Library Prep Kit v.3.0 (BGI, Shenzhen). Transcriptome sequencing was performed by cPAS (Combinatorial probe-anchor synthesis) using the high-throughput sequencing platform DNBSEQ-T7 (MGI) and the RNA-seq data can be obtained from the Genome Sequence Archive of the China National Center for Bioinformation project PRJCA023025. Low-quality sequences and splice sequences in raw reads were filtered using the software fastp.v.0.21.0 (Chen *et al*., 2018) to obtain clean reads. The filtered reads were aligned to the *Musella lasiocarpa* genome (https://banana-genome-hub.southgreen.fr/node/50/942) by STAR (Dobin *et al*., 2015), and the raw number of aligned reads was calculated using the feature Counts software (Liao *et al*., 2014). The TPM (transcripts per million) value was calculated using the edgeR package (Robinson *et al*., 2010) and normalized using the TMM (trimmed mean of M-values) method. A heat map of the flavonoid biosynthetic gene expression pattern was constructed based on log_2_ (TPM + 1) values.

## Results

### Identification and number of flavonoid biosynthetic genes

To identify the flavonoid biosynthetic genes and regulatory sequences (transcription factors) in Musaceae, we gathered the conserved domains of these gene families and used HMMER software (Pucker *et al*., 2020) to search for corresponding domains in genome assemblies (removing any allelic variants at single loci) from six Musaceae species (*Ensete glaucum, Musella lasiocarpa*, *Musa beccarii*, *M. acuminata*, *M. balbisiana*, and *M. schizocarpa*). For comparisons, we also used flavonoid genes identified in the genome assemblies of Zingiberaceae (*Zingiber officinale,* ginger), four species of Poaceae (*Oryza sativa*, rice; *Brachypodium distachyon*, Brachypodium; *Avena longiglumis*; *Sorghum bicolor*, Sorghum; and the dicot *Arabidopsis thaliana,* Arabidopsis, all diploid species. Almost all genes and transcription factors were found in all analysed plant families (Table 1). The number of biosynthetic structural genes in the Musaceae ranged from 65 to 74 and the number of regulatory genes ranged from 6 to 7, for a total of 71 to 80 flavonoid biosynthetic genes and transcription factors. These numbers are broadly similar to the numbers found in *Zingiber* (84) and Poaceae (64–103) species but two to three times greater than the number found in Arabidopsis (Tables 1, S1; detailed characteristics, including protein properties and subcellular localization prediction, are provided in Table S2). There were more early-than late-stage biosynthetic genes in each Musaceae species, with early/late ratios ranging from 1.25–2.82 (Table 1). This finding is similar to the situation in Arabidopsis. In contrast, ginger and three of the four Poaceae species have fewer early biosynthetic genes than late ones, with early/late ratios of 0.55 in ginger and 0.29–0.56 in the three grasses. The fourth grass species, sorghum, is an exception, with a ratio of 1.10.

Although the total numbers of genes were similar across the monocot species represented, there were marked differences in the copy numbers of individual genes; this phenomenon was especially pronounced among the structural genes but less pronounced among the regulatory genes. Although there has been extensive multiplication of PAL and ANR in *Avena longiglumis* (Table 1), within the Musaceae, although the copy numbers of each gene are relatively constant and even invariant among the taxa, there is evidence of multiplication in the CHS and especially F3’,5’H gene families when compared with the Poaceae. In contrast, there are fewer copies of the F3’H and UGT75C1 families in Poaceae than in Zingiberales. Among the transcription factor families, Musaceae had two or three members, while Poaceae usually had fewer. LAR and MYB (C1) have not been reported in Arabidopsis (Table 1).

### Chromosomal distribution of flavonoid biosynthetic genes

The flavonoid biosynthetic genes are distributed across all chromosomes in the six Musaceae species (Fig. S1), with some tandem duplications (shown in boxes in Fig. S1; in some cases, the genes were adjacent but not recognized as tandem duplications). Most of the flavonoid biosynthetic genes were distributed near the ends of the chromosome arms, except for the F3’5’H genes. There is a cluster of F3’5’H genes near the centromere of chromosome 2 in the *x*=11 *Musa* species, *M. acuminata, M. balbisiana, and M. schizocarpa*.

### Phylogenetic analysis of flavonoid biosynthetic genes in Musaceae and outgroups

Phylogenetic trees were constructed using the amino acid sequences of 17 flavonoid biosynthetic genes in six Musaceae species, ginger, four grasses, and Arabidopsis (Figs 2 and S2). For Musaceae, 50 out of 72 terminal branches had a gene from all six species, showing multi-copy gene duplication and that some diversification occurred before speciation. The phylogeny of the *F3’5’H* gene family (amplified in Musaceae; Table 1) was more complex, revealing a new clade of *Musella lasiocarpa* genes with tandem duplication of the genes (Fig. 2). Around this clade, there were notable clusters of *F3’5’H* around the chromosome 2 centromere in *Musa acuminata*, *M. balbisiana*, and *M. schizocarpa* (Fig. 2).

**Fig. 2.**
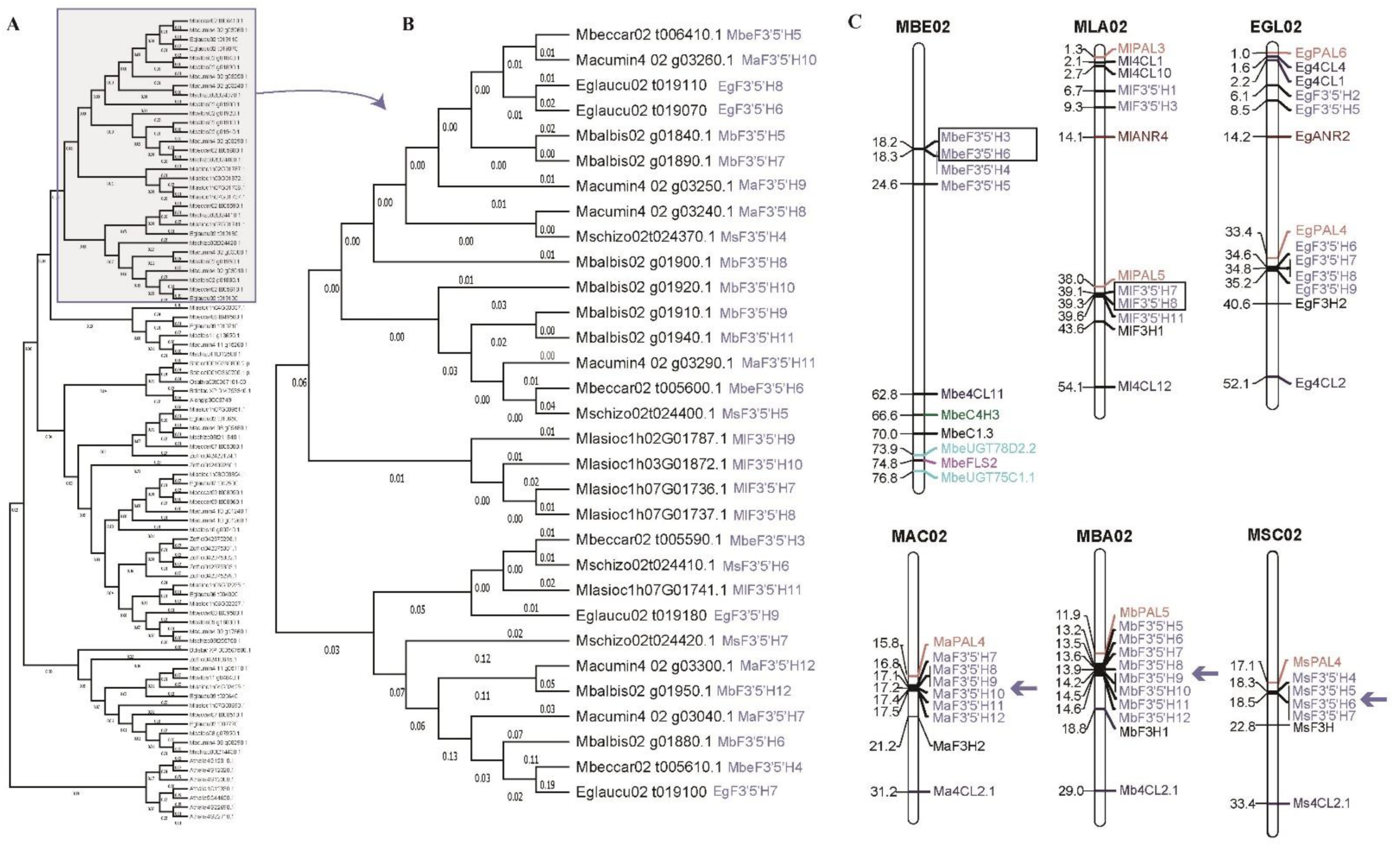
Phylogenetic tree and chromosomal location of *F3’5’Hs.* (**A**) Phylogenetic tree of the F3’5’H gene family of Musaceae plus rice, Brachypodium, *Avena longiglumis*, sorghum, ginger and Arabidopsis as outgroups. (**B**) One clade of the F3’5’H phylogenetic tree terminal with genus (initial letter) and species (six letter) abbreviations and (**C**) Chromosome 2 gene location maps in Musaceae. Genes from different families are shown in different colors (F3’5’H in purple). Horizontal bars represent the gene locations (Mb, megabases) on syntenic chromosome 2 in Fig. 2 **C**. Genes with tandem duplications are framed by black boxes; F3’5’H clusters near the chromosome 2 centromere are indicated by arrows. MBE, Mbeccar, *M. beccarii*; MLA, Mlasioc, *M. lasiocarpa*; EGL, Eglaucu, *E. glaucum*; MAC, Macumin, *M. acuminata*; MBA, Mbalbis, *M. balbisiana*; MSC, Mschizo, *M. schizocarpa*

### Identification of duplicated (paralogous) gene pairs of flavonoid biosynthetic genes in Musaceae

The Musaceae family shares ancestral whole-genome duplication events, with three copies of most of the genes being found (D’Hont *et al*., 2012; Wang *et al*., 2022). We analysed the locations and changes in the copy numbers of paralogous flavonoid biosynthetic genes with respect to genome reorganization and chromosome recombination and identified 17 to 25 segmental gene duplications and 3 to 5 tandemly duplicated gene pairs in the *E. glaucum, M. lasiocarpa, M. beccarii*, *M. acuminata*, *M. balbisiana,* and *M. schizocarpa* genomes (Fig. 3 and Table S4). In these six species, genes in the PAL, C4H, 4CL, CHS, and UFGT families always exhibited segmental duplication events. Tandem duplication events also played a part in UFGT amplification in Musaceae.

**Fig. 3.**
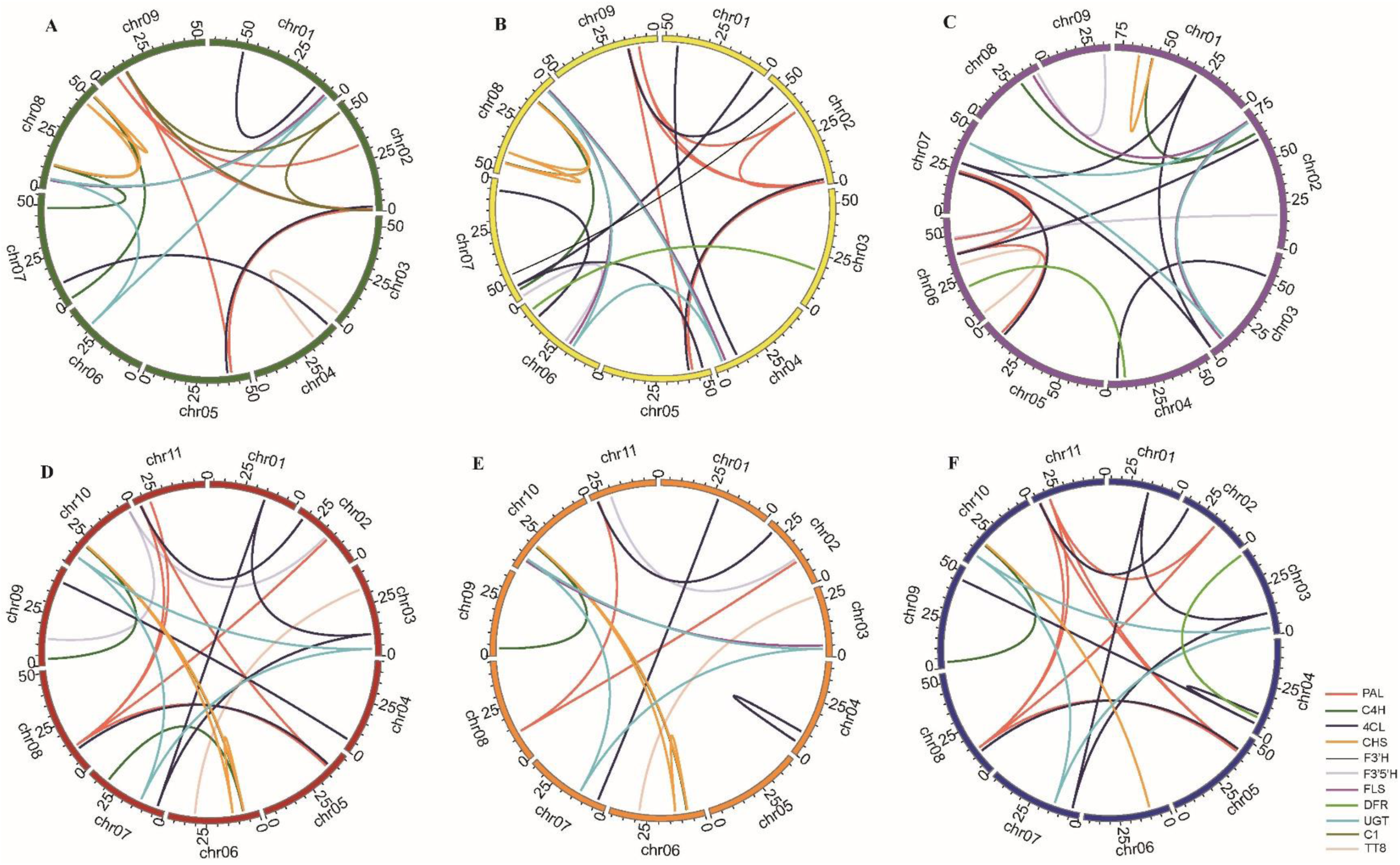
Circos plot of flavonoid biosynthetic gene paralogue locations on the chromosomes of six Musaceae species. (**A**) *E. glaucum* (green). (**B**) *M. lasiocarpa* (yellow). (**C**) *M. beccarii* (purple). (**D**) *M. acuminata* (red). (**E**) *M. balbisiana* (orange). (**F**) *M. schizocarpa* (blue). Different gene families are represented by different colored lines

### Identification of orthologous gene pairs of flavonoid biosynthetic genes in Musaceae

On average, 77 orthologous gene pairs were detected between the Musaceae species (Fig. S3 and Table 1). There is well characterized and extensive chromosomal recombination (translocations, inversions, and fusion/fission events) between the Musaceae species with chromosome basic number *x*=9 and 11 (Droc *et al*., 2022; Wang *et al*., 2022): the genes for the flavonoid biosynthetic pathway are within syntenic blocks, and their rearrangements across species follow the pattern of other genes [red lines linking flavonoid pathway genes; the synteny (collinearity) of 27,021 genes is shown by the grey underlying lines].

The *ANR* gene family has higher copy numbers (4 or 5) in *Musella lasiocarpa* and *Musa acuminata* than in other Musaceae species due to tandem triplication on chromosome 6 and chromosome 7 (Fig. 4, S1, and Table 1). Interestingly, this triplicated collinear block identified in *M. lasiocarpa* and *M. acuminata* was not found in other Musaceae species, suggesting that the tandem triplication event may have occurred independently in *M. lasiocarpa* and *M. acuminata* (Fig. 4B).

**Fig. 4.**
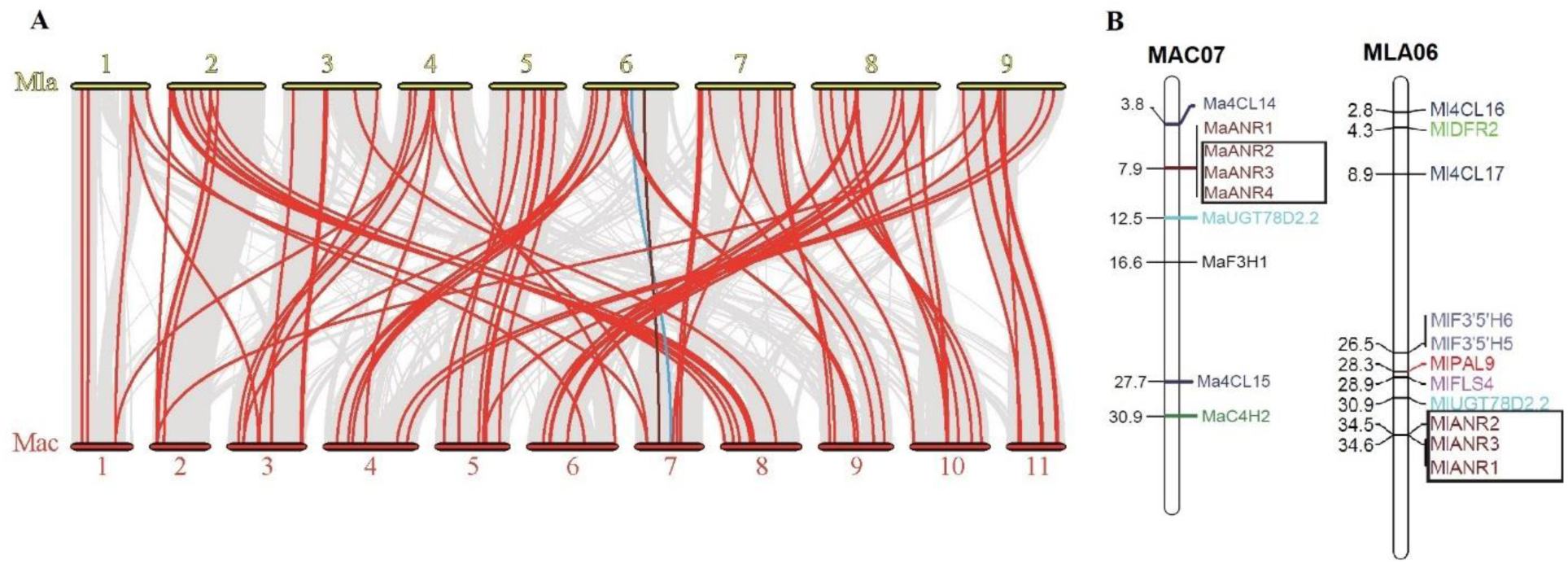
Synteny analysis and chromosomal location of *ANRs.* (**A**) Synteny analysis of flavonoid biosynthetic genes identified between MLA (*M. lasiocarpa*) and MAC (*M. acuminata*). The gray lines indicate all collinear gene blocks, while the red lines highlight the syntenic flavonoid biosynthetic gene pairs. The brown and blue lines indicate the ANR and UGT syntenic blocks, respectively. (**B**) Chromosome localization of flavonoid biosynthetic genes identified in *M. lasiocarpa* and *M. acuminata*. Genes from different families are shown in different colours. The chromosome number is shown at the top of each bar. The horizontal bars represent the gene locations on each chromosome (Mb). *ANR* genes with tandem duplications are framed by black boxes

For Musaceae, we calculated the Ka/Ks ratios and divergence times for orthologous genes (Table S3). The results showed that most (1166 out of 1169 comparisons) of the Ka/Ks ratios were less than 1, consistent with ongoing stabilizing or purifying selection with no positive selection in any of the Musaceae. The Ka/Ks ratios of three gene pairs in the UGT gene family were greater than 1 (*MaUGT75C1* vs. *MbUGT75C1*, *MaUGT79B1* vs. *MsUGT79B1*, and *MbUGT75C1* vs. *MsUGT75C1.4*), perhaps indicating slight (beneficial) positive selection pressure.

### Prediction of cis-acting elements in the putative gene promoters of flavonoid biosynthetic genes in Musaceae

Sequences (1.5 kb) upstream of the translation initiation codon for the flavonoid biosynthetic genes were examined for the presence of cis-acting elements using the PLANTCARE online database. In addition to cis-acting elements that are characteristic of eukaryotic promoters, various cis-acting elements, including those associated with phytohormones, light responses, plant growth, development, and stress responses (Alam *et al*., 2019), have been found among flavonoid biosynthetic genes. The cis-acting elements in the promoter region associated with stress responses (MYB and MYC elements), light responses (Box 4 and G-box), and phytohormones (ABRE, CGTCA-motif, and TGACG-motif) were present in almost all the flavonoid biosynthetic genes of Musaceae (Fig. S4). In addition to the *4CL* genes, all *CHS* gene promoters contain candidate ABREs and G-box elements, namely, the stress responsive elements (Xia *et al*., 2021).

### Expression of flavonoid biosynthetic genes under low temperature

The expression levels of flavonoid biosynthetic genes were analysed in *Musella lasiocarpa* var. *lasiocarpa* (yellow bracts) and *M. lasiocarpa* var. *rubribracteata* (red bracts) grown under low-temperature (15 ℃) and at 25 ℃. Among the 80 flavonoid biosynthetic genes, under both conditions, 18 were expressed at very low levels (transcripts per million TPM value < 1) in both the yellow- and red-bracted varieties and all *F3’5’H* genes and the transcription factors TTG and C1 had low TPMs (< 21), while 11 genes (*CHI*, *C4H*, *F3H*, *F3’H*, *DFR* and *LDOX* genes) were more strongly expressed (most with TPM value > 100) (Fig. 5 and Table S5). In other gene families, some members were less expressed than others in all four samples, with some being more strongly expressed in var. *lasiocarpa* (Y) than in var. *rubribraceata* (R) (eg. *PAL1*, *4CL13*, *FLS5*), or vice-versa (eg. *PAL2*, *PAL9*, *4CL10*, *C1.3*, and, most strongly, *TT8.2*). There were few notable trends in expression between temperatures, but after low-temperature treatment, the expression levels of *LAR* decreased by half while *MlDFR2* and *3* expression levels increased.

**Fig. 5.**
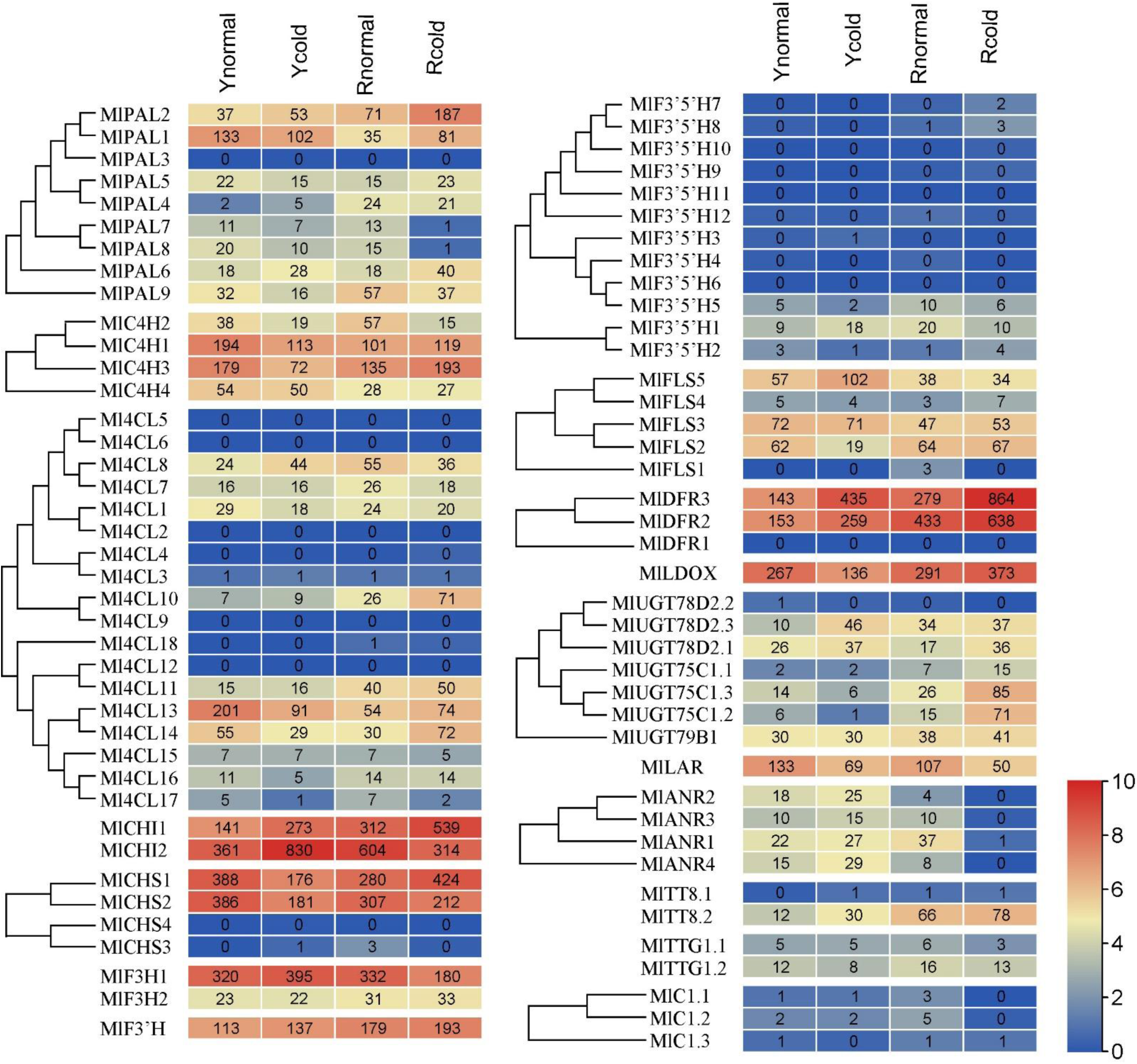
The expression of flavonoid biosynthetic genes in *Musella lasiocarpa* var. *lasiocarpa* (circles labelled Y) and *M. lasiocarpa* var. *rubribracteata* (labelled R); 15°C-treated groups labelled with ‘cold’, and control groups marked with ‘normal’. The flavonoid biosynthetic gene expression levels were calculated as transcripts per million (TPM) values. Different log_2_(TPM+1) values labelled with different colours, red to blue colours indicate expression levels from high to low

## Discussion

### Gene identification

Musaceae, with well-defined species in three genera with Asian and African natives and high-quality chromosome-level genome assemblies, is a good model taxon to understand the evolution of flavonoid gene diversity. All the structural gene families annotated in Poaceae and Arabidopsis are found in Musaceae (Table 1). Based on a syntenic homology analysis, 74, 80, 74, 80, 76, and 71 flavonoid biosynthetic genes were identified in *E. glaucum, Musella lasiocarpa, Musa beccarii, M. acuminata*, *M. balbisiana*, and *M. schizocarpa*, respectively, representing homologues of 34 out of the 40 flavonoid biosynthetic genes in Arabidopsis (Fig. 1, Table 1). The analysis of both structural and regulatory genes in the flavonoid pathway enables us to understand a variety of evolutionary and selective processes. The role of gene duplication, the nature of evolutionary rate variation among genes, the relative importance of structural and regulatory genes in the evolution of ecologically important characters (Rausher 2006), and the consequences of structural organization changes in the gene locations between species are important evolutionary questions. On average, 69 (65–74) structural genes encoding flavonoid biosynthetic enzymes and 7 (6–7) regulatory genes encoding transcription factors were identified in each species in the Musaceae. We identified an average of 29 (25–34) flavonoid biosynthetic genes with segmental duplications and 9 (6–13) flavonoid biosynthetic genes with tandem events (Table S4). Among these duplicated genes, regulatory genes were found for only two TT8 bHLH transcription factors (required for normal expression of some flavonoid biosynthetic genes) in *E. glaucum, M. beccarii, M. acuminata*, and *M. balbisiana*. These findings indicate that both tandem duplication and segmental duplication contribute to the amplification of structural gene number, while segmental duplication contributes to the amplification of key regulatory gene number. These results have implications for taxonomy and phylogeny (Hollingsworth *et al*., 1999), evolutionary mechanisms, and selection.

### Number of genes in a comparative context

The expression of flavonoid biosynthetic genes has expanded in Musaceae species compared with Arabidopsis, Brachypodium, and rice. In Arabidopsis, 32 structural genes that encode flavonoid biosynthetic enzymes and 2 regulatory genes encoding transcription factors have been identified as homologous to genes in Musaceae (Guo *et al*., 2014; Li *et al*., 2019). The greater number of flavonoid biosynthetic genes in Musaceae than in Arabidopsis presumably reflects the ancient genome triplication that occurred in the family (D’Hont *et al*., 2012; Zhang *et al*., 2020). Compared with those of Musaceae, Arabidopsis, sorghum, and ginger lack a reported functional LAR gene (Table 1); consequently, proanthocyanin synthesis only results in the production of epicatechin (2,3-cis-flavan-3-ol), which is found in the seed coat by ANR (encoded by the BANYULS gene) (Xie *et al*., 2003; Dixon *et al*., 2005; Li *et al*., 2019). Sequence and copy number evolution of F3’5’H is discussed below.

### Gene locations and clustering in the flavonoid pathway

There were three chromosomal clusters of tandem duplication/triplication events in every species (Figs. 2, S1 and Table S3). The genes exhibited a widespread distribution across chromosomes. Some closely linked genes encoding enzymes involved in different steps of the flavonoid pathway, such as FLS and UGT, 4CL and PAL, CHS, and C4H, were observed in the chromosome localization map (Fig. S1). Interestingly, segmental duplications were found in PAL, C4H, 4CL, CHS and UGT families in Musaceae, and there are tandem duplications in UGT families (Table S3). Gene amplifications, such as those involving enzymes at branch points upstream and downstream of metabolic pathways, are known to occur at bifurcation points (Dall’Olio *et al*., 2012; Wheeler *et al*., 2022). All the upstream genes underwent segmental duplications.

According to the phylogenetic trees (Fig. S2), flavonoid biosynthetic genes in Musaceae usually clustered together, which indicates that the duplications leading to their amplification occurred before the divergence of *E. glaucum*, *Musella lasiocarpa, Musa beccarii, M. acuminata*, *M. balbisiana*, and *M. schizocarpa*. Notably, the UGT genes of Arabidopsis, Poaceae, and ginger were clustered into UGT75C1, UGT78D2, and UGT79B1 groups (Fig. S2), implying that the duplications leading to the amplification of the UGT75C1, UGT78D2, and UGT79B1 genes were ancient events, pre-dating the separation of Musaceae from the outgroups.

### Sequence variation, phylogeny and adaptation

Analysis of nucleotide variation of flavonoid genes (Table S3) shows that most (1166 out of 1169 comparisons) of the non-synonymous to synonymous (Ka/Ks) ratios were less than 1, consistent with ongoing stabilizing or purifying selection with no positive selection at sequence level in Musaceae. Here, most of the diploid Musaceae accessions are non-domesticated wild species. There is no evidence that the species selected as crops (that is, *M. acuminata* as the desert banana, A genome, and *M. balbisiana*, B genome, often in hybrids with *M. acuminata*, for starchy fruits), show pre-adaptation before domestication, with no increase in flavonoid biosynthetic pathway genes, which could lead to increased stress tolerance. An increase in flavonoid pathway genes might reflect increased protection against biotic and abiotic stresses (Zhang *et al*., 2019), and colouration might be a quality attribute. Cenci *et al*. (2019) suggested that the presence of the B genome was associated with increased expression of genes related to flavonoid biosynthesis. The extreme plasticity of transcriptional regulation, leading to novel colouration phenotypes (Nair *et al*., 2005), has been shown by Recinos and Pucker (2023), so variation in genes and regulation is likely to be more important than gene copy number. Nevertheless, the 5- to 10-fold increase in the copy number of *F3’5’H* genes in Zingiberales compared with that in Poaceae suggested the selection of multiple copies in monocots, as proven by the greater number of 4CL gene copies than other genes in the pathway. In a genome-wide study, Rijzaani *et al*. (2022) proposed that a selective sweep through certain genomic regions might explain the genetic differentiation between *Ensete* and *Musa*; for example, some genes (GO terms) in highly differentiated regions are related to flavonoid biosynthesis and defense responses.

### Balance of structural and regulatory genes

Both structural and regulatory gene sequences are highly conserved among Musaceae species (Fig. 2 and S2). Changes in biosynthesis of flavonoids would be expected to alter the fitness of the plant (Guo *et al*., 2014). Transcription factors show conserved functions based on amino acids in the active center, and may be conserved because of involvement in several pathways, having pleiotropic effects (Dwivedi *et al*., 2016). Although some researchers reported that transcription factors evolved faster than their structural genes in the flavonoid pigment pathway (Rausher *et al*., 1999; Rausher 2006; Wheeler *et al*., 2022), our study suggest that the adaptive evolution of flavonoid gene families is mediated by both regulatory and structural genes.

The flavonoid gene copy number ratio of Musaceae/Poaceae is 1.16 for the early biosynthetic genes and 0.74 for the late biosynthetic genes, respectively (using data from Table 1; average early/late biosynthetic gene copy number in Musaceae divided by that of Poaceae, largely because of the high copy number of the *F3’5’H* gene). Upstream genes (phenylpropanoid pathway and early biosynthetic genes; Table 1) in the flavonoid pathway have evolved more slowly (Fig. S2) than the late biosynthetic genes, and these different evolutionary rates may be caused by upstream genes being more conserved due to involvement in multiple biochemical pathways (Fig. 1; Nair *et al*., 2005).

### Evolution of flavonoid 3’,4’,5’-hydroxylase *F3’5’H* genes

F3’5’H is a key branch-point in the flavonoid biosynthetic pathway (Fig. 1), generating 3’,4’,5’-hydroxylated flavonoids (Wang *et al*., 2014), which are regarded as giving quality attributes, e.g., for example to tea (Jin *et al*., 2017). F3’H and F3’5’H are members of the cytochrome P450 enzyme family (Park *et al*., 2021; Zhang *et al*., 2022). McClean *et al*. (2022) reported that their “phylogenetic analysis determined F3’5’H first appeared in the Streptophyta and is present in only 41% of angiosperm reference genomes” while F3’H is found universally in angiosperms, due to its critical role in abiotic stress, *F3’5’H* was reported as a major gene for the flower and seed coat colour regulation in plants (McClean *et al*., 2022), with MYB and MYC elements in the promoter region (McClean *et al*., 2022). Nevertheless, multiple copies of the F3’5’H family genes were found in Arabidopsis, Brachypodium, sorghum, ginger, *E. glaucum, Musella lasiocarpa, Musa beccarii, M. acuminata, M. balbisiana*, and *M. schizocarpa* (Fig. S2), with amplification in monocots before separation of Poales and Zingiberales. Further proliferation of F3’5’Hs occurred at one of the loci after the separation of Musaceae and gingers; one clade of *F3’5’H* genes was specific to Musaceae (Fig. 2). The ratio of gene number of *F3’5’H* (average Musaceae/average Poaceae) was 7.33, while the ratio of gene number of *F3’H* is 0.31. The centromeric locations of duplicated *MaF3’5’H*, *MbF3’5’H*, and *MsF3’5’H* on chromosome 2 suggest tandem duplication by unequal crossing over (Fig. 2): clusters of *F3’5’H* are near the chromosome 2 centromere of *M. acuminata, M. balbisiana* and *M. schizocarpa*. The non-centromeric *F3’5’H* genes in Musaceae are close to those of species in other families, while the centromeric *F3’5’H* genes in Musaceae are relatively distant from the others in the phylogeny (Fig. 2), perhaps suggesting different origins of the centromeric *F3’5’H* genes. Seitz *et al*. (2006 and 2015) reported multiple origins of *F3’5’H* from F3’H precursors at least four times in dicotyledonous plants, while Negi *et al*. (2023) reported that the functional divergence of the 5’ regulator region of *F3’5’H* related to stress response flavonoid phenotype of the Musaceae is underexplored, particularly with respect to non-anthocyanin flavonoids, and very little can be said about any relationship between *F3’5’H* copy number and 3’5’-O flavonoid diversity. Available data show that B-ring 3’5’-oxygenation (as myricetin or delphinidin and its derivatives) occurs in two species of *Ensete* and nine species of *Musa*, most of which have not been investigated (Gornall unpublished).

### Gene expression in *Musella lasiocarpa*

Flavonoid biosynthetic genes were expressed at different levels in the bract tissues of *Musella lasiocarpa*. Similar to the gene expression levels in *Musa acuminata* bracts (Pandey *et al*., 2016), *MlF3H1*, *MlF3’H*, *MlCHI1/2*, *MlFLS2/3/5* and *MlLDOX* genes were expressed at high levels in the bracts of *M. lasiocarpa* (Fig. 5). In particular, the *LDOX* (one copy only) was expressed at a high level (TPM values of 135.6-373.1) (Fig. 5): the spatial correlation between *LDOX* gene expression and anthocyanin metabolites in banana (cv. Grand Naine) bract tissue supports the importance of LDOX enzyme in the conversion of leucoanthocyanidins to anthocyanidins (Busche *et al*., 2021). Because of their powerful antioxidant activity, accumulation of flavonoids is likely to counteract oxidative damage induced by various stresses (Reginata *et al*., 2014). The accumulation of anthocyanins gives the bracts of Musaceae yellow or red colours (Pazmiño-Durán *et al*., 2001), with relatively high expression of *MlF3H1* and *MlF3’H* (TPM > 110, Fig. 5) regulating synthesis of orange-yellow pelargonidin and the purplish-red cyanidin (Fig. 1) pigment. The colour of the bracts of plantain species, related to the composition of anthocyanins, has taxonomic value: the bracts of *Musa coccinea* contain both delphinidin and pelargonidin, and the bracts of *Musa balbisiana* contain both delphinidin and cyanidin (Pazmiño-Durán *et al*., 2001; Chandra *et al*., 2021). In terms of other flavonoid compounds, Pandey *et al*. (2016) reported that the content of isoquercetin in banana bracts was greater than that in tissues of roots, leaves, pseudostems, and pericarp. After low-temperature treatment, the expression level of *MlDFR2* and *3* increased, presumably because the content of leucoanthocyanidin increased in the bracts of *M. lasiocarpa* (Fig. 5; see Fig. 1), while *MlLAR* expression level decreased under cold conditions. It is possible that most leucoanthocyanidins were synthesized from the highly expressed *MlLDOX* and *MlUGT* (*MlUFGT*) genes to synthesize anthocyanidins rather than *MlLAR* to synthesize proanthocyanidins (Figs. 1 and 5; Wang *et al*., 2013; León-Chan *et al*., 2020). The expression levels of the *C1* genes from the MYB family, a positive regulator of the flavonoid biosynthetic pathway, was always decreased or unchanged in the yellow and red bract tissues of *M. lasiocarpa* (Zhang *et al*., 2015). The cold domestication process of vesicular mosses (*Physcomitrella patens*) is dependent on the phytohormone abscisic acid (ABA) for signalling (Bhyan *et al*., 2012). Negi et al. (2023) concluded that *F3’5’H* is a stress-responsive gene that is induced in banana by the application of salicylic acid and methyl jasmonate; whereas under drought, salt and ABA treatments, the *F3’5’H* expression level showed a sharp decrease.

### Flavonoid pathway in evolution and crop improvement

The first pangenome of banana was described by Rijzaani *et al*. (2022), and the Banana Genome Hub (Droc *et al*., 2022) provides access to curated data and high-quality chromosome-level genome assemblies. Here, flavonoid pathway analysis revealed the value and mechanism through which such pangenome data can be used. Knowledge of the range of diversity of genes in the flavonoid pathway has implications for taxonomy and phylogeny, potentially identifying traits selected from the wild for domesticated cultivars and suggesting where intraspecific variation may be valuable. Musaceae has three well-defined genera (genus *Musa* in two taxonomic sections);: therefore, this family is a concise model for understanding the range and nature of flavonoid gene diversity. In addition to the extensive use of ensete as human food and animal feed, some ensete landraces play a well-known and important role in traditional medicine to cure diverse ailments (Nuraga *et al*., 2022), while ginger rhizomes and extracts are widely considered to have ethno-medicinal and nutritional value (Dissanayake *et al*., 2020). Flavonoids possess a number of medicinal benefits, and their effectiveness has been proven for a variety of disease treatments (Ullah *et al*., 2020). Although the mechanisms of action of flavonoids are not fully understood (Panche *et al*., 2016), access to gene sequence and copy number information can assist in their enhancement in crops. As such, knowledge of the diverse flavonoid genes is valuable for functional and phylogenetic studies across all angiosperm species and can be used for the development of approaches to modify flavonoid or anthocyanin contents related to stress resistance and nutritional or medical applications.

## Abbreviations

4CL: 4 coumarate
CoA: ligase
ANR: anthocyanidin reductase
ANS/LDOX: anthocyanidin synthase/leucoanthocyanidin dioxygenase
C4H: cinnamic acid 4-hydroxylase
CHI: chalcone isomerase
CHS: chalcone synthase
DFR: dihydroflavonol reductase
EBG: early biosynthetic gene
F3’5’H: flavanone 3’5’-hydroxylase
F3’H: flavanone 3’-hydroxylase
F3H: flavanone 3-hydroxylase
FBG: flavonoid biosynthetic gene
FLS: flavonol synthase
FNS: flavone synthase
GPP: general phenylpropanoid pathway
IFS: isoflavone synthase
Ka: non-synonymous
Ks: synonymous
LAR: leucoanthocyanidin reductase
LBG: late biosynthetic gene
NJ: neighbor joining
PAL: phenylalanine ammonia lyase
SMART: simple modular architecture research tool
TF: transcription factor
TMM: trimmed mean of M-values
TPM: transcripts per million
UGP/UFGT: UDP-flavonoid glucosyl transferase

## Ethics approval and consent to participate

Not applicable

## Consent for publication

Not applicable

## Availability of data and materials

The genome data of *O. sativa*, *S. bicolor*, *Z. officinale*, *E. glaucum*, *Musella lasiocarpa*, *Musa beccarii*, *M. acuminata*, *M. balbisiana*, and *M. schizocarpa* were retrieved from https://banana-genome-hub.southgreen.fr/content/download. The *B. distachyon* (PRJNA32607 at the China National Center for Bioinformation) and *A. longiglumis* (PRJNA956334 at the National Center for Biotechnology Information) genome data and flavonoid biosynthetic pathway genes of *A. thaliana*, *O. sativa* and *B. distachyon* were retrieved from NCBI at https://www.ncbi.nlm.nih.gov/. The RNA-seq data of *Musella* bracts can be obtained in the Genome Sequence Archive for project PRJCA023025 (https://ngdc.cncb.ac.cn/gsa/browse/CRA014572).

## Authors’ Contributions

Conceptualization, DC, PHH, and RJG; methodology, DC and PHH; software, DC; validation, DC, GX, LY, RJG, PHH, and QL; formal analysis, DC; resources, ZW; original draft preparation, DC; review and editing, DC, GX, LY, RJG, PHH, and QL; visualization, DC; supervision, PHH, QL, and RJG; project administration, PHH, and QL; funding acquisition, PHH and QL All authors have read and agreed to the published version of the manuscript.

## Supporting Information

### Additional file 1

**Figure S1.** Chromosome localization of flavonoid biosynthetic genes identified in Musaceae.

**Figure S2.** Phylogenetic trees (Neighbour-Joining method) showing the genetic relationship of flavonoid biosynthetic genes in Arabidopsis, *B. distachyon*, *A. longiglumis*, *O. sativa*, *S. bicolor*, *Z. officinale*, *E. glaucum*, *Musella lasiocarpa*, *Musa beccarii*, *M. acuminata*, *M. balbisiana*, and *M. schizocarpa*.

**Figure S3.** Synteny analysis of FBGs between six Musaceae species.

**Figure S4.** Cis-acting elements in flavonoid biosynthetic genes of Musaceae.

### Additional file 2

**Table S1.** Flavonoid biosynthetic genes (FBGs) in *A. thaliana*, *B. distachyon*, *O. sativa*, *S. bicolor*, *Z. officinale*, *E. glaucum*, *Musella lasiocarpa, Musa beccarii, M. acuminata*, *M. balbisiana* and *M. schizocharpa*.

**Table S2.** Information on flavonoid biosynthetic genes identified in Musaceae.

**Table S3.** Divergence time, Ka, Ks, and Ka/Ks values of orthologous flavonoid biosynthetic genes pairs between Musaceae.

**Table S4.** Intra-species pairs of duplicated genes.

### Additional file 3

**Table S5.** Expression of several flavonoid biosynthetic genes in *Musella lasiocarpa*.

## References

Alam I, Cui DL, Batool K, Yang YQ, Lu YH. Comprehensive genomic survey, characterization and expression analysis of the HECT gene family in *Brassica rapa* L. and *Brassica oleracea* L. Genes. 2019;10:400. 10.3390/genes10050400.

Alseekh S, Perez de Souza L, Benina M, Fernie AR. The style and substance of plant flavonoid decoration, towards defining both structure and function. Phytochemistry. 2020;174:112347. 10.1016/j.phytochem.2020.112347.

Amah D, Turner D, Gibbs DJ, Waniale A., Gram G, Swennen R. Overcoming the fertility crisis in bananas (*Musa* spp.). In: Achieving sustainable cultivation of bananas. Burleigh Dodds Science Publishing; 2020;p. 257–306.

Bartas M, Volna A, Cerven J, Pucker B. Identification of annotation artifacts concerning the *chalcone synthase* (*CHS*). BMC Research Notes. 2018;16:109. 10.1186/s13104-023-06386-z.

Bate-Smith EC. Flavonoid compounds in foods. Advances in Food Research. 1954;55:261–300.

Bhyan SB, Minami A, Kaneko Y, Suzuki S, Arakawa K, Sakata Y, Takezawa D. Cold acclimation in the moss *Physcomitrella patens* involves abscisic acid-dependent signaling. Journal of Plant Physiology. 2012;169:137–45. 10.1016/j.jplph.2011.08.004.

Borrell JS, Biswas MK, Goodwin M, Blomme G, Schwarzacher T, Heslop-Harrison JS, Wendawek AM, Berhanu A, Kallow S, Janssens S, et al. Enset in Ethiopia: a poorly characterized but resilient starch staple. Annals of Botany. 2019;123:747–66. 10.1093/aob/mcy214.

Busche M, Acatay C, Martens S, Weisshaar B, Stracke R. Functional characterisation of banana (*Musa* spp.) 2-oxoglutarate-dependent dioxygenases involved in flavonoid biosynthesis. Frontiers in Plant Science. 2021;12:701780. 10.3389/fpls.2021.701780.

Busche M, Pucker B, Weisshaar B, Stracke R. Three R2R3-MYB transcription factors from banana (*Musa acuminata*) activate structural anthocyanin biosynthesis genes as part of an MBW complex. BMC Research Notes. 2023;16:103. 10.1186/s13104-023-06375-2.

Cenci A, Hueber Y, Zorrilla-Fontanesi Y, van Wesemael J, Kissel E, Gislard M, Sardos J, Swennen R, Roux N, Carpentier SC, et al. Effect of paleopolyploidy and allopolyploidy on gene expression in banana. BMC Genomics. 2019;20:244. 10.1186/s12864-019-5618-0.

Chandra RD, Prihastyanti MNU, Lukitasari DM. Effects of pH, high pressure processing, and ultraviolet light on carotenoids, chlorophylls, and anthocyanins of fresh fruit and vegetable juices. EFood. 2021;2:113–24. 10.2991/efood.k.210630.001.

Cheesman EE. Classification of the bananas: the genus *Ensete* Horan. Kew Bulletin. 1947;2:97–106. 10.2307/4109206.

Chen, S, Zhou, Y, Chen, Y, Gu j. Fastp: an ultra-fast all-in-one FASTQ preprocessor. Bioinformatics. 2018;34:884–90. 10.1093/bioinformatics/bty560.

Christelová P, Valárik M, Hřibová E, De Langhe E, Doležel J. A multi gene sequence-based phylogeny of the Musaceae (banana) family. BMC Evolutionary Biology. 2011;11:103. 10.1186/1471-2148-11-103.

Chu S, Wang J, Zhu Y, Liu S, Zhou X, Zhang H, Wang C, Yang W, Tian Z, Cheng H, et al. An R2R3-type MYB transcription factor, GmMYB29, regulates isoflavone biosynthesis in soybean. PLoS Genetics. 2017;13:e1006770. 10.1371/journal.pgen.1006770.

D’Hont A, Denoeud F, Aury JM, Baurens FC, Carreel F, Garsmeur O, Noel B, Bocs S, Droc G, Rouard M, et al. The banana (*Musa acuminata*) genome and the evolution of monocotyledonous plants. Nature. 2012;488:213–7. 10.1038/nature11241.

Dall’Olio GM, Laayouni H, Luisi P, Sikora M, Montanucci L, Bertranpetit J. Distribution of events of positive selection and population differentiation in a metabolic pathway: the case of asparagine N-glycosylation. BMC Evolutionary Biology. 2012;12:98. 10.1186/1471-2148-12-98.

Davies KM, Jibran R, Zhou Y, Albert NW, Brummell DA, Jordan BR, Bowman JL, Schwinn KE. The evolution of flavonoid biosynthesis: a bryophyte perspective. Frontiers in Plant Science. 2020;11:7. 10.3389/fpls.2020.00007

Dissanayake KGC, Waliwita W, Liyanage RP. A review on medicinal uses of *Zingiber officinale* (ginger). International Journal of Health Sciences and Research. 2020;10:142–8.

Dixon RA, Xie DY, Sharma SB. Proanthocyanidins – a final frontier in flavonoid research? New Phytologist. 2005;165:9–28. 10.1111/j.1469-8137.2004.01217.

Dobin A, Gingeras TR. Mapping RNA-seq reads with STAR. Current Protocols in Bioinformatics. 2015;51:11–4. 10.1002/0471250953.bi1114s51.

Droc G, Martin G, Guignon V, Summo M, SempéréG, Durant E, Soriano A, Baurens FC, Cenci A, Breton C, et al. The banana genome hub: a community database for genomics in the Musaceae. Horticulture Research. 2022;9:uhac221. 10.1093/hr/uhac221.

Dwivedi SL, Upadhyaya HD, Chung IM, De Vita P, García-Lara S, Guajardo-Flores D, Gutiérrez-Uribe JA, Serna-Saldívar SO, Rajakumar G, Sahrawat KL, et al. Exploiting phenylpropanoid derivatives to enhance the nutraceutical values of cereals and legumes. Frontiers in Plant Science. 2016;7:763. 10.3389/fpls.2016.00763.

Falcone Ferreyra ML, Rius S, Casati P. Flavonoids: biosynthesis, biological functions, and biotechnological applications. Frontiers in Plant Science. 2012;3:222. 10.3389/fpls.2012.00222.

Ferreyra MLF, Serra P, Casati P. Recent advances on the roles of flavonoids as plant protective molecules after UV and high light exposure. Physiologia Plantarum. 2021;173:736–49. 10.1111/ppl.13543.

Finn RD, Clements J, Eddy SR. HMMER web server: interactive sequence similarity searching. Nucleic Acids Research. 2011;39:W29–37. 10.1093/nar/gkr367.

Giusti MM, Miyagusuku–Cruzado G, Wallace TC. Flavonoids as natural pigments. Handbook of Natural Colorants. 2023:371–90. 10.1002/9781119811749.ch17.

Goiris K, Muylaert K, Voorspoels S, Noten B, De Paepe D, Baart E GJ, De Cooman L. Detection of flavonoids in microalgae from different evolutionary lineages. Journal of Phycology. 2014;50:483–92.

Gornall RJ, Bohm BA, Dahlgren R. The distribution of flavonoids in the angiosperms. Botaniska Notiser. 1979;132:1–30.

Guo N, Cheng F, Wu J, Liu B, Zheng S, Liang J, Wang X. Anthocyanin biosynthetic genes in *Brassica rapa*. BMC Genomics. 2014;15:426. 10.1186/1471-2164-15-426.

Häkkinen M. Reappraisal of sectional taxonomy in *Musa* (Musaceae). Taxon. 2013;62:809–13. 10.12705/624.3.

Heslop-Harrison JS, Schwarzacher T. Domestication, genomics and the future for banana. Annals of Botany. 2007;100:1073–84. 10.1093/aob/mcm191.

Hollingsworth PM, Bateman RM, Gornall RJ. Molecular systematics and plant evolution. CRC Press, UK; 1999.

Janssens SB, Vandelook F, De Langhe E, Verstraete B, Smets E, Vandenhouwe I, Swennen R. Evolutionary dynamics and biogeography of Musaceae reveal a correlation between the diversification of the banana family and the geological and climatic history of Southeast Asia. New Phytologist. 2016;210:1453–65. 10.1111/nph.13856.

Jayachandran M, Zhang T, Wu Z, Liu Y, Xu B. Isoquercetin regulates SREBP-1C via AMPK pathway in skeletal muscle to exert antihyperlipidemic and anti-inflammatory effects in STZ induced diabetic rats. Molecular Biology Reports. 2020;47:593– 602. 10.1007/s11033-019-05166-y.

Jin JQ, Ma JQ, Yao MZ, Ma CL, Chen L. Functional natural allelic variants of *flavonoid 3’,5’-hydroxylase* gene governing catechin traits in tea plant and its relatives. Planta. 2017;245:523–38. 10.1007/s00425-016-2620-5.

Krzywinski M, Schein J, Birol I, Connors J, Gascoyne R, Horsman D, Jones SJ, Marra MA. Circos: an information aesthetic for comparative genomics. Genome Research. 2009;19:1639–45. 10.1101/gr.092759.109.

Kumar S, Stecher G, Li M, Knyaz C, Tamura K. MEGA X: molecular evolutionary genetics analysis across computing platforms. Molecular Biology and Evolution. 2018;35:1547–49. 10.1093/molbev/msy096.

León-Chan RG, Lightbourn-Rojas LA, López-Meyer M, Amarillas L, Heredia JB, Martínez-Bastidas T, Villicaña C, León-Félix J. Differential gene expression of anthocyanin biosynthetic genes under low temperature and ultraviolet-B radiation in bell pepper (*Capsicum annuum*). International Journal of Agriculture and Biology. 2020;23:501–8. 10.17957/IJAB/15.1315.

Lepiniec L, Debeaujon I, Routaboul JM, Baudry A, Pourcel L, Nesi N, Caboche M. Genetics and biochemistry of seed flavonoids. Annual Review of Plant Biology. 2006;57:405–30. 10.1146/annurev.arplant.57.032905.105252.

Lescot M, Déhais P, Thijs G, Marchal K, Moreau Y, Van de Peer Y, RouzéP, Rombauts S. PlantCARE, a database of plant cis-acting regulatory elements and a portal to tools for *in silico* analysis of promoter sequences. Nucleic Acids Research. 2002;30:325–7 10.1093/nar/30.1.325.

Lescot M, Piffanelli P, Ciampi AY, Ruiz M, Blanc G, Leebens-Mack J, da Silva FR, Santos CMR, D’Hont A, Garsmeur O, et al. Insights into the *Musa* genome: syntenic relationships to rice and between Musa species. BMC Genomics. 2008;9:58. 10.1186/1471-2164-9-58.

Li H, Tian J, Yao Y, Zhang J, Song T, Li K, Yao Y. Identification of leucoanthocyanidin reductase and anthocyanidin reductase genes involved in proanthocyanidin biosynthesis in *Malus* crabapple plants. Plant Physiology and Biochemistry. 2019;139:141–51. 10.1016/j.plaphy.2019.03.003.

Li HW. The Musaceae of Yunnan. Acta Phytotaxonomica Sinica. 1978;16:54–64.

Li HW. Musaceae. In: Wu CY, editor, Flora of Yunnan, Vol. 2. Science Press, 1979. p.725–33.

Liao Y, Smyth GK, Shi W. FeatureCounts: an efficient general purpose program for assigning sequence reads to genomic features. Bioinformatics. 2014;30:923–30. 10.1093/bioinformatics/btt656.

Liu F, Dou T, Hu C, Zhong Q, Sheng O, Yang Q, Deng G, He W, Gao H, Li C, Dong T. WRKY transcription factor MaWRKY49 positively regulates pectate lyase genes during fruit ripening of *Musa acuminata*. Plant Physiology and Biochemistry. 2023;194:643–50.

Liu Q, Yuan HY, Li MZ, Wang ZW, Cui DL, Ye YS, Sun ZY, Tan XK, Schwarzacher T, Heslop-Harrison JS. Chromosome-scale genome assembly of the diploid oat *Avena longiglumis* reveals the landscape of repetitive sequences, genes and chromosome evolution in grasses. bioRxiv. 2022:479819.

Liu S, Ju J, Xia G. Identification of the *flavonoid 3’-hydroxylase* and *flavonoid 3’*, *5’-hydroxylase* genes from Antarctic moss and their regulation during abiotic stress. Gene. 2014;543:145–52. 10.1016/j.gene.2014.03.026.

Mahajan M, Ahuja PS, Yadav SK. Post-transcriptional silencing of flavonol synthase mRNA in tobacco leads to fruits with arrested seed set. PLoS One. 2011;6:e28315. 10.1371/journal.pone.0028315.

Manzoor MA, Sabir IA, Shah IH, Riaz MW, Rehman S, Song C, Gouhui L, Malik MS, Ashraf GA, Haider MS. Flavonoids: a review on biosynthesis and transportation mechanism in plants. Functional and Integrative Genomics. 2023;23:212

McClean PE, Lee R, Howe K, Osborne C, Grimwood J, Levy S, Haugrud AP, Plott C, Robinson M, Skiba RM, et al. The common bean V gene encodes flavonoid 3’5’hydroxylase: a major mutational target for flavonoid diversity in angiosperms. Frontiers in Plant Science. 2022;13:869582. 10.3389/fpls.2022.869582.

Mingmanit Y, Boonsrangsom T, Sujipuli K, Ratanasut K, Inthima P. Pollen viabilities and gene expression profiles across *Musa* genomes. AoB Plants. 2023;15:plad052.

Mistry J, Chuguransky S, Williams L, Qureshi M, Salazar Gustavo A, Sonnhammer ELL, Tosatto SCE, Paladin L, Raj S, Richardson LJ, et al. Pfam: the protein families database in 2021. Nucleic Acids Research. 2021;49: D412–9. 10.1093/nar/gkaa913.

Nair AS, Teo CH, Schwarzacher T, Harrison PH. Genome classification of banana cultivars from South India using IRAP markers. Euphytica. 2005;165:285–90. 10.1007/s10681-005-7321-2.

Negi S, Tak H, Madari S, Bhakta S, Ganapathi TR. Functional characterization of 5’-regulatory region of flavonoid 3’,5’- hydroxylase-1 gene of banana plants. Protoplasma. 2023;260:391–403. 10.1007/s00709-022-01785-8.

Neumann K, Hildebrand E. Early bananas in Africa: the state of the art. Ethnobotany Research and Applications. 2009;7:353–62. Retrieved from https://ethnobotanyjournal.org/era/index.php/era/article/view/361.

Ng J, Smith SD. How to make a red flower: the combinatorial effect of pigments. AoB Plants. 2016;8:plw013. 10.1093/aobpla/plw013.

Nugraha A, Darsono D, Marwanti S. Analysis of indonesian banana export performance in major export destination countries. SVU-International Journal of Agricultural Sciences. 2023;5:127–36. 10.21608/SVUIJAS.2023.219439.1294.

Nuraga GW, Feyissa T, Tesfaye K, Biswas MK, Schwarzacher T, Borrell JS, Wilkin P, Demissew S, Tadele Z, Heslop-Harrison JS. The genetic diversity of enset (*Ensete ventricosum*) landraces used in traditional medicine is similar to the diversity found in non-medicinal landraces. Frontiers in Plant Science. 2022;12:756182. 10.3389/fpls.2021.756182.

Panche AN, Diwan AD, Chandra SR. Flavonoids: an overview. Journal of Nutritional Science. 2016;5:e47. 10.1017/jns.2016.41.

Pandey A, Alok A, Lakhwani D, Singh J, Asif MH, Trivedi PK. Genome-wide expression analysis and metabolite profiling elucidate transcriptional regulation of flavonoid biosynthesis and modulation under abiotic stresses in banana. Scientific Reports. 2016;6:31361. 10.1038/srep31361.

Park S, Lee H, Min MK, Ha J, Song J, Lim CJ, Oh J, Lee SB, Lee JY, Kim BG. Functional characterization of BrF3’H, which determines the typical flavonoid profile of purple Chinese cabbage. Frontiers in Plant Science. 2021;12: 793589. 10.3389/fpls.2021.793589.

Passos MAN, de Oliveira Cruz V, Emediato FL, de Camargo Teixeira C, Souza MT, Matsumoto T, Rennó Azevedo VC, Ferreira CF, Amorim EP, de Alencar Figueiredo LF, et al., Development of expressed sequence tag and expressed sequence tag-simple sequence repeat marker resources for *Musa acuminata*. AoB Plants. 2012;2012:pls030. 10.1093/aobpla/pls030.

Paysan-Lafosse T, Blum M, Chuguransky S, Grego T, Pinto BL, Salazar Gustavo A, Bileschi Maxwell L, Bork P, Bridge A, Colwell L, et al., InterPro in 2022. Nucleic Acids Research. 2023;51:D418–27. 10.1093/nar/gkac993.

Pazmiño-Durán EA, Giusti MM, Wrolstad RE, Glória MBA. Anthocyanins from banana bracts (*Musa* ×*paradisiaca*) as potential food colorants. Food Chemistry. 2001;73:327–32. 10.1016/S0308-8146(00)00305-8.

Peng Y, Thrimawithana AH, Cooney JM, Jensen DJ, Espley RV, Allan AC. The proanthocyanin-related transcription factors MYBC1 and WRKY44 regulate branch points in the kiwifruit anthocyanin pathway. Scientific Reports. 2020;10:14161.

Petroni K, Tonelli C. Recent advances on the regulation of anthocyanin synthesis in reproductive organs. Plant Science. 2011;181:219–29. 10.1016/j.plantsci.2011.05.009.

Pucker B, Reiher F, Schilbert HM. Automatic identification of players in the flavonoid biosynthesis with application on the biomedicinal plant *Croton tiglium*. Plants. 2020;9:1103. 10.3390/plants9091103.

Pucker B. Automatic identification and annotation of MYB gene family members in plants. BMC Genomics. 2022;23:220. 10.1186/s12864-022-08452-5.

Qian Y, Cheng Y, Cheng X, Jiang H, Zhu S, Cheng B. Identification and characterization of Dicer-like, Argonaute and RNA-dependent RNA polymerase gene families in maize. Plant Cell Reports. 2011;30:1347–63. 10.1007/s00299-011-1046-6.

Ranjha MMAN, Irfan S, Nadeem M, Mahmood S. A comprehensive review on nutritional value, medicinal uses, and processing of banana. Food Reviews International. 2022;38:199–225. 10.1080/87559129.2020.1725890.

Rausher MD, Miller RE, Tiffin P. Patterns of evolutionary rate variation among genes of the anthocyanin biosynthetic pathway. Molecular Biology and Evolution. 1999;16:266–74. 10.1093/oxfordjournals.molbev.a026108.

Rausher MD. The evolution of flavonoids and their genes. In Grotewold E, editon, The science of flavonoids, Springer New York: New York, NY; 2006. p. 175–211.

Recinos MFM, Pucker B. Genetic factors explaining anthocyanin pigmentation differences. bioRxiv. 2023;543820. 10.1101/2023.06.05.543820.

Reginato MA, Castagna A, Furlán A, Castro S, Ranieri A, Luna V. Physiological responses of a halophytic shrub to salt stress by Na2SO4 and NaCl: oxidative damage and the role of polyphenols in antioxidant protection. AoB Plants. 2014;6:plu042.

Rijzaani H, Bayer PE, Rouard M, Doležel J, Batley J, Edwards D. The pangenome of banana highlights differences between genera and genomes. Plant Genome. 2022;15:e20100. 10.1002/tpg2.20100.

Robinson MD, McCarthy DJ, Smyth GK. edgeR: a bioconductor package for differential expression analysis of digital gene expression data. Bioinformatics. 2010;26:139–40. 10.1093/bioinformatics/btp616.

Saito K, Yonekura-Sakakibara K, Nakabayashi R, Higashi Y, Yamazaki M, Tohge T, Fernie AR. The flavonoid biosynthetic pathway in Arabidopsis: structural and genetic diversity. Plant Physiology and Biochemistry. 2013;72:21–34. 10.1016/j.plaphy.2013.02.001.

Samanta A, Das G, Das SK. Roles of flavonoids in plants. Carbon. 2011;100:12–35.

Seitz C, Eder C, Deiml B, Kellner S, Martens S, Forkmann G. Cloning, functional identification and sequence analysis of flavonoid 3′-hydroxylase and flavonoid 3’, 5’-hydroxylase cDNAs reveals independent evolution of flavonoid 3’, 5’-hydroxylase in the Asteraceae family. Plant Molecular Biology. 2006;61:365–81. 10.1007/s11103-006-0012-0.

Seitz C, Ameres S, Schlangen K, Forkmann G, Halbwirth H.: Multiple evolution of flavonoid 3’, 5’-hydroxylase. Planta. 2015;245:561–73. 10.1007/s00425-015-2293-5.

Shirley BW. Flavonoid biosynthesis:‘new’functions for an ‘old’pathway. Trends in Plant Science. 1996;1:377–82. 10.1016/S1360-1385(96)80312-8.

Someya S, Yoshiki Y, Okubo K. Antioxidant compounds from bananas (*Musa Cavendish*). Food Chemistry. 2002;79:351–4. 10.1016/S0308-8146(02)00186-3.

Tiku AR. Antimicrobial compounds (phytoanticipins and phytoalexins) and their role in plant defense. In: Mérillon JM, Ramawat KG, editors. Co-Evolution secondary metabolites. Springer Nature, Switzerland; 2020. p. 845–68.

Ullah A, Munir S, Badshah SL, Forkmann G, Halbwirth H. Multiple evolution of flavonoid 3’, 5’-hydroxylase. Molecules. 2020;25:5243. 10.1007/s00425-015-2293-5.

van der Kooi CJ, Dyer AG, Kevan PG, Lunau K. Functional significance of the optical properties of flowers for visual signalling. Annals of Botany. 2019;123:263–76. 10.1093/aob/mcy119.

Voorrips RE. MapChart: software for the graphical presentation of linkage maps and QTLs. Journal of Heredity. 2022;93:77–8. 10.1093/jhered/93.1.77

Wang D, Zhang Y, Zhang Z, Zhu J, Yu J. KaKs_Calculator 2.0: a toolkit incorporating gamma-series methods and sliding window strategies. Genomics, Proteomics and Bioinformatics. 2010;8:77–80. 10.1016/S1672-0229(10)60008-3.

Wang H, Fan W, Li H, Yang J, Huang J, Zhang P. Functional characterization of dihydroflavonol-4-reductase in anthocyanin biosynthesis of purple sweet potato underlies the direct evidence of anthocyanin function against abiotic stresses. PLoS One. 2013;8:1‒14. 10.1371/journal.pone.0078484.

Wang L, Ying Lam LP, Lui ACW, Zhu FY, Chen MX, Liu H, Zhang J, Lo C. Flavonoids are indispensable for complete male fertility in rice. Journal of Experimental Botany. 2020;71:4715–28. 10.1093/jxb/eraa204.

Wang YS, Xu YJ, Gao LP, Yu O, Wang XZ, He XJ, Jiang XL, Liu YJ, Xia T. Functional analysis of flavonoid 3’, 5’-hydroxylase from tea plant (*Camellia sinensis*): critical role in the accumulation of catechins. BMC Plant Biology. 2014;14:1–14. 10.1186/s12870-014-0347-7.

Wang Z, Miao H, Liu J, Xu B, Yao X, Xu C, Zhao S, Fang X, Jia C, Wang J. *Musa balbisiana* genome reveals subgenome evolution and functional divergence. Nature Plants. 2019;5:810–21 10.1038/s41477-019-0452-6.

Wang Z, Rouard M, Biswas MK, Droc G, Cui D, Roux N, Baurens F.C, Ge X.J, Schwarzacher T, Heslop-Harrison P, et al. A chromosome-level reference genome of *Ensete glaucum* gives insight into diversity and chromosomal and repetitive sequence evolution in the Musaceae. GigaScience. 2022;11:giac027. 10.1093/gigascience/giac027.

Wang ZF, Rouard M, Droc G, Heslop-Harrison P, Ge XJ. Genome assembly of *Musa beccarii* shows extensive chromosomal rearrangements and genome expansion during evolution of Musaceae genomes. GigaScience. 2023;12:giad005. 10.1093/gigascience/giad005.

Wei L, Wang W, Li T, Chen O, Yao S, Deng L, Zeng K. Genome-wide identification of the *CsPAL* gene family and functional analysis for strengthening green mold resistance in citrus fruit. Postharvest Biology and Technology. 2023;196:112178. 10.1016/j.postharvbio.2022.112178.

Wen W, Alseekh S, Fernie AR. Conservation and diversification of flavonoid metabolism in the plant kingdom. Current Opinion in Plant Biology. 2020;55:100–8. 10.1016/j.pbi.2020.04.004.

Wheeler LC, Walker JF, Ng J, Deanna R, Dunbar-Wallis A, Backes A, Pezzi PH, Palchetti MV, Robertson HM, Monaghan A, et al., Transcription factors evolve faster than their structural gene targets in the flavonoid pigment pathway. Molecular Biology and Evolution. 2022;39:msac044. 10.1093/molbev/msac044.

Wilkins O, Nahal H, Foong J, Provart NJ, Campbell MM. Expansion and diversification of the *Populus* R2R3-MYB family of transcription factors. Plant Physiology. 2009;149:981–93.

Williams CA., Grayer RJ. Anthocyanins and other flavonoids. Natural Product Reports. 2004;21:539–73. 10.1039/B311404J.

Xia D, Zhou H, Wang Y, Li P, Fu P, Wu B, He Y. How rice organs are colored: The genetic basis of anthocyanin biosynthesis in rice. The Crop Journal. 2021;9:598–608. 10.1016/j.cj.2021.03.013.

Xie DY, Sharma SB, Paiva NL, Ferreir D, Dixon RA. Role of anthocyanidin reductase, encoded by BANYULS in plant flavonoid biosynthesis. Science. 2003;299:396–9. 10.1126/science.1078540.

Xie T, Chen C, Li C, Liu J, Liu C, He Y. Genome-wide investigation of WRKY gene family in pineapple: evolution and expression profiles during development and stress. BMC Genomics. 2018;19:490. 10.1186/s12864-018-4880-x.

Yu X, Zhu Y, Fan J, Wang D, Gong X, Ouyang Z. Accumulation of flavonoid glycosides and UFGT gene expression in mulberry leaves (*Morus alba* L.) before and after frost. Chemistry and Biodiversity. 2017;14:e1600496. 10.1002/cbdv.201600496.

Zhang L, Guo X, Wang Z, Wang M, Hu Q. Characterization of the complete chloroplast genome of *Musella lasiocarpa*. Mitochondrial DNA Part B. 2018;3:728–9. 10.1080/23802359.2018.1456367.

Zhang L, Cenci A, Rouard M, Zhang D, Wang Y, Tang W, Zheng SJ. Transcriptomic analysis of resistant and susceptible banana corms in response to infection by *Fusarium oxysporum* f. sp. *cubense* tropical race 4. Scientific Reports. 2019;9:8199. 10.1038/s41598-019-44637-x.

Zhang L, Wu S, Chang X, Wang X, Zhao Y, Xia Y, Trigiano RN, Jiao Y, Chen F. The ancient wave of polyploidization events in flowering plants and their facilitated adaptation to environmental stress. Plant Cell Environment. 2020;43:2847–56. 10.1111/pce.13898.

Zhang S, Gao S, Chen Y, Xu S, Yu S, Zhou J. Identification of hydroxylation enzymes and the metabolic analysis of dihydromyricetin synthesis in *Ampelopsis grossedentata*. Genes. 2022;13:2318. 10.3390/genes13122318.

Zhang Z, Li DW, Jin JH, Yin YX, Zhang HX, Chai WG, Gong ZH. VIGS approach reveals the modulation of anthocyanin biosynthetic genes by *CaMYB* in chili pepper leaves. Frontiers in Plant Science. 2015;6:1‒10. 10.3389/fpls.2015.00500.

Zhao C, Wang F, Lian Y, Xiao H, Zheng J. Biosynthesis of citrus flavonoids and their health effects. Critical Reviews in Food Science and Nutrition. 2020;60:566–83. 10.1080/10408398.2018.1544885.

Zhao J. Flavonoid transport mechanisms: how to go, and with whom. Trends Plant Science. 2015;20:576–85. 10.1016/j.tplants.2015.06.007.

Zheng J, Wu H, Zhu H, Huang C, Liu C, Chang Y, Kong Z, Zhou Z, Wang G, Lin Y, et al. Determining factors, regulation system, and domestication of anthocyanin biosynthesis in rice leaves. New Phytologist. 2019;223:705–21. 10.1111/nph.15807.

