## Supplemental Figure 1-4 for "Genome-wide analysis of flavonoid biosynthetic genes in Musaceae (*Ensete*, *Musella*, and *Musa* species) reveals amplification of flavonoid 3’,5’-hydroxylase"

**Additional file 1:**

**Figure S1.** Chromosome localization of flavonoid biosynthetic genes identified in Musaceae.

**Figure S2.** Phylogenetic trees show the genetic relationship of flavonoid biosynthetic genes in Arabidopsis, *B. distachyon*, *A. longiglumis*, *O. sativa*, *S. bicolor*, *Z. officinale*, *E. glaucum*, *M. lasiocarpa*, *M. beccarii*, *M. acuminata*, *M. balbisiana*, and *M. schizocarpa*.

**Figure S3.** Synteny analysis of FBGs between six Musaceae species.

**Figure S4.** Cis-acting elements in flavonoid biosynthetic genes of Musaceae.

**A** *M. acuminata*

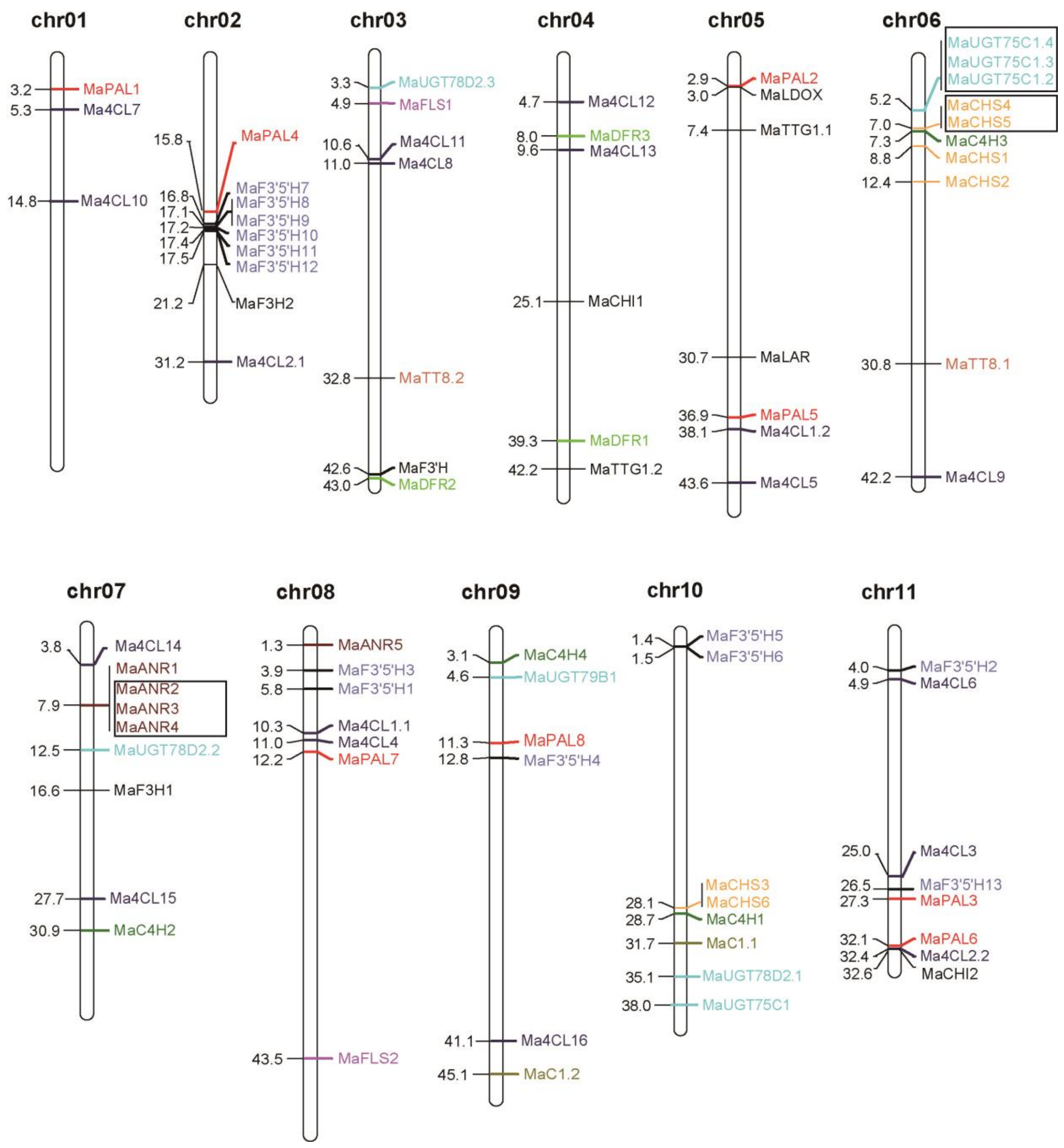


Flavonoid biosynthetic genes in Musaceae.

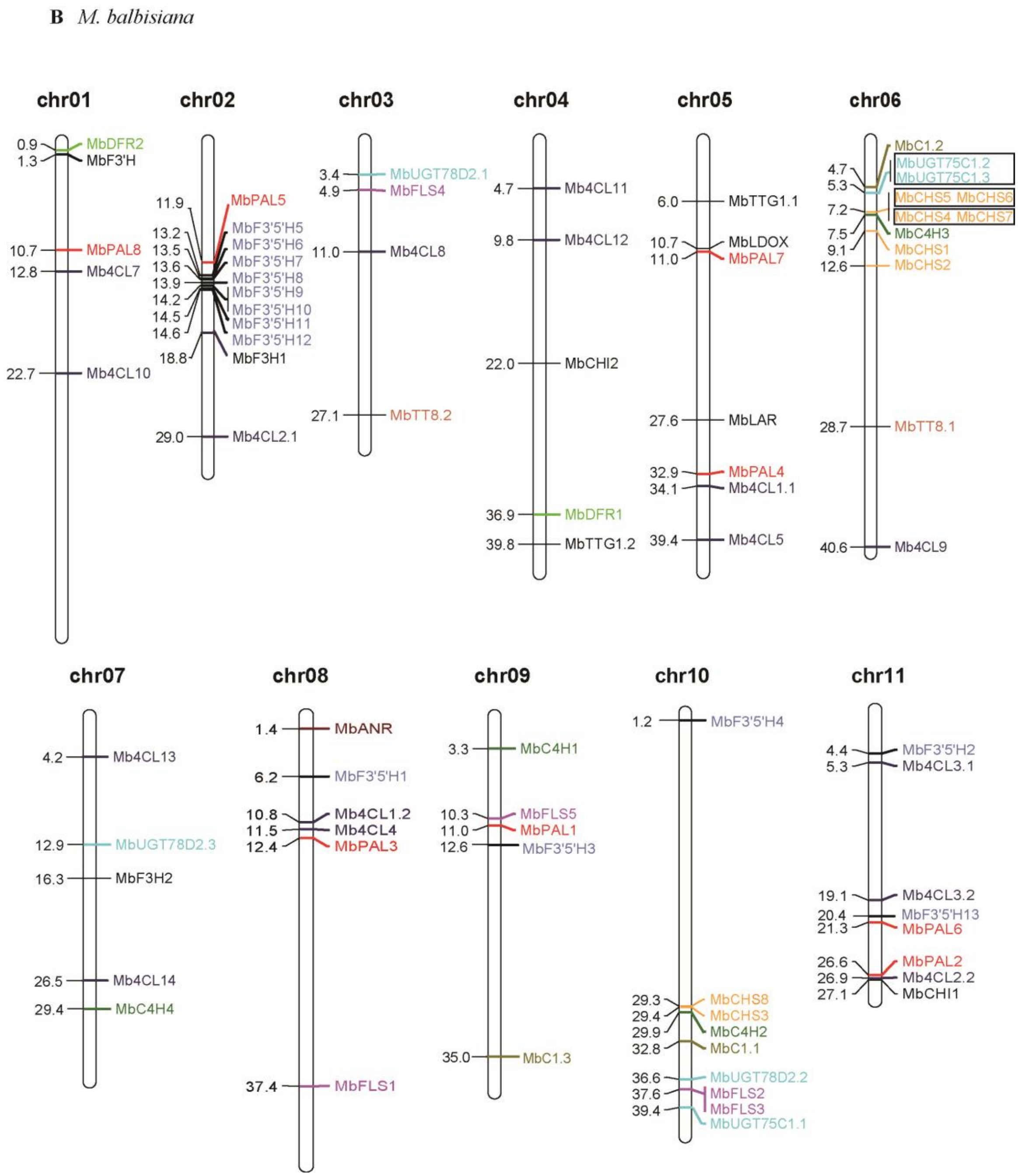


14

Flavonoid biosynthetic genes in Musaceae.

**C** *M. schizocharpa*

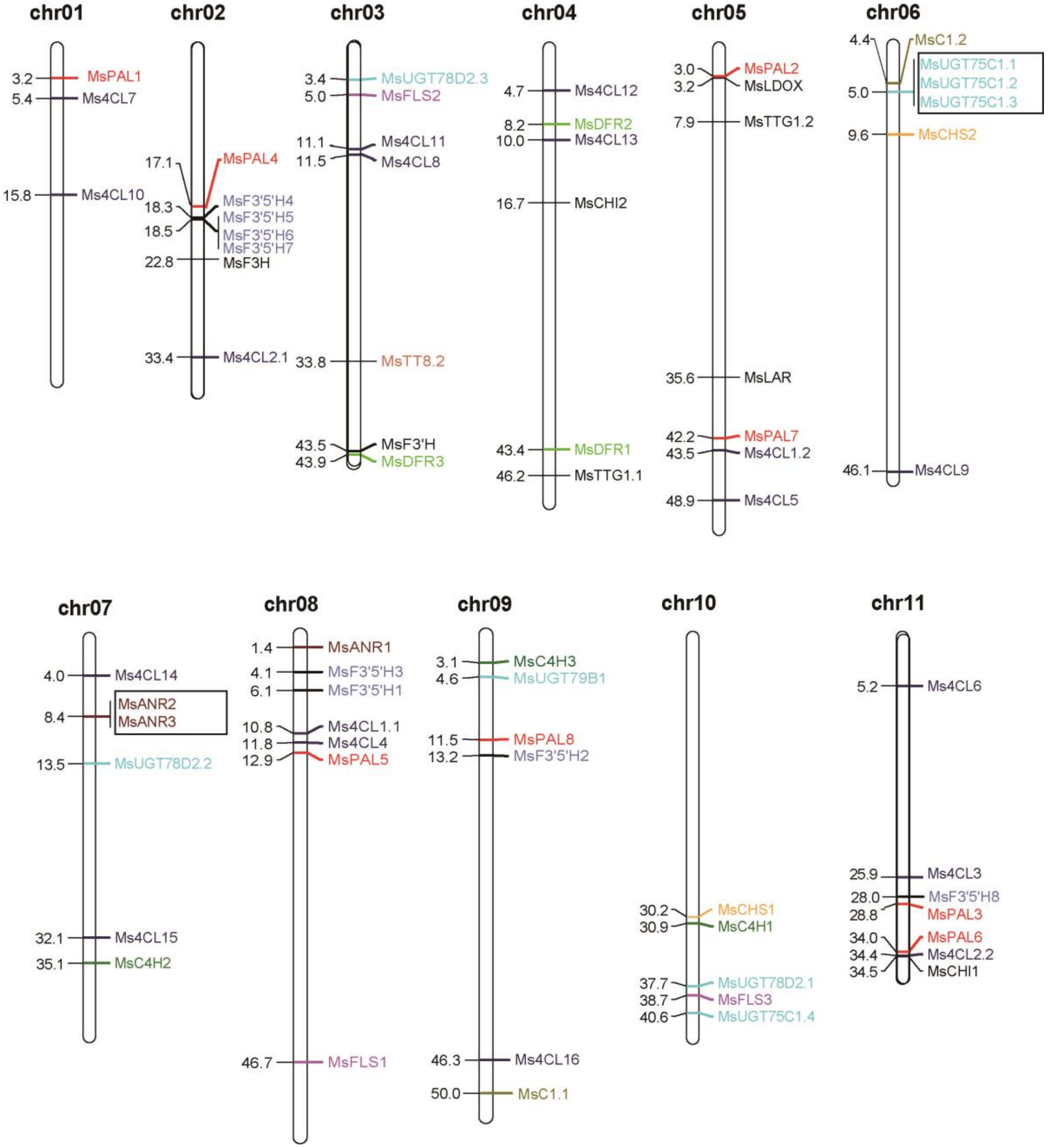


15

Flavonoid biosynthetic genes in Musaceae.

**D** *M. beccarii*

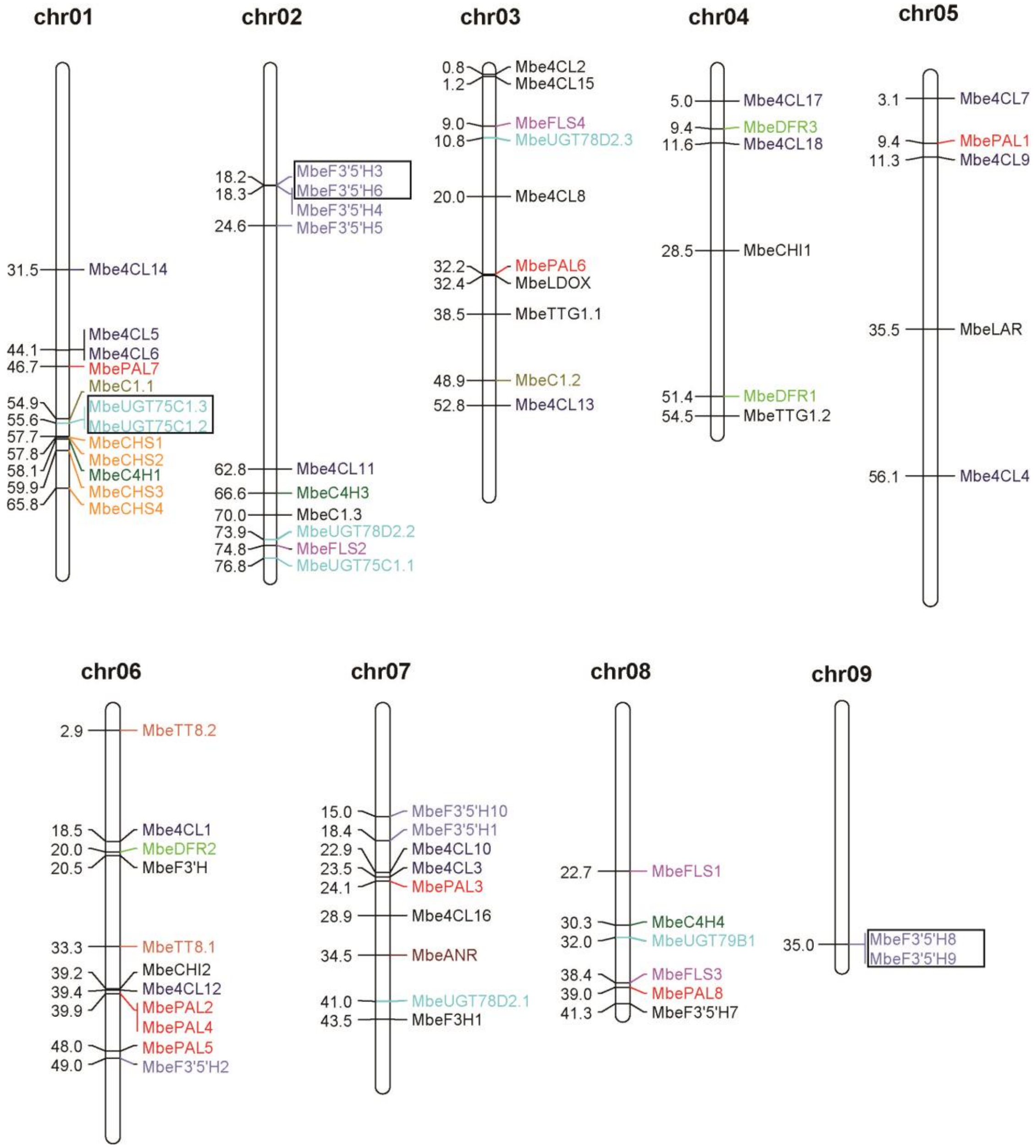


16

Flavonoid biosynthetic genes in Musaceae.

**E** *M. lasiocarpa*

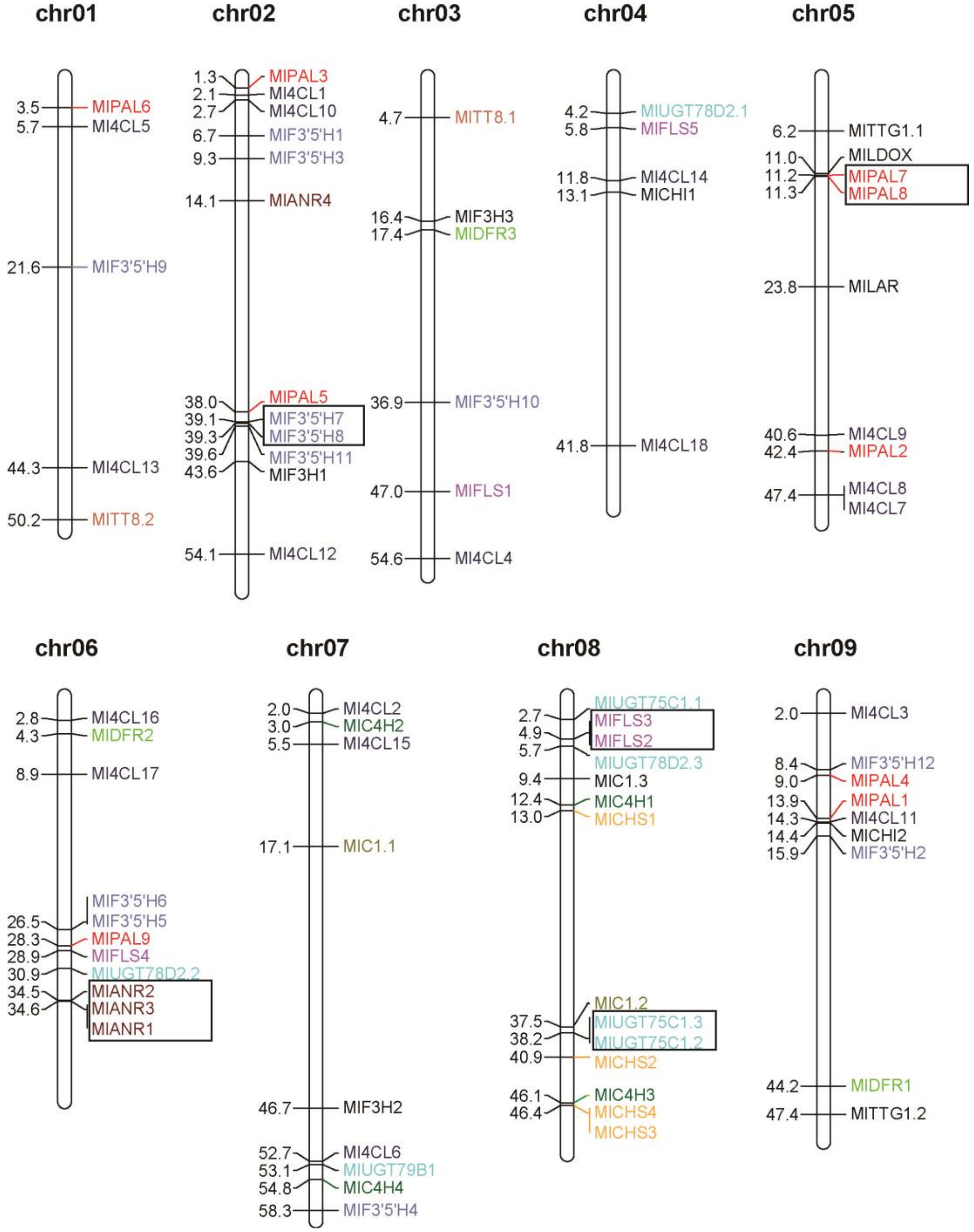


17

Flavonoid biosynthetic genes in Musaceae.

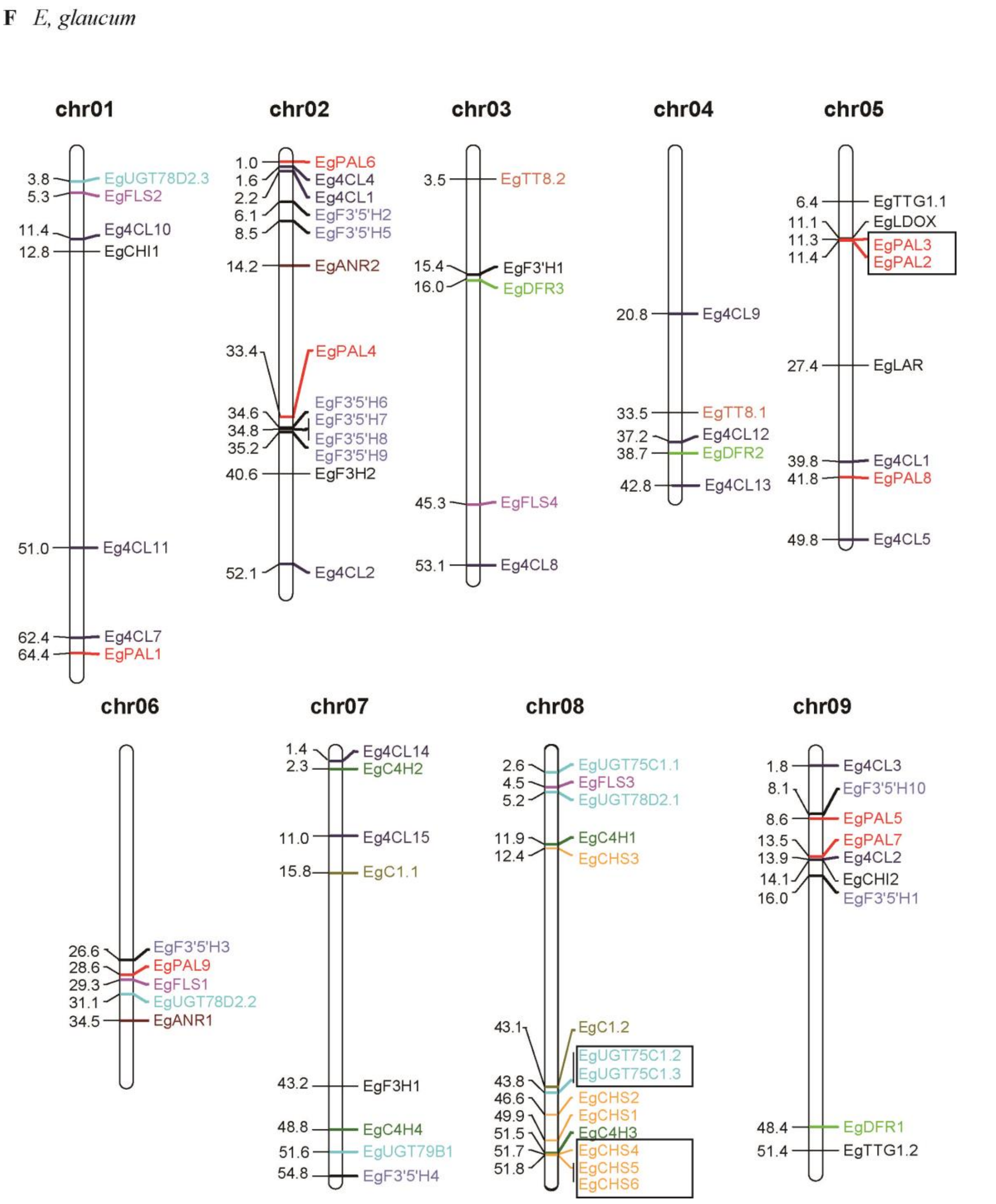


**Figure S1. Chromosome localization of flavonoid biosynthetic genes (FBGs) identified in Musaceae.** (**A**) FBGs in *Musa acuminata* (*x*=11). (**B**) FBGs in *M. balbisiana* (*x*=11). (**C**) FBGs in *M. schizocarpa* (*x*=11). (**D**) FBGs in *M. beccarii* (*x*=9). (**E**) FBGs in *Musella lasiocarpa* (*x*=9). (**F**) FBGs in *Ensete glaucum* (*x*=9).

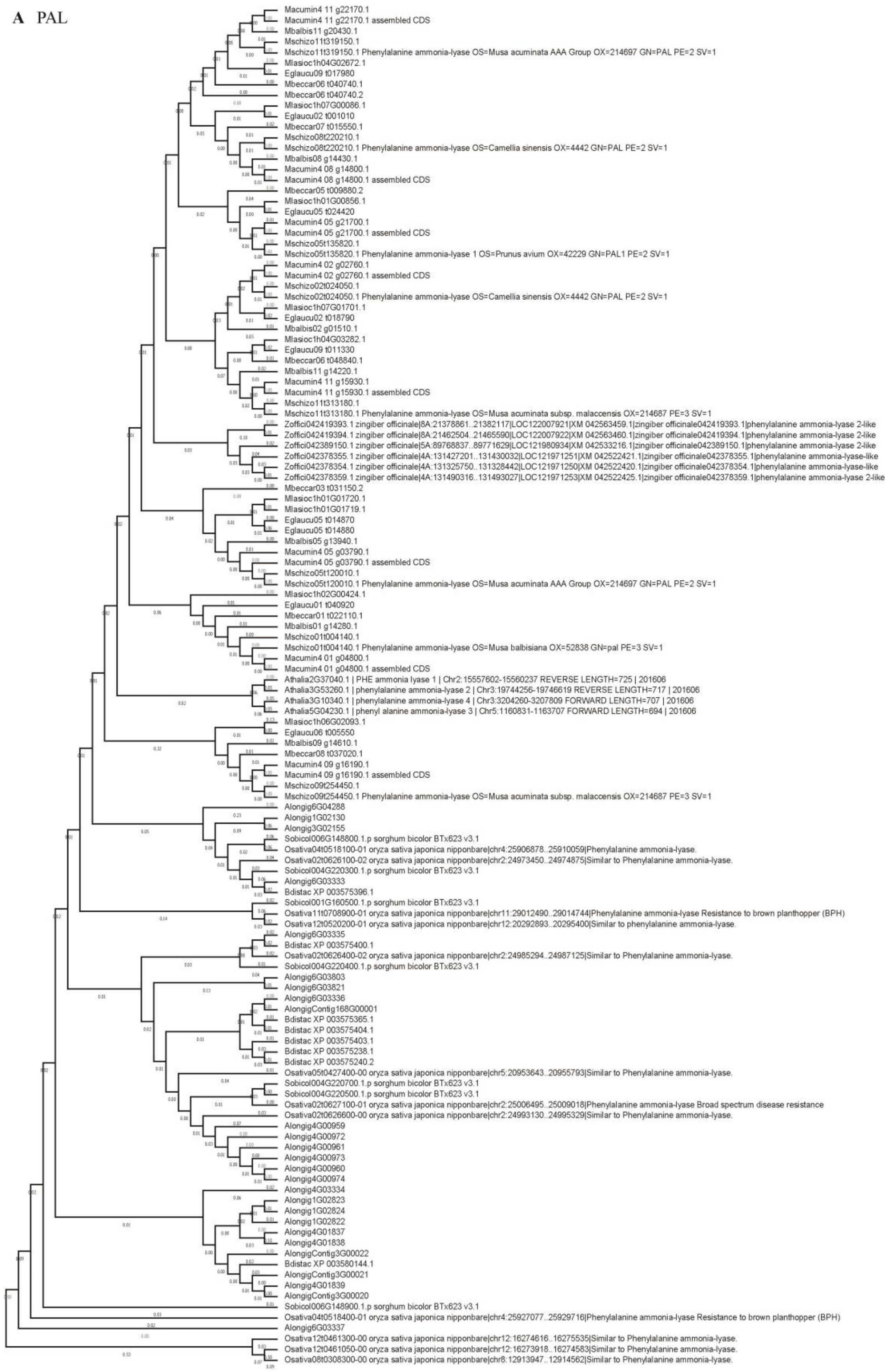
A PAL
Macumin4 11 g22170.1
Macumin4 11 g22170.1 assembled CDS
Mbalbis11 g20430.1
Mschizo11t319150.1
Mschizo11t319150.1 Phenylalanine ammonia-lyase OS=Musa acuminata AAA Group OX=214697 GN=PAL PE=2 SV=1
Mlasioc1h04G02672.1
Eglaucu09 t017980
Mbeccar06 t040740.1
Mbeccar06 t040740.2
Mlasioc1h07G00086.1
Eglaucu02 t001010
Mbeccar07 t015550.1
Mschizo08t220210.1
Mschizo08t220210.1 Phenylalanine ammonia-lyase OS=Camellia sinensis OX=4442 GN=PAL PE=2 SV=1
Mbalbis08 g14430.1
Macumin4 08 g14800.1
Macumin4 08 g14800.1 assembled CDS
Mbeccar05 t009880.2
Mlasioc1h01G00856.1
Eglaucu05 t024420
Macumin4 05 g21700.1
Macumin4 05 g21700.1 assembled CDS
Mschizo05t135820.1
Mschizo05t135820.1 Phenylalanine ammonia-lyase 1 OS=Prunus avium OX=42229 GN=PAL1 PE=2 SV=1
Macumin4 02 g02760.1
Macumin4 02 g02760.1 assembled CDS
Mschizo02t024050.1
Mschizo02t024050.1 Phenylalanine ammonia-lyase OS=Camellia sinensis OX=4442 GN=PAL PE=2 SV=1
Mlasioc1h07G01701.1
Eglaucu02 t018790
Mbalbis02 g01510.1
Mlasioc1h04G03282.1
Eglaucu09 t011330
Mbeccar06 t048840.1
Mbalbis11 g14220.1
Macumin4 11 g15930.1
Macumin4 11 g15930.1 assembled CDS
Mschizo11t313180.1
Mschizo11t313180.1 Phenylalanine ammonia-lyase OS=Musa acuminata subsp. malaccensis OX=214687 PE=3 SV=1
Zoffici042419393.1 zingiber officinale|8A:21378861..21382117|LOC122007921|XM 042563459.1|zingiber officinale042419393.1|phenylalanine ammonia-lyase 2-like
Zoffici042419394.1 zingiber officinale|8A:21462504..21465590|LOC122007922|XM 042563460.1|zingiber officinale042419394.1|phenylalanine ammonia-lyase 2-like
Zoffici042389150.1 zingiber officinale|5A:89768837..89771629|LOC121980934|XM 042533216.1|zingiber officinale042389150.1|phenylalanine ammonia-lyase 2-like
Zoffici042378355.1 zingiber officinale|4A:131427201..131430032|LOC121971251|XM 042522421.1|zingiber officinale042378355.1|phenylalanine ammonia-lyase-like
Zoffici042378354.1 zingiber officinale|4A:131325750..131328442|LOC121971250|XM 042522420.1|zingiber officinale042378354.1|phenylalanine ammonia-lyase-like
Zoffici042378359.1 zingiber officinale|4A:131490316..131493027|LOC121971253|XM 042522425.1|zingiber officinale042378359.1|phenylalanine ammonia-lyase 2-like
Mbeccar03 t031150.2
Mlasioc1h01G01720.1
Mlasioc1h01G01719.1
Eglaucu05 t014870
Eglaucu05 t014880
Mbalbis05 g13940.1
Macumin4 05 g03790.1
Macumin4 05 g03790.1 assembled CDS
Mschizo05t120010.1
Mschizo05t120010.1 Phenylalanine ammonia-lyase OS=Musa acuminata AAA Group OX=214697 GN=PAL PE=2 SV=1
Mlasioc1h02G00424.1
Eglaucu01 t040920
Mbeccar01 t022110.1
Mbalbis01 g14280.1
Mschizo01t004140.1
Mschizo01t004140.1 Phenylalanine ammonia-lyase OS=Musa balbisiana OX=52838 GN=pal PE=3 SV=1
Macumin4 01 g04800.1
Macumin4 01 g04800.1 assembled CDS
Athalia2G37040.1 | PHE ammonia lyase 1 | Chr2:15557602-15560237 REVERSE LENGTH=725 | 201606
Athalia3G53260.1 | phenylalanine ammonia-lyase 2 | Chr3:19744256-19746619 REVERSE LENGTH=717 | 201606
Athalia3G10340.1 | phenylalanine ammonia-lyase 4 | Chr3:3204260-3207809 FORWARD LENGTH=707 | 201606
Athalia5G04230.1 | phenyl alanine ammonia-lyase 3 | Chr5:1160831-1163707 FORWARD LENGTH=694 | 201606
Mlasioc1h06G02093.1
Eglaucu06 t005550
Mbalbis09 g14610.1
Mbeccar08 t037020.1
Macumin4 09 g16190.1
Macumin4 09 g16190.1 assembled CDS
Mschizo09t254450.1
Mschizo09t254450.1 Phenylalanine ammonia-lyase OS=Musa acuminata subsp. malaccensis OX=214687 PE=3 SV=1
Alongig6G04288
Alongig1G02130
Alongig3G02155
Sobicol006G148800.1.p sorghum bicolor BTx623 v3.1
Osativa04t0518100-01 oryza sativa japonica nipponbare|chr4:25906878..25910059|Phenylalanine ammonia-lyase.
Osativa02t0626100-02 oryza sativa japonica nipponbare|chr2:24973450..24974875|Similar to Phenylalanine ammonia-lyase.
Sobicol004G220300.1.p sorghum bicolor BTx623 v3.1
Alongig6G03333
Bdistac XP 003575396.1
Sobicol001G160500.1.p sorghum bicolor BTx623 v3.1
Osativa11t0708900-01 oryza sativa japonica nipponbare|chr11:29012490..29014744|Phenylalanine ammonia-lyase Resistance to brown planthopper (BPH)
Osativa12t0520200-01 oryza sativa japonica nipponbare|chr12:20292893..20295400|Similar to phenylalanine ammonia-lyase.
Alongig6G03335
Bdistac XP 003575400.1
Osativa02t0626400-02 oryza sativa japonica nipponbare|chr2:24985294..24987125|Similar to Phenylalanine ammonia-lyase.
Sobicol004G220400.1.p sorghum bicolor BTx623 v3.1
Alongig6G03803
Alongig6G03821
Alongig6G03336
AlongigContig168G00001
Bdistac XP 003575365.1
Bdistac XP 003575404.1
Bdistac XP 003575403.1
Bdistac XP 003575238.1
Bdistac XP 003575240.2
Osativa05t0427400-00 oryza sativa japonica nipponbare|chr5:20953643..20955793|Similar to Phenylalanine ammonia-lyase.
Sobicol004G220700.1.p sorghum bicolor BTx623 v3.1
Sobicol004G220500.1.p sorghum bicolor BTx623 v3.1
Osativa02t0627100-01 oryza sativa japonica nipponbare|chr2:25006495..25009018|Phenylalanine ammonia-lyase Broad spectrum disease resistance
Osativa02t0626600-00 oryza sativa japonica nipponbare|chr2:24993130..24995329|Similar to Phenylalanine ammonia-lyase.
Alongig4G00959
Alongig4G00972
Alongig4G00961
Alongig4G00973
Alongig4G00960
Alongig4G00974
Alongig4G03334
Alongig1G02823
Alongig1G02824
Alongig1G02822
Alongig4G01837
Alongig4G01838
AlongigContig3G00022
Bdistac XP 003580144.1
AlongigContig3G00021
Alongig4G01839
AlongigContig3G00020
Sobicol006G148900.1.p sorghum bicolor BTx623 v3.1
Osativa04t0518400-01 oryza sativa japonica nipponbare|chr4:25927077..25929716|Phenylalanine ammonia-lyase Resistance to brown planthopper (BPH)
Alongig6G03337
Osativa12t0461300-00 oryza sativa japonica nipponbare|chr12:16274616..16275535|Similar to Phenylalanine ammonia-lyase.
Osativa12t0461050-00 oryza sativa japonica nipponbare|chr12:16273918..16274583|Similar to Phenylalanine ammonia-lyase.
Osativa08t0308300-00 oryza sativa japonica nipponbare|chr8:12913947..12914562|Similar to Phenylalanine ammonia-lyase.

24

**B** 4CL

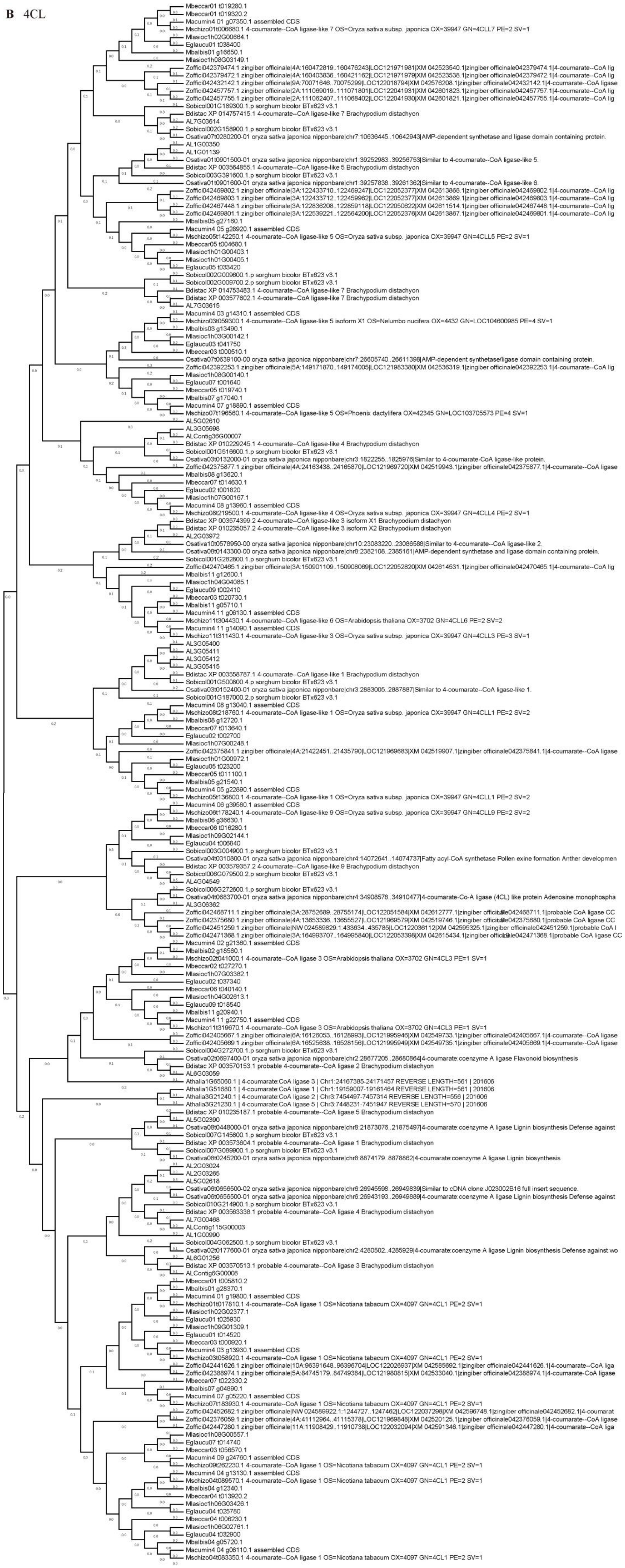
Mbeccar01 t019280.1
Mbeccar01 t019320.2
Macumin4 01 g07350.1 assembled CDS
Mschizo01t006680.1 4-coumarate--CoA ligase-like 7 OS=Oryza sativa subsp. japonica OX=39947 GN=4CLL7 PE=2 SV=1
Mlasioc1h02G00664.1
Eglaucu01 t038400
Mbalbis01 g16650.1
Mlasioc1h08G03149.1
Zoffici042379474.1 zingiber officinale|4A:160472819..160476243|LOC121971981|XM 042523540.1|zingiber officinale042379474.1|4-coumarate--CoA lig
Zoffici042379472.1 zingiber officinale|4A:160403836..160421162|LOC121971979|XM 042523538.1|zingiber officinale042379472.1|4-coumarate--CoA lig
Zoffici042432142.1 zingiber officinale|9A:70071646..70075299|LOC122018794|XM 042576208.1|zingiber officinale042432142.1|4-coumarate--CoA ligase
Zoffici042457757.1 zingiber officinale|2A:111069019..111071801|LOC122041931|XM 042601823.1|zingiber officinale042457757.1|4-coumarate--CoA lig
Zoffici042457755.1 zingiber officinale|2A:111062407..111068402|LOC122041930|XM 042601821.1|zingiber officinale042457755.1|4-coumarate--CoA lig
Sobicol001G189300.1.p sorghum bicolor BTx623 v3.1
Bdistac XP 014757415.1 4-coumarate--CoA ligase-like 7 Brachypodium distachyon
AL7G03614
Sobicol002G158900.1.p sorghum bicolor BTx623 v3.1
Osativa07t0280200-01 oryza sativa japonica nipponbare|chr7:10636445..10642943|AMP-dependent synthetase and ligase domain containing protein.
AL1G00350
AL1G01139
Osativa01t0901500-01 oryza sativa japonica nipponbare|chr1:39252983..39256753|Similar to 4-coumarate--CoA ligase-like 5.
Bdistac XP 003564855.1 4-coumarate--CoA ligase-like 5 Brachypodium distachyon
Sobicol003G391600.1.p sorghum bicolor BTx623 v3.1
Osativa01t0901600-01 oryza sativa japonica nipponbare|chr1:39257838..39261362|Similar to 4-coumarate--CoA ligase-like 6.
Zoffici042469802.1 zingiber officinale|3A:122433710..122469247|LOC122052377|XM 042613868.1|zingiber officinale042469802.1|4-coumarate--CoA lig
Zoffici042469803.1 zingiber officinale|3A:122433712..122459962|LOC122052377|XM 042613869.1|zingiber officinale042469803.1|4-coumarate--CoA lig
Zoffici042467448.1 zingiber officinale|3A:122836208..122859118|LOC122050622|XM 042611514.1|zingiber officinale042467448.1|4-coumarate--CoA lig
Zoffici042469801.1 zingiber officinale|3A:122539221..122564200|LOC122052376|XM 042613867.1|zingiber officinale042469801.1|4-coumarate--CoA lig
Mbalbis05 g27160.1
Macumin4 05 g28920.1 assembled CDS
Mschizo05t142250.1 4-coumarate--CoA ligase-like 5 OS=Oryza sativa subsp. japonica OX=39947 GN=4CLL5 PE=2 SV=1
Mbeccar05 t004680.1
Mlasioc1h01G00403.1
Mlasioc1h01G00405.1
Eglaucu05 t033420
Sobicol002G009600.1.p sorghum bicolor BTx623 v3.1
Sobicol002G009700.2.p sorghum bicolor BTx623 v3.1
Bdistac XP 014753483.1 4-coumarate--CoA ligase-like 7 Brachypodium distachyon
Bdistac XP 003577602.1 4-coumarate--CoA ligase-like 7 Brachypodium distachyon
AL7G03615
Macumin4 03 g14310.1 assembled CDS
Mschizo03t059300.1 4-coumarate--CoA ligase-like 5 isoform X1 OS=Nelumbo nucifera OX=4432 GN=LOC104600985 PE=4 SV=1
Mbalbis03 g13490.1
Mlasioc1h03G00142.1
Eglaucu03 t041750
Mbeccar03 t000510.1
Osativa07t0639100-00 oryza sativa japonica nipponbare|chr7:26605740..26611398|AMP-dependent synthetase/ligase domain containing protein.
Zoffici042392253.1 zingiber officinale|5A:149171870..149174005|LOC121983380|XM 042536319.1|zingiber officinale042392253.1|4-coumarate--CoA lig
Mlasioc1h08G00140.1
Eglaucu07 t001640
Mbeccar05 t019740.1
Mbalbis07 g17040.1
Macumin4 07 g18890.1 assembled CDS
Mschizo07t196560.1 4-coumarate--CoA ligase-like 5 OS=Phoenix dactylifera OX=42345 GN=LOC103705573 PE=4 SV=1
AL5G02610
AL3G05698
ALContig36G00007
Bdistac XP 010229245.1 4-coumarate--CoA ligase-like 4 Brachypodium distachyon
Sobicol001G516600.1.p sorghum bicolor BTx623 v3.1
Osativa03t0132000-01 oryza sativa japonica nipponbare|chr3:1822255..1825976|Similar to 4-coumarate-CoA ligase-like protein.
Zoffici042375877.1 zingiber officinale|4A:24163438..24165870|LOC121969720|XM 042519943.1|zingiber officinale042375877.1|4-coumarate--CoA ligase
Mbalbis08 g13620.1
Mbeccar07 t014630.1
Eglaucu02 t001820
Mlasioc1h07G00167.1
Macumin4 08 g13960.1 assembled CDS
Mschizo08t219500.1 4-coumarate--CoA ligase-like 4 OS=Oryza sativa subsp. japonica OX=39947 GN=4CLL4 PE=2 SV=1
Bdistac XP 003574399.2 4-coumarate--CoA ligase-like 3 isoform X1 Brachypodium distachyon
Bdistac XP 010235057.2 4-coumarate--CoA ligase-like 3 isoform X2 Brachypodium distachyon
AL2G03972
Osativa10t0578950-00 oryza sativa japonica nipponbare|chr10:23083220..23086588|Similar to 4-coumarate--CoA ligase-like 2.
Osativa08t0143300-00 oryza sativa japonica nipponbare|chr8:2382108..2385161|AMP-dependent synthetase and ligase domain containing protein.
Sobicol001G282800.1.p sorghum bicolor BTx623 v3.1
Zoffici042470465.1 zingiber officinale|3A:150901109..150908069|LOC122052820|XM 042614531.1|zingiber officinale042470465.1|4-coumarate--CoA lig
Mbalbis11 g12600.1
Mlasioc1h04G04085.1
Eglaucu09 t002410
Mbeccar03 t020730.1
Mbalbis11 g05710.1
Macumin4 11 g06130.1 assembled CDS
Mschizo11t304430.1 4-coumarate--CoA ligase-like 6 OS=Arabidopsis thaliana OX=3702 GN=4CLL6 PE=2 SV=2
Macumin4 11 g14090.1 assembled CDS
Mschizo11t311430.1 4-coumarate--CoA ligase-like 3 OS=Oryza sativa subsp. japonica OX=39947 GN=4CLL3 PE=3 SV=1
AL3G05400
AL3G05411
AL3G05412
AL3G05415
Bdistac XP 003558787.1 4-coumarate--CoA ligase-like 1 Brachypodium distachyon
Sobicol001G500800.4.p sorghum bicolor BTx623 v3.1
Osativa03t0152400-01 oryza sativa japonica nipponbare|chr3:2883005..2887887|Similar to 4-coumarate--CoA ligase-like 1.
Sobicol001G187000.2.p sorghum bicolor BTx623 v3.1
Macumin4 08 g13040.1 assembled CDS
Mschizo08t218760.1 4-coumarate--CoA ligase-like 1 OS=Oryza sativa subsp. japonica OX=39947 GN=4CLL1 PE=2 SV=2
Mbalbis08 g12720.1
Mbeccar07 t013640.1
Eglaucu02 t002700
Mlasioc1h07G00248.1
Zoffici042375841.1 zingiber officinale|4A:21422451..21435790|LOC121969683|XM 042519907.1|zingiber officinale042375841.1|4-coumarate--CoA ligase
Mlasioc1h01G00972.1
Eglaucu05 t023200
Mbeccar05 t011100.1
Mbalbis05 g21540.1
Macumin4 05 g22890.1 assembled CDS
Mschizo05t136800.1 4-coumarate--CoA ligase-like 1 OS=Oryza sativa subsp. japonica OX=39947 GN=4CLL1 PE=2 SV=2
Macumin4 06 g39580.1 assembled CDS
Mschizo06t178240.1 4-coumarate--CoA ligase-like 9 OS=Oryza sativa subsp. japonica OX=39947 GN=4CLL9 PE=2 SV=2
Mbalbis06 g36630.1
Mbeccar06 t016280.1
Mlasioc1h09G02144.1
Eglaucu04 t006840
Sobicol003G004900.1.p sorghum bicolor BTx623 v3.1
Osativa04t0310800-01 oryza sativa japonica nipponbare|chr4:14072641..14074737|Fatty acyl-CoA synthetase Pollen exine formation Anther developmen
Bdistac XP 003579357.2 4-coumarate--CoA ligase-like 9 Brachypodium distachyon
Sobicol006G079500.2.p sorghum bicolor BTx623 v3.1
AL4G04549
Sobicol006G272600.1.p sorghum bicolor BTx623 v3.1
Osativa04t0683700-01 oryza sativa japonica nipponbare|chr4:34908578..34910477|4-coumarate-Co-A ligase (4CL) like protein Adenosine monophospha
AL3G06362
Zoffici042468711.1 zingiber officinale|3A:28752689..28755174|LOC122051584|XM 042612777.1|zingiber officinale042468711.1|probable CoA ligase CC
Zoffici042375680.1 zingiber officinale|4A:13653336..13655527|LOC121969579|XM 042519746.1|zingiber officinale042375680.1|probable CoA ligase CC
Zoffici042451259.1 zingiber officinale|NW 024589829.1:433634..435785|LOC122036112|XM 042595325.1|zingiber officinale042451259.1|probable CoA l
Zoffici042471368.1 zingiber officinale|3A:164993707..164995840|LOC122053398|XM 042615434.1|zingiber officinale042471368.1|probable CoA ligase CC
Macumin4 02 g21360.1 assembled CDS
Mbalbis02 g18560.1
Mschizo02t041000.1 4-coumarate--CoA ligase 3 OS=Arabidopsis thaliana OX=3702 GN=4CL3 PE=1 SV=1
Mbeccar02 t027270.1
Mlasioc1h07G03382.1
Eglaucu02 t037340
Mbeccar06 t040140.1
Mlasioc1h04G02613.1
Eglaucu09 t018540
Mbalbis11 g20940.1
Macumin4 11 g22750.1 assembled CDS
Mschizo11t319670.1 4-coumarate--CoA ligase 3 OS=Arabidopsis thaliana OX=3702 GN=4CL3 PE=1 SV=1
Zoffici042405667.1 zingiber officinale|6A:16126053..16128993|LOC121995946|XM 042549733.1|zingiber officinale042405667.1|4-coumarate--CoA ligase
Zoffici042405669.1 zingiber officinale|6A:16525638..16528156|LOC121995949|XM 042549735.1|zingiber officinale042405669.1|4-coumarate--CoA ligase
Sobicol004G272700.1.p sorghum bicolor BTx623 v3.1
Osativa02t0697400-01 oryza sativa japonica nipponbare|chr2:28677205..28680864|4-coumarate:coenzyme A ligase Flavonoid biosynthesis
Bdistac XP 003570153.1 probable 4-coumarate--CoA ligase 2 Brachypodium distachyon
AL6G03059
Athalia1G65060.1 | 4-coumarate:CoA ligase 3 | Chr1:24167385-24171457 REVERSE LENGTH=561 | 201606
Athalia1G51680.1 | 4-coumarate:CoA ligase 1 | Chr1:19159007-19161464 REVERSE LENGTH=561 | 201606
Athalia3G21240.1 | 4-coumarate:CoA ligase 2 | Chr3:7454497-7457314 REVERSE LENGTH=556 | 201606
Athalia3G21230.1 | 4-coumarate:CoA ligase 5 | Chr3:7448231-7451947 REVERSE LENGTH=570 | 201606
Bdistac XP 010235187.1 probable 4-coumarate--CoA ligase 5 Brachypodium distachyon
AL5G02390
Osativa08t0448000-01 oryza sativa japonica nipponbare|chr8:21873076..21875497|4-coumarate:coenzyme A ligase Lignin biosynthesis Defense against
Sobicol007G145600.1.p sorghum bicolor BTx623 v3.1
Bdistac XP 003573604.1 probable 4-coumarate--CoA ligase 1 Brachypodium distachyon
Sobicol007G089900.1.p sorghum bicolor BTx623 v3.1
Osativa08t0245200-01 oryza sativa japonica nipponbare|chr8:8874179..8878862|4-coumarate:coenzyme A ligase Lignin biosynthesis
AL2G03024
AL2G03265
AL5G02618
Osativa06t0656500-02 oryza sativa japonica nipponbare|chr6:26945598..26949839|Similar to cDNA clone:J023002B16 full insert sequence.
Osativa06t0656500-01 oryza sativa japonica nipponbare|chr6:26943193..26949889|4-coumarate:coenzyme A ligase Lignin biosynthesis Defense against
Sobicol010G214900.1.p sorghum bicolor BTx623 v3.1
Bdistac XP 003563338.1 probable 4-coumarate--CoA ligase 4 Brachypodium distachyon
AL7G00468
ALContig115G00003
AL1G00990
Sobicol004G062500.1.p sorghum bicolor BTx623 v3.1
Osativa02t0177600-01 oryza sativa japonica nipponbare|chr2:4280502..4285929|4-coumarate:coenzyme A ligase Lignin biosynthesis Defense against wo
AL6G01256
Bdistac XP 003570513.1 probable 4-coumarate--CoA ligase 3 Brachypodium distachyon
ALContig6G00008
Mbeccar01 t005810.2
Mbalbis01 g28370.1
Macumin4 01 g19800.1 assembled CDS
Mschizo01t017810.1 4-coumarate--CoA ligase 1 OS=Nicotiana tabacum OX=4097 GN=4CL1 PE=2 SV=1
Mlasioc1h08G02377.1
Eglaucu01 t025930
Mlasioc1h09G01309.1
Eglaucu01 t014520
Mbeccar03 t000920.1
Macumin4 03 g13930.1 assembled CDS
Mschizo03t058920.1 4-coumarate--CoA ligase 1 OS=Nicotiana tabacum OX=4097 GN=4CL1 PE=2 SV=1
Zoffici042441626.1 zingiber officinale|10A:96391648..96396704|LOC122026937|XM 042585692.1|zingiber officinale042441626.1|4-coumarate--CoA liga
Zoffici042388974.1 zingiber officinale|5A:84745179..84749384|LOC121980815|XM 042533040.1|zingiber officinale042388974.1|4-coumarate-CoA ligase
Mbeccar07 t022330.2
Mbalbis07 g04890.1
Macumin4 07 g05220.1 assembled CDS
Mschizo07t183930.1 4-coumarate--CoA ligase 1 OS=Nicotiana tabacum OX=4097 GN=4CL1 PE=2 SV=1
Zoffici042452682.1 zingiber officinale|NW 024589922.1:1244727..1247462|LOC122037298|XM 042596748.1|zingiber officinale042452682.1|4-coumarat
Zoffici042376059.1 zingiber officinale|4A:41112964..41115378|LOC121969848|XM 042520125.1|zingiber officinale042376059.1|4-coumarate--CoA ligase
Zoffici042447280.1 zingiber officinale|11A:11908429..11910738|LOC122032094|XM 042591346.1|zingiber officinale042447280.1|4-coumarate--CoA liga
Mlasioc1h08G00557.1
Eglaucu07 t014740
Mbeccar03 t056570.1
Macumin4 09 g24760.1 assembled CDS
Mschizo09t262230.1 4-coumarate--CoA ligase 1 OS=Nicotiana tabacum OX=4097 GN=4CL1 PE=2 SV=1
Macumin4 04 g13130.1 assembled CDS
Mschizo04t089570.1 4-coumarate--CoA ligase 1 OS=Nicotiana tabacum OX=4097 GN=4CL1 PE=2 SV=1
Mbalbis04 g12340.1
Mbeccar04 t013920.2
Mlasioc1h06G03426.1
Eglaucu04 t025780
Mbeccar04 t006230.1
Mlasioc1h06G02761.1
Eglaucu04 t032900
Mbalbis04 g05720.1
Macumin4 04 g06110.1 assembled CDS
Mschizo04t083350.1 4-coumarate--CoA ligase 1 OS=Nicotiana tabacum OX=4097 GN=4CL1 PE=2 SV=1

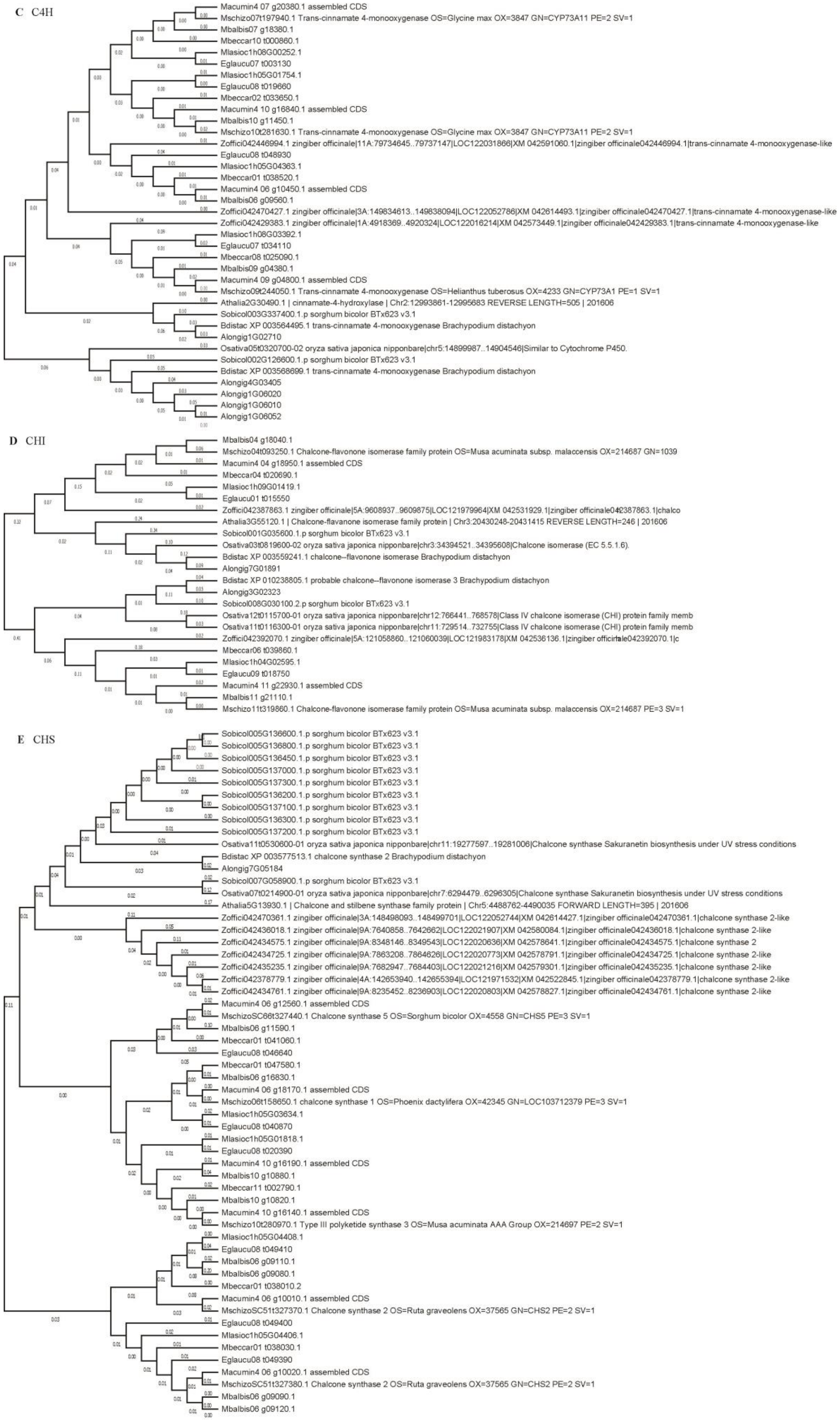
C C4H
Macumin4 07 g20380.1 assembled CDS
Mschizo07t197940.1 Trans-cinnamate 4-monooxygenase OS=Glycine max OX=3847 GN=CYP73A11 PE=2 SV=1
Mbalbis07 g18380.1
Mbeccar10 t000860.1
Mlasioc1h08G00252.1
Eglaucu07 t003130
Mlasioc1h05G01754.1
Eglaucu08 t019660
Mbeccar02 t033650.1
Macumin4 10 g16840.1 assembled CDS
Mbalbis10 g11450.1
Mschizo10t281630.1 Trans-cinnamate 4-monooxygenase OS=Glycine max OX=3847 GN=CYP73A11 PE=2 SV=1
Zoffici042446994.1 zingiber officinale|11A:79734645..79737147|LOC122031866|XM 042591060.1|zingiber officinale042446994.1|trans-cinnamate 4-monooxygenase-like
Eglaucu08 t048930
Mlasioc1h05G04363.1
Mbeccar01 t038520.1
Macumin4 06 g10450.1 assembled CDS
Mbalbis06 g09560.1
Zoffici042470427.1 zingiber officinale|3A:149834613..149838094|LOC122052786|XM 042614493.1|zingiber officinale042470427.1|trans-cinnamate 4-monooxygenase-like
Zoffici042429383.1 zingiber officinale|1A:4918369..4920324|LOC122016214|XM 042573449.1|zingiber officinale042429383.1|trans-cinnamate 4-monooxygenase-like
Mlasioc1h08G03392.1
Eglaucu07 t034110
Mbeccar08 t025090.1
Mbalbis09 g04380.1
Macumin4 09 g04800.1 assembled CDS
Mschizo09t244050.1 Trans-cinnamate 4-monooxygenase OS=Helianthus tuberosus OX=4233 GN=CYP73A1 PE=1 SV=1
Athalia2G30490.1 | cinnamate-4-hydroxylase | Chr2:12993861-12995683 REVERSE LENGTH=505 | 201606
Sobicol003G337400.1.p sorghum bicolor BTx623 v3.1
Bdistac XP 003564495.1 trans-cinnamate 4-monooxygenase Brachypodium distachyon
Alongig1G02710
Osativa05t0320700-02 oryza sativa japonica nipponbare|chr5:14899987..14904546|Similar to Cytochrome P450.
Sobicol002G126600.1.p sorghum bicolor BTx623 v3.1
Bdistac XP 003568699.1 trans-cinnamate 4-monooxygenase Brachypodium distachyon
Alongig4G03405
Alongig1G06020
Alongig1G06010
Alongig1G06052
D CHI
Mbalbis04 g18040.1
Mschizo04t093250.1 Chalcone-flavonone isomerase family protein OS=Musa acuminata subsp. malaccensis OX=214687 GN=1039
Macumin4 04 g18950.1 assembled CDS
Mbeccar04 t020690.1
Mlasioc1h09G01419.1
Eglaucu01 t015550
Zoffici042387863.1 zingiber officinale|5A:9608937..9609875|LOC121979964|XM 042531929.1|zingiber officinale042387863.1|chalco
Athalia3G55120.1 | Chalcone-flavanone isomerase family protein | Chr3:20430248-20431415 REVERSE LENGTH=246 | 201606
Sobicol001G035600.1.p sorghum bicolor BTx623 v3.1
Osativa03t0819600-02 oryza sativa japonica nipponbare|chr3:34394521..34395608|Chalcone isomerase (EC 5.5.1.6).
Bdistac XP 003559241.1 chalcone--flavonone isomerase Brachypodium distachyon
Alongig7G01891
Bdistac XP 010238805.1 probable chalcone--flavonone isomerase 3 Brachypodium distachyon
Alongig3G02323
Sobicol008G030100.2.p sorghum bicolor BTx623 v3.1
Osativa12t0115700-01 oryza sativa japonica nipponbare|chr12:766441..768578|Class IV chalcone isomerase (CHI) protein family memb
Osativa11t0116300-01 oryza sativa japonica nipponbare|chr11:729514..732755|Class IV chalcone isomerase (CHI) protein family memb
Zoffici042392070.1 zingiber officinale|5A:121058860..121060039|LOC121983178|XM 042536136.1|zingiber officinale042392070.1|c
Mbeccar06 t039860.1
Mlasioc1h04G02595.1
Eglaucu09 t018750
Macumin4 11 g22930.1 assembled CDS
Mbalbis11 g21110.1
Mschizo11t319860.1 Chalcone-flavonone isomerase family protein OS=Musa acuminata subsp. malaccensis OX=214687 PE=3 SV=1
E CHS
Sobicol005G136600.1.p sorghum bicolor BTx623 v3.1
Sobicol005G136800.1.p sorghum bicolor BTx623 v3.1
Sobicol005G136450.1.p sorghum bicolor BTx623 v3.1
Sobicol005G137000.1.p sorghum bicolor BTx623 v3.1
Sobicol005G137300.1.p sorghum bicolor BTx623 v3.1
Sobicol005G136200.1.p sorghum bicolor BTx623 v3.1
Sobicol005G137100.1.p sorghum bicolor BTx623 v3.1
Sobicol005G136300.1.p sorghum bicolor BTx623 v3.1
Sobicol005G137200.1.p sorghum bicolor BTx623 v3.1
Osativa11t0530600-01 oryza sativa japonica nipponbare|chr11:19277597..19281006|Chalcone synthase Sakuranetin biosynthesis under UV stress conditions
Bdistac XP 003577513.1 chalcone synthase 2 Brachypodium distachyon
Alongig7G05184
Sobicol007G058900.1.p sorghum bicolor BTx623 v3.1
Osativa07t0214900-01 oryza sativa japonica nipponbare|chr7:6294479..6296305|Chalcone synthase Sakuranetin biosynthesis under UV stress conditions
Athalia5G13930.1 | Chalcone and stilbene synthase family protein | Chr5:4488762-4490035 FORWARD LENGTH=395 | 201606
Zoffici042470361.1 zingiber officinale|3A:148498093..148499701|LOC122052744|XM 042614427.1|zingiber officinale042470361.1|chalcone synthase 2-like
Zoffici042436018.1 zingiber officinale|9A:7640858..7642662|LOC122021907|XM 042580084.1|zingiber officinale042436018.1|chalcone synthase 2-like
Zoffici042434575.1 zingiber officinale|9A:8348146..8349543|LOC122020636|XM 042578641.1|zingiber officinale042434575.1|chalcone synthase 2
Zoffici042434725.1 zingiber officinale|9A:7863208..7864626|LOC122020773|XM 042578791.1|zingiber officinale042434725.1|chalcone synthase 2-like
Zoffici042435235.1 zingiber officinale|9A:7682947..7684403|LOC122021216|XM 042579301.1|zingiber officinale042435235.1|chalcone synthase 2-like
Zoffici042378779.1 zingiber officinale|4A:142653940..142655394|LOC121971532|XM 042522845.1|zingiber officinale042378779.1|chalcone synthase 2-like
Zoffici042434761.1 zingiber officinale|9A:8235452..8236903|LOC122020803|XM 042578827.1|zingiber officinale042434761.1|chalcone synthase 2-like
Macumin4 06 g12560.1 assembled CDS
MschizoSC66t327440.1 Chalcone synthase 5 OS=Sorghum bicolor OX=4558 GN=CHS5 PE=3 SV=1
Mbalbis06 g11590.1
Mbeccar01 t041060.1
Eglaucu08 t046640
Mbeccar01 t047580.1
Mbalbis06 g16830.1
Macumin4 06 g18170.1 assembled CDS
Mschizo06t158650.1 chalcone synthase 1 OS=Phoenix dactylifera OX=42345 GN=LOC103712379 PE=3 SV=1
Mlasioc1h05G03634.1
Eglaucu08 t040870
Mlasioc1h05G01818.1
Eglaucu08 t020390
Macumin4 10 g16190.1 assembled CDS
Mbalbis10 g10880.1
Mbeccar11 t002790.1
Mbalbis10 g10820.1
Macumin4 10 g16140.1 assembled CDS
Mschizo10t280970.1 Type III polyketide synthase 3 OS=Musa acuminata AAA Group OX=214697 PE=2 SV=1
Mlasioc1h05G04408.1
Eglaucu08 t049410
Mbalbis06 g09110.1
Mbalbis06 g09080.1
Mbeccar01 t038010.2
Macumin4 06 g10010.1 assembled CDS
MschizoSC51t327370.1 Chalcone synthase 2 OS=Ruta graveolens OX=37565 GN=CHS2 PE=2 SV=1
Eglaucu08 t049400
Mlasioc1h05G04406.1
Mbeccar01 t038030.1
Eglaucu08 t049390
Macumin4 06 g10020.1 assembled CDS
MschizoSC51t327380.1 Chalcone synthase 2 OS=Ruta graveolens OX=37565 GN=CHS2 PE=2 SV=1
Mbalbis06 g09090.1
Mbalbis06 g09120.1

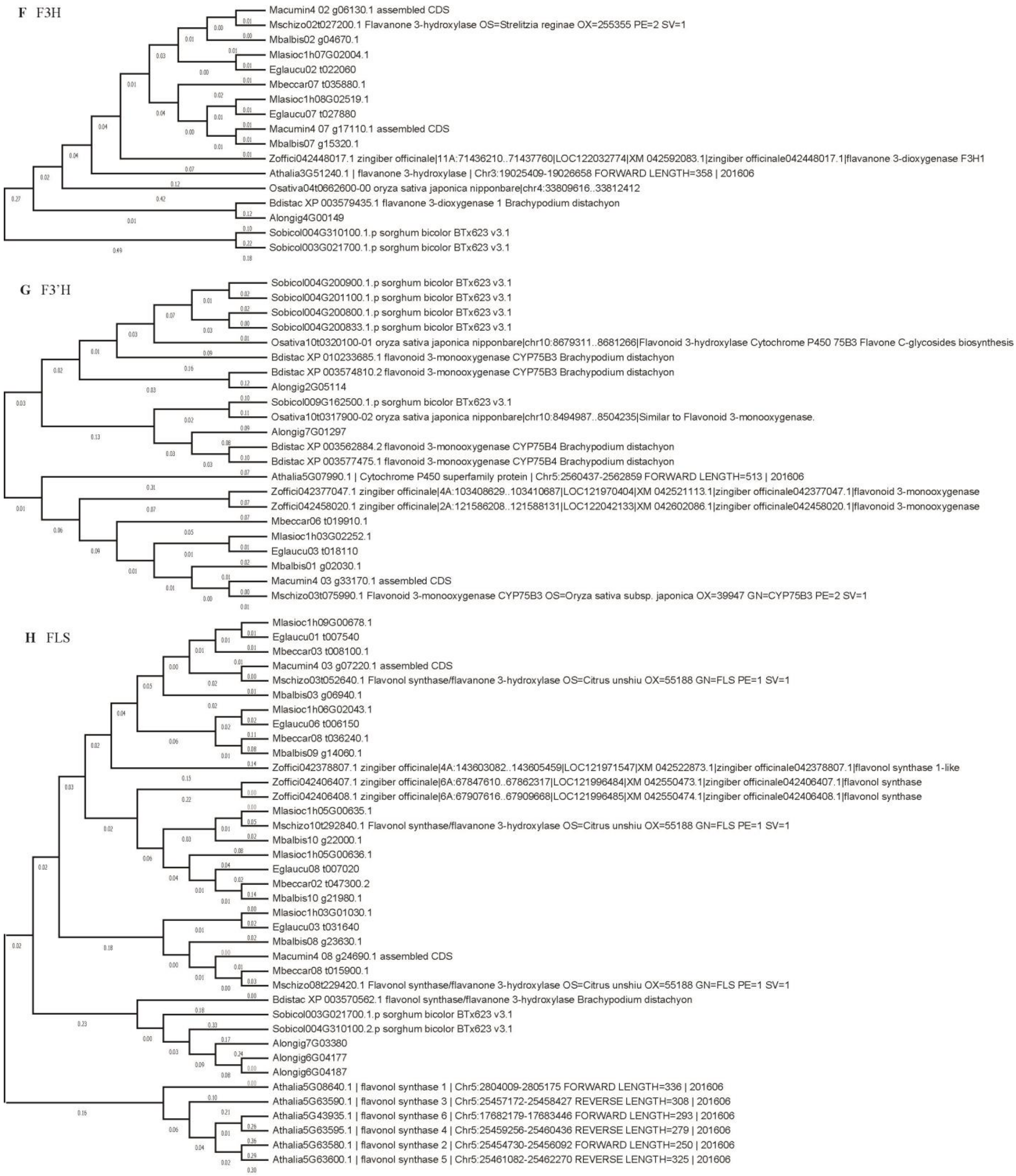
F F3H
Macumin4 02 g06130.1 assembled CDS
Mschizo02t027200.1 Flavanone 3-hydroxylase OS=Strelitzia reginae OX=255355 PE=2 SV=1
Mbalbis02 g04670.1
Mlasioc1h07G02004.1
Eglaucu02 t022060
Mbeccar07 t035880.1
Mlasioc1h08G02519.1
Eglaucu07 t027880
Macumin4 07 g17110.1 assembled CDS
Mbalbis07 g15320.1
Zoffici042448017.1 zingiber officinale|11A:71436210..71437760|LOC122032774|XM 042592083.1|zingiber officinale042448017.1|flavanone 3-dioxygenase F3H1
Athalia3G51240.1 | flavanone 3-hydroxylase | Chr3:19025409-19026658 FORWARD LENGTH=358 | 201606
Osativa04t0662600-00 oryza sativa japonica nipponbare|chr4:33809616..33812412
Bdistac XP 003579435.1 flavanone 3-dioxygenase 1 Brachypodium distachyon
Alongig4G00149
Sobicol004G310100.1.p sorghum bicolor BTx623 v3.1
Sobicol003G021700.1.p sorghum bicolor BTx623 v3.1
G F3'H
Sobicol004G200900.1.p sorghum bicolor BTx623 v3.1
Sobicol004G201100.1.p sorghum bicolor BTx623 v3.1
Sobicol004G200800.1.p sorghum bicolor BTx623 v3.1
Sobicol004G200833.1.p sorghum bicolor BTx623 v3.1
Osativa10t0320100-01 oryza sativa japonica nipponbare|chr10:8679311..8681266|Flavonoid 3-hydroxylase Cytochrome P450 75B3 Flavone C-glycosides biosynthesis
Bdistac XP 010233685.1 flavonoid 3-monooxygenase CYP75B3 Brachypodium distachyon
Bdistac XP 003574810.2 flavonoid 3-monooxygenase CYP75B3 Brachypodium distachyon
Alongig2G05114
Sobicol009G162500.1.p sorghum bicolor BTx623 v3.1
Osativa10t0317900-02 oryza sativa japonica nipponbare|chr10:8494987..8504235|Similar to Flavonoid 3-monooxygenase.
Alongig7G01297
Bdistac XP 003562884.2 flavonoid 3-monooxygenase CYP75B4 Brachypodium distachyon
Bdistac XP 003577475.1 flavonoid 3-monooxygenase CYP75B4 Brachypodium distachyon
Athalia5G07990.1 | Cytochrome P450 superfamily protein | Chr5:2560437-2562859 FORWARD LENGTH=513 | 201606
Zoffici042377047.1 zingiber officinale|4A:103408629..103410687|LOC121970404|XM 042521113.1|zingiber officinale042377047.1|flavonoid 3-monooxygenase
Zoffici042458020.1 zingiber officinale|2A:121586208..121588131|LOC122042133|XM 042602086.1|zingiber officinale042458020.1|flavonoid 3-monooxygenase
Mbeccar06 t019910.1
Mlasioc1h03G02252.1
Eglaucu03 t018110
Mbalbis01 g02030.1
Macumin4 03 g33170.1 assembled CDS
Mschizo03t075990.1 Flavonoid 3-monooxygenase CYP75B3 OS=Oryza sativa subsp. japonica OX=39947 GN=CYP75B3 PE=2 SV=1
H FLS
Mlasioc1h09G00678.1
Eglaucu01 t007540
Mbeccar03 t008100.1
Macumin4 03 g07220.1 assembled CDS
Mschizo03t052640.1 Flavonol synthase/flavanone 3-hydroxylase OS=Citrus unshiu OX=55188 GN=FLS PE=1 SV=1
Mbalbis03 g06940.1
Mlasioc1h06G02043.1
Eglaucu06 t006150
Mbeccar08 t036240.1
Mbalbis09 g14060.1
Zoffici042378807.1 zingiber officinale|4A:143603082..143605459|LOC121971547|XM 042522873.1|zingiber officinale042378807.1|flavonol synthase 1-like
Zoffici042406407.1 zingiber officinale|6A:67847610..67862317|LOC121996484|XM 042550473.1|zingiber officinale042406407.1|flavonol synthase
Zoffici042406408.1 zingiber officinale|6A:67907616..67909668|LOC121996485|XM 042550474.1|zingiber officinale042406408.1|flavonol synthase
Mlasioc1h05G00635.1
Mschizo10t292840.1 Flavonol synthase/flavanone 3-hydroxylase OS=Citrus unshiu OX=55188 GN=FLS PE=1 SV=1
Mbalbis10 g22000.1
Mlasioc1h05G00636.1
Eglaucu08 t007020
Mbeccar02 t047300.2
Mbalbis10 g21980.1
Mlasioc1h03G01030.1
Eglaucu03 t031640
Mbalbis08 g23630.1
Macumin4 08 g24690.1 assembled CDS
Mbeccar08 t015900.1
Mschizo08t229420.1 Flavonol synthase/flavanone 3-hydroxylase OS=Citrus unshiu OX=55188 GN=FLS PE=1 SV=1
Bdistac XP 003570562.1 flavonol synthase/flavanone 3-hydroxylase Brachypodium distachyon
Sobicol003G021700.1.p sorghum bicolor BTx623 v3.1
Sobicol004G310100.2.p sorghum bicolor BTx623 v3.1
Alongig7G03380
Alongig6G04177
Alongig6G04187
Athalia5G08640.1 | flavonol synthase 1 | Chr5:2804009-2805175 FORWARD LENGTH=336 | 201606
Athalia5G63590.1 | flavonol synthase 3 | Chr5:25457172-25458427 REVERSE LENGTH=308 | 201606
Athalia5G43935.1 | flavonol synthase 6 | Chr5:17682179-17683446 FORWARD LENGTH=293 | 201606
Athalia5G63595.1 | flavonol synthase 4 | Chr5:25459256-25460436 REVERSE LENGTH=279 | 201606
Athalia5G63580.1 | flavonol synthase 2 | Chr5:25454730-25456092 FORWARD LENGTH=250 | 201606
Athalia5G63600.1 | flavonol synthase 5 | Chr5:25461082-25462270 REVERSE LENGTH=325 | 201606

26

**I** F3'5'H

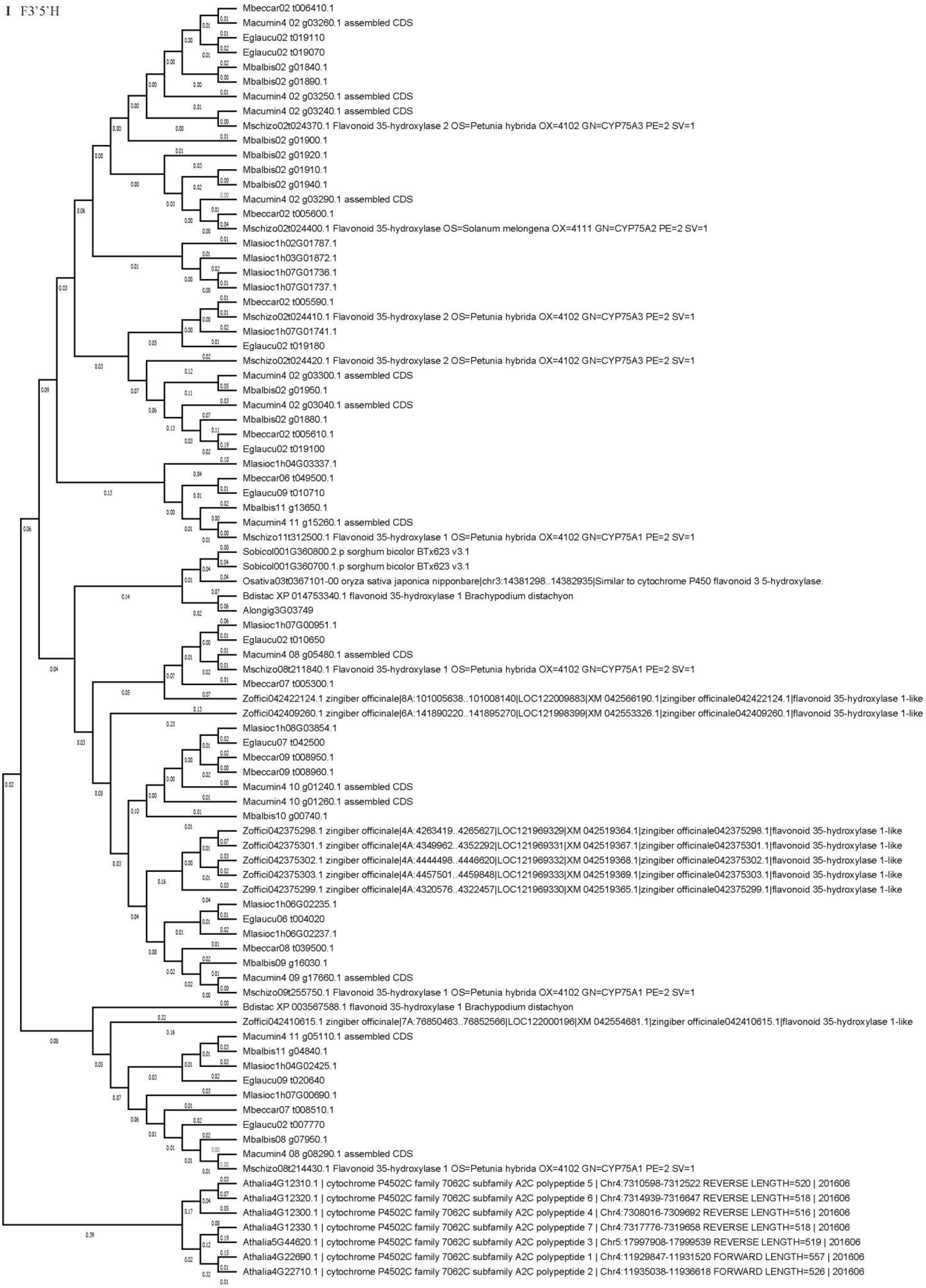
Mbeccar02 t006410.1
Macumin4 02 g03260.1 assembled CDS
Eglaucu02 t019110
Eglaucu02 t019070
Mbalbis02 g01840.1
Mbalbis02 g01890.1
Macumin4 02 g03250.1 assembled CDS
Macumin4 02 g03240.1 assembled CDS
Mschizo02t024370.1 Flavonoid 35-hydroxylase 2 OS=Petunia hybrida OX=4102 GN=CYP75A3 PE=2 SV=1
Mbalbis02 g01900.1
Mbalbis02 g01920.1
Mbalbis02 g01910.1
Mbalbis02 g01940.1
Macumin4 02 g03290.1 assembled CDS
Mbeccar02 t005600.1
Mschizo02t024400.1 Flavonoid 35-hydroxylase OS=Solanum melongena OX=4111 GN=CYP75A2 PE=2 SV=1
Mlasioc1h02G01787.1
Mlasioc1h03G01872.1
Mlasioc1h07G01736.1
Mlasioc1h07G01737.1
Mbeccar02 t005590.1
Mschizo02t024410.1 Flavonoid 35-hydroxylase 2 OS=Petunia hybrida OX=4102 GN=CYP75A3 PE=2 SV=1
Mlasioc1h07G01741.1
Eglaucu02 t019180
Mschizo02t024420.1 Flavonoid 35-hydroxylase 2 OS=Petunia hybrida OX=4102 GN=CYP75A3 PE=2 SV=1
Macumin4 02 g03300.1 assembled CDS
Mbalbis02 g01950.1
Macumin4 02 g03040.1 assembled CDS
Mbalbis02 g01880.1
Mbeccar02 t005610.1
Eglaucu02 t019100
Mlasioc1h04G03337.1
Mbeccar06 t049500.1
Eglaucu09 t010710
Mbalbis11 g13650.1
Macumin4 11 g15260.1 assembled CDS
Mschizo11t312500.1 Flavonoid 35-hydroxylase 1 OS=Petunia hybrida OX=4102 GN=CYP75A1 PE=2 SV=1
Sobicol001G360800.2.p sorghum bicolor BTx623 v3.1
Sobicol001G360700.1.p sorghum bicolor BTx623 v3.1
Osativa03t0367101-00 oryza sativa japonica nipponbare|chr3:14381298..14382935|Similar to cytochrome P450 flavonoid 3 5-hydroxylase.
Bdistac XP 014753340.1 flavonoid 35-hydroxylase 1 Brachypodium distachyon
Alongig3G03749
Mlasioc1h07G00951.1
Eglaucu02 t010650
Macumin4 08 g05480.1 assembled CDS
Mschizo08t211840.1 Flavonoid 35-hydroxylase 1 OS=Petunia hybrida OX=4102 GN=CYP75A1 PE=2 SV=1
Mbeccar07 t005300.1
Zoffici042422124.1 zingiber officinale|8A:101005638..101008140|LOC122009883|XM 042566190.1|zingiber officinale042422124.1|flavonoid 35-hydroxylase 1-like
Zoffici042409260.1 zingiber officinale|6A:141890220..141895270|LOC121998399|XM 042553326.1|zingiber officinale042409260.1|flavonoid 35-hydroxylase 1-like
Mlasioc1h08G03854.1
Eglaucu07 t042500
Mbeccar09 t008950.1
Mbeccar09 t008960.1
Macumin4 10 g01240.1 assembled CDS
Macumin4 10 g01260.1 assembled CDS
Mbalbis10 g00740.1
Zoffici042375298.1 zingiber officinale|4A:4263419..4265627|LOC121969329|XM 042519364.1|zingiber officinale042375298.1|flavonoid 35-hydroxylase 1-like
Zoffici042375301.1 zingiber officinale|4A:4349962..4352292|LOC121969331|XM 042519367.1|zingiber officinale042375301.1|flavonoid 35-hydroxylase 1-like
Zoffici042375302.1 zingiber officinale|4A:4444498..4446620|LOC121969332|XM 042519368.1|zingiber officinale042375302.1|flavonoid 35-hydroxylase 1-like
Zoffici042375303.1 zingiber officinale|4A:4457501..4459848|LOC121969333|XM 042519369.1|zingiber officinale042375303.1|flavonoid 35-hydroxylase 1-like
Zoffici042375299.1 zingiber officinale|4A:4320576..4322457|LOC121969330|XM 042519365.1|zingiber officinale042375299.1|flavonoid 35-hydroxylase 1-like
Mlasioc1h06G02235.1
Eglaucu06 t004020
Mlasioc1h06G02237.1
Mbeccar08 t039500.1
Mbalbis09 g16030.1
Macumin4 09 g17660.1 assembled CDS
Mschizo09t255750.1 Flavonoid 35-hydroxylase 1 OS=Petunia hybrida OX=4102 GN=CYP75A1 PE=2 SV=1
Bdistac XP 003567588.1 flavonoid 35-hydroxylase 1 Brachypodium distachyon
Zoffici042410615.1 zingiber officinale|7A:76850463..76852566|LOC122000196|XM 042554681.1|zingiber officinale042410615.1|flavonoid 35-hydroxylase 1-like
Macumin4 11 g05110.1 assembled CDS
Mbalbis11 g04840.1
Mlasioc1h04G02425.1
Eglaucu09 t020640
Mlasioc1h07G00690.1
Mbeccar07 t008510.1
Eglaucu02 t007770
Mbalbis08 g07950.1
Macumin4 08 g08290.1 assembled CDS
Mschizo08t214430.1 Flavonoid 35-hydroxylase 1 OS=Petunia hybrida OX=4102 GN=CYP75A1 PE=2 SV=1
Athalia4G12310.1 | cytochrome P4502C family 7062C subfamily A2C polypeptide 5 | Chr4:7310598-7312522 REVERSE LENGTH=520 | 201606
Athalia4G12320.1 | cytochrome P4502C family 7062C subfamily A2C polypeptide 6 | Chr4:7314939-7316647 REVERSE LENGTH=518 | 201606
Athalia4G12300.1 | cytochrome P4502C family 7062C subfamily A2C polypeptide 4 | Chr4:7308016-7309692 REVERSE LENGTH=516 | 201606
Athalia4G12330.1 | cytochrome P4502C family 7062C subfamily A2C polypeptide 7 | Chr4:7317776-7319658 REVERSE LENGTH=518 | 201606
Athalia5G44620.1 | cytochrome P4502C family 7062C subfamily A2C polypeptide 3 | Chr5:17997908-17999539 REVERSE LENGTH=519 | 201606
Athalia4G22690.1 | cytochrome P4502C family 7062C subfamily A2C polypeptide 1 | Chr4:11929847-11931520 FORWARD LENGTH=557 | 201606
Athalia4G22710.1 | cytochrome P4502C family 7062C subfamily A2C polypeptide 2 | Chr4:11935038-11936618 FORWARD LENGTH=526 | 201606

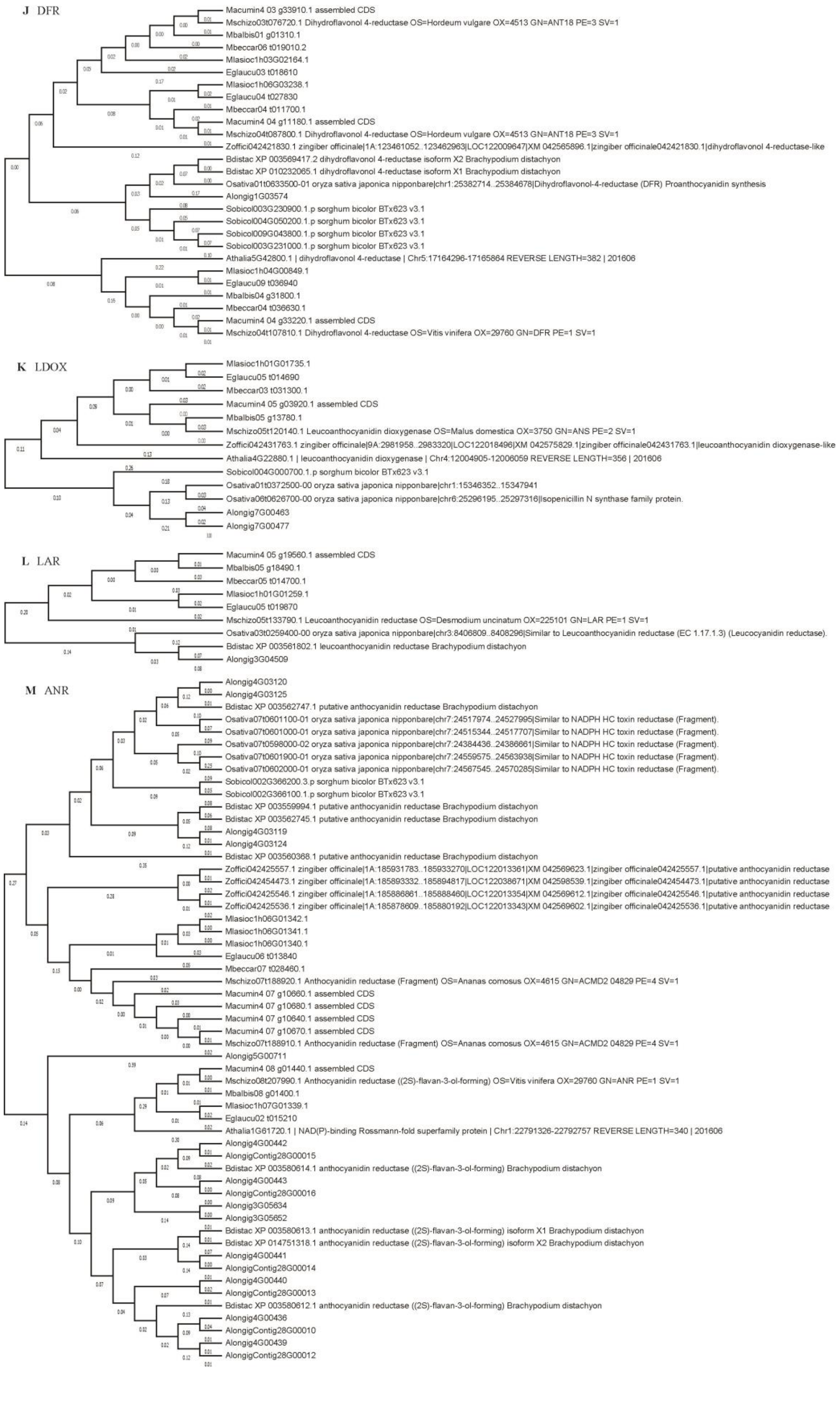
J DFR
Macumin4 03 g33910.1 assembled CDS
Mschizo03t076720.1 Dihydroflavonol 4-reductase OS=Hordeum vulgare OX=4513 GN=ANT18 PE=3 SV=1
Mbalbis01 g01310.1
Mbeccar06 t019010.2
Mlasioc1h03G02164.1
Eglaucu03 t018610
Mlasioc1h06G03238.1
Eglaucu04 t027830
Mbeccar04 t011700.1
Macumin4 04 g11180.1 assembled CDS
Mschizo04t087800.1 Dihydroflavonol 4-reductase OS=Hordeum vulgare OX=4513 GN=ANT18 PE=3 SV=1
Zoffici042421830.1 zingiber officinale|1A:123461052..123462963|LOC122009647|XM 042565896.1|zingiber officinale042421830.1|dihydroflavonol 4-reductase-like
Bdistac XP 003569417.2 dihydroflavonol 4-reductase isoform X2 Brachypodium distachyon
Bdistac XP 010232065.1 dihydroflavonol 4-reductase isoform X1 Brachypodium distachyon
Osativa01t0633500-01 oryza sativa japonica nipponbare|chr1:25382714..25384678|Dihydroflavonol-4-reductase (DFR) Proanthocyanidin synthesis
Alongig1G03574
Sobicol003G230900.1.p sorghum bicolor BTx623 v3.1
Sobicol004G050200.1.p sorghum bicolor BTx623 v3.1
Sobicol009G043800.1.p sorghum bicolor BTx623 v3.1
Sobicol003G231000.1.p sorghum bicolor BTx623 v3.1
Athalia5G42800.1 | dihydroflavonol 4-reductase | Chr5:17164296-17165864 REVERSE LENGTH=382 | 201606
Mlasioc1h04G00849.1
Eglaucu09 t036940
Mbalbis04 g31800.1
Mbeccar04 t036630.1
Macumin4 04 g33220.1 assembled CDS
Mschizo04t107810.1 Dihydroflavonol 4-reductase OS=Vitis vinifera OX=29760 GN=DFR PE=1 SV=1
K LDOX
Mlasioc1h01G01735.1
Eglaucu05 t014690
Mbeccar03 t031300.1
Macumin4 05 g03920.1 assembled CDS
Mbalbis05 g13780.1
Mschizo05t120140.1 Leucoanthocyanidin dioxygenase OS=Malus domestica OX=3750 GN=ANS PE=2 SV=1
Zoffici042431763.1 zingiber officinale|9A:2981958..2983320|LOC122018496|XM 042575829.1|zingiber officinale042431763.1|leucoanthocyanidin dioxygenase-like
Athalia4G22880.1 | leucoanthocyanidin dioxygenase | Chr4:12004905-12006059 REVERSE LENGTH=356 | 201606
Sobicol004G000700.1.p sorghum bicolor BTx623 v3.1
Osativa01t0372500-00 oryza sativa japonica nipponbare|chr1:15346352..15347941
Osativa06t0626700-00 oryza sativa japonica nipponbare|chr6:25296195..25297316|Isopenicillin N synthase family protein.
Alongig7G00463
Alongig7G00477
L LAR
Macumin4 05 g19560.1 assembled CDS
Mbalbis05 g18490.1
Mbeccar05 t014700.1
Mlasioc1h01G01259.1
Eglaucu05 t019870
Mschizo05t133790.1 Leucoanthocyanidin reductase OS=Desmodium uncinatum OX=225101 GN=LAR PE=1 SV=1
Osativa03t0259400-00 oryza sativa japonica nipponbare|chr3:8406809..8408296|Similar to Leucoanthocyanidin reductase (EC 1.17.1.3) (Leucocyanidin reductase).
Bdistac XP 003561802.1 leucoanthocyanidin reductase Brachypodium distachyon
Alongig3G04509
M ANR
Alongig4G03120
Alongig4G03125
Bdistac XP 003562747.1 putative anthocyanidin reductase Brachypodium distachyon
Osativa07t0601100-01 oryza sativa japonica nipponbare|chr7:24517974..24527995|Similar to NADPH HC toxin reductase (Fragment).
Osativa07t0601000-01 oryza sativa japonica nipponbare|chr7:24515344..24517707|Similar to NADPH HC toxin reductase (Fragment).
Osativa07t0598000-02 oryza sativa japonica nipponbare|chr7:24384436..24386661|Similar to NADPH HC toxin reductase (Fragment).
Osativa07t0601900-01 oryza sativa japonica nipponbare|chr7:24559575..24563938|Similar to NADPH HC toxin reductase (Fragment).
Osativa07t0602000-01 oryza sativa japonica nipponbare|chr7:24567545..24570285|Similar to NADPH HC toxin reductase (Fragment).
Sobicol002G366200.3.p sorghum bicolor BTx623 v3.1
Sobicol002G366100.1.p sorghum bicolor BTx623 v3.1
Bdistac XP 003559994.1 putative anthocyanidin reductase Brachypodium distachyon
Bdistac XP 003562745.1 putative anthocyanidin reductase Brachypodium distachyon
Alongig4G03119
Alongig4G03124
Bdistac XP 003560368.1 putative anthocyanidin reductase Brachypodium distachyon
Zoffici042425557.1 zingiber officinale|1A:185931783..185933270|LOC122013361|XM 042569623.1|zingiber officinale042425557.1|putative anthocyanidin reductase
Zoffici042454473.1 zingiber officinale|1A:185893332..185894817|LOC122038671|XM 042598539.1|zingiber officinale042454473.1|putative anthocyanidin reductase
Zoffici042425546.1 zingiber officinale|1A:185886861..185888460|LOC122013354|XM 042569612.1|zingiber officinale042425546.1|putative anthocyanidin reductase
Zoffici042425536.1 zingiber officinale|1A:185878609..185880192|LOC122013343|XM 042569602.1|zingiber officinale042425536.1|putative anthocyanidin reductase
Mlasioc1h06G01342.1
Mlasioc1h06G01341.1
Mlasioc1h06G01340.1
Eglaucu06 t013840
Mbeccar07 t028460.1
Mschizo07t188920.1 Anthocyanidin reductase (Fragment) OS=Ananas comosus OX=4615 GN=ACMD2 04829 PE=4 SV=1
Macumin4 07 g10660.1 assembled CDS
Macumin4 07 g10680.1 assembled CDS
Macumin4 07 g10640.1 assembled CDS
Macumin4 07 g10670.1 assembled CDS
Mschizo07t188910.1 Anthocyanidin reductase (Fragment) OS=Ananas comosus OX=4615 GN=ACMD2 04829 PE=4 SV=1
Alongig5G00711
Macumin4 08 g01440.1 assembled CDS
Mschizo08t207990.1 Anthocyanidin reductase ((2S)-flavan-3-ol-forming) OS=Vitis vinifera OX=29760 GN=ANR PE=1 SV=1
Mbalbis08 g01400.1
Mlasioc1h07G01339.1
Eglaucu02 t015210
Athalia1G61720.1 | NAD(P)-binding Rossmann-fold superfamily protein | Chr1:22791326-22792757 REVERSE LENGTH=340 | 201606
Alongig4G00442
AlongigContig28G00015
Bdistac XP 003580614.1 anthocyanidin reductase ((2S)-flavan-3-ol-forming) Brachypodium distachyon
Alongig4G00443
AlongigContig28G00016
Alongig3G05634
Alongig3G05652
Bdistac XP 003580613.1 anthocyanidin reductase ((2S)-flavan-3-ol-forming) isoform X1 Brachypodium distachyon
Bdistac XP 014751318.1 anthocyanidin reductase ((2S)-flavan-3-ol-forming) isoform X2 Brachypodium distachyon
Alongig4G00441
AlongigContig28G00014
Alongig4G00440
AlongigContig28G00013
Bdistac XP 003580612.1 anthocyanidin reductase ((2S)-flavan-3-ol-forming) Brachypodium distachyon
Alongig4G00436
AlongigContig28G00010
Alongig4G00439
AlongigContig28G00012

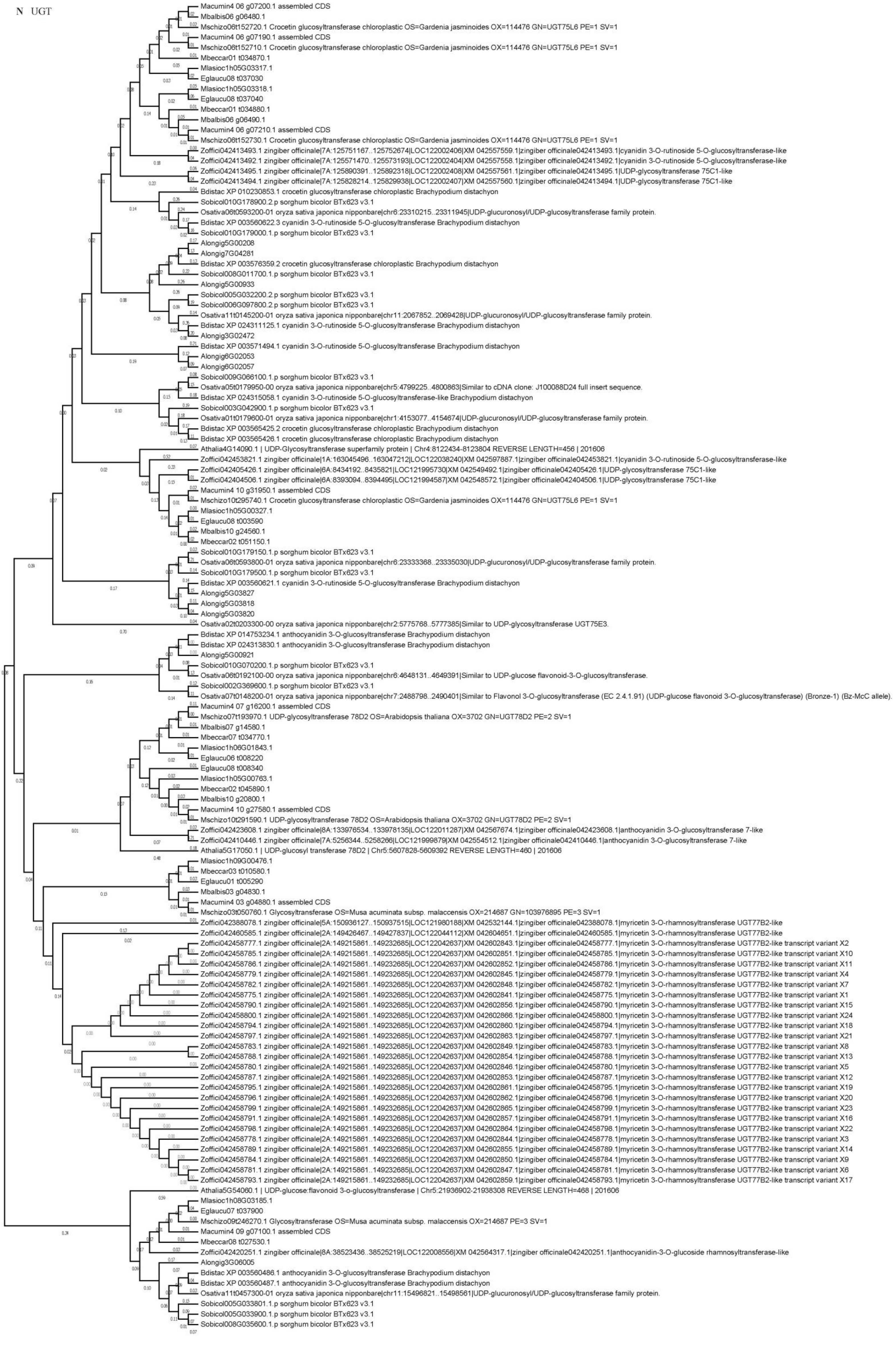


Flavonoid biosynthetic genes in Musaceae.

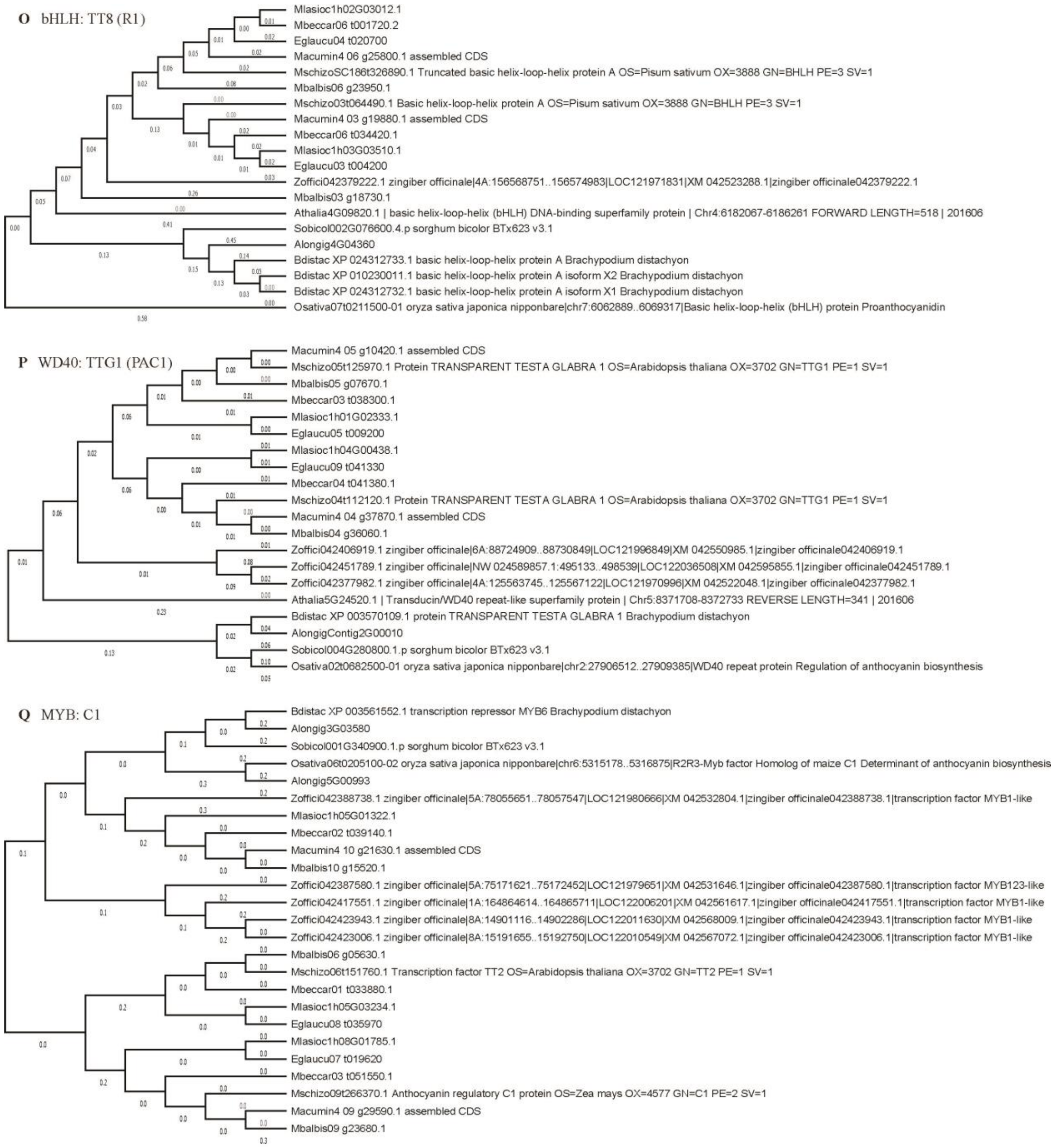


**Figure S2 Phylogenetic trees shows the genetic relationship of flavonoid biosynthetic genes in *Arabidopsis*, *B. distachyon*, *A. longiglumis*, *O. sativa*, *S. bicolor*, *Z. officinale*, *E. glaucum*, *M. lasiocarpa*, *M. beccarii*, *M. acuminata*, *M. balbisiana*, and *M. schizocarpa*.** (**A**) PAL family. (**B**) 4CL family. (**C**) C4H family. (**D**) CHI family. (**E**) CHS family. (**F**) F3H family. (**G**) F3'H family. (**H**) FLS family. (**I**) F3'5'H family. (**J**) DFR family. (**K**) LDOX family. (**L**) LAR family. (**M**) ANR family. (**N**) UGT family. (**O**) bHLH: TT8 (R1) family. (**P**) WD40: TTG1 (PAC1) family. (**Q**) MYB: C1 family.

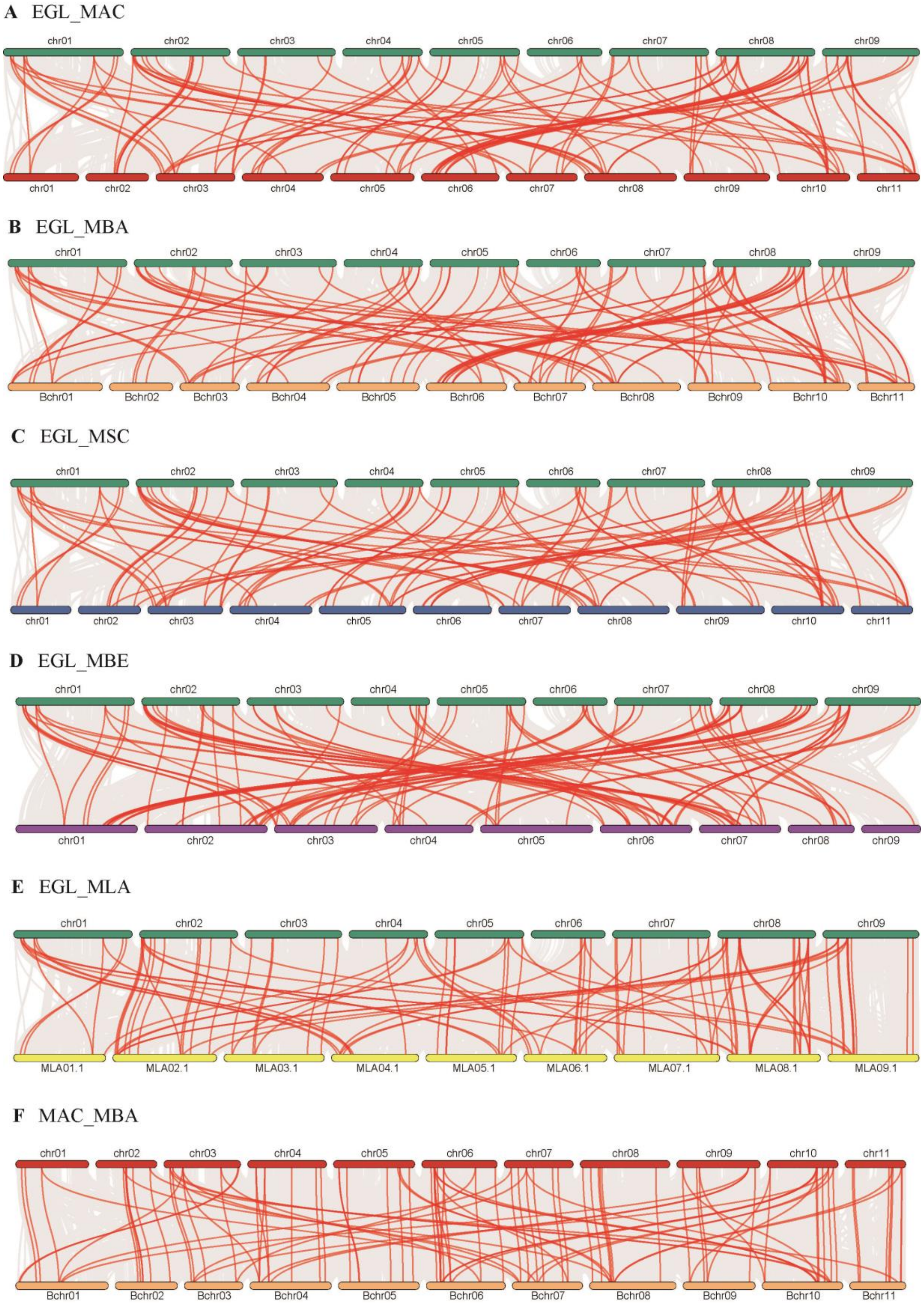
A EGL_MAC
chr01
chr02
chr03
chr04
chr05
chr06
chr07
chr08
chr09
chr01
chr02
chr03
chr04
chr05
chr06
chr07
chr08
chr09
chr10
chr11
B EGL_MBA
chr01
chr02
chr03
chr04
chr05
chr06
chr07
chr08
chr09
Bchr01
Bchr02
Bchr03
Bchr04
Bchr05
Bchr06
Bchr07
Bchr08
Bchr09
Bchr10
Bchr11
C EGL_MSC
chr01
chr02
chr03
chr04
chr05
chr06
chr07
chr08
chr09
chr01
chr02
chr03
chr04
chr05
chr06
chr07
chr08
chr09
chr10
chr11
D EGL_MBE
chr01
chr02
chr03
chr04
chr05
chr06
chr07
chr08
chr09
chr01
chr02
chr03
chr04
chr05
chr06
chr07
chr08
chr09
E EGL_MLA
chr01
chr02
chr03
chr04
chr05
chr06
chr07
chr08
chr09
MLA01.1
MLA02.1
MLA03.1
MLA04.1
MLA05.1
MLA06.1
MLA07.1
MLA08.1
MLA09.1
F MAC_MBA
chr01
chr02
chr03
chr04
chr05
chr06
chr07
chr08
chr09
chr10
chr11
Bchr01
Bchr02
Bchr03
Bchr04
Bchr05
Bchr06
Bchr07
Bchr08
Bchr09
Bchr10
Bchr11

38

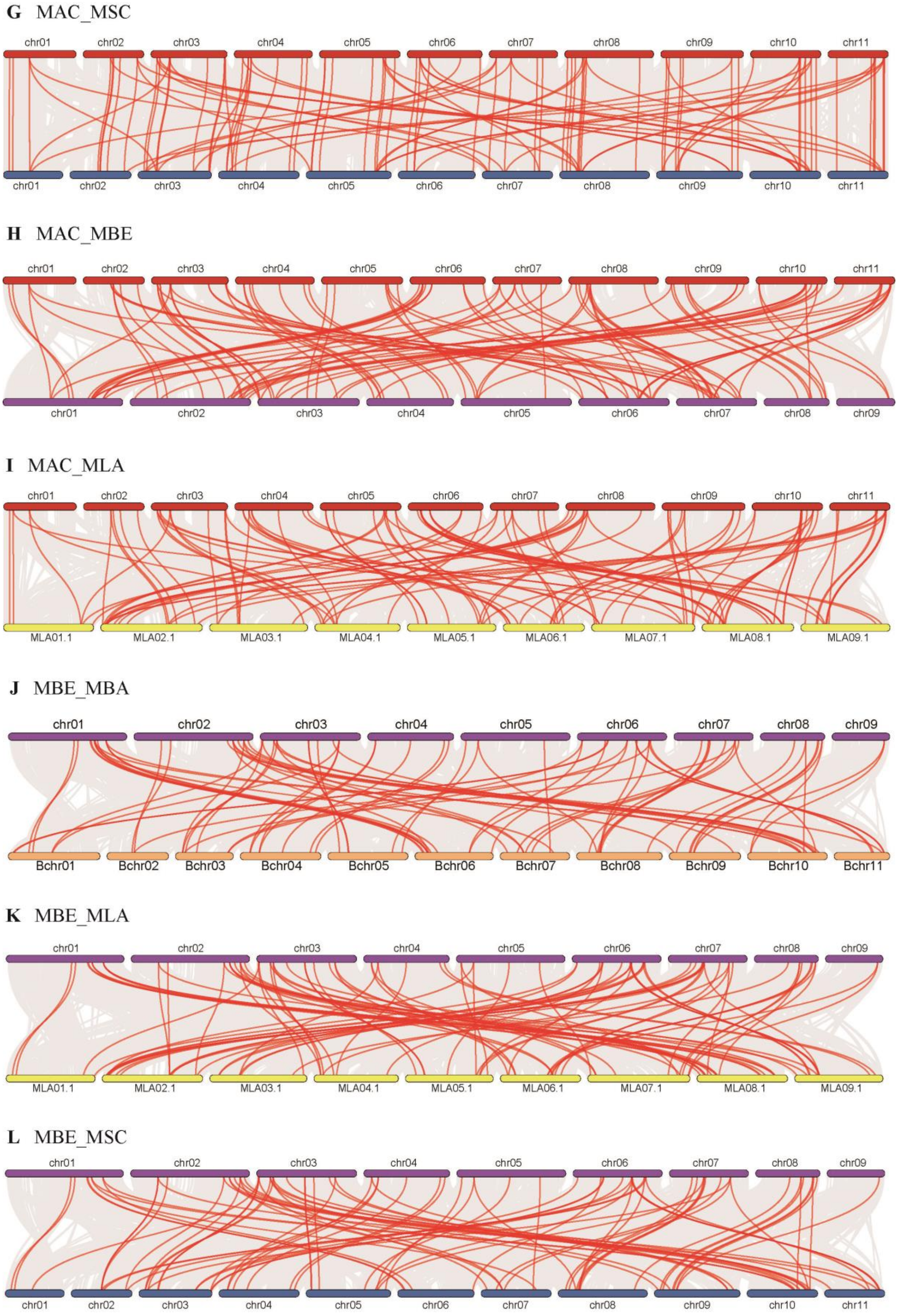
G MAC_MSC
chr01 chr02 chr03 chr04 chr05 chr06 chr07 chr08 chr09 chr10 chr11
chr01 chr02 chr03 chr04 chr05 chr06 chr07 chr08 chr09 chr10 chr11
H MAC_MBE
chr01 chr02 chr03 chr04 chr05 chr06 chr07 chr08 chr09 chr10 chr11
chr01 chr02 chr03 chr04 chr05 chr06 chr07 chr08 chr09
I MAC_MLA
chr01 chr02 chr03 chr04 chr05 chr06 chr07 chr08 chr09 chr10 chr11
MLA01.1 MLA02.1 MLA03.1 MLA04.1 MLA05.1 MLA06.1 MLA07.1 MLA08.1 MLA09.1
J MBE_MBA
chr01 chr02 chr03 chr04 chr05 chr06 chr07 chr08 chr09
Bchr01 Bchr02 Bchr03 Bchr04 Bchr05 Bchr06 Bchr07 Bchr08 Bchr09 Bchr10 Bchr11
K MBE_MLA
chr01 chr02 chr03 chr04 chr05 chr06 chr07 chr08 chr09
MLA01.1 MLA02.1 MLA03.1 MLA04.1 MLA05.1 MLA06.1 MLA07.1 MLA08.1 MLA09.1
L MBE_MSC
chr01 chr02 chr03 chr04 chr05 chr06 chr07 chr08 chr09
chr01 chr02 chr03 chr04 chr05 chr06 chr07 chr08 chr09 chr10 chr11

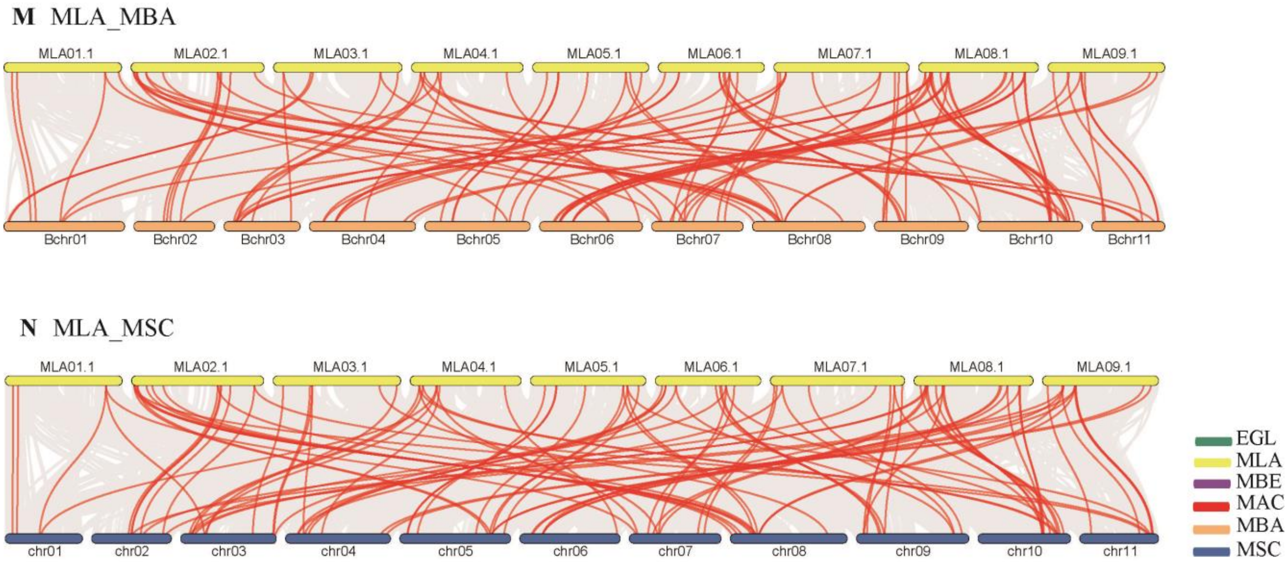


**Figure S3. Synteny analysis of FBGs between six Musaceae species.** Gray lines in the background indicate the collinear blocks within Musaceae genomes, while the red lines highlight the syntenic FBG pairs. The specie names “EGL”, “MLA”, “MBE”, “MAC”, “MBA”, and “MSC” indicate *E. glaucum*, *M. lasiocarpa*, *M. beccarii*, *M. acuminata*, *M. balbisiana*, and *M. schizocarpa*. (**A**) *E. glaucum* with *M. acuminata*. (**B**) *E. glaucum* with *M. balbisiana*. (**C**) *E. glaucum* with *M. schizocarpa*. (**D**) *E. glaucum* with *M. beccarii*. (**E**) *E. glaucum* with *M. lasiocarpa*. (**F**) *M. acuminata* with *M. balbisiana*. (**G**) *M. acuminata* with *M. schizocarpa*. (**H**) *M. acuminata* with *M. beccarii.* (**I**) *M. acuminata.* with *M. lasiocarpa* (**J**) *M. beccarii* with *M. balbisiana* (**K**) *M. beccarii* with *M. lasiocarpa.* (**L**) *M. beccarii* with *M. schizocarpa* (**M**) *M. lasiocarpa* with *M. balbisiana* (**N**) *M. lasiocarpa* with *M. schizocarp*.

**A**

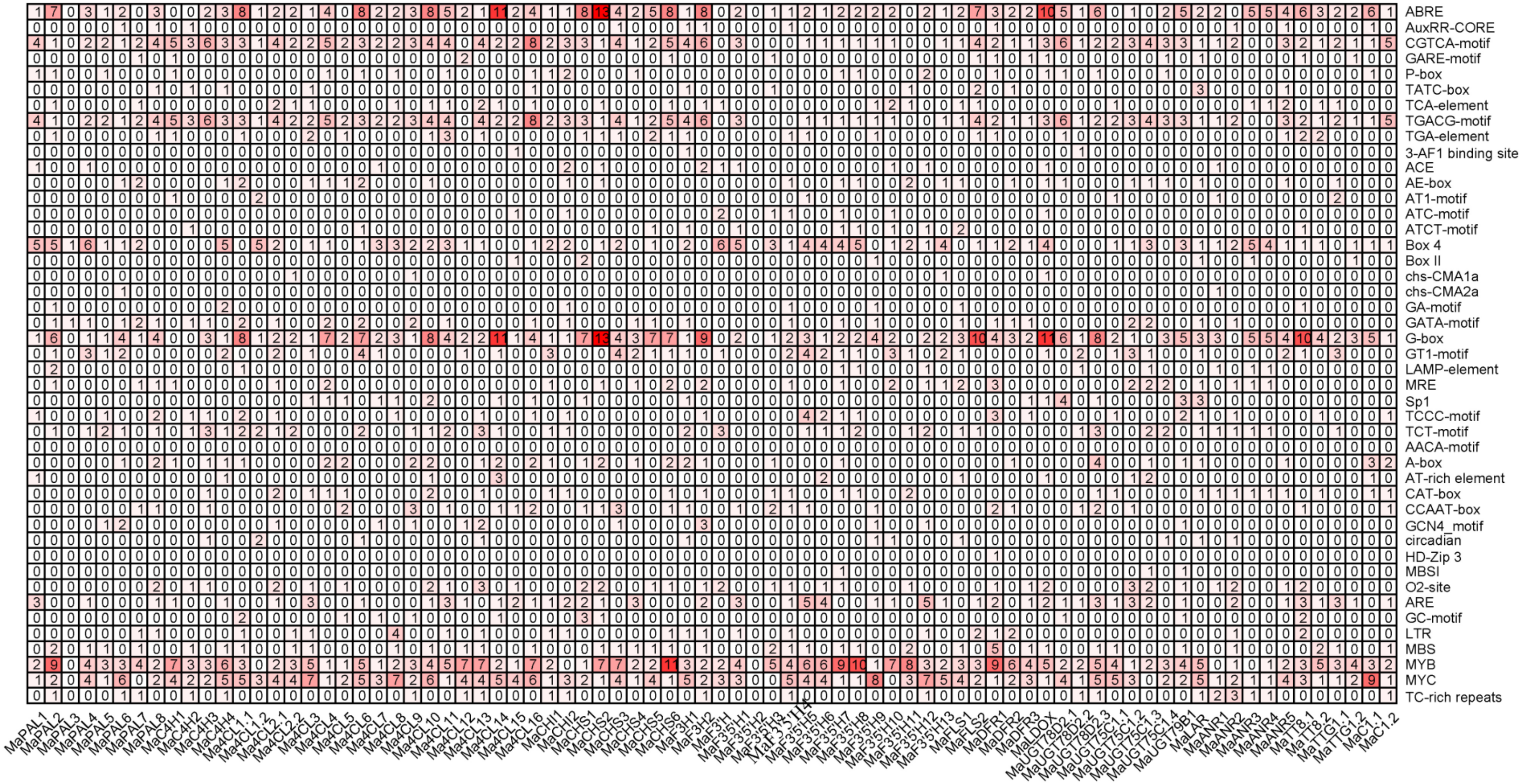


Flavonoid biosynthetic genes in Musaceae.

**B**

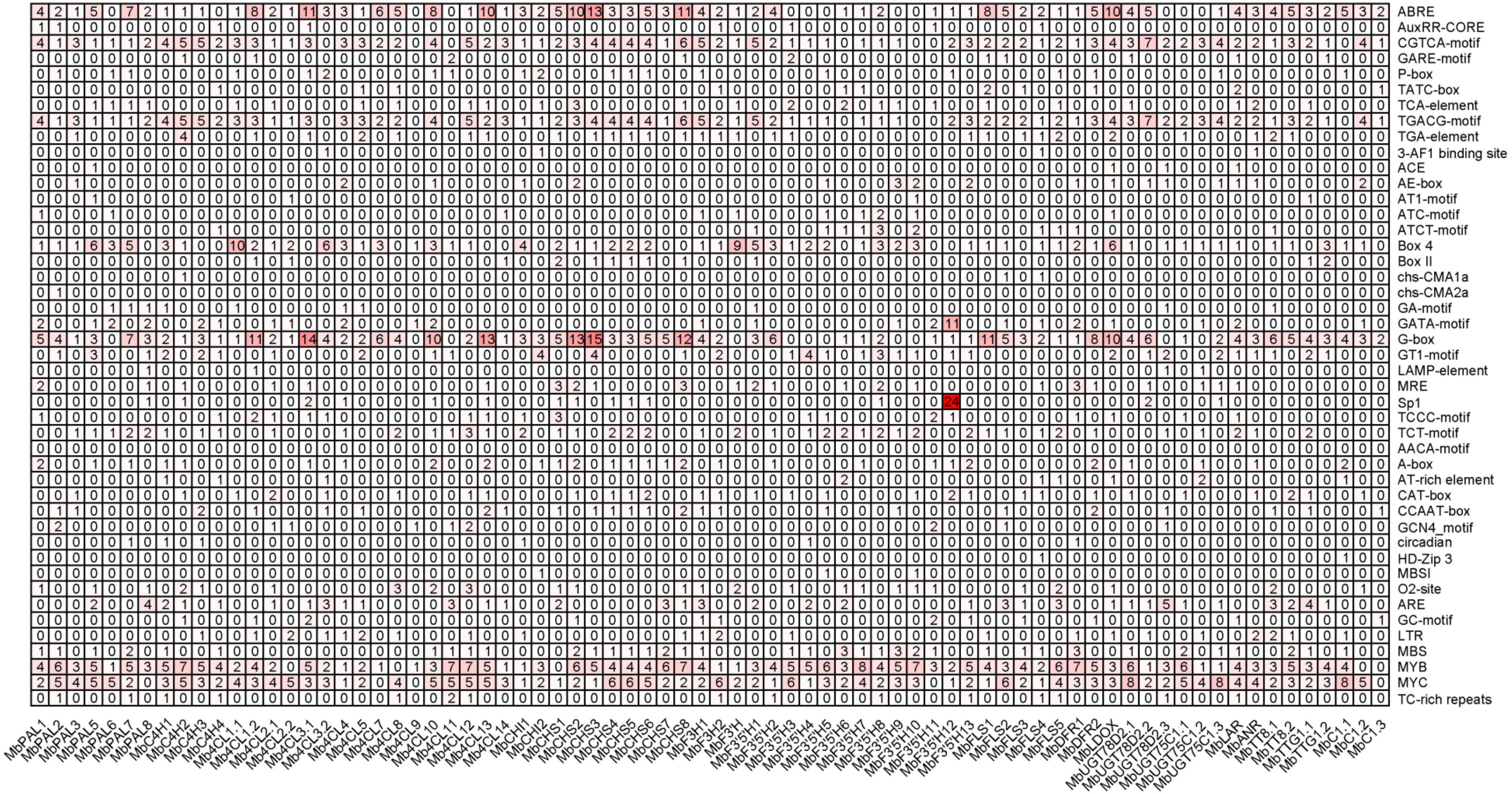


Flavonoid biosynthetic genes in Musaceae.

**C**

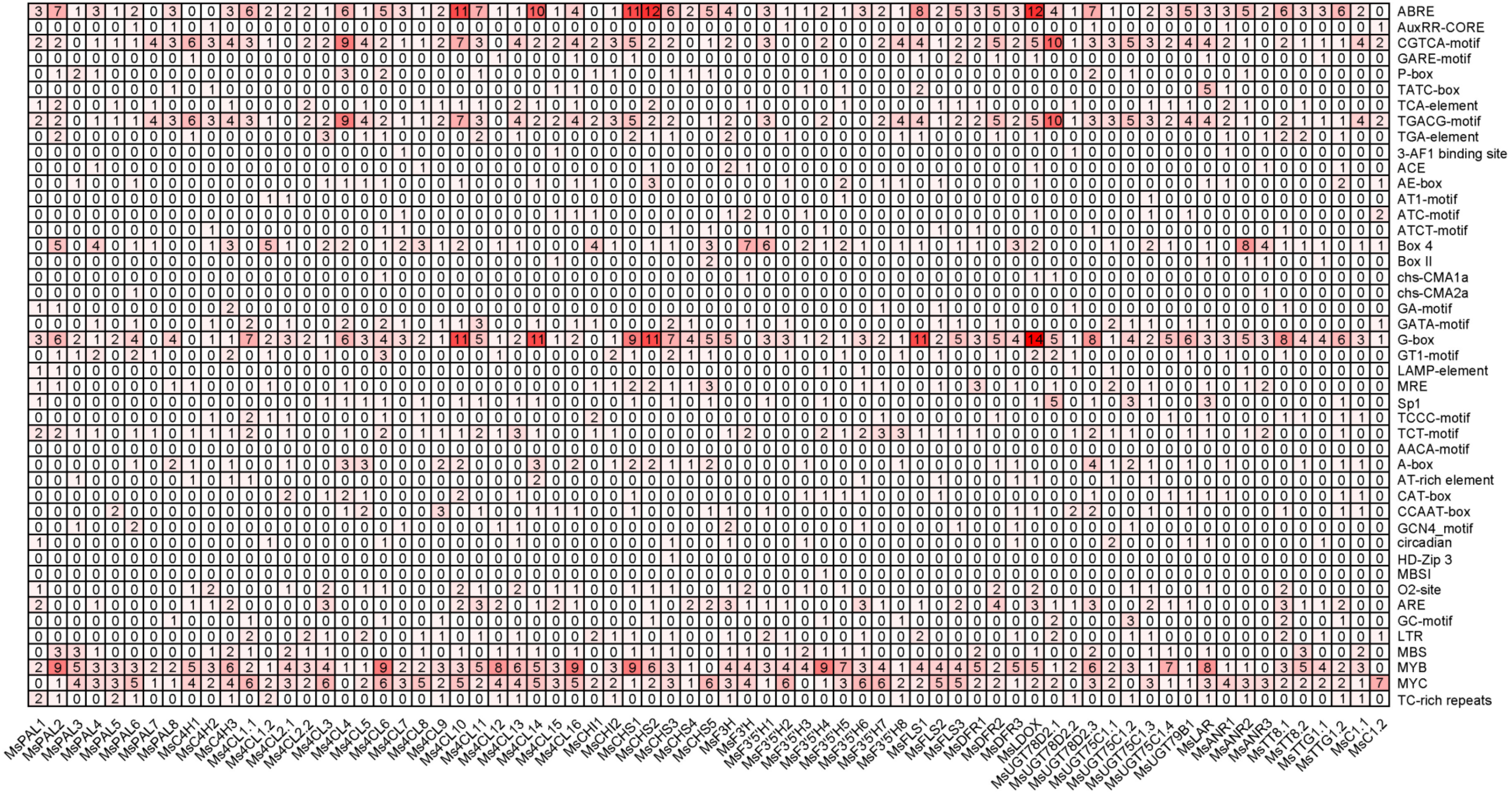


Flavonoid biosynthetic genes in Musaceae.

D

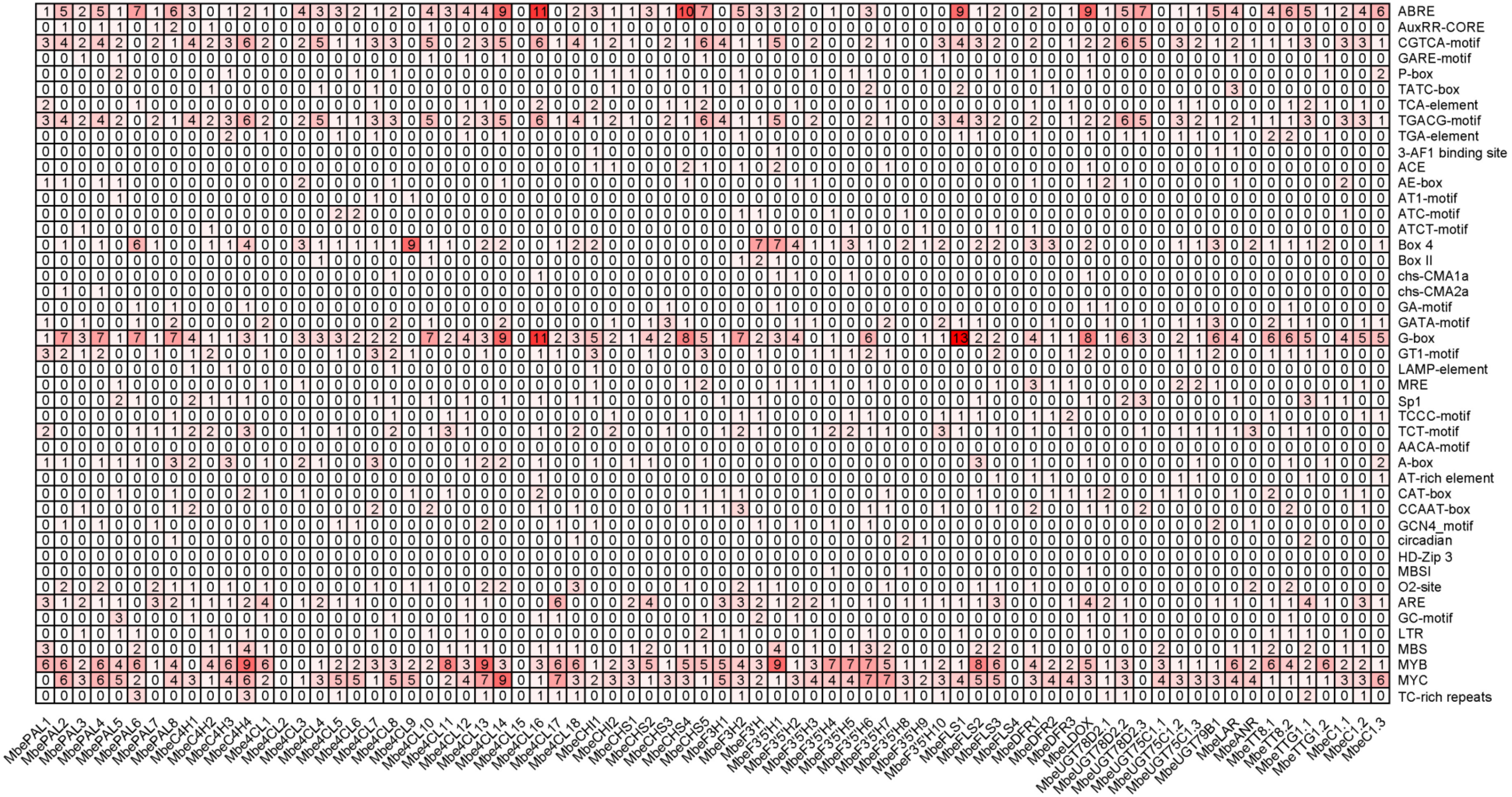


Flavonoid biosynthetic genes in Musaceae.

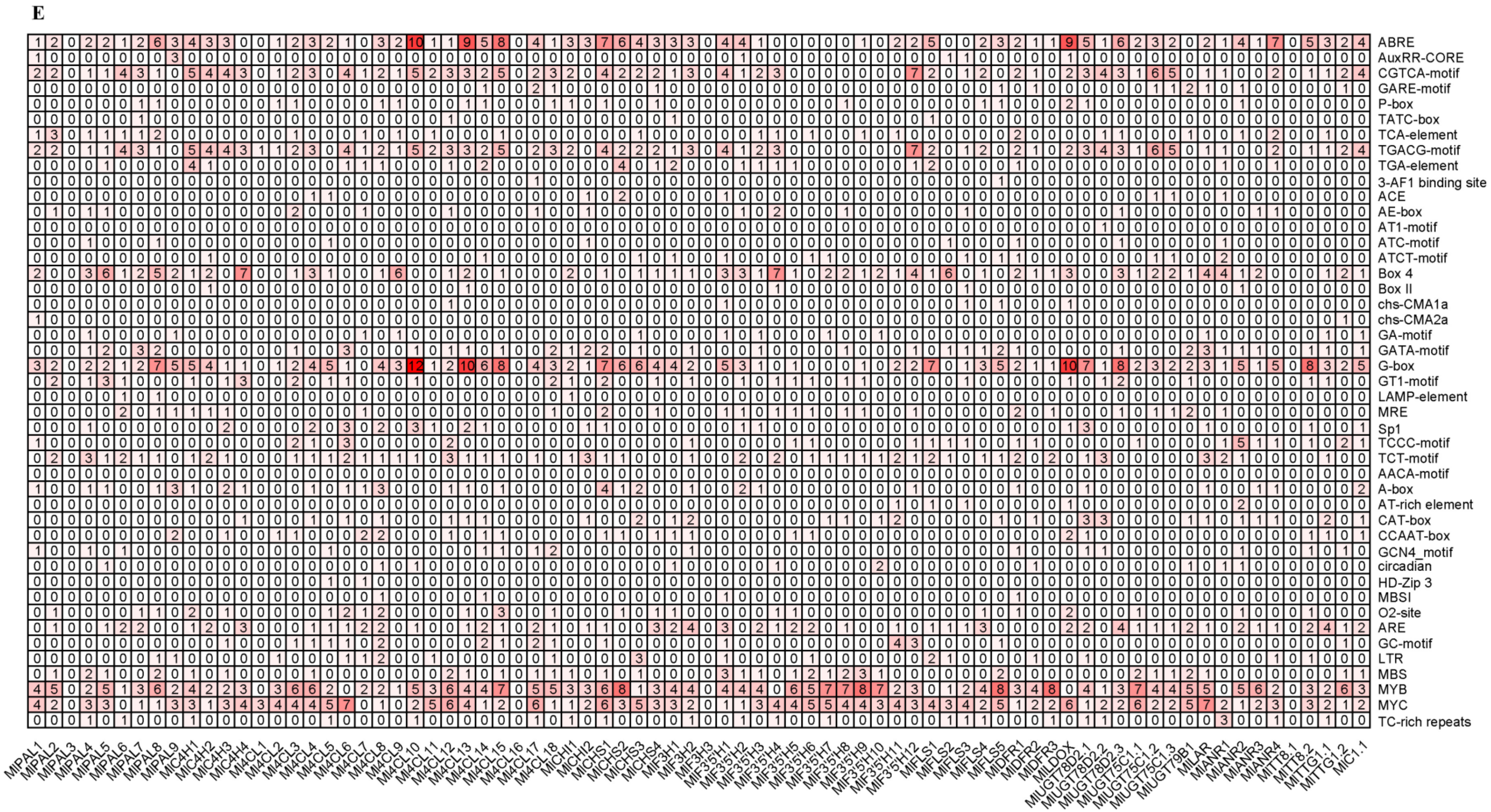


Flavonoid biosynthetic genes in Musaceae.

F

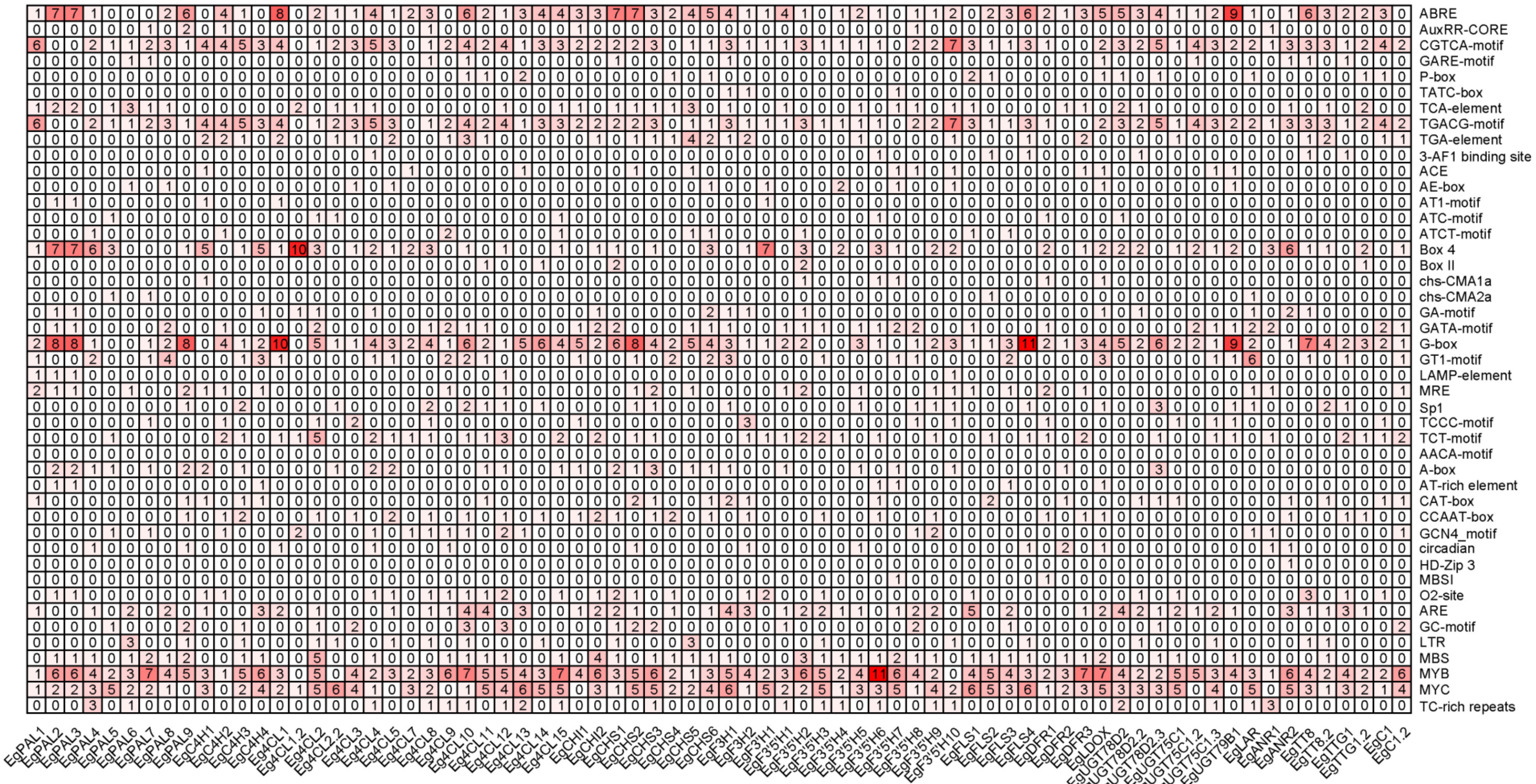


**Figure S4. Cis-acting elements in flavonoid biosynthetic genes of Musaceae.** **(A)** *Musa acuminata* (*x*=11). **(B)** *M. balbisiana* (*x*=11). **(C)** *M. schizocarpa* (*x*=11). **(D)** *M. beccarii* (*x*=9). **(E)** *Musella lasiocarpa* (*x*=9). **(F)** *Ensete glaucum* (*x*=9)

Flavonoid biosynthetic genes in Musaceae.
