## Supplemental Table 1-4 for "Genome-wide analysis of flavonoid biosynthetic genes in Musaceae (*Ensete*, *Musella*, and *Musa* species) reveals amplification of flavonoid 3’,5’-hydroxylase"

**Additional file 2:**

**Table S1.** Flavonoid biosynthetic genes (FBGs) in *A. thaliana*, *B. distachyon*, *O. sativa*, *S. bicolor*, *Z. officinale*, *M. beccarii*, *M. lasiocarpa*, *E. glaucum*, *M. acuminata*, *M. balbisiana* and *M. schizocharpa*.

**Table S2.** The information of flavonoid biosynthetic genes identified in Musaceae.

**Table S3.** Divergence time, Ka, Ks, and Ka/Ks values of orthologous flavonoid biosynthetic genes pairs between Musaceae.

**Table S4.** Duplication pairs intra Musaceae species.

**Table S1.** Flavonoid biosynthetic genes (FBGs) in *A. thaliana*, *B. distachyon*, *O. sativa*, *S. bicolor*, *Z. officinale*, *M. beccarii*, *M. lasiocarpa*, *E. glaucum*, *M. acuminata*, *M. balbisiana*, and *M. schizocharpa*.

| | ***A. thaliana*** | ***B. distachyon*** | ***A. longiglumis*** | ***O. sativa*** | ***S. bicolor*** | ***Z. officinale*** | ***E. glaucum*** | ***M. lasiocarpa*** | ***M. beccarii*** | ***M. acuminata*** | ***M. balbisiana*** | ***M. schizocharpa*** |
|---|---|---|---|---|---|---|---|---|---|---|---|---|
| **PAL** | AT2G37040 | XP_003575396.1 | AL6G04288 | Os04t0518100-01 | Sobic.006G148800.1.p | zingiber_officinale042419393.1 | Eg01_t040920 | ML1h04G02672.1 | Mbe05_t009880.2 | Macma4_01_g04800.1 | Mba09_g14610 | Ms01t004140 |
| | AT3G53260 | XP_003575400.1 | AL1G02130 | Os02t0626100-02 | Sobic.004G220300.1.p | zingiber_officinale042419394.1 | Eg05_t014880 | ML1h01G00856.1 | Mbe06_t040740.1 | Macma4_05_g03790.1 | Mba11_g20430 | Ms05t120010 |
| | AT5G04230 | XP_003580144.1 | AL3G02155 | Os11t0708900-01 | Sobic.001G160500.1.p | zingiber_officinale042389150.1 | Eg05_t014870 | ML1h07G00086.1 | Mbe07_t015550.1 | Macma4_11_g15930.1 | Mba08_g14430 | Ms11t313180 |
| | AT3G10340 | XP_003575404.1 | AL6G03333 | Os12t0520200-01 | Sobic.006G148900.1.p | zingiber_officinale042378355.1 | Eg02_t018790 | ML1h04G03282.1 | Mbe06_t040740.2 | Macma4_02_g02760.1 | Mba02_g01510 | Ms02t024050 |
| | | XP_003575365.1 | AL6G03335 | Os04t0518400-01 | Sobic.004G220700.1.p | zingiber_officinale042378354.1 | Eg09_t011330 | ML1h07G01701.1 | Mbe06_t048840.1 | Macma4_05_g21700.1 | Mba11_g14220 | Ms08t220210 |
| | | XP_003575403.1 | AL6G03803 | Os05t0427400-00 | Sobic.004G220600.1.p | zingiber_officinale042378359.1 | Eg02_t001010 | ML1h02G00424.1 | Mbe03_t031150.2 | Macma4_11_g22170.1 | Mba05_g13940 | Ms11t319150 |
| | | XP_003575240.2 | AL6G03821 | Os02t0626600-00 | Sobic.004G220500.1.p | | Eg09_t017980 | ML1h01G01720.1 | Mbe01_t022110.1 | Macma4_08_g14800.1 | Mba01_g14280 | Ms05t135820 |
| | | XP_003575238.1 | AL6G03336 | Os02t0627100-01 | Sobic.004G220400.1.p | | Eg05_t024420 | ML1h01G01719.1 | Mbe08_t037020.1 | Macma4_09_g16190.1 | | Ms09t254450 |

Musaceae flavonoid biosynthetic genes

| | | | | | |
|---|---|---|---|---|---|
| **PAL** | ALContig168G00001<br>AL4G00959<br>AL4G00972<br>AL4G00961<br>AL4G00973<br>AL4G00960<br>AL4G00974<br>AL4G03334<br>ALContig3G00022<br>AL1G02823<br>AL1G02824<br>AL1G02822<br>AL4G01837<br>AL4G01838<br>ALContig3G00021<br>AL4G01839<br>ALContig3G00020<br>AL6G03337 | Os02t0626400-02<br>Os12t0461300-00<br>Os12t0461050-00<br>Os08t0308300-00 | | Eg06_t005550 | ML1h06G02093.1 |

| | | | | | | | | | | | | |
|---|---|---|---|---|---|---|---|---|---|---|---|---|
| **C4H** | AT2G30490 | XP_003564495.1 | AL1G02710 | Os05t0320700-02 | Sobic.003G337400.1.p | zingiber_officinale042446994.1 | Eg08_t019660 | ML1h05G01754.1 | Mbe01_t038520.1 | Macma4_10_g16840.1 | Mba09_g04380 | Ms10t281630 |
| | | XP_003568699.1 | AL4G03405 | | Sobic.002G126600.1.p | zingiber_officinale042470427.1 | Eg07_t003130 | ML1h08G00252.1 | Mbe10_t000860.1 | Macma4_07_g20380.1 | Mba10_g11450 | Ms07t197940 |
| | | | AL1G06020 | | | zingiber_officinale042429383.1 | Eg08_t048930 | ML1h05G04363.1 | Mbe02_t033650.1 | Macma4_06_g10450.1 | Mba06_g09560 | Ms09t244050 |
| | | | AL1G06010 | | | | Eg07_t034110 | ML1h08G03392.1 | Mbe08_t025090.1 | Macma4_09_g04800.1 | Mba07_g18380 | |
| | | | AL1G06052 | | | | | | | | | |
| **4CL** | AT1G51680 | XP_010229245.1 | AL7G03614 | Os04t0310800-01 | Sobic.003G004900.1.p | zingiber_officinale042379474.1 | Eg02_t002700 | ML1h07G00167.1 | Mbe06_t016280.1 | Macma4_08_g13040.1 | Mba05_g21540 | Ms08t218760 |
| | AT3G21240 | XP_003574399.2 | AL1G00350 | Os03t0132000-01 | Sobic.006G079500.2.p | zingiber_officinale042379472.1 | Eg05_t023200 | ML1h08G00140.1 | Mbe03_t000510.1 | Macma4_05_g22890.1 | Mba08_g12720 | Ms05t136800 |
| | AT1G65060 | XP_010235057.2 | AL1G01139 | Os07t0280200-01 | Sobic.001G516600.1.p | zingiber_officinale042432142.1 | Eg02_t037340 | ML1h04G04085.1 | Mbe07_t014630.1 | Macma4_02_g21360.1 | Mba02_g18560 | Ms02t041000 |
| | AT3G21230 | XP_014757415.1 | AL7G03615 | Os01t0901600-01 | Sobic.002G158900.1.p | zingiber_officinale042457757.1 | Eg09_t018540 | ML1h03G00142.1 | Mbe05_t019740.1 | Macma4_11_g22750.1 | Mba11_g20940 | Ms11t319670 |
| | | XP_003577602.1 | AL5G02610 | Os01t0901500-01 | Sobic.001G189300.1.p | zingiber_officinale042457755.1 | Eg09_t002410 | ML1h02G00664.1 | Mbe01_t019280.1 | Macma4_11_g14090.1 | Mba11_g05710 | Ms11t311430 |
| | | XP_014753483.1 | AL3G05698 | Os10t0578950-00 | Sobic.003G391600.1.p | zingiber_officinale042469802.1 | Eg02_t001820 | ML1h08G03149.1 | Mbe01_t019320.2 | Macma4_08_g13960.1 | Mba11_g12600 | Ms08t219500 |
| | | XP_003564855.1 | ALContig36G00007 | Os08t0143300-00 | Sobic.002G009600.1.p | zingiber_officinale042469803.1 | Eg05_t033420 | ML1h01G00403.1 | Mbe05_t004680.1 | Macma4_05_g28920.1 | Mba08_g13620 | Ms05t142250 |
| | | XP_003558787.1 | AL2G03972 | Os03t0152400-01 | Sobic.002G009700.2.p | zingiber_officinale042467448.1 | Eg01_t038400 | ML1h01G00405.1 | Mbe03_t020730.1 | Macma4_11_g06130.1 | Mba05_g27160 | Ms11t304430 |

| | | | | | | | | | | | |
|---|---|---|---|---|---|---|---|---|---|---|---|
| **4CL** | XP_003570153.1 | AL3G05400 | Os02t0697400-01 | Sobic.001G282800.1.p | zingiber_officinale042469801.1 | Eg03_t041750 | ML1h01G00972.1 | Mbe05_t011100.1 | Macma4_01_g07350.1 | Mba01_g16650 | Ms01t006680 |
| | XP_010235187.1 | AL3G05411 | Os06t0656500-01 | Sobic.001G187000.2.p | zingiber_officinale042392253.1 | Eg04_t006840 | ML1h07G00248.1 | Mbe07_t013640.1 | Macma4_03_g14310.1 | Mba03_g13490 | Ms03t059300 |
| | XP_003570513.1 | AL3G05412 | Os06t0656500-02 | Sobic.001G500800.4.p | zingiber_officinale042375877.1 | Eg01_t014520 | ML1h04G02613.1 | Mbe02_t027270.1 | Macma4_06_g39580.1 | Mba06_g36630 | Ms06t178240 |
| | XP_003563338.1 | AL3G05415 | Os02t0177600-01 | Sobic.004G272700.1.p | zingiber_officinale042470465.1 | Eg01_t025930 | ML1h07G03382.1 | Mbe06_t040140.1 | Macma4_01_g19800.1 | Mba01_g28370.1 | Ms01t017810.1 |
| | XP_003573604.1 | AL4G04549 | Os08t0245200-01 | Sobic.010G214900.1.p | zingiber_officinale042375841.1 | Eg04_t025780 | ML1h02G02377.1 | Mbe03_t056570.1 | Macma4_03_g13930.1 | Mba04_g05720.1 | Ms03t058920.1 |
| | XP_003579357.2 | AL3G06362 | Os08t0448000-01 | Sobic.004G062500.1.p | zingiber_officinale042468711.1 | Eg04_t032900 | ML1h09G01309.1 | Mbe01_t005810.2 | Macma4_04_g06110.1 | Mba04_g12340.1 | Ms04t083350.1 |
| | | AL6G03059 | Os04t0683700-01 | Sobic.007G089900.1.p | zingiber_officinale042375680.1 | Eg07_t001640 | ML1h08G00557.1 | Mbe03_t000920.1 | Macma4_04_g13130.1 | Mba07_g04890.1 | Ms04t089570.1 |
| | | AL5G02390 | Os07t0639100-00 | Sobic.007G145600.1.p | zingiber_officinale042451259.1 | Eg07_t014740 | ML1h06G03426.1 | Mbe07_t022330.2 | Macma4_07_g05220.1 | Mba07_g17040.1 | Ms07t183930.1 |
| | | AL2G03024 | | Sobic.006G272600.1.p | zingiber_officinale042471368.1 | | ML1h06G02761.1 | Mbe04_t006230.1 | Macma4_07_g18890.1 | | Ms07t196560.1 |
| | | AL2G03265 | | | zingiber_officinale042405667.1 | | ML1h09G02144.1 | Mbe04_t013920.2 | Macma4_09_g24760.1 | | Ms09t262230.1 |
| | | AL5G02618 | | | zingiber_officinale042405669.1 | | | | | | |
| | | AL7G00468 | | | zingiber_officinale042441626.1 | | | | | | |

| | | | | | | | | | | | | |
|---|---|---|---|---|---|---|---|---|---|---|---|---|
| **4CL** | | | ALContig115G00003 | | | zingiber_officinale042388974.1 | | | | | | |
| | | | AL1G00990 | | | zingiber_officinale042452682.1 | | | | | | |
| | | | AL6G01256 | | | zingiber_officinale042376059.1 | | | | | | |
| | | | ALContig6G00008 | | | zingiber_officinale042447280.1 | | | | | | |
| **CHI** | AT3G55120 | XP_003559241.1 | AL3G02323 | Os03t0819600-02 | Sobic.001G035600.1.p | zingiber_officinale042387863.1 | Eg01_t015550 | ML1h09G01419.1 | Mbe04_t020690.1 | Macma4_04_g18950.1 | Mba11_g21110 | Ms11t319860 |
| | | XP_010238805.1 | AL7G01891 | Os12t0115700-01 | Sobic.008G030100.2.p | zingiber_officinale042392070.1 | Eg09_t018750 | ML1h04G02595.1 | Mbe06_t039860.1 | Macma4_11_g22930.1 | Mba04_g18040 | Ms04t093250 |
| | | | | Os11t0116300-01 | | | | | | | | |
| **CHS** | AT5G13930 | XP_003577513.1 | AL7G05184 | Os07t0214900-01 | Sobic.007G058900.1.p | zingiber_officinale042470361.1 | Eg08_t046640 | ML1h05G01818.1 | Mbe01_t038010.2 | Macma4_06_g12560.1 | Mba06_g11590 | Ms10t280970 |
| | | | | Os11t0530600-01 | Sobic.005G137200.1.p | zingiber_officinale042436018.1 | Eg08_t040870 | ML1h05G03634.1 | Mbe01_t038030.1 | Macma4_06_g18170.1 | Mba06_g16830 | Ms06t158650 |
| | | | | | Sobic.005G137000.1.p | zingiber_officinale042434575.1 | Eg08_t020390 | ML1h05G04408.1 | Mbe01_t041060.1 | Macma4_10_g16190.1 | Mba10_g10880 | MsSC51t327370 |
| | | | | | Sobic.005G136450.1.p | zingiber_officinale042434725.1 | Eg08_t049390 | ML1h05G04406.1 | Mbe01_t047580.1 | Macma4_06_g10020.1 | Mba06_g09110 | MsSC51t327380 |
| | | | | | Sobic.005G136600.1.p | zingiber_officinale042435235.1 | Eg08_t049400 | | Mbe11_t002790.1 | Macma4_06_g10010.1 | Mba06_g09080 | MsSC66t327440 |

Musaceae flavonoid biosynthetic genes

| | | | | | | | | | | | | |
|---|---|---|---|---|---|---|---|---|---|---|---|---|
| **CHS** | | | | | Sobic.005G136800.1.p | zingiber_officinale042378779.1 | Eg08_t049410 | | | Macma4_10_g16140.1 | Mba06_g09090 | |
| | | | | | Sobic.005G137100.1.p | zingiber_officinale042434761.1 | | | | | Mba06_g09120 | |
| | | | | | Sobic.005G136200.1.p | | | | | | Mba10_g10820 | |
| | | | | | Sobic.005G137300.1.p | | | | | | | |
| | | | | | Sobic.005G136300.1.p | | | | | | | |
| **F3H** | AT3G51240 | XP_003579435.1 | AL4G00149 | Os04t0662600-00 | Sobic.003G021700.1.p | zingiber_officinale042448017.1 | Eg07_t027880 | ML1h07G02004.1 | Mbe07_t035880.1 | Macma4_07_g17110.1 | Mba02_g04670 | Ms02t027200 |
| | | | | | Sobic.004G310100.1.p | | Eg02_t022060 | ML1h08G02519.1 | Mbe02_t010150.1 | Macma4_02_g06130.1 | Mba07_g15320 | |
| **F3'H** | AT5G07990 | XP_003574810.2 | AL7G01297 | Os10t0317900-02 | Sobic.009G162500.1.p | zingiber_officinale042377047.1 | Eg03_t018110 | ML1h03G02252.1 | Mbe06_t019910.1 | Macma4_03_g33170.1 | Mba01_g02030 | Ms03t075990 |
| | | XP_010233685.1 | AL2G05114 | Os10t0320100-01 | Sobic.004G200900.1.p | zingiber_officinale042458020.1 | | | | | | |
| | | XP_003562884.2 | | | Sobic.004G201100.1.p | | | | | | | |
| | | XP_003577475.1 | | | Sobic.004G200800.1.p | | | | | | | |
| | | | | | Sobic.004G200833.1.p | | | | | | | |

Musaceae flavonoid biosynthetic genes

| | | | | | | | | | | | | |
|---|---|---|---|---|---|---|---|---|---|---|---|---|
| **F3'5'H** | AT4G12300 | XP_014753340.1 | AL3G03749 | Os03t0367101-00 | Sobic.001G360800.2.p | zingiber_officinale042422124.1 | Eg09_t020640 | ML1h07G00690.1 | Mbe07_t008510.1 | Macma4_08_g08290.1 | Mba08_g07950 | Ms08t214430 |
| | AT4G12320 | XP_003567588.1 | | | Sobic.001G360700.1.p | zingiber_officinale042409260.1 | Eg02_t007770 | ML1h04G02425.1 | Mbe06_t049500.1 | Macma4_11_g05110.1 | Mba11_g04840 | Ms09t255750 |
| | AT4G12310 | | | | | zingiber_officinale042375298.1 | Eg06_t004020 | ML1h07G00951.1 | Mbe02_t005590.1 | Macma4_08_g05480.1 | Mba09_g16030 | Ms08t211840 |
| | AT4G12330 | | | | | zingiber_officinale042375301.1 | Eg07_t042500 | ML1h08G03854.1 | Mbe02_t005610.1 | Macma4_09_g17660.1 | Mba10_g00740 | Ms02t024370.1 |
| | AT5G44620 | | | | | zingiber_officinale042375302.1 | Eg02_t010650 | ML1h06G02235.1 | Mbe02_t006410.1 | Macma4_10_g01240.1 | Mba02_g01840.1 | Ms02t024400.1 |
| | AT4G22690 | | | | | zingiber_officinale042375303.1 | Eg02_t019070 | ML1h06G02237.1 | Mbe02_t005600.1 | Macma4_10_g01260.1 | Mba02_g01880.1 | Ms02t024410.1 |
| | AT4G22710 | | | | | zingiber_officinale042375299.1 | Eg02_t019100 | ML1h07G01736.1 | Mbe08_t039500.1 | Macma4_02_g03040.1 | Mba02_g01890.1 | Ms02t024420.1 |
| | | | | | | zingiber_officinale042410615.1 | Eg02_t019110 | ML1h07G01737.1 | Mbe09_t008950.1 | Macma4_02_g03240.1 | Mba02_g01900.1 | Ms11t312500.1 |
| | | | | | | | Eg02_t019180 | ML1h02G01787.1 | Mbe09_t008960.1 | Macma4_02_g03250.1 | Mba02_g01910.1 | |
| | | | | | | | Eg09_t010710 | ML1h03G01872.1 | Mbe07_t005300.1 | Macma4_02_g03260.1 | Mba02_g01920.1 | |
| | | | | | | | | ML1h07G01741.1 | | Macma4_02_g03290.1 | Mba02_g01940.1 | |
| | | | | | | | | ML1h04G03337.1 | | Macma4_02_g03300.1 | Mba02_g01950.1 | |

| | | | | | | | | | | | | |
|---|---|---|---|---|---|---|---|---|---|---|---|---|
| | | | | | | | | | | Macma4_11_g15260.1 | Mba11_g13650.1 | |
| **FLS** | AT5G08640 | XP_003570562.1 | AL7G03380 | Os02t0767300-01(OsFLS) | Sobic.003G021700.1.p | zingiber_officinale042378807.1 | Eg06_t006150 | ML1h03G01030.1 | Mbe08_t015900.1 | Macma4_03_g07220.1 | Mba08_g23630 | Ms08t229420 |
| | AT5G63580 | | AL6G04177 | | Sobic.004G310100.2.p | zingiber_officinale042406407.1 | Eg01_t007540 | ML1h05G00636.1 | Mbe02_t047300.2 | Macma4_08_g24690.1 | Mba10_g22000 | Ms03t052640 |
| | AT5G63590 | | AL6G04187 | | | zingiber_officinale042406408.1 | Eg08_t007020 | ML1h05G00635.1 | Mbe08_t036240.1 | | Mba10_g21980 | Ms10t292840 |
| | AT5G63595 | | | | | | Eg03_t031640 | ML1h06G02043.1 | Mbe03_t008100.1 | | Mba03_g06940 | |
| | AT5G63600 | | | | | | | ML1h09G00678.1 | | | Mba09_g14060 | |
| | AT5G43935 | | | | | | | | | | | |
| **DFR** | AT5G42800 | XP_010232065.1 | AL1G03574 | Os01t0633500-01 | Sobic.004G050200.1.p | zingiber_officinale042421830.1 | Eg09_t036940 | ML1h04G00849.1 | Mbe04_t036630.1 | Macma4_04_g33220.1 | Mba04_g31800 | Ms04t107810 |
| | | XP_003569417.2 | | | Sobic.003G230900.1.p | | Eg04_t027830 | ML1h06G03238.1 | Mbe06_t019010.2 | Macma4_03_g33910.1 | Mba01_g01310 | Ms04t087800 |
| | | | | | Sobic.003G231000.1.p | | Eg03_t018610 | ML1h03G02164.1 | Mbe04_t011700.1 | Macma4_04_g11180.1 | | Ms03t076720 |
| | | | | | Sobic.009G043800.1.p | | | | | | | |
| **LDOX/ANS** | AT4G22880 | | AL7G00463 | Os01t0372500-00 | Sobic.004G000700.1.p | zingiber_officinale042431763.1 | Eg05_t014690 | ML1h01G01735.1 | Mbe03_t031300.1 | Macma4_05_g03920.1 | Mba05_g13780 | Ms05t120140 |
| | | | AL7G00477 | Os06t0626700-00 | | | | | | | | |

Musaceae flavonoid biosynthetic genes

| | | | | | | | | | | | | |
|---|---|---|---|---|---|---|---|---|---|---|---|---|
| **UGT78D2** | AT5G17050 | XP_014753234.1 | AL5G00921 | Os06t0192100-00 | Sobic.010G070200.1.p | zingiber_officinale042423608.1 | Eg08_t008340 | ML1h09G00476.1 | Mbe07_t034770.1 | Macma4_10_g27580.1 | Mba03_g04830 | Ms10t291590 |
| | | XP_024313830.1 | | Os07t0148200-01 | Sobic.002G369600.1.p | zingiber_officinale042410446.1 | Eg06_t008220 | ML1h06G01843.1 | Mbe02_t045890.1 | Macma4_07_g16200.1 | Mba10_g20800 | Ms07t193970 |
| | | | | | | zingiber_officinale042388078.1 | Eg01_t005290 | ML1h05G00763.1 | Mbe03_t010580.1 | Macma4_03_g04880.1 | Mba07_g14580 | Ms03t050760 |
| | | | | | | zingiber_officinale042460585.1 | | | | | | |
| | | | | | | zingiber_officinale042458777.1 | | | | | | |
| | | | | | | zingiber_officinale042458785.1 | | | | | | |
| | | | | | | zingiber_officinale042458786.1 | | | | | | |
| | | | | | | zingiber_officinale042458779.1 | | | | | | |
| | | | | | | zingiber_officinale042458782.1 | | | | | | |
| | | | | | | zingiber_officinale042458775.1 | | | | | | |
| | | | | | | zingiber_officinale042458790.1 | | | | | | |
| | | | | | | zingiber_officinale042458800.1 | | | | | | |

| | |
|---|---|
| **UGT78D2** | zingiber_officinale042458794.1<br>zingiber_officinale042458797.1<br>zingiber_officinale042458783.1<br>zingiber_officinale042458788.1<br>zingiber_officinale042458780.1<br>zingiber_officinale042458787.1<br>zingiber_officinale042458795.1<br>zingiber_officinale042458796.1<br>zingiber_officinale042458799.1<br>zingiber_officinale042458791.1<br>zingiber_officinale042458798.1<br>zingiber_officinale042458778.1 |

Musaceae flavonoid biosynthetic genes

| | | | | | | | | | | | | |
|---|---|---|---|---|---|---|---|---|---|---|---|---|
| **UGT78D2** | | | | | | zingiber_officinale042458789.1<br>zingiber_officinale042458784.1<br>zingiber_officinale042458781.1<br>zingiber_officinale042458793.1 | | | | | | |
| **UGT75C1** | AT4G14090 | XP_003576359.2 | AL5G03818 | Os06t0593800-01 | Sobic.010G179150.1.p | zingiber_officinale042413493.1 | Eg08_t003590 | ML1h05G00327.1 | Mbe02_t051150.1 | Macma4_10_g31950.1 | Mba10_g24560 | Ms06t152710 |
| | | XP_024311125.1 | AL5G03820 | Os01t0179600-01 | Sobic.010G179500.1.p | zingiber_officinale042413492.1 | Eg08_t037030 | ML1h05G03318.1 | Mbe01_t034880.1 | Macma4_06_g07210.1 | Mba06_g06480 | Ms06t152720 |
| | | XP_003560621.1 | AL5G03827 | Os05t0179950-00 | Sobic.003G042900.1.p | zingiber_officinale042413495.1 | Eg08_t037040 | ML1h05G03317.1 | Mbe01_t034870.1 | Macma4_06_g07200.1 | Mba06_g06490 | Ms06t152730 |
| | | XP_003571494.1 | AL6G02053 | Os06t0593200-01 | Sobic.009G066100.1.p | zingiber_officinale042413494.1 | | | | Macma4_06_g07190.1 | | Ms10t295740 |
| | | XP_003560622.3 | AL6G02057 | Os11t0145200-01 | Sobic.010G178900.2.p | zingiber_officinale042453821.1 | | | | | | |
| | | XP_010230853.1 | AL5G00208 | Os02t0203300-00 | Sobic.010G179000.1.p | zingiber_officinale042405426.1 | | | | | | |
| | | XP_024315058.1 | AL7G04281 | | Sobic.005G032200.2.p | zingiber_officinale042404506.1 | | | | | | |
| | | XP_003565425.2 | AL5G00933 | | Sobic.006G097800.2.p | | | | | | | |

| | | | | | | | | | | | | |
|---|---|---|---|---|---|---|---|---|---|---|---|---|
| | | XP_003565426.1 | AL3G02472 | | Sobic.008G011700.1.p | | | | | | | |
| **UGT79B1** | AT5G54060 | XP_003560487.1 | AL3G06005 | Os11t0457300-01 | Sobic.008G035600.1.p | zingiber_officinale042420251.1 | Eg07_t037900 | ML1h08G03185.1 | Mbe08_t027530.1 | Macma4_09_g07100.1 | | Ms09t246270 |
| | | XP_003560486.1 | | | Sobic.005G033900.1.p | | | | | | | |
| | | | | | Sobic.005G033801.1.p | | | | | | | |
| **LAR** | 0 | XP_003561802.1 | AL3G04509 | Os03t0259400-00 | 0 | 0 | Eg05_t019870 | ML1h01G01259.1 | Mbe05_t014700.1 | Macma4_05_g19560.1 | Mba05_g18490 | Ms05t133790 |
| **ANR** | AT1G61720 | XP_003560368.1 | AL4G03119 | Os07t0601900-01 | Sobic.002G366200.3.p | zingiber_officinale042454473.1 | Eg06_t013840 | ML1h06G01340.1 | Mbe07_t028460.1 | Macma4_07_g10640.1 | Mba08_g01400 | Ms08t207990 |
| | | XP_003562747.1 | AL4G03124 | Os07t0602000-01 | Sobic.002G366100.1.p | zingiber_officinale042425557.1 | Eg02_t015210 | ML1h06G01342.1 | | Macma4_07_g10660.1 | | Ms07t188920 |
| | | XP_003559994.1 | AL4G03120 | Os07t0598000-02 | | zingiber_officinale042425546.1 | | ML1h06G01341.1 | | Macma4_07_g10670.1 | | Ms07t188910 |
| | | XP_003562745.1 | AL4G03125 | Os07t0601100-01 | | zingiber_officinale042425536.1 | | ML1h07G01339.1 | | Macma4_07_g10680.1 | | |
| | | XP_003580613.1 | AL5G00711 | Os07t0601000-01 | | | | | | Macma4_08_g01440.1 | | |
| | | XP_014751318.1 | AL4G00442 | | | | | | | | | |
| | | XP_003580612.1 | ALContig28G00015 | | | | | | | | | |

Musaceae flavonoid biosynthetic genes

| | | | | | | | | | | | | |
|---|---|---|---|---|---|---|---|---|---|---|---|---|
| **ANR** | | XP_003580614.1 | AL4G00443<br>ALContig28G00016<br>AL3G05634<br>AL3G05652<br>AL4G00441<br>ALContig28G00014<br>AL4G00440<br>ALContig28G00013<br>AL4G00436<br>ALContig28G00010<br>AL4G00439<br>ALContig28G00012 | | | | | | | | | |
| **TT8** | AT4G09820 | XP_024312733.1 | AL4G04360 | Os07t0211500-01 | Sobic.002G076600.4.p | zingiber_officinale042379222.1 | Eg04_t020700 | ML1h03G03510.1 | Mbe06_t034420.1 | Macma4_06_g25800.1 | Mba06_g23950 | MsSC186t326890 |
| | | XP_010230011.1 | | | | | Eg03_t004200 | ML1h02G03012.1 | Mbe06_t001720.2 | Macma4_03_g19880.1 | Mba03_g18730 | Ms03t064490 |
| | | XP_024312732.1 | | | | | | | | | | |
| | AT5G24520 | XP_003570109.1 | ALContig2G00010 | Os02t0682500-01 | Sobic.004G280800.1.p | zingiber_officinale042406919.1 | Eg05_t009200 | ML1h01G02333.1 | Mbe03_t038300.1 | Macma4_05_g10420.1 | Mba05_g07670 | Ms04t112120 |

Musaceae flavonoid biosynthetic genes

| | | | | | | | | | | | | |
|---|---|---|---|---|---|---|---|---|---|---|---|---|
| **GLABRA1(TTG1)** | | | | | | zingiber_officinale042451789.1 | Eg09_t041330 | ML1h04G00438.1 | Mbe04_t041380.1 | Macma4_04_g37870.1 | Mba04_g36060 | Ms05t125970 |
| | | | | | | zingiber_officinale042377982.1 | | | | | | |
| **C1** | | XP_003561552.1 | AL3G03580 | Os06t0205100-02 | Sobic.001G340900.1.p | zingiber_officinale042388738.1 | Eg07_t019620 | ML1h08G01785.1 | Mbe01_t033880.1 | Macma4_10_g21630.1 | Mba10_g15520 | Ms09t266370.1 |
| | | | AL5G00993 | | | zingiber_officinale042387580.1 | Eg08_t035970 | ML1h05G03234.1 | Mbe03_t051550.1 | Macma4_09_g29590.1 | Mba06_g05630 | Ms06t151760.1 |
| | | | | | | zingiber_officinale042417551.1 | | ML1h05G01322.1 | Mbe02_t039140.1 | | Mba09_g23680 | |
| | | | | | | zingiber_officinale042423943.1 | | | | | | |
| | | | | | | zingiber_officinale042423006.1 | | | | | | |

**Table S2a**. The information of flavonoid biosynthetic genes identified in *M. beccarii*.

| ID | Name | length | MW(Da) | pI | Localizations |
|---|---|---|---|---|---|
| Mbe05_t009880.2 | *MbePAL1* | 557 | 60494.9 | 6.35 | Cytoplasm |
| Mbe06_t040740.1 | *MbePAL2* | 713 | 77095.3 | 6.6 | Cytoplasm |
| Mbe07_t015550.1 | *MbePAL3* | 709 | 76790 | 6.56 | Cytoplasm |
| Mbe06_t040740.2 | *MbePAL4* | 557 | 60665 | 6.57 | Cytoplasm |
| Mbe06_t048840.1 | *MbePAL5* | 701 | 75831.8 | 5.77 | Cytoplasm |
| Mbe03_t031150.2 | *MbePAL6* | 423 | 46990.1 | 6.58 | Cytoplasm |
| Mbe01_t022110.1 | *MbePAL7* | 713 | 77124.6 | 6.26 | Cytoplasm |
| Mbe08_t037020.1 | *MbePAL8* | 757 | 83240.1 | 5.59 | Cytoplasm |
| Mbe01_t038520.1 | *MbeC4H1* | 503 | 57545.6 | 9.56 | Endoplasmic reticulum |
| Mbe10_t000860.1 | *MbeC4H2* | 503 | 57405.4 | 9.66 | Endoplasmic reticulum |
| Mbe02_t033650.1 | *MbeC4H3* | 503 | 57436.4 | 9.69 | Endoplasmic reticulum |
| Mbe08_t025090.1 | *MbeC4H4* | 503 | 57928 | 9.36 | Endoplasmic reticulum |
| Mbe06_t016280.1 | *Mbe4CL1* | 545 | 59022.8 | 6.74 | Cytoplasm |
| Mbe03_t000510.1 | *Mbe4CL2* | 650 | 70159.3 | 8.85 | Peroxisome |
| Mbe07_t014630.1 | *Mbe4CL3* | 549 | 59243.6 | 8.73 | Peroxisome |
| Mbe05_t019740.1 | *Mbe4CL4* | 625 | 66173.6 | 7.81 | Peroxisome |
| Mbe01_t019280.1 | *Mbe4CL5* | 555 | 59588.4 | 8.79 | Peroxisome |
| Mbe01_t019320.2 | *Mbe4CL6* | 503 | 53550.1 | 6.65 | Cytoplasm |
| Mbe05_t004680.1 | *Mbe4CL7* | 550 | 58618.9 | 7.02 | Peroxisome |
| Mbe03_t020730.1 | *Mbe4CL8* | 548 | 59184.8 | 8.21 | Peroxisome |
| Mbe05_t011100.1 | *Mbe4CL9* | 544 | 58594.2 | 6.99 | Peroxisome |
| Mbe07_t013640.1 | *Mbe4CL10* | 543 | 58425.9 | 8.34 | Peroxisome |

| | | | | | |
|---|---|---|---|---|---|
| Mbe02_t027270.1 | *Mbe4CL11* | 552 | 59673.4 | 6.2 | Cytoplasm |
| Mbe06_t040140.1 | *Mbe4CL12* | 568 | 61167.2 | 5.37 | Cytoplasm |
| Mbe03_t056570.1 | *Mbe4CL13* | 543 | 59202.6 | 5.65 | Cytoplasm |
| Mbe01_t005810.2 | *Mbe4CL14* | 471 | 50634.8 | 4.78 | Cytoplasm |
| Mbe03_t000920.1 | *Mbe4CL15* | 548 | 59482.1 | 5.52 | Cytoplasm |
| Mbe07_t022330.2 | *Mbe4CL16* | 400 | 43002.4 | 5.63 | Cytoplasm |
| Mbe04_t006230.1 | *Mbe4CL17* | 543 | 59060.5 | 5.66 | Cytoplasm |
| Mbe04_t013920.2 | *Mbe4CL18* | 404 | 43697 | 6.01 | Cytoplasm |
| Mbe04_t020690.1 | *MbeCHI1* | 255 | 26702.3 | 4.84 | Cytoplasm |
| Mbe06_t039860.1 | *MbeCHI2* | 209 | 23445.7 | 4.81 | Cytoplasm |
| Mbe01_t038010.2 | *MbeCHS1* | 520 | 57925.8 | 8.55 | Nucleus |
| Mbe01_t038030.1 | *MbeCHS2* | 392 | 43202.4 | 7.06 | Cytoplasm |
| Mbe01_t041060.1 | *MbeCHS3* | 493 | 53761.4 | 6.64 | Cytoplasm |
| Mbe01_t047580.1 | *MbeCHS4* | 480 | 52053.9 | 7.97 | Plastid |
| Mbe11_t002790.1 | *MbeCHS5* | 392 | 42951.2 | 6.43 | Cytoplasm |
| Mbe07_t035880.1 | *MbeF3H1* | 373 | 41367.6 | 5.1 | Cytoplasm |
| Mbe06_t019910.1 | *MbeF3'H* | 520 | 57128.1 | 7.69 | Endoplasmic reticulum |
| Mbe07_t008510.1 | *MbeF3'5'H1* | 503 | 56483.5 | 9.49 | Endoplasmic reticulum |
| Mbe06_t049500.1 | *MbeF3'5'H2* | 505 | 56211.9 | 9.25 | Endoplasmic reticulum |
| Mbe02_t005590.1 | *MbeF3'5'H3* | 505 | 56293 | 8.46 | Endoplasmic reticulum |
| Mbe02_t005610.1 | *MbeF3'5'H4* | 370 | 40785.5 | 10.29 | Endoplasmic reticulum |
| Mbe02_t006410.1 | *MbeF3'5'H5* | 540 | 59944.2 | 8.68 | Endoplasmic reticulum |
| Mbe02_t005600.1 | *MbeF3'5'H6* | 208 | 23306.7 | 6.63 | Cytoplasm |
| Mbe08_t039500.1 | *MbeF3'5'H7* | 505 | 56209.8 | 7.85 | Endoplasmic reticulum |
| Mbe09_t008950.1 | *MbeF3'5'H8* | 507 | 56012.4 | 8.47 | Endoplasmic reticulum |

| | | | | | |
|---|---|---|---|---|---|
| Mbe09_t008960.1 | *MbeF3'5'H9* | 507 | 56005.4 | 8.47 | Endoplasmic reticulum |
| Mbe07_t005300.1 | *MbeF3'5'H10* | 523 | 57292.8 | 8.16 | Endoplasmic reticulum |
| Mbe08_t015900.1 | *MbeFLS1* | 398 | 44295.4 | 7.57 | Nucleus |
| Mbe02_t047300.2 | *MbeFLS2* | 291 | 32583.2 | 6.13 | Cytoplasm |
| Mbe08_t036240.1 | *MbeFLS3* | 406 | 45798.9 | 8.64 | Cytoplasm |
| Mbe03_t008100.1 | *MbeFLS4* | 333 | 37372.4 | 5.43 | Cytoplasm |
| Mbe04_t036630.1 | *MbeDFR1* | 342 | 38256.6 | 6 | Cytoplasm |
| Mbe06_t019010.2 | *MbeDFR2* | 378 | 42879.6 | 6.07 | Cytoplasm |
| Mbe04_t011700.1 | *MbeDFR3* | 358 | 40360.1 | 6.29 | Cytoplasm |
| Mbe03_t031300.1 | *MbeLDOX* | 398 | 44831.2 | 5.69 | Cytoplasm\|Nucleus |
| Mbe07_t034770.1 | *MbeUGT78D2.1* | 458 | 47917.9 | 6.45 | Cytoplasm |
| Mbe02_t045890.1 | *MbeUGT78D2.2* | 455 | 47753.5 | 6.55 | Cytoplasm |
| Mbe03_t010580.1 | *MbeUGT78D2.3* | 361 | 38355 | 6.05 | Cytoplasm |
| Mbe02_t051150.1 | *MbeUGT75C1.1* | 501 | 54513 | 5.31 | Cytoplasm |
| Mbe01_t034880.1 | *MbeUGT75C1.2* | 579 | 64964.9 | 6.66 | Cytoplasm |
| Mbe01_t034870.1 | *MbeUGT75C1.3* | 473 | 52570.3 | 5.1 | Cytoplasm |
| Mbe08_t027530.1 | *MbeUGT79B1* | 471 | 51052.7 | 7.07 | Cytoplasm |
| Mbe05_t014700.1 | *MbeLAR* | 358 | 38681.8 | 6.84 | Cytoplasm |
| Mbe07_t028460.1 | *MbeANR* | 341 | 37711.7 | 5.43 | Cytoplasm |
| Mbe06_t034420.1 | *MbeTT8.1* | 693 | 77470.9 | 6.14 | Nucleus |
| Mbe06_t001720.2 | *MbeTT8.2* | 643 | 72324 | 5.03 | Nucleus |
| Mbe03_t038300.1 | *MbeTTG1.1* | 378 | 41003.5 | 4.55 | Cytoplasm\|Nucleus |
| Mbe04_t041380.1 | *MbeTTG1.2* | 340 | 36953 | 4.67 | Cytoplasm\|Nucleus |
| Mbe01_t033880.1 | *MbeC1.1* | 274 | 30268.9 | 7.92 | Nucleus |
| Mbe03_t051550.1 | *MbeC1.2* | 260 | 28985.5 | 7.47 | Nucleus |

| | | | | | |
|---|---|---|---|---|---|
| Mbe02_t039140.1 | *MbeC1.3* | 278 | 30575.2 | 7.71 | Nucleus |

**Table S2b.** The information of flavonoid biosynthetic genes identified in *M. lasiocarpa*.

| ID | Name | length | MW(Da) | pI | Localizations |
|---|---|---|---|---|---|
| ML1h04G02672.1 | *MlPAL1* | 713 | 76972 | 6.22 | Cytoplasm |
| ML1h01G00856.1 | *MlPAL2* | 711 | 76675 | 6.43 | Cytoplasm |
| ML1h07G00086.1 | *MlPAL3* | 709 | 77027.1 | 6.66 | Cytoplasm |
| ML1h04G03282.1 | *MlPAL4* | 703 | 76163.1 | 5.66 | Cytoplasm |
| ML1h07G01701.1 | *MlPAL5* | 711 | 76896.7 | 5.98 | Cytoplasm |
| ML1h02G00424.1 | *MlPAL6* | 712 | 77188.7 | 6.32 | Cytoplasm |
| ML1h01G01720.1 | *MlPAL7* | 709 | 76855.9 | 6.51 | Cytoplasm\|Nucleus |
| ML1h01G01719.1 | *MlPAL8* | 709 | 76908.9 | 6.63 | Cytoplasm\|Nucleus |
| ML1h06G02093.1 | *MlPAL9* | 757 | 83181.1 | 6.04 | Cytoplasm |
| ML1h05G01754.1 | *MlC4H1* | 536 | 61135.7 | 9.79 | Endoplasmic reticulum |
| ML1h08G00252.1 | *MlC4H2* | 481 | 55154.7 | 9.75 | Endoplasmic reticulum |
| ML1h05G04363.1 | *MlC4H3* | 503 | 57631.6 | 9.67 | Endoplasmic reticulum |
| ML1h08G03392.1 | *MlC4H4* | 503 | 57911.1 | 9.22 | Endoplasmic reticulum |
| ML1h07G00167.1 | *Ml4CL1* | 549 | 59273.6 | 8.93 | Peroxisome |
| ML1h08G00140.1 | *Ml4CL2* | 563 | 59603.8 | 6.52 | Peroxisome |
| ML1h04G04085.1 | *Ml4CL3* | 548 | 59500.3 | 9.04 | Peroxisome |
| ML1h03G00142.1 | *Ml4CL4* | 547 | 58681.3 | 7.5 | Peroxisome |
| ML1h02G00664.1 | *Ml4CL5* | 557 | 59598.3 | 8.22 | Peroxisome |
| ML1h08G03149.1 | *Ml4CL6* | 555 | 58892.2 | 8.21 | Peroxisome |
| ML1h01G00403.1 | *Ml4CL7* | 550 | 58556.7 | 6.89 | Peroxisome |
| ML1h01G00405.1 | *Ml4CL8* | 550 | 58810.2 | 7.2 | Peroxisome |

| | | | | | |
|---|---|---|---|---|---|
| ML1h01G00972.1 | *Ml4CL9* | 546 | 59097.7 | 7.85 | Peroxisome |
| ML1h07G00248.1 | *Ml4CL10* | 515 | 55430.6 | 6.94 | Peroxisome |
| ML1h04G02613.1 | *Ml4CL11* | 568 | 61229.3 | 5.37 | Cytoplasm |
| ML1h07G03382.1 | *Ml4CL12* | 563 | 60560.5 | 5.64 | Cytoplasm |
| ML1h02G02377.1 | *Ml4CL13* | 548 | 59367.9 | 5.16 | Cytoplasm |
| ML1h09G01309.1 | *Ml4CL14* | 548 | 59368 | 5.07 | Cytoplasm |
| ML1h08G00557.1 | *Ml4CL15* | 543 | 59144.4 | 5.1 | Cytoplasm |
| ML1h06G03426.1 | *Ml4CL16* | 543 | 59167.6 | 5.66 | Cytoplasm |
| ML1h06G02761.1 | *Ml4CL17* | 550 | 59656.4 | 6.38 | Cytoplasm |
| ML1h09G02144.1 | *Ml4CL18* | 548 | 59473.2 | 6.4 | Cytoplasm\|Peroxisome |
| ML1h09G01419.1 | *MlCHI1* | 236 | 24960.1 | 5.12 | Cytoplasm |
| ML1h04G02595.1 | *MlCHI2* | 211 | 23702.1 | 4.79 | Cytoplasm |
| ML1h05G01818.1 | *MlCHS1* | 392 | 42787.9 | 6.25 | Cytoplasm |
| ML1h05G03634.1 | *MlCHS2* | 392 | 42997.2 | 6.52 | Cytoplasm |
| ML1h05G04408.1 | *MlCHS3* | 777 | 85121.1 | 6.34 | Cytoplasm |
| ML1h05G04406.1 | *MlCHS4* | 392 | 43055.3 | 7.31 | Cytoplasm |
| ML1h07G02004.1 | *MlF3H1* | 373 | 41487 | 5.52 | Cytoplasm |
| ML1h08G02519.1 | *MlF3H2* | 373 | 41386.7 | 5.35 | Cytoplasm |
| ML1h03G02252.1 | *MlF3H3* | 520 | 57489.4 | 6.99 | Endoplasmic reticulum |
| ML1h07G00690.1 | *MlF3'5'H1* | 445 | 49816.4 | 8.45 | Mitochondrion |
| ML1h04G02425.1 | *MlF3'5'H2* | 506 | 56209.3 | 8.9 | Endoplasmic reticulum |
| ML1h07G00951.1 | *MlF3'5'H3* | 505 | 55905.5 | 7.84 | Endoplasmic reticulum |
| ML1h08G03854.1 | *MlF3'5'H4* | 507 | 56073.5 | 8.08 | Endoplasmic reticulum |
| ML1h06G02235.1 | *MlF3'5'H5* | 505 | 56222.8 | 7.59 | Endoplasmic reticulum |
| ML1h06G02237.1 | *MlF3'5'H6* | 505 | 56373.9 | 7.85 | Endoplasmic reticulum |

| | | | | | |
|---|---|---|---|---|---|
| ML1h07G01736.1 | *MlF3'5'H7* | 505 | 56088.7 | 7.72 | Endoplasmic reticulum |
| ML1h07G01737.1 | *MlF3'5'H8* | 504 | 56084.8 | 8.58 | Endoplasmic reticulum |
| ML1h02G01787.1 | *MlF3'5'H9* | 505 | 56248.9 | 8.04 | Endoplasmic reticulum |
| ML1h03G01872.1 | *MlF3'5'H10* | 505 | 56131.8 | 8.59 | Endoplasmic reticulum |
| ML1h07G01741.1 | *MlF3'5'H11* | 505 | 56186.8 | 9.02 | Endoplasmic reticulum |
| ML1h04G03337.1 | *MlF3'5'H12* | 454 | 50867 | 7.22 | Mitochondrion |
| ML1h03G01030.1 | *MlFLS1* | 333 | 37327.5 | 6.64 | Cytoplasm |
| ML1h05G00636.1 | *MlFLS2* | 335 | 37718.1 | 5.94 | Cytoplasm\|Nucleus |
| ML1h05G00635.1 | *MlFLS3* | 700 | 76851.6 | 8.14 | Mitochondrion |
| ML1h06G02043.1 | *MlFLS4* | 348 | 38764 | 6.74 | Cytoplasm\|Nucleus |
| ML1h09G00678.1 | *MlFLS5* | 333 | 37444.5 | 5.14 | Cytoplasm |
| ML1h04G00849.1 | *MlDFR1* | 342 | 38190.6 | 6.29 | Cytoplasm |
| ML1h06G03238.1 | *MlDFR2* | 358 | 40584.3 | 6.14 | Cytoplasm |
| ML1h03G02164.1 | *MlDFR3* | 356 | 40126.6 | 4.96 | Cytoplasm |
| ML1h01G01735.1 | *MlLDOX* | 360 | 40563.4 | 5.66 | Cytoplasm |
| ML1h09G00476.1 | *MlUGT78D2.1* | 459 | 48291.3 | 6.14 | Cytoplasm |
| ML1h06G01843.1 | *MlUGT78D2.2* | 458 | 47589.5 | 6.27 | Cytoplasm |
| ML1h05G00763.1 | *MlUGT78D2.3* | 455 | 47828.7 | 6.61 | Cytoplasm |
| ML1h05G00327.1 | *MlUGT75C1.1* | 477 | 52172.3 | 4.98 | Cytoplasm |
| ML1h05G03318.1 | *MlUGT75C1.2* | 464 | 51172.8 | 4.77 | Cytoplasm |
| ML1h05G03317.1 | *MlUGT75C1.3* | 466 | 51540 | 5.25 | Cytoplasm |
| ML1h08G03185.1 | *MlUGT79B1* | 694 | 75152.1 | 5.43 | Endoplasmic reticulum |
| ML1h01G01259.1 | *MlLAR* | 353 | 38132.1 | 6.84 | Cytoplasm |
| ML1h06G01340.1 | *MlANR1* | 337 | 37083.1 | 5.26 | Cytoplasm |
| ML1h06G01342.1 | *MlANR2* | 336 | 37121.1 | 5.43 | Cytoplasm |

| | | | | | |
|---|---|---|---|---|---|
| ML1h06G01341.1 | *MlANR3* | 337 | 37240.3 | 5.43 | Cytoplasm |
| ML1h07G01339.1 | *MlANR4* | 350 | 37948.2 | 6.52 | Cytoplasm |
| ML1h03G03510.1 | *MlTT8.1* | 641 | 71812.2 | 6.05 | Nucleus |
| ML1h02G03012.1 | *MlTT8.2* | 657 | 73615.5 | 5.18 | Nucleus |
| ML1h01G02333.1 | *MlTTG1.1* | 378 | 41058.5 | 4.67 | Cytoplasm\|Nucleus |
| ML1h04G00438.1 | *MlTTG1.2* | 316 | 34344.3 | 5.02 | Cytoplasm\|Nucleus |
| ML1h08G01785.1 | *MlC1.1* | 260 | 28947.5 | 7.82 | Nucleus |
| ML1h05G03234.1 | *MlC1.2* | 277 | 30559.2 | 7.97 | Nucleus |
| ML1h05G01322.1 | *MlC1.3* | 278 | 30511.3 | 8.27 | Nucleus |

**Table S2c.** The information of flavonoid biosynthetic genes identified in *E. glaucum*

| ID | Name | length | MW(Da) | pI | Localizations |
|---|---|---|---|---|---|
| Eg01_t040920 | *EgPAL1* | 713 | 77092.6 | 6.26 | Cytoplasm |
| Eg05_t014880 | *EgPAL2* | 710 | 76801.6 | 6.51 | Cytoplasm\|Nucleus |
| Eg05_t014870 | *EgPAL3* | 682 | 73719.7 | 6.59 | Cytoplasm\|Nucleus |
| Eg02_t018790 | *EgPAL4* | 712 | 77302.3 | 6.19 | Cytoplasm |
| Eg09_t011330 | *EgPAL5* | 709 | 76467.5 | 6.15 | Cytoplasm |
| Eg02_t001010 | *EgPAL6* | 710 | 76853.9 | 6.61 | Cytoplasm |
| Eg09_t017980 | *EgPAL7* | 714 | 76873 | 6.39 | Cytoplasm |
| Eg05_t024420 | *EgPAL8* | 712 | 76639.9 | 6.51 | Cytoplasm |
| Eg06_t005550 | *EgPAL9* | 758 | 83171.1 | 6.04 | Cytoplasm |
| Eg08_t019660 | *EgC4H1* | 504 | 57589.5 | 9.6 | Endoplasmic reticulum |
| Eg07_t003130 | *EgC4H2* | 504 | 57346.3 | 9.44 | Endoplasmic reticulum |
| Eg08_t048930 | *EgC4H3* | 504 | 57389.3 | 9.57 | Endoplasmic reticulum |
| Eg07_t034110 | *EgC4H4* | 504 | 57849.9 | 8.66 | Endoplasmic reticulum |

| | | | | | |
|---|---|---|---|---|---|
| Eg02_t002700 | *Eg4CL1.1* | 544 | 58412 | 8.53 | Peroxisome |
| Eg05_t023200 | *Eg4CL1.2* | 547 | 59105.8 | 8.52 | Peroxisome |
| Eg02_t037340 | *Eg4CL2.1* | 564 | 60678.7 | 5.83 | Cytoplasm |
| Eg09_t018540 | *Eg4CL2.2* | 570 | 61371.5 | 5.5 | Cytoplasm |
| Eg09_t002410 | *Eg4CL3* | 550 | 59306.1 | 8.55 | Peroxisome |
| Eg02_t001820 | *Eg4CL4* | 550 | 59197.5 | 8.94 | Peroxisome |
| Eg05_t033420 | *Eg4CL5* | 551 | 58611.8 | 6.69 | Peroxisome |
| Eg01_t038400 | *Eg4CL7* | 556 | 59297.9 | 8.31 | Peroxisome |
| Eg03_t041750 | *Eg4CL8* | 545 | 58062.5 | 7.3 | Peroxisome |
| Eg04_t006840 | *Eg4CL9* | 549 | 59476.2 | 6.51 | Cytoplasm\|Peroxisome |
| Eg01_t014520 | *Eg4CL10* | 549 | 59490.3 | 6.43 | Cytoplasm |
| Eg01_t025930 | *Eg4CL11* | 549 | 59318.9 | 5.15 | Cytoplasm |
| Eg04_t025780 | *Eg4CL12* | 544 | 59003.4 | 5.81 | Cytoplasm |
| Eg04_t032900 | *Eg4CL13* | 544 | 58907.4 | 6.15 | Cytoplasm |
| Eg07_t001640 | *Eg4CL14* | 563 | 59753 | 6.36 | Peroxisome |
| Eg07_t014740 | *Eg4CL15* | 544 | 59315.6 | 5.29 | Cytoplasm |
| Eg01_t015550 | *EgCHI1* | 237 | 24932.1 | 4.81 | Cytoplasm |
| Eg09_t018750 | *EgCHI2* | 212 | 23585.9 | 4.79 | Cytoplasm |
| Eg08_t046640 | *EgCHS1* | 393 | 42788.9 | 6.94 | Cytoplasm |
| Eg08_t040870 | *EgCHS2* | 393 | 42822.2 | 6.94 | Cytoplasm |
| Eg08_t020390 | *EgCHS3* | 393 | 42952.2 | 6.07 | Cytoplasm |
| Eg08_t049390 | *EgCHS4* | 393 | 43183.4 | 7.31 | Cytoplasm |
| Eg08_t049400 | *EgCHS5* | 393 | 43094.3 | 7.06 | Cytoplasm |
| Eg08_t049410 | *EgCHS6* | 393 | 43044.2 | 7.31 | Cytoplasm |
| Eg07_t027880 | *EgF3H1* | 374 | 41398.8 | 5.35 | Cytoplasm |

| | | | | | |
|---|---|---|---|---|---|
| Eg02_t022060 | *EgF3H2* | 374 | 41633.2 | 5.36 | Cytoplasm |
| Eg03_t018110 | *EgF3'H1* | 521 | 57527.6 | 7.48 | Endoplasmic reticulum |
| Eg09_t020640 | *EgF3'5'H1* | 508 | 56310.1 | 8.83 | Endoplasmic reticulum |
| Eg02_t007770 | *EgF3'5'H2* | 504 | 56228.2 | 9.53 | Endoplasmic reticulum |
| Eg06_t004020 | *EgF3'5'H3* | 506 | 56223.7 | 7.39 | Endoplasmic reticulum |
| Eg07_t042500 | *EgF3'5'H4* | 508 | 56049.6 | 8.47 | Endoplasmic reticulum |
| Eg02_t010650 | *EgF3'5'H5* | 506 | 55952.6 | 7.81 | Endoplasmic reticulum |
| Eg02_t019070 | *EgF3'5'H6* | 506 | 56107.7 | 7 | Endoplasmic reticulum |
| Eg02_t019100 | *EgF3'5'H7* | 341 | 37198.9 | 10.07 | Endoplasmic reticulum |
| Eg02_t019110 | *EgF3'5'H8* | 209 | 23498.9 | 5.33 | Cytoplasm |
| Eg02_t019180 | *EgF3'5'H9* | 506 | 56483.2 | 8.33 | Endoplasmic reticulum |
| Eg09_t010710 | *EgF3'5'H10* | 506 | 56233.7 | 8.59 | Endoplasmic reticulum |
| Eg06_t006150 | *EgFLS1* | 302 | 33251.6 | 6.35 | Cytoplasm\|Nucleus |
| Eg01_t007540 | *EgFLS2* | 334 | 37412.3 | 5.14 | Cytoplasm |
| Eg08_t007020 | *EgFLS3* | 336 | 37815.4 | 6.19 | Cytoplasm\|Nucleus |
| Eg03_t031640 | *EgFLS4* | 334 | 37332.6 | 6.63 | Cytoplasm |
| Eg09_t036940 | *EgDFR1* | 343 | 38149.3 | 5.24 | Cytoplasm |
| Eg04_t027830 | *EgDFR2* | 360 | 40467 | 5.98 | Cytoplasm |
| Eg03_t018610 | *EgDFR3* | 299 | 34129.2 | 5.42 | Cytoplasm |
| Eg05_t014690 | *EgLDOX* | 361 | 40632.4 | 5.66 | Cytoplasm |
| Eg08_t008340 | *EgUGT78D2.1* | 457 | 47910.6 | 6.65 | Cytoplasm |
| Eg06_t008220 | *EgUGT78D2.2* | 459 | 47656.6 | 6.39 | Cytoplasm |
| Eg01_t005290 | *EgUGT78D2.3* | 460 | 48329.3 | 6.29 | Cytoplasm |
| Eg08_t003590 | *EgUGT75C1.1* | 477 | 52031.3 | 5.1 | Cytoplasm |
| Eg08_t037030 | *EgUGT75C1.2* | 495 | 54906.7 | 5.88 | Cytoplasm |

| | | | | | |
|---|---|---|---|---|---|
| Eg08_t037040 | *EgUGT75C1.3* | 474 | 52377.2 | 5.17 | Cytoplasm |
| Eg07_t037900 | *EgUGT79B1* | 472 | 51214.8 | 6.67 | Cytoplasm |
| Eg05_t019870 | *EgLAR* | 355 | 38258.2 | 6.84 | Cytoplasm |
| Eg06_t013840 | *EgANR1* | 333 | 36445.3 | 5.92 | Cytoplasm |
| Eg02_t015210 | *EgANR2* | 343 | 37129.4 | 6.07 | Cytoplasm |
| Eg04_t020700 | *EgTT8.1* | 651 | 73039.9 | 5.52 | Nucleus |
| Eg03_t004200 | *EgTT8.2* | 642 | 71979.5 | 6.13 | Nucleus |
| Eg05_t009200 | *EgTTG1.1* | 385 | 41845.3 | 4.67 | Cytoplasm\|Nucleus |
| Eg09_t041330 | *EgTTG1.2* | 342 | 37149.2 | 4.75 | Cytoplasm\|Nucleus |
| Eg07_t019620 | *EgC1.1* | 261 | 28848.4 | 8.59 | Nucleus |
| Eg08_t035970 | *EgC1.2* | 279 | 30806.5 | 7.97 | Nucleus |

**Table S2d**. The information of flavonoid biosynthetic genes identified in *M. acuminata.*

| **ID** | **Name** | **length** | **MW(Da)** | **pI** | **Localizations** |
|---|---|---|---|---|---|
| Macma4_01_g04800.1 | *MaPAL1* | 713 | 77111.5 | 6.15 | Cytoplasm |
| Macma4_05_g03790.1 | *MaPAL2* | 710 | 76888.9 | 6.68 | Cytoplasm\|Nucleus |
| Macma4_11_g15930.1 | *MaPAL3* | 821 | 89696 | 6.86 | Cytoplasm\|Nucleus |
| Macma4_02_g02760.1 | *MaPAL4* | 712 | 76823.8 | 6 | Cytoplasm |
| Macma4_05_g21700.1 | *MaPAL5* | 712 | 76936.3 | 6.39 | Cytoplasm |
| Macma4_11_g22170.1 | *MaPAL6* | 714 | 77034.2 | 6.3 | Cytoplasm |
| Macma4_08_g14800.1 | *MaPAL7* | 710 | 76913.9 | 6.18 | Cytoplasm |
| Macma4_09_g16190.1 | *MaPAL8* | 783 | 85950.4 | 6.31 | Plastid |
| Macma4_10_g16840.1 | *MaC4H1* | 504 | 57549.5 | 9.6 | Endoplasmic reticulum |
| Macma4_07_g20380.1 | *MaC4H2* | 504 | 57454.5 | 9.77 | Endoplasmic reticulum |
| Macma4_06_g10450.1 | *MaC4H3* | 504 | 57587.5 | 9.34 | Endoplasmic reticulum |

| | | | | | |
|---|---|---|---|---|---|
| Macma4_09_g04800.1 | *MaC4H4* | 504 | 57918.9 | 8.66 | Endoplasmic reticulum |
| Macma4_08_g13040.1 | *Ma4CL1.1* | 544 | 58411.8 | 7.94 | Peroxisome |
| Macma4_05_g22890.1 | *Ma4CL1.2* | 545 | 58653.3 | 7.58 | Peroxisome |
| Macma4_02_g21360.1 | *Ma4CL2.1* | 916 | 100037.3 | 6.17 | Cytoplasm |
| Macma4_11_g22750.1 | *Ma4CL2.2* | 569 | 61158.2 | 5.26 | Cytoplasm |
| Macma4_11_g14090.1 | *Ma4CL3* | 341 | 37269.2 | 9 | Peroxisome |
| Macma4_08_g13960.1 | *Ma4CL4* | 553 | 59587.1 | 9.25 | Peroxisome |
| Macma4_05_g28920.1 | *Ma4CL5* | 555 | 59148.5 | 6.73 | Peroxisome |
| Macma4_11_g06130.1 | *Ma4CL6* | 576 | 61756.7 | 8.84 | Peroxisome |
| Macma4_01_g07350.1 | *Ma4CL7* | 556 | 59390.9 | 8.31 | Peroxisome |
| Macma4_03_g14310.1 | *Ma4CL8* | 545 | 58542.2 | 8.03 | Peroxisome |
| Macma4_06_g39580.1 | *Ma4CL9* | 543 | 58654.2 | 6.2 | Cytoplasm |
| Macma4_01_g19800.1 | *Ma4CL10* | 549 | 59198.7 | 5 | Cytoplasm |
| Macma4_03_g13930.1 | *Ma4CL11* | 656 | 71400.8 | 5.86 | Endoplasmic reticulum |
| Macma4_04_g06110.1 | *Ma4CL12* | 544 | 59039.5 | 5.94 | Cytoplasm |
| Macma4_04_g13130.1 | *Ma4CL13* | 544 | 59221.7 | 5.79 | Cytoplasm |
| Macma4_07_g05220.1 | *Ma4CL14* | 544 | 58642.1 | 5.64 | Cytoplasm |
| Macma4_07_g18890.1 | *Ma4CL15* | 563 | 59412.6 | 6.42 | Peroxisome |
| Macma4_09_g24760.1 | *Ma4CL16* | 544 | 59345.8 | 5.28 | Cytoplasm |
| Macma4_04_g18950.1 | *MaCHI1* | 237 | 25003.3 | 4.72 | Cytoplasm |
| Macma4_11_g22930.1 | *MaCHI2* | 212 | 23695 | 4.72 | Cytoplasm |
| Macma4_06_g12560.1 | *MaCHS1* | 434 | 47100.1 | 7.78 | Cytoplasm |
| Macma4_06_g18170.1 | *MaCHS2* | 416 | 45664.5 | 6.98 | Cytoplasm |
| Macma4_10_g16190.1 | *MaCHS3* | 393 | 42939.3 | 6.61 | Cytoplasm |
| Macma4_06_g10020.1 | *MaCHS4* | 407 | 44537.1 | 7.27 | Cytoplasm |

| | | | | | |
|---|---|---|---|---|---|
| Macma4_06_g10010.1 | *MaCHS5* | 424 | 46536.2 | 6.86 | Cytoplasm |
| Macma4_10_g16140.1 | *MaCHS6* | 393 | 42888.2 | 6.79 | Cytoplasm |
| Macma4_07_g17110.1 | *MaF3H1* | 374 | 41382.5 | 4.92 | Cytoplasm |
| Macma4_02_g06130.1 | *MaF3H2* | 373 | 41427.9 | 5.11 | Cytoplasm |
| Macma4_03_g33170.1 | *MaF3'H* | 542 | 59767.2 | 8.01 | Endoplasmic reticulum |
| Macma4_08_g08290.1 | *MaF3'5'H1* | 504 | 56530.4 | 9.61 | Endoplasmic reticulum |
| Macma4_11_g05110.1 | *MaF3'5'H2* | 505 | 55839.7 | 9.38 | Endoplasmic reticulum |
| Macma4_08_g05480.1 | *MaF3'5'H3* | 506 | 56055.8 | 7.85 | Endoplasmic reticulum |
| Macma4_09_g17660.1 | *MaF3'5'H4* | 506 | 56302.8 | 6.7 | Endoplasmic reticulum |
| Macma4_10_g01240.1 | *MaF3'5'H5* | 453 | 49898.5 | 9.01 | Endoplasmic reticulum |
| Macma4_10_g01260.1 | *MaF3'5'H6* | 453 | 49897.4 | 8.83 | Endoplasmic reticulum |
| Macma4_02_g03040.1 | *MaF3'5'H7* | 417 | 45829.5 | 8.23 | Endoplasmic reticulum |
| Macma4_02_g03240.1 | *MaF3'5'H8* | 506 | 56237 | 7.97 | Endoplasmic reticulum |
| Macma4_02_g03250.1 | *MaF3'5'H9* | 506 | 56182.9 | 7.97 | Endoplasmic reticulum |
| Macma4_02_g03260.1 | *MaF3'5'H10* | 209 | 23585.2 | 6.16 | Cytoplasm |
| Macma4_02_g03290.1 | *MaF3'5'H11* | 506 | 56011.6 | 7.7 | Endoplasmic reticulum |
| Macma4_02_g03300.1 | *MaF3'5'H12* | 573 | 63705 | 8.86 | Peroxisome |
| Macma4_11_g15260.1 | *MaF3'5'H13* | 506 | 56128.6 | 9.25 | Endoplasmic reticulum |
| Macma4_03_g07220.1 | *MaFLS1* | 334 | 37416.5 | 5.79 | Cytoplasm |
| Macma4_08_g24690.1 | *MaFLS2* | 334 | 37063 | 5.93 | Cytoplasm |
| Macma4_04_g33220.1 | *MaDFR1* | 353 | 39360.9 | 6.13 | Cytoplasm |
| Macma4_03_g33910.1 | *MaDFR2* | 357 | 40207.7 | 4.89 | Cytoplasm |
| Macma4_04_g11180.1 | *MaDFR3* | 359 | 40477.2 | 6.35 | Cytoplasm |
| Macma4_05_g03920.1 | *MaLDOX* | 361 | 40674.6 | 5.68 | Cytoplasm |
| Macma4_10_g27580.1 | *MaUGT78D2.1* | 457 | 47869.8 | 6.99 | Cytoplasm |

| | | | | | |
|---|---|---|---|---|---|
| Macma4_07_g16200.1 | *MaUGT78D2.2* | 459 | 47723.6 | 6.34 | Cytoplasm |
| Macma4_03_g04880.1 | *MaUGT78D2.3* | 460 | 48317.5 | 6.29 | Cytoplasm |
| Macma4_10_g31950.1 | *MaUGT75C1.1* | 479 | 52486.7 | 5.31 | Cytoplasm |
| Macma4_06_g07210.1 | *MaUGT75C1.2* | 474 | 52394.4 | 5.28 | Cytoplasm |
| Macma4_06_g07200.1 | *MaUGT75C1.3* | 505 | 55611.5 | 5.62 | Cytoplasm |
| Macma4_06_g07190.1 | *MaUGT75C1.4* | 470 | 52105.7 | 5.1 | Cytoplasm |
| Macma4_09_g07100.1 | *MaUGT79B1* | 468 | 50614.2 | 7.2 | Cytoplasm |
| Macma4_05_g19560.1 | *MaLAR* | 352 | 37887.7 | 6.78 | Cytoplasm |
| Macma4_07_g10640.1 | *MaANR1* | 339 | 37253.2 | 5.64 | Cytoplasm |
| Macma4_07_g10660.1 | *MaANR2* | 336 | 36926 | 5.62 | Cytoplasm |
| Macma4_07_g10670.1 | *MaANR3* | 339 | 37358.3 | 5.29 | Cytoplasm |
| Macma4_07_g10680.1 | *MaANR4* | 339 | 37203.2 | 5.28 | Cytoplasm |
| Macma4_08_g01440.1 | *MaANR5* | 352 | 37982.2 | 5.76 | Cytoplasm |
| Macma4_06_g25800.1 | *MaTT8.1* | 652 | 72922.9 | 5.39 | Nucleus |
| Macma4_03_g19880.1 | *MaTT8.2* | 642 | 71624 | 5.8 | Nucleus |
| Macma4_05_g10420.1 | *MaTTG1.1* | 379 | 41179.7 | 4.67 | Cytoplasm\|Nucleus |
| Macma4_04_g37870.1 | *MaTTG1.2* | 357 | 38819 | 4.63 | Cytoplasm\|Nucleus |
| Macma4_10_g21630.1 | *MaC1.1* | 279 | 30553.2 | 7.71 | Nucleus |
| Macma4_09_g29590.1 | *MaC1.2* | 261 | 28854.3 | 7.14 | Nucleus |

**Table S2e.** The information of flavonoid biosynthetic genes identified in *M. balbisiana*.

| ID | Name | length | MW(Da) | pI | Localizations |
|---|---|---|---|---|---|
| Mba09_g14610.1 | *MbPAL1* | 758 | 83243.3 | 5.93 | Cytoplasm |
| Mba11_g20430.1 | *MbPAL2* | 784 | 85216.6 | 6.68 | Cytoplasm |
| Mba08_g14430.1 | *MbPAL3* | 710 | 76854 | 6.66 | Cytoplasm\|Nucleus |

| | | | | | |
|---|---|---|---|---|---|
| Mba02_g01510.1 | *MbPAL5* | 495 | 52879.7 | 7.7 | Cytoplasm |
| Mba11_g14220.1 | *MbPAL6* | 683 | 73969.3 | 7.92 | Cytoplasm\|Nucleus |
| Mba05_g13940.1 | *MbPAL7* | 710 | 76894.9 | 6.59 | Cytoplasm\|Nucleus |
| Mba01_g14280.1 | *MbPAL8* | 713 | 77137.6 | 6.36 | Cytoplasm |
| Mba09_g04380.1 | *MbC4H1* | 504 | 57779.6 | 7.96 | Endoplasmic reticulum |
| Mba10_g11450.1 | *MbC4H2* | 470 | 53564 | 10.19 | Endoplasmic reticulum |
| Mba06_g09560.1 | *MbC4H3* | 504 | 57550.5 | 9.34 | Endoplasmic reticulum |
| Mba07_g18380.1 | *MbC4H4* | 504 | 57433.5 | 9.74 | Endoplasmic reticulum |
| Mba05_g21540.1 | *Mb4CL1.1* | 545 | 58665.3 | 7.6 | Peroxisome |
| Mba08_g12720.1 | *Mb4CL1.2* | 479 | 51433 | 9.2 | Peroxisome |
| Mba02_g18560.1 | *Mb4CL2.1* | 916 | 99925.1 | 6.31 | Cytoplasm |
| Mba11_g20940.1 | *Mb4CL2.2* | 569 | 61081.1 | 5.49 | Cytoplasm |
| Mba11_g05710.1 | *Mb4CL3.1* | 1028 | 110546.8 | 8.85 | Lysosome/Vacuole |
| Mba11_g12600.1 | *Mb4CL3.2* | 282 | 30353 | 8.85 | Cytoplasm\|Nucleus |
| Mba08_g13620.1 | *Mb4CL4* | 638 | 69166.8 | 9.39 | Plastid |
| Mba05_g27160.1 | *Mb4CL5* | 551 | 58825 | 6.73 | Peroxisome |
| Mba01_g16650.1 | *Mb4CL7* | 289 | 31830.6 | 9.78 | Peroxisome |
| Mba03_g13490.1 | *Mb4CL8* | 590 | 63467.2 | 8.48 | Peroxisome |
| Mba06_g36630.1 | *Mb4CL9* | 543 | 58869.5 | 6.07 | Cytoplasm |
| Mba01_g28370.1 | *Mb4CL10* | 549 | 59225.7 | 5.07 | Cytoplasm |
| Mba04_g05720.1 | *Mb4CL11* | 544 | 59041.4 | 6.21 | Cytoplasm |
| Mba04_g12340.1 | *Mb4CL12* | 544 | 59147.6 | 5.79 | Cytoplasm |
| Mba07_g04890.1 | *Mb4CL13* | 512 | 55129.1 | 4.98 | Cytoplasm |
| Mba07_g17040.1 | *Mb4CL14* | 563 | 59524.8 | 6.61 | Peroxisome |
| Mba11_g21110.1 | *MbCHI1* | 210 | 23524.8 | 4.72 | Cytoplasm |

| | | | | | |
|---|---|---|---|---|---|
| Mba04_g18040.1 | *MbCHI2* | 205 | 21682.7 | 6.02 | Cytoplasm |
| Mba06_g11590.1 | *MbCHS1* | 393 | 42806 | 6.72 | Cytoplasm |
| Mba06_g16830.1 | *MbCHS2* | 393 | 42950.2 | 6.72 | Cytoplasm |
| Mba10_g10880.1 | *MbCHS3* | 399 | 43644 | 6.52 | Cytoplasm |
| Mba06_g09110.1 | *MbCHS4* | 270 | 29676.9 | 10.08 | Cytoplasm |
| Mba06_g09080.1 | *MbCHS5* | 339 | 37144.6 | 6.29 | Cytoplasm |
| Mba06_g09090.1 | *MbCHS6* | 318 | 34448.5 | 6.17 | Cytoplasm |
| Mba06_g09120.1 | *MbCHS7* | 342 | 37079.5 | 6.11 | Cytoplasm |
| Mba10_g10820.1 | *MbCHS8* | 393 | 42860.1 | 6.6 | Cytoplasm |
| Mba02_g04670.1 | *MbF3H1* | 373 | 41366.9 | 5.22 | Cytoplasm |
| Mba07_g15320.1 | *MbF3H2* | 374 | 41371.6 | 4.92 | Cytoplasm |
| Mba01_g02030.1 | *MbF3'H* | 542 | 59798.1 | 7.08 | Endoplasmic reticulum |
| Mba08_g07950.1 | *MbF3'5'H1* | 504 | 56368.2 | 9.37 | Endoplasmic reticulum |
| Mba11_g04840.1 | *MbF3'5'H2* | 864 | 94437.6 | 8.97 | Endoplasmic reticulum |
| Mba09_g16030.1 | *MbF3'5'H3* | 506 | 56216.6 | 6.7 | Endoplasmic reticulum |
| Mba10_g00740.1 | *MbF3'5'H4* | 457 | 50366 | 8.83 | Endoplasmic reticulum |
| Mba02_g01840.1 | *MbF3'5'H5* | 506 | 56169.8 | 7.68 | Endoplasmic reticulum |
| Mba02_g01880.1 | *MbF3'5'H6* | 323 | 35617.3 | 10.17 | Endoplasmic reticulum |
| Mba02_g01890.1 | *MbF3'5'H7* | 506 | 56224.9 | 8.04 | Endoplasmic reticulum |
| Mba02_g01900.1 | *MbF3'5'H8* | 506 | 56288.1 | 8.24 | Endoplasmic reticulum |
| Mba02_g01910.1 | *MbF3'5'H9* | 506 | 56017.8 | 9.09 | Endoplasmic reticulum |
| Mba02_g01920.1 | *MbF3'5'H10* | 561 | 62396 | 7.91 | Endoplasmic reticulum |
| Mba02_g01940.1 | *MbF3'5'H11* | 209 | 23404 | 6.51 | Cytoplasm |
| Mba02_g01950.1 | *MbF3'5'H12* | 1217 | 134294 | 9.9 | Endoplasmic reticulum |
| Mba11_g13650.1 | *MbF3'5'H13* | 506 | 56182.6 | 9.06 | Endoplasmic reticulum |

| | | | | | |
|---|---|---|---|---|---|
| Mba08_g23630.1 | *MbFLS1* | 302 | 33559.2 | 6.26 | Cytoplasm\|Nucleus |
| Mba10_g22000.1 | *MbFLS2* | 305 | 33869 | 5.83 | Cytoplasm\|Nucleus |
| Mba10_g21980.1 | *MbFLS3* | 336 | 37799.4 | 6.35 | Cytoplasm |
| Mba03_g06940.1 | *MbFLS4* | 334 | 37542.6 | 5.44 | Cytoplasm |
| Mba09_g14060.1 | *MbFLS5* | 336 | 37001.8 | 5.45 | Cytoplasm |
| Mba04_g31800.1 | *MbDFR1* | 322 | 36000.2 | 6.24 | Cytoplasm |
| Mba01_g01310.1 | *MbDFR2* | 357 | 40174.7 | 4.97 | Cytoplasm |
| Mba05_g13780.1 | *MbLDOX* | 413 | 45918.6 | 5.66 | Cytoplasm |
| Mba03_g04830.1 | *MbUGT78D2.1* | 499 | 52468.3 | 6.24 | Cytoplasm |
| Mba10_g20800.1 | *MbUGT78D2.2* | 456 | 47887.8 | 7.47 | Cytoplasm |
| Mba07_g14580.1 | *MbUGT78D2.3* | 459 | 47791.9 | 6.45 | Cytoplasm |
| Mba10_g24560.1 | *MbUGT75C1.1* | 480 | 52673.9 | 5.45 | Cytoplasm |
| Mba06_g06480.1 | *MbUGT75C1.2* | 470 | 51986.6 | 5.08 | Cytoplasm |
| Mba06_g06490.1 | *MbUGT75C1.3* | 477 | 52963.9 | 5.12 | Cytoplasm |
| Mba05_g18490.1 | *MbLAR* | 355 | 38226.2 | 6.83 | Cytoplasm |
| Mba08_g01400.1 | *MbANR* | 352 | 38139.4 | 5.77 | Cytoplasm |
| Mba06_g23950.1 | *MbTT8.1* | 197 | 21951.2 | 8.92 | Cytoplasm\|Nucleus |
| Mba03_g18730.1 | *MbTT8.2* | 173 | 19324.2 | 8.47 | Nucleus |
| Mba05_g07670.1 | *MbTTG1.1* | 402 | 44000 | 4.77 | Cytoplasm\|Nucleus |
| Mba04_g36060.1 | *MbTTG1.2* | 383 | 41633.2 | 4.77 | Cytoplasm\|Nucleus |
| Mba10_g15520.1 | *MbC1.1* | 279 | 30582.2 | 7.18 | Nucleus |
| Mba06_g05630.1 | *MbC1.2* | 276 | 30433 | 8.23 | Nucleus |
| Mba09_g23680.1 | *MbC1.3* | 228 | 24926.7 | 4.68 | Nucleus |

**Table S2f.** The information of flavonoid biosynthetic genes identified in *M. schizocharpa*

| ID | Name | length | MW(Da) | pI | Localizations |
|---|---|---|---|---|---|
| Ms01t004140.1 | *MsPAL1* | 713 | 77111.6 | 6.26 | Cytoplasm |
| Ms05t120010.1 | *MsPAL2* | 710 | 76874.9 | 6.68 | Cytoplasm\|Nucleus |
| Ms11t313180.1 | *MsPAL3* | 644 | 69769 | 6.36 | Cytoplasm\|Nucleus |
| Ms02t024050.1 | *MsPAL4* | 712 | 76767.6 | 5.77 | Cytoplasm |
| Ms08t220210.1 | *MsPAL5* | 710 | 76899.1 | 6.56 | Cytoplasm |
| Ms11t319150.1 | *MsPAL6* | 714 | 77060.2 | 6.39 | Cytoplasm |
| Ms05t135820.1 | *MsPAL7* | 712 | 76757.1 | 6.47 | Cytoplasm |
| Ms09t254450.1 | *MsPAL8* | 758 | 83149.1 | 5.7 | Cytoplasm |
| Ms10t281630.1 | *MsC4H1* | 505 | 57631.6 | 9.49 | Endoplasmic reticulum |
| Ms07t197940.1 | *MsC4H2* | 504 | 57465.5 | 9.75 | Endoplasmic reticulum |
| Ms09t244050.1 | *MsC4H3* | 504 | 57960.9 | 8.66 | Endoplasmic reticulum |
| Ms08t218760.1 | *Ms4CL1.1* | 544 | 58385.8 | 7.63 | Peroxisome |
| Ms05t136800.1 | *Ms4CL1.2* | 545 | 58688.2 | 7.59 | Peroxisome |
| Ms02t041000.1 | *Ms4CL2.1* | 540 | 58326.1 | 6.46 | Cytoplasm |
| Ms11t319670.1 | *Ms4CL2.2* | 569 | 61167.2 | 5.37 | Cytoplasm |
| Ms11t311430.1 | *Ms4CL3* | 341 | 37481.3 | 9.29 | Peroxisome |
| Ms08t219500.1 | *Ms4CL4* | 550 | 59269.7 | 9.26 | Peroxisome |
| Ms05t142250.1 | *Ms4CL5* | 587 | 62546.4 | 7.36 | Peroxisome |
| Ms11t304430.1 | *Ms4CL6* | 559 | 61313.7 | 9.28 | Peroxisome |
| Ms01t006680.1 | *Ms4CL7* | 572 | 61161.8 | 7.4 | Peroxisome |
| Ms03t059300.1 | *Ms4CL8* | 528 | 56948.5 | 8.66 | Peroxisome |
| Ms06t178240.1 | *Ms4CL9* | 543 | 58759.4 | 6.04 | Cytoplasm |
| Ms01t017810.1 | *Ms4CL10* | 549 | 59239.8 | 5.16 | Cytoplasm |
| Ms03t058920.1 | *Ms4CL11* | 549 | 59491.1 | 5.27 | Cytoplasm |

Musaceae flavonoid biosynthetic genes

| | | | | | |
|---|---|---|---|---|---|
| Ms04t083350.1 | *Ms4CL12* | 544 | 59032.5 | 6.07 | Cytoplasm |
| Ms04t089570.1 | *Ms4CL13* | 544 | 59155.6 | 5.51 | Cytoplasm |
| Ms07t183930.1 | *Ms4CL14* | 544 | 58730.2 | 5.5 | Cytoplasm |
| Ms07t196560.1 | *Ms4CL15* | 512 | 53831.1 | 6.12 | Cytoplasm |
| Ms09t262230.1 | *Ms4CL16* | 544 | 59429 | 5.28 | Cytoplasm |
| Ms11t319860.1 | *MsCHI1* | 288 | 32488.4 | 6.81 | Plastid |
| Ms04t093250.1 | *MsCHI2* | 237 | 25028.4 | 5.12 | Cytoplasm |
| Ms10t280970.1 | *MsCHS1* | 393 | 42878.1 | 6.6 | Cytoplasm |
| Ms06t158650.1 | *MsCHS2* | 392 | 42857.1 | 6.71 | Cytoplasm |
| MsSC51t327370.1 | *MsCHS3* | 369 | 40956.8 | 6.91 | Cytoplasm |
| MsSC51t327380.1 | *MsCHS4* | 410 | 44989.7 | 7.93 | Cytoplasm |
| MsSC66t327440.1 | *MsCHS5* | 389 | 41971.8 | 9.06 | Cytoplasm |
| Ms02t027200.1 | *MsF3H* | 373 | 41440.9 | 5.1 | Cytoplasm |
| Ms03t075990.1 | *MsF3'H* | 523 | 57540.6 | 7.28 | Endoplasmic reticulum |
| Ms08t214430.1 | *MsF3'5'H1* | 445 | 49617.4 | 9.02 | Endoplasmic reticulum |
| Ms09t255750.1 | *MsF3'5'H2* | 474 | 52543.3 | 6.11 | Endoplasmic reticulum |
| Ms08t211840.1 | *MsF3'5'H3* | 506 | 56115.9 | 8 | Endoplasmic reticulum |
| Ms02t024370.1 | *MsF3'5'H4* | 321 | 36104.3 | 5.81 | Plastid |
| Ms02t024400.1 | *MsF3'5'H5* | 506 | 55805.4 | 8.47 | Endoplasmic reticulum |
| Ms02t024410.1 | *MsF3'5'H6* | 421 | 46389.7 | 8.65 | Endoplasmic reticulum |
| Ms02t024420.1 | *MsF3'5'H7* | 476 | 52619.6 | 9.26 | Endoplasmic reticulum |
| Ms11t312500.1 | *MsF3'5'H8* | 506 | 56176.6 | 9.06 | Endoplasmic reticulum |
| Ms08t229420.1 | *MsFLS1* | 334 | 37144.2 | 5.71 | Cytoplasm |
| Ms03t052640.1 | *MsFLS2* | 334 | 37487.6 | 5.61 | Cytoplasm |
| Ms10t292840.1 | *MsFLS3* | 336 | 37491 | 6.45 | Cytoplasm\|Nucleus |

| | | | | | |
|---|---|---|---|---|---|
| Ms04t107810.1 | *MsDFR1* | 353 | 39425.1 | 6.18 | Cytoplasm |
| Ms04t087800.1 | *MsDFR2* | 354 | 39767.3 | 6.14 | Cytoplasm |
| Ms03t076720.1 | *MsDFR3* | 357 | 40143.6 | 4.89 | Cytoplasm |
| Ms05t120140.1 | *MsLDOX* | 361 | 40674.6 | 5.68 | Cytoplasm |
| Ms10t291590.1 | *MsUGT78D2.1* | 457 | 48126.1 | 6.99 | Cytoplasm |
| Ms07t193970.1 | *MsUGT78D2.2* | 455 | 47311 | 6.23 | Cytoplasm |
| Ms03t050760.1 | *MsUGT78D2.3* | 460 | 48360.4 | 5.97 | Cytoplasm |
| Ms06t152710.1 | *MsUGT75C1.1* | 470 | 52002.5 | 5.09 | Cytoplasm |
| Ms06t152720.1 | *MsUGT75C1.2* | 470 | 51872.5 | 5.37 | Cytoplasm |
| Ms06t152730.1 | *MsUGT75C1.3* | 474 | 52482.5 | 5.51 | Cytoplasm |
| Ms10t295740.1 | *MsUGT75C1.4* | 471 | 51659.9 | 5.55 | Cytoplasm |
| Ms09t246270.1 | *MsUGT79B1* | 471 | 51074.6 | 6.67 | Cytoplasm |
| Ms05t133790.1 | *MsLAR* | 325 | 35058.6 | 6.62 | Cytoplasm |
| Ms08t207990.1 | *MsANR1* | 354 | 38292.7 | 7.42 | Cytoplasm |
| Ms07t188920.1 | *MsANR2* | 351 | 38307.5 | 5.61 | Cytoplasm |
| Ms07t188910.1 | *MsANR3* | 339 | 37321.2 | 5.15 | Cytoplasm |
| MsSC186t326890.1 | *MsTT8.1* | 385 | 42678.3 | 4.39 | Cytoplasm\|Nucleus |
| Ms03t064490.1 | *MsTT8.2* | 565 | 62784.4 | 6.42 | Nucleus |
| Ms04t112120.1 | *MsTTG1.1* | 341 | 36987.1 | 4.75 | Cytoplasm\|Nucleus |
| Ms05t125970.1 | *MsTTG1.2* | 356 | 38547.9 | 4.63 | Cytoplasm\|Nucleus |
| Ms09t266370.1 | *MsC1.1* | 261 | 29000.5 | 8.07 | Nucleus |
| Ms06t151760.1 | *MsC1.2* | 276 | 30410 | 8.02 | Nucleus |

**Table S3a.** Divergence time, Ka, Ks and Ka/Ks values of orthologous flavonoid biosynthetic genes pairs between *E. glaucum* and *M. acuminata*

| **Seq_1** | **Name1** | **Seq_2** | **Name2** | **Ka** | **Ks** | **Ka_Ks** | **Diversity_year** |
|---|---|---|---|---|---|---|---|
| Eg02_t002700 | Eg4CL1 | Parent=Macma4_08_g13040 | Ma4CL1 | 0.030448156 | 0.087782748 | 0.346858085 | 19.50727735 |
| Eg05_t023200 | Eg4CL1.2 | Parent=Macma4_05_g22890 | Ma4CL1.2 | 0.028867818 | 0.102914267 | 0.280503555 | 22.86983716 |
| Eg01_t014520 | Eg4CL10 | Parent=Macma4_01_g19800 | Ma4CL10 | 0.050934779 | 0.351209955 | 0.145026581 | 78.04665669 |
| Eg01_t014520 | Eg4CL10 | Parent=Macma4_03_g13930 | Ma4CL11 | 0.022718058 | 0.088092637 | 0.257888276 | 19.57614153 |
| Eg01_t025930 | Eg4CL11 | Parent=Macma4_01_g19800 | Ma4CL10 | 0.012306286 | 0.089448569 | 0.137579456 | 19.87745987 |
| Eg01_t025930 | Eg4CL11 | Parent=Macma4_03_g13930 | Ma4CL11 | 0.04613291 | 0.361629763 | 0.127569452 | 80.36216945 |
| Eg01_t025930 | Eg4CL11 | Parent=Macma4_07_g05220 | Ma4CL14 | 0.083495624 | 0.454381606 | 0.183756612 | 100.9736902 |
| Eg04_t025780 | Eg4CL12 | Parent=Macma4_04_g13130 | Ma4CL13 | 0.013220751 | 0.08282486 | 0.159622984 | 18.40552448 |
| Eg04_t025780 | Eg4CL12 | Parent=Macma4_04_g06110 | Ma4CL12 | 0.042717676 | 0.361825668 | 0.118061485 | 80.40570393 |
| Eg04_t025780 | Eg4CL12 | Parent=Macma4_09_g24760 | Ma4CL16 | 0.057442296 | 0.416504423 | 0.137915213 | 92.55653853 |
| Eg04_t032900 | Eg4CL13 | Parent=Macma4_04_g06110 | Ma4CL12 | 0.015828425 | 0.091971342 | 0.172101707 | 20.43807607 |
| Eg04_t032900 | Eg4CL13 | Parent=Macma4_04_g13130 | Ma4CL13 | 0.049643453 | 0.336560674 | 0.147502239 | 74.79126088 |
| Eg07_t001640 | Eg4CL14 | Parent=Macma4_07_g18890 | Ma4CL15 | 0.033652834 | 0.109779242 | 0.30655007 | 24.39538711 |
| Eg07_t014740 | Eg4CL15 | Parent=Macma4_09_g24760 | Ma4CL16 | 0.020341637 | 0.095221058 | 0.213625405 | 21.16023509 |
| Eg09_t018540 | Eg4CL2.2 | Parent=Macma4_11_g22750 | Ma4CL2.2 | 0.011830676 | 0.063646732 | 0.185880335 | 14.14371818 |
| Eg09_t002410 | Eg4CL3 | Parent=Macma4_11_g06130 | Ma4CL6 | 0.046396851 | 0.103003864 | 0.450437966 | 22.88974756 |
| Eg02_t001820 | Eg4CL4 | Parent=Macma4_08_g13960 | Ma4CL4 | 0.014301169 | 0.092773033 | 0.154152224 | 20.6162295 |
| Eg05_t033420 | Eg4CL5 | Parent=Macma4_05_g28920 | Ma4CL5 | 0.021000952 | 0.096247208 | 0.218198039 | 21.38826854 |
| Eg01_t038400 | Eg4CL7 | Parent=Macma4_01_g07350 | Ma4CL7 | 0.022725881 | 0.120256988 | 0.188977637 | 26.72377501 |
| Eg03_t041750 | Eg4CL8 | Parent=Macma4_03_g14310 | Ma4CL8 | 0.033998936 | 0.107672474 | 0.315762565 | 23.92721635 |
| Eg04_t006840 | Eg4CL9 | Parent=Macma4_06_g39580 | Ma4CL9 | 0.024011986 | 0.076482549 | 0.31395379 | 16.9961219 |
| Eg06_t013840 | EgANR1 | Parent=Macma4_07_g10660 | MaANR2 | 0.051307976 | 0.11652155 | 0.440330357 | 25.89367784 |

| | | | | | | | |
|---|---|---|---|---|---|---|---|
| Eg02_t015210 | EgANR2 | Parent=Macma4_08_g01440 | MaANR5 | 0.029759249 | 0.067715135 | 0.439477065 | 15.04780778 |
| Eg07_t019620 | EgC1 | Parent=Macma4_09_g29590 | MaC1.2 | 0.047662584 | 0.095228596 | 0.500507049 | 21.16191029 |
| Eg08_t019660 | EgC4H1 | Parent=Macma4_06_g10450 | MaC4H3 | 0.033843676 | 0.295141444 | 0.114669344 | 65.58698754 |
| Eg08_t019660 | EgC4H1 | Parent=Macma4_10_g16840 | MaC4H1 | 0.007887314 | 0.075517183 | 0.104443969 | 16.78159633 |
| Eg07_t003130 | EgC4H2 | Parent=Macma4_07_g20380 | MaC4H2 | 0.009631654 | 0.088445917 | 0.108898794 | 19.65464829 |
| Eg08_t048930 | EgC4H3 | Parent=Macma4_06_g10450 | MaC4H3 | 0.012307969 | 0.084679048 | 0.145348452 | 18.81756618 |
| Eg08_t048930 | EgC4H3 | Parent=Macma4_10_g16840 | MaC4H1 | 0.038660467 | 0.355247937 | 0.108826718 | 78.94398603 |
| Eg07_t034110 | EgC4H4 | Parent=Macma4_09_g04800 | MaC4H4 | 0.01400274 | 0.086319142 | 0.162220571 | 19.18203146 |
| Eg01_t015550 | EgCHI1 | Parent=Macma4_04_g18950 | MaCHI1 | 0.052121717 | 0.066655272 | 0.781959408 | 14.81228265 |
| Eg09_t018750 | EgCHI2 | Parent=Macma4_11_g22930 | MaCHI2 | 0.01648418 | 0.105436463 | 0.156342309 | 23.4303251 |
| Eg08_t046640 | EgCHS1 | Parent=Macma4_06_g12560 | MaCHS1 | 0.029252417 | 0.075016776 | 0.389945011 | 16.67039458 |
| Eg08_t040870 | EgCHS2 | Parent=Macma4_06_g18170 | MaCHS2 | 0.017616114 | 0.060086713 | 0.293178197 | 13.35260298 |
| Eg08_t040870 | EgCHS2 | Parent=Macma4_06_g10010 | MaCHS5 | 0.106008972 | 0.350319314 | 0.302606701 | 77.84873651 |
| Eg08_t040870 | EgCHS2 | Parent=Macma4_10_g16140 | MaCHS6 | 0.032099655 | 0.329646032 | 0.097376131 | 73.25467381 |
| Eg08_t020390 | EgCHS3 | Parent=Macma4_06_g18170 | MaCHS2 | 0.041153619 | 0.29886016 | 0.137701924 | 66.41336893 |
| Eg08_t020390 | EgCHS3 | Parent=Macma4_10_g16140 | MaCHS6 | 0.011301777 | 0.069946424 | 0.161577628 | 15.5436497 |
| Eg08_t049400 | EgCHS5 | Parent=Macma4_06_g10010 | MaCHS5 | 0.028067422 | 0.079020387 | 0.35519216 | 17.56008594 |
| Eg08_t049400 | EgCHS5 | Parent=Macma4_06_g18170 | MaCHS2 | 0.102873973 | 0.357535564 | 0.287730743 | 79.45234746 |
| Eg08_t049400 | EgCHS5 | Parent=Macma4_10_g16140 | MaCHS6 | 0.107293231 | 0.373124864 | 0.287553153 | 82.91663637 |
| Eg09_t036940 | EgDFR1 | Parent=Macma4_04_g33220 | MaDFR1 | 0.020420277 | 0.106454841 | 0.191821023 | 23.65663143 |
| Eg04_t027830 | EgDFR2 | Parent=Macma4_04_g11180 | MaDFR3 | 0.023229241 | 0.068816148 | 0.337555086 | 15.29247727 |
| Eg09_t020640 | EgF3'5'H1 | Parent=Macma4_11_g05110 | MaF3'5'H2 | 0.04171204 | 0.104813195 | 0.397965545 | 23.29182113 |
| Eg09_t010710 | EgF3'5'H10 | Parent=Macma4_11_g15260 | MaF3'5'H13 | 0.020904177 | 0.074203628 | 0.281713683 | 16.48969508 |
| Eg02_t007770 | EgF3'5'H2 | Parent=Macma4_08_g08290 | MaF3'5'H1 | 0.019630833 | 0.097024186 | 0.202329272 | 21.5609302 |
| Eg06_t004020 | EgF3'5'H3 | Parent=Macma4_09_g17660 | MaF3'5'H4 | 0.023059979 | 0.113561102 | 0.203062306 | 25.23580042 |

Musaceae flavonoid biosynthetic genes

| | | | | | | | |
|---|---|---|---|---|---|---|---|
| Eg07_t042500 | EgF3'5'H4 | Parent=Macma4_09_g17660 | MaF3'5'H4 | 0.124106248 | 0.464327144 | 0.267281915 | 103.1838097 |
| Eg07_t042500 | EgF3'5'H4 | Parent=Macma4_10_g01260 | MaF3'5'H6 | 0.020876099 | 0.10088637 | 0.206926854 | 22.41919342 |
| Eg02_t010650 | EgF3'5'H5 | Parent=Macma4_08_g05480 | MaF3'5'H3 | 0.02504705 | 0.072077584 | 0.347501246 | 16.01724098 |
| Eg02_t019070 | EgF3'5'H6 | Parent=Macma4_02_g03040 | MaF3'5'H7 | 0.050319405 | 0.165735594 | 0.303612543 | 36.83013189 |
| Eg02_t019070 | EgF3'5'H6 | Parent=Macma4_02_g03290 | MaF3'5'H11 | 0.03908655 | 0.17604095 | 0.222031012 | 39.12021122 |
| Eg02_t019070 | EgF3'5'H6 | Parent=Macma4_02_g03300 | MaF3'5'H12 | 0.17390456 | 0.592788456 | 0.293366982 | 131.7307679 |
| Eg02_t019110 | EgF3'5'H8 | Parent=Macma4_02_g03260 | MaF3'5'H10 | 0.019112696 | 0.125522379 | 0.152265246 | 27.89386197 |
| Eg07_t027880 | EgF3H1 | Parent=Macma4_07_g17110 | MaF3H1 | 0.021999199 | 0.151758777 | 0.144961625 | 33.72417276 |
| Eg03_t018110 | EgF3'H1 | Parent=Macma4_03_g33170 | MaF3'H | 0.021851509 | 0.07919438 | 0.275922472 | 17.59875122 |
| Eg02_t022060 | EgF3H2 | Parent=Macma4_02_g06130 | MaF3H2 | 0.012978399 | 0.089385328 | 0.145196076 | 19.86340612 |
| Eg01_t007540 | EgFLS2 | Parent=Macma4_03_g07220 | MaFLS1 | 0.017187047 | 0.090779349 | 0.189327719 | 20.17318876 |
| Eg03_t031640 | EgFLS4 | Parent=Macma4_08_g24690 | MaFLS2 | 0.022583065 | 0.071429408 | 0.316159215 | 15.87320167 |
| Eg05_t019870 | EgLAR | Parent=Macma4_05_g19560 | MaLAR | 0.023227663 | 0.096131395 | 0.241624111 | 21.36253224 |
| Eg05_t014690 | EgLDOX | Parent=Macma4_05_g03920 | MaLDOX | 0.020728814 | 0.067319162 | 0.307918476 | 14.95981384 |
| Eg01_t040920 | EgPAL1 | Parent=Macma4_01_g04800 | MaPAL1 | 0.012837552 | 0.079218668 | 0.162052104 | 17.60414851 |
| Eg05_t014880 | EgPAL2 | Parent=Macma4_05_g03790 | MaPAL2 | 0.010621858 | 0.088488942 | 0.120035996 | 19.66420934 |
| Eg02_t018790 | EgPAL4 | Parent=Macma4_02_g02760 | MaPAL4 | 0.021509118 | 0.105258317 | 0.204346017 | 23.39073706 |
| Eg02_t018790 | EgPAL4 | Parent=Macma4_08_g14800 | MaPAL7 | 0.058940904 | 0.401943438 | 0.146639796 | 89.3207639 |
| Eg09_t011330 | EgPAL5 | Parent=Macma4_02_g02760 | MaPAL4 | 0.066969936 | 0.380667201 | 0.175927782 | 84.5927114 |
| Eg09_t011330 | EgPAL5 | Parent=Macma4_08_g14800 | MaPAL7 | 0.081008797 | 0.385794925 | 0.209978907 | 85.73220561 |
| Eg09_t011330 | EgPAL5 | Parent=Macma4_11_g15930 | MaPAL3 | 0.02180223 | 0.081532772 | 0.267404497 | 18.11839375 |
| Eg02_t001010 | EgPAL6 | Parent=Macma4_02_g02760 | MaPAL4 | 0.066617058 | 0.381236394 | 0.174739502 | 84.71919863 |
| Eg02_t001010 | EgPAL6 | Parent=Macma4_08_g14800 | MaPAL7 | 0.014431063 | 0.058203717 | 0.247940577 | 12.93415936 |
| Eg09_t017980 | EgPAL7 | Parent=Macma4_05_g21700 | MaPAL5 | 0.043020243 | 0.376043694 | 0.114402247 | 83.56526526 |
| Eg09_t017980 | EgPAL7 | Parent=Macma4_11_g22170 | MaPAL6 | 0.010017366 | 0.085679392 | 0.116916866 | 19.03986485 |

Musaceae flavonoid biosynthetic genes

| | | | | | | | |
|---|---|---|---|---|---|---|---|
| Eg05_t024420 | EgPAL8 | Parent=Macma4_05_g21700 | MaPAL5 | 0.013827449 | 0.063289053 | 0.218480891 | 14.06423403 |
| Eg06_t005550 | EgPAL9 | Parent=Macma4_09_g16190 | MaPAL8 | 0.008264953 | 0.062411775 | 0.132426182 | 13.86928325 |
| Eg04_t020700 | EgTT8 | Parent=Macma4_06_g25800 | MaTT8 | 0.024548516 | 0.092329963 | 0.265878109 | 20.51776961 |
| Eg03_t004200 | EgTT8.2 | Parent=Macma4_03_g19880 | MaTT8.2 | 0.03276383 | 0.132223082 | 0.247792063 | 29.38290703 |
| Eg05_t009200 | EgTTG1 | Parent=Macma4_05_g10420 | MaTTG1 | 0.016593269 | 0.098783089 | 0.167976822 | 21.95179757 |
| Eg09_t041330 | EgTTG1.2 | Parent=Macma4_04_g37870 | MaTTG1.2 | 0.015821285 | 0.124125967 | 0.127461528 | 27.58354822 |
| Eg08_t037030 | EgUGT75C1.2 | Parent=Macma4_06_g07200 | MaUGT75C1.3 | 0.07564968 | 0.159282386 | 0.474940653 | 35.39608579 |
| Eg08_t037040 | EgUGT75C1.3 | Parent=Macma4_06_g07210 | MaUGT75C1.2 | 0.039690535 | 0.080912142 | 0.490538682 | 17.98047597 |
| Eg08_t008340 | EgUGT78D2 | Parent=Macma4_10_g27580 | MaUGT78D2 | 0.046476545 | 0.127836889 | 0.363561295 | 28.40819753 |
| Eg06_t008220 | EgUGT78D2.2 | Parent=Macma4_07_g16200 | MaUGT78D2.2 | 0.02597446 | 0.086340581 | 0.300837211 | 19.18679588 |
| Eg01_t005290 | EgUGT78D2.3 | Parent=Macma4_03_g04880 | MaUGT78D2.3 | 0.03218191 | 0.100593663 | 0.319919856 | 22.35414734 |
| Eg07_t037900 | EgUGT79B1 | Parent=Macma4_09_g07100 | MaUGT79B1 | 0.029682758 | 0.075957716 | 0.390780025 | 16.87949236 |

**Table S3b.** Divergence time, Ka, Ks and Ka/Ks values of orthologous flavonoid biosynthetic genes pairs between *E. glaucum* and *M. balbisiana*

| **Seq_1** | **Name1** | **Seq_2** | **Name2** | **Ka** | **Ks** | **Ka_Ks** | **Diversity_year** |
|---|---|---|---|---|---|---|---|
| Eg02_t002700 | Eg4CL1 | Mba08_g12720 | Mb4CL1.2 | 0.027861033 | 0.097001086 | 0.287223923 | 21.55579698 |
| Eg05_t023200 | Eg4CL1.2 | Mba05_g21540 | Mb4CL1 | 0.030607937 | 0.102579665 | 0.298382108 | 22.79548109 |
| Eg01_t014520 | Eg4CL10 | Mba01_g28370 | Mb4CL10 | 0.051097917 | 0.361717414 | 0.141264741 | 80.38164766 |
| Eg01_t025930 | Eg4CL11 | Mba01_g28370 | Mb4CL10 | 0.011483478 | 0.105799832 | 0.108539661 | 23.51107385 |
| Eg04_t025780 | Eg4CL12 | Mba04_g12340 | Mb4CL12 | 0.014885748 | 0.096718544 | 0.153907901 | 21.49300975 |
| Eg04_t025780 | Eg4CL12 | Mba04_g05720 | Mb4CL11 | 0.04620431 | 0.35410953 | 0.130480279 | 78.69100661 |
| Eg04_t032900 | Eg4CL13 | Mba04_g05720 | Mb4CL11 | 0.020048528 | 0.075854067 | 0.264303926 | 16.85645937 |
| Eg04_t032900 | Eg4CL13 | Mba04_g12340 | Mb4CL12 | 0.051385593 | 0.356223906 | 0.144250828 | 79.16086793 |
| Eg07_t001640 | Eg4CL14 | Mba07_g17040 | Mb4CL14 | 0.033655165 | 0.119994493 | 0.28047258 | 26.66544294 |
| Eg07_t014740 | Eg4CL15 | Mba04_g05720 | Mb4CL11 | 0.06433892 | 0.57664255 | 0.111575048 | 128.1427888 |

| | | | | | | | |
|---|---|---|---|---|---|---|---|
| Eg09_t018540 | Eg4CL2.2 | Mba11_g20940 | Mb4CL2.2 | 0.014224053 | 0.05596434 | 0.254162801 | 12.43651995 |
| Eg02_t001820 | Eg4CL4 | Mba08_g13620 | Mb4CL4 | 0.020965115 | 0.113939276 | 0.184002532 | 25.31983903 |
| Eg05_t033420 | Eg4CL5 | Mba05_g27160 | Mb4CL5 | 0.02442664 | 0.085944481 | 0.284214172 | 19.09877361 |
| Eg01_t038400 | Eg4CL7 | Mba01_g16650 | Mb4CL7 | 0.011060108 | 0.075662594 | 0.1461767 | 16.81390977 |
| Eg03_t041750 | Eg4CL8 | Mba03_g13490 | Mb4CL8 | 0.054629005 | 0.126001261 | 0.433559192 | 28.00028025 |
| Eg04_t006840 | Eg4CL9 | Mba06_g36630 | Mb4CL9 | 0.025671841 | 0.071168367 | 0.360719826 | 15.81519269 |
| Eg02_t015210 | EgANR2 | Mba08_g01400 | MbANR | 0.027041839 | 0.063606107 | 0.425145322 | 14.13469045 |
| Eg07_t019620 | EgC1 | Mba09_g23680 | MbC1.3 | 0.181505951 | 0.262138392 | 0.692405068 | 58.25297596 |
| Eg08_t035970 | EgC1.2 | Mba06_g05630 | MbC1.2 | 0.037411197 | 0.13693676 | 0.273200544 | 30.43039116 |
| Eg07_t003130 | EgC4H2 | Mba07_g18380 | MbC4H4 | 0.006991021 | 0.09164931 | 0.076280123 | 20.36651334 |
| Eg08_t048930 | EgC4H3 | Mba06_g09560 | MbC4H3 | 0.011431305 | 0.096885056 | 0.117988321 | 21.53001244 |
| Eg07_t034110 | EgC4H4 | Mba09_g04380 | MbC4H1 | 0.023783555 | 0.07691463 | 0.30922017 | 17.09213992 |
| Eg01_t015550 | EgCHI1 | Mba04_g18040 | MbCHI2 | 0.075505527 | 0.113254769 | 0.666687394 | 25.16772652 |
| Eg09_t018750 | EgCHI2 | Mba11_g21110 | MbCHI1 | 0.018750977 | 0.106507748 | 0.1760527 | 23.6683885 |
| Eg08_t046640 | EgCHS1 | Mba06_g11590 | MbCHS1 | 0.024013485 | 0.057842565 | 0.415152495 | 12.85390337 |
| Eg08_t040870 | EgCHS2 | Mba06_g16830 | MbCHS2 | 0.010176355 | 0.073520709 | 0.138414811 | 16.33793541 |
| Eg08_t040870 | EgCHS2 | Mba10_g10820 | MbCHS8 | 0.028602131 | 0.334742724 | 0.085445118 | 74.38727194 |
| Eg08_t020390 | EgCHS3 | Mba06_g16830 | MbCHS2 | 0.033281421 | 0.318481924 | 0.104500189 | 70.77376093 |
| Eg08_t020390 | EgCHS3 | Mba10_g10820 | MbCHS8 | 0.011304971 | 0.085461979 | 0.132280713 | 18.99155085 |
| Eg08_t049400 | EgCHS5 | Mba06_g09120 | MbCHS7 | 0.020097217 | 0.100039483 | 0.20089285 | 22.23099614 |
| Eg08_t049400 | EgCHS5 | Mba06_g16830 | MbCHS2 | 0.095795195 | 0.366288822 | 0.261529124 | 81.39751606 |
| Eg08_t049400 | EgCHS5 | Mba10_g10820 | MbCHS8 | 0.103446314 | 0.378471643 | 0.273326458 | 84.10480954 |
| Eg09_t036940 | EgDFR1 | Mba04_g31800 | MbDFR1 | 0.017625938 | 0.103740921 | 0.169903431 | 23.05353792 |
| Eg09_t020640 | EgF3'5'H1 | Mba11_g04840 | MbF3'5'H2 | 0.040891344 | 0.114974818 | 0.355654782 | 25.54995961 |
| Eg09_t010710 | EgF3'5'H10 | Mba11_g13650 | MbF3'5'H13 | 0.017749682 | 0.08463644 | 0.209716785 | 18.80809786 |

Musaceae flavonoid biosynthetic genes

| | | | | | | | |
|---|---|---|---|---|---|---|---|
| Eg02_t007770 | EgF3'5'H2 | Mba08_g07950 | MbF3'5'H1 | 0.015117222 | 0.088065101 | 0.171659618 | 19.57002254 |
| Eg06_t004020 | EgF3'5'H3 | Mba09_g16030 | MbF3'5'H3 | 0.024002048 | 0.119205831 | 0.201349616 | 26.49018469 |
| Eg07_t042500 | EgF3'5'H4 | Mba10_g00740 | MbF3'5'H4 | 0.023234681 | 0.085363352 | 0.272185664 | 18.96963374 |
| Eg02_t019070 | EgF3'5'H6 | Mba02_g01950 | MbF3'5'H12 | 0.094948398 | 0.521089619 | 0.182211263 | 115.7976931 |
| Eg02_t019110 | EgF3'5'H8 | Mba02_g01940 | MbF3'5'H11 | 0.0420671 | 0.121290554 | 0.346829152 | 26.95345645 |
| Eg07_t027880 | EgF3H1 | Mba07_g15320 | MbF3H2 | 0.021391824 | 0.168182034 | 0.127194469 | 37.37378523 |
| Eg03_t018110 | EgF3'H1 | Mba01_g02030 | MbF3'H | 0.022735638 | 0.079229271 | 0.286960076 | 17.60650473 |
| Eg02_t022060 | EgF3H2 | Mba02_g04670 | MbF3H1 | 0.012975847 | 0.072369457 | 0.179300047 | 16.08210161 |
| Eg06_t006150 | EgFLS1 | Mba09_g14060 | MbFLS5 | 0.141119552 | 0.204327392 | 0.690654106 | 45.40608714 |
| Eg01_t007540 | EgFLS2 | Mba03_g06940 | MbFLS4 | 0.017871778 | 0.078576468 | 0.227444401 | 17.46143742 |
| Eg01_t007540 | EgFLS2 | Mba09_g14060 | MbFLS5 | 0.152797668 | 0.501688536 | 0.304566792 | 111.4863413 |
| Eg08_t007020 | EgFLS3 | Mba10_g21980 | MbFLS3 | 0.025120779 | 0.094755958 | 0.26511029 | 21.05687949 |
| Eg03_t031640 | EgFLS4 | Mba08_g23630 | MbFLS1 | 0.016049245 | 0.11824346 | 0.135730506 | 26.27632435 |
| Eg05_t019870 | EgLAR | Mba05_g18490 | MbLAR | 0.030210936 | 0.080750184 | 0.374128379 | 17.94448542 |
| Eg05_t014690 | EgLDOX | Mba05_g13780 | MbLDOX | 0.026947173 | 0.085016608 | 0.316963632 | 18.89257956 |
| Eg01_t040920 | EgPAL1 | Mba01_g14280 | MbPAL8 | 0.011264879 | 0.073850538 | 0.152536185 | 16.41123068 |
| Eg05_t014880 | EgPAL2 | Mba05_g13940 | MbPAL7 | 0.015037313 | 0.09511955 | 0.158088559 | 21.13767777 |
| Eg02_t018790 | EgPAL4 | Mba02_g01510 | MbPAL5 | 0.052573992 | 0.130658518 | 0.402377077 | 29.03522614 |
| Eg02_t018790 | EgPAL4 | Mba08_g14430 | MbPAL3 | 0.058947294 | 0.40177258 | 0.146718061 | 89.28279545 |
| Eg09_t011330 | EgPAL5 | Mba02_g01510 | MbPAL5 | 0.089161023 | 0.383754621 | 0.232338631 | 85.27880467 |
| Eg09_t011330 | EgPAL5 | Mba11_g14220 | MbPAL6 | 0.030510176 | 0.104883377 | 0.2908962 | 23.30741718 |
| Eg02_t001010 | EgPAL6 | Mba08_g14430 | MbPAL3 | 0.014432573 | 0.058184244 | 0.248049496 | 12.92983211 |
| Eg09_t017980 | EgPAL7 | Mba11_g20430 | MbPAL2 | 0.011277819 | 0.083599292 | 0.134903286 | 18.57762038 |
| Eg06_t005550 | EgPAL9 | Mba09_g14610 | MbPAL1 | 0.008859243 | 0.070097065 | 0.126385359 | 15.57712554 |
| Eg04_t020700 | EgTT8 | Mba06_g23950 | MbTT8 | 0.082164693 | 0.167282093 | 0.491174466 | 37.17379843 |

Musaceae flavonoid biosynthetic genes

| | | | | | | | |
|---|---|---|---|---|---|---|---|
| Eg03_t004200 | EgTT8.2 | Mba03_g18730 | MbTT8.2 | 0.102720217 | 0.160248606 | 0.641005374 | 35.61080123 |
| Eg03_t004200 | EgTT8.2 | Mba06_g23950 | MbTT8 | 0.106584956 | 0.417975642 | 0.255002794 | 92.88347593 |
| Eg05_t009200 | EgTTG1 | Mba05_g07670 | MbTTG1 | 0.018999631 | 0.106847011 | 0.177820891 | 23.74378025 |
| Eg09_t041330 | EgTTG1.2 | Mba04_g36060 | MbTTG1.2 | 0.015776217 | 0.12833498 | 0.122929984 | 28.51888451 |
| Eg08_t037030 | EgUGT75C1.2 | Mba06_g06480 | MbUGT75C1.2 | 0.063079795 | 0.113481846 | 0.555858032 | 25.21818792 |
| Eg08_t037040 | EgUGT75C1.3 | Mba06_g06490 | MbUGT75C1.3 | 0.038006966 | 0.064169735 | 0.592288025 | 14.25994113 |
| Eg08_t008340 | EgUGT78D2 | Mba10_g20800 | MbUGT78D2.2 | 0.037957164 | 0.098576801 | 0.385051686 | 21.90595588 |
| Eg06_t008220 | EgUGT78D2.2 | Mba07_g14580 | MbUGT78D2.3 | 0.02493788 | 0.074006256 | 0.336969898 | 16.4458346 |
| Eg01_t005290 | EgUGT78D2.3 | Mba03_g04830 | MbUGT78D2 | 0.02910064 | 0.110207788 | 0.264052485 | 24.49061962 |

**Table S3c.** Divergence time, Ka, Ks and Ka/Ks values of orthologous flavonoid biosynthetic genes pairs between *E. glaucum* and *M. schizocharpa*

| **Seq_1** | **Name1** | **Seq_2** | **Name2** | **Ka** | **Ks** | **Ka_Ks** | **Diversity_year** |
|---|---|---|---|---|---|---|---|
| Eg02_t002700 | Eg4CL1 | Ms08t218760 | Ms4CL1 | 0.029593807 | 0.093045064 | 0.318058856 | 20.67668097 |
| Eg05_t023200 | Eg4CL1.2 | Ms05t136800 | Ms4CL1.2 | 0.025479271 | 0.105505243 | 0.24149767 | 23.44560966 |
| Eg01_t014520 | Eg4CL10 | Ms01t017810 | Ms4CL10 | 0.050963394 | 0.362087034 | 0.140749017 | 80.46378532 |
| Eg01_t014520 | Eg4CL10 | Ms03t058920 | Ms4CL11 | 0.023547545 | 0.093651284 | 0.251438572 | 20.81139651 |
| Eg01_t025930 | Eg4CL11 | Ms01t017810 | Ms4CL10 | 0.01231302 | 0.108506079 | 0.113477699 | 24.11246205 |
| Eg01_t025930 | Eg4CL11 | Ms03t058920 | Ms4CL11 | 0.045249323 | 0.366113716 | 0.123593629 | 81.35860356 |
| Eg01_t025930 | Eg4CL11 | Ms07t183930 | Ms4CL14 | 0.083875315 | 0.458289121 | 0.183018341 | 101.842027 |
| Eg04_t025780 | Eg4CL12 | Ms04t089570 | Ms4CL13 | 0.014889853 | 0.088318133 | 0.168593384 | 19.62625169 |
| Eg04_t025780 | Eg4CL12 | Ms04t083350 | Ms4CL12 | 0.045744592 | 0.372479318 | 0.122811094 | 82.77318172 |
| Eg04_t032900 | Eg4CL13 | Ms04t083350 | Ms4CL12 | 0.019193037 | 0.09212845 | 0.208329097 | 20.47298891 |
| Eg04_t032900 | Eg4CL13 | Ms04t089570 | Ms4CL13 | 0.051400151 | 0.344218686 | 0.149324117 | 76.49304125 |
| Eg07_t001640 | Eg4CL14 | Ms07t196560 | Ms4CL15 | 0.041898569 | 0.122895352 | 0.340928832 | 27.3100782 |

| | | | | | | | |
|---|---|---|---|---|---|---|---|
| Eg07_t014740 | Eg4CL15 | Ms04t089570 | Ms4CL13 | 0.048195771 | 0.500649897 | 0.096266416 | 111.2555326 |
| Eg07_t014740 | Eg4CL15 | Ms09t262230 | Ms4CL16 | 0.023704239 | 0.092585382 | 0.256025722 | 20.57452931 |
| Eg02_t037340 | Eg4CL2 | Ms02t041000 | Ms4CL2 | 0.029372329 | 0.093308542 | 0.314787143 | 20.73523147 |
| Eg09_t018540 | Eg4CL2.2 | Ms11t319670 | Ms4CL2.2 | 0.011036132 | 0.063646732 | 0.173396674 | 14.14371818 |
| Eg02_t001820 | Eg4CL4 | Ms08t219500 | Ms4CL4 | 0.015150968 | 0.100408305 | 0.15089357 | 22.31295678 |
| Eg05_t033420 | Eg4CL5 | Ms05t142250 | Ms4CL5 | 0.024419783 | 0.093646658 | 0.260765129 | 20.81036834 |
| Eg01_t038400 | Eg4CL7 | Ms01t006680 | Ms4CL7 | 0.019797685 | 0.108206949 | 0.18296131 | 24.04598865 |
| Eg03_t041750 | Eg4CL8 | Ms03t059300 | Ms4CL8 | 0.040158291 | 0.101935694 | 0.3939571 | 22.65237655 |
| Eg04_t006840 | Eg4CL9 | Ms06t178240 | Ms4CL9 | 0.025694895 | 0.087691004 | 0.293016319 | 19.48688968 |
| Eg06_t013840 | EgANR1 | Ms07t188910 | MsANR3 | 0.051590353 | 0.112628525 | 0.458057613 | 25.02856116 |
| Eg02_t015210 | EgANR2 | Ms08t207990 | MsANR1 | 0.033810409 | 0.063475789 | 0.532650471 | 14.10573086 |
| Eg07_t019620 | EgC1 | Ms09t266370 | MsC1 | 0.05487863 | 0.095091209 | 0.577115702 | 21.13137979 |
| Eg08_t035970 | EgC1.2 | Ms06t151760 | MsC1.2 | 0.045822019 | 0.124285518 | 0.368683493 | 27.61900409 |
| Eg08_t019660 | EgC4H1 | Ms10t281630 | MsC4H1 | 0.009657021 | 0.081495162 | 0.118498093 | 18.11003592 |
| Eg07_t003130 | EgC4H2 | Ms07t197940 | MsC4H2 | 0.009631654 | 0.097892382 | 0.098390228 | 21.7538626 |
| Eg08_t048930 | EgC4H3 | Ms10t281630 | MsC4H1 | 0.036387942 | 0.361054381 | 0.100782441 | 80.2343069 |
| Eg07_t034110 | EgC4H4 | Ms09t244050 | MsC4H3 | 0.015773913 | 0.083129986 | 0.189749975 | 18.47333024 |
| Eg01_t015550 | EgCHI1 | Ms04t093250 | MsCHI2 | 0.050194071 | 0.060140597 | 0.83461211 | 13.3645771 |
| Eg09_t018750 | EgCHI2 | Ms11t319860 | MsCHI1 | 0.027678418 | 0.11903611 | 0.232521187 | 26.45246895 |
| Eg08_t046640 | EgCHS1 | MsSC66t327440 | MsCHS5 | 0.09887286 | 0.158904877 | 0.622214133 | 35.31219485 |
| Eg08_t040870 | EgCHS2 | Ms06t158650 | MsCHS2 | 0.021703399 | 0.085435496 | 0.254032576 | 18.98566568 |
| Eg08_t040870 | EgCHS2 | Ms10t280970 | MsCHS1 | 0.032099655 | 0.340620165 | 0.094238857 | 75.69336996 |
| Eg08_t020390 | EgCHS3 | Ms06t158650 | MsCHS2 | 0.045425972 | 0.32402875 | 0.140191179 | 72.00638892 |
| Eg08_t020390 | EgCHS3 | Ms10t280970 | MsCHS1 | 0.011301777 | 0.077703442 | 0.145447576 | 17.26743149 |
| Eg08_t049400 | EgCHS5 | MsSC51t327380 | MsCHS4 | 0.019392715 | 0.1121488 | 0.172919508 | 24.92195566 |

| | | | | | | | |
|---|---|---|---|---|---|---|---|
| Eg08_t049400 | EgCHS5 | Ms06t158650 | MsCHS2 | 0.108974557 | 0.367529742 | 0.296505411 | 81.673276 |
| Eg08_t049400 | EgCHS5 | Ms10t280970 | MsCHS1 | 0.107293231 | 0.384724902 | 0.278882991 | 85.49442257 |
| Eg09_t036940 | EgDFR1 | Ms04t107810 | MsDFR1 | 0.021692102 | 0.092034121 | 0.235696304 | 20.45202689 |
| Eg04_t027830 | EgDFR2 | Ms04t087800 | MsDFR2 | 0.021110665 | 0.065448478 | 0.32255395 | 14.54410629 |
| Eg09_t010710 | EgF3'5'H10 | Ms11t312500 | MsF3'5'H8 | 0.018194813 | 0.083203331 | 0.218678899 | 18.48962916 |
| Eg02_t007770 | EgF3'5'H2 | Ms08t214430 | MsF3'5'H1 | 0.023348189 | 0.096440853 | 0.242098532 | 21.43130064 |
| Eg06_t004020 | EgF3'5'H3 | Ms09t255750 | MsF3'5'H2 | 0.024679914 | 0.111286276 | 0.221769611 | 24.73028347 |
| Eg02_t010650 | EgF3'5'H5 | Ms08t211840 | MsF3'5'H3 | 0.022321415 | 0.066295721 | 0.33669466 | 14.73238253 |
| Eg02_t019070 | EgF3'5'H6 | Ms02t024400 | MsF3'5'H5 | 0.039101193 | 0.172440151 | 0.226752255 | 38.32003345 |
| Eg02_t019100 | EgF3'5'H7 | Ms02t024420 | MsF3'5'H7 | 0.160630791 | 0.486120138 | 0.330434348 | 108.0266973 |
| Eg02_t019110 | EgF3'5'H8 | Ms02t024370 | MsF3'5'H4 | 0.021300686 | 0.078363027 | 0.271820614 | 17.41400609 |
| Eg02_t019180 | EgF3'5'H9 | Ms02t024400 | MsF3'5'H5 | 0.089851232 | 0.362465952 | 0.24788875 | 80.54798922 |
| Eg03_t018110 | EgF3'H1 | Ms03t075990 | MsF3'H | 0.025866522 | 0.069406088 | 0.372683767 | 15.42357501 |
| Eg02_t022060 | EgF3H2 | Ms02t027200 | MsF3H | 0.011791537 | 0.102373304 | 0.115181757 | 22.74962304 |
| Eg01_t007540 | EgFLS2 | Ms03t052640 | MsFLS2 | 0.019866519 | 0.090779349 | 0.218844029 | 20.17318876 |
| Eg08_t007020 | EgFLS3 | Ms10t292840 | MsFLS3 | 0.065775128 | 0.235719582 | 0.279039727 | 52.38212934 |
| Eg03_t031640 | EgFLS4 | Ms08t229420 | MsFLS1 | 0.017185154 | 0.057763548 | 0.297508627 | 12.83634391 |
| Eg05_t019870 | EgLAR | Ms05t133790 | MsLAR | 0.035804646 | 0.113213008 | 0.316259118 | 25.15844618 |
| Eg05_t014690 | EgLDOX | Ms05t120140 | MsLDOX | 0.020733028 | 0.058525502 | 0.354256296 | 13.0056671 |
| Eg01_t040920 | EgPAL1 | Ms01t004140 | MsPAL1 | 0.012203659 | 0.075050065 | 0.16260691 | 16.67779215 |
| Eg05_t014880 | EgPAL2 | Ms05t120010 | MsPAL2 | 0.011247868 | 0.084219714 | 0.133553855 | 18.71549207 |
| Eg02_t018790 | EgPAL4 | Ms02t024050 | MsPAL4 | 0.022144164 | 0.088160077 | 0.251181321 | 19.59112811 |
| Eg02_t018790 | EgPAL4 | Ms08t220210 | MsPAL5 | 0.060504209 | 0.428112827 | 0.141327719 | 95.13618367 |
| Eg09_t011330 | EgPAL5 | Ms02t024050 | MsPAL4 | 0.069354685 | 0.373271387 | 0.185802307 | 82.94919717 |
| Eg09_t011330 | EgPAL5 | Ms08t220210 | MsPAL5 | 0.083072531 | 0.405778322 | 0.204723925 | 90.17296041 |

| | | | | | | | |
|---|---|---|---|---|---|---|---|
| Eg09_t011330 | EgPAL5 | Ms11t313180 | MsPAL3 | 0.021701333 | 0.079063813 | 0.274478705 | 17.56973631 |
| Eg02_t001010 | EgPAL6 | Ms02t024050 | MsPAL4 | 0.068309675 | 0.358209276 | 0.190697671 | 79.60206123 |
| Eg02_t001010 | EgPAL6 | Ms05t135820 | MsPAL7 | 0.067189621 | 0.401239167 | 0.167455288 | 89.16425941 |
| Eg02_t001010 | EgPAL6 | Ms08t220210 | MsPAL5 | 0.016964962 | 0.068769718 | 0.246692333 | 15.28215965 |
| Eg09_t017980 | EgPAL7 | Ms05t135820 | MsPAL7 | 0.04010364 | 0.385167969 | 0.104119873 | 85.59288196 |
| Eg09_t017980 | EgPAL7 | Ms11t319150 | MsPAL6 | 0.008760594 | 0.085594024 | 0.102350532 | 19.02089427 |
| Eg05_t024420 | EgPAL8 | Ms05t135820 | MsPAL7 | 0.011308523 | 0.075472919 | 0.149835502 | 16.77175982 |
| Eg05_t024420 | EgPAL8 | Ms11t319150 | MsPAL6 | 0.03417666 | 0.367210948 | 0.093070918 | 81.60243296 |
| Eg06_t005550 | EgPAL9 | Ms09t254450 | MsPAL8 | 0.008266173 | 0.06429884 | 0.128558666 | 14.28863105 |
| Eg03_t004200 | EgTT8.2 | Ms03t064490 | MsTT8.2 | 0.027791829 | 0.138726736 | 0.200335063 | 30.82816356 |
| Eg05_t009200 | EgTTG1 | Ms05t125970 | MsTTG1.2 | 0.013921513 | 0.091322181 | 0.152443944 | 20.29381791 |
| Eg09_t041330 | EgTTG1.2 | Ms04t112120 | MsTTG1 | 0.015821285 | 0.110290063 | 0.143451594 | 24.50890299 |
| Eg08_t037030 | EgUGT75C1.2 | Ms06t152710 | MsUGT75C1 | 0.058871564 | 0.120601552 | 0.488149304 | 26.80034496 |
| Eg08_t037040 | EgUGT75C1.3 | Ms06t152730 | MsUGT75C1.3 | 0.037713776 | 0.077793828 | 0.484791358 | 17.28751744 |
| Eg08_t008340 | EgUGT78D2 | Ms10t291590 | MsUGT78D2 | 0.049072609 | 0.123049247 | 0.398804625 | 27.34427709 |
| Eg06_t008220 | EgUGT78D2.2 | Ms07t193970 | MsUGT78D2.2 | 0.031696342 | 0.079762984 | 0.397381593 | 17.72510752 |
| Eg01_t005290 | EgUGT78D2.3 | Ms03t050760 | MsUGT78D2.3 | 0.035211831 | 0.10109291 | 0.348311579 | 22.46509108 |
| Eg07_t037900 | EgUGT79B1 | Ms09t246270 | MsUGT79B1 | 0.027679762 | 0.072992315 | 0.379214742 | 16.22051446 |

**Table S3d.** Divergence time, Ka, Ks and Ka/Ks values of orthologous flavonoid biosynthetic genes pairs between *M. acuminata* and *M. balbisiana*

| **Seq_1** | **Name1** | **Seq_2** | **Name2** | **Ka** | **Ks** | **Ka_Ks** | **Diversity_year** |
|---|---|---|---|---|---|---|---|
| Parent=Macma4_08_g13040 | Ma4CL1 | Mba08_g12720 | Mb4CL1.2 | 0.016183398 | 0.038593723 | 0.419327206 | 8.576382879 |
| Parent=Macma4_05_g22890 | Ma4CL1.2 | Mba05_g21540 | Mb4CL1 | 0.013224394 | 0.052863141 | 0.250162847 | 11.74736468 |
| Parent=Macma4_01_g19800 | Ma4CL10 | Mba01_g28370 | Mb4CL10 | 0.00572639 | 0.044466612 | 0.128779532 | 9.881469403 |

| | | | | | | | |
|---|---|---|---|---|---|---|---|
| Parent=Macma4_03_g13930 | Ma4CL11 | Mba01_g28370 | Mb4CL10 | 0.039447019 | 0.339818038 | 0.116082771 | 75.5151196 |
| Parent=Macma4_04_g06110 | Ma4CL12 | Mba04_g05720 | Mb4CL11 | 0.0107923 | 0.065299533 | 0.165273765 | 14.51100737 |
| Parent=Macma4_04_g06110 | Ma4CL12 | Mba04_g12340 | Mb4CL12 | 0.04307707 | 0.382206809 | 0.11270618 | 84.93484643 |
| Parent=Macma4_04_g13130 | Ma4CL13 | Mba04_g12340 | Mb4CL12 | 0.003277798 | 0.019967902 | 0.164153362 | 4.437311452 |
| Parent=Macma4_07_g05220 | Ma4CL14 | Mba07_g04890 | Mb4CL13 | 0.014573212 | 0.043576611 | 0.334427378 | 9.683691409 |
| Parent=Macma4_07_g18890 | Ma4CL15 | Mba07_g17040 | Mb4CL14 | 0.010623262 | 0.050142078 | 0.211863224 | 11.14268409 |
| Parent=Macma4_02_g21360 | Ma4CL2 | Mba02_g18560 | Mb4CL2 | 0.006261325 | 0.042061637 | 0.148860699 | 9.347030536 |
| Parent=Macma4_11_g22750 | Ma4CL2.2 | Mba11_g20940 | Mb4CL2.2 | 0.005497047 | 0.031160863 | 0.176408692 | 6.924636186 |
| Parent=Macma4_08_g13960 | Ma4CL4 | Mba08_g13620 | Mb4CL4 | 0.01354341 | 0.045519034 | 0.297532902 | 10.11534098 |
| Parent=Macma4_05_g28920 | Ma4CL5 | Mba05_g27160 | Mb4CL5 | 0.02014107 | 0.041838887 | 0.481395943 | 9.297530461 |
| Parent=Macma4_01_g07350 | Ma4CL7 | Mba01_g16650 | Mb4CL7 | 0.012625232 | 0.060334024 | 0.2092556 | 13.40756087 |
| Parent=Macma4_03_g14310 | Ma4CL8 | Mba03_g13490 | Mb4CL8 | 0.034549 | 0.068179503 | 0.506735875 | 15.15100068 |
| Parent=Macma4_06_g39580 | Ma4CL9 | Mba06_g36630 | Mb4CL9 | 0.013125848 | 0.033539552 | 0.391354293 | 7.453233823 |
| Parent=Macma4_08_g01440 | MaANR5 | Mba08_g01400 | MbANR | 0.012717558 | 0.043547851 | 0.292036405 | 9.677300159 |
| Parent=Macma4_10_g21630 | MaC1 | Mba10_g15520 | MbC1 | 0.009472564 | 0.063670558 | 0.14877463 | 14.14901298 |
| Parent=Macma4_09_g29590 | MaC1.2 | Mba09_g23680 | MbC1.3 | 0.17018449 | 0.251325074 | 0.677148872 | 55.85001634 |
| Parent=Macma4_10_g16840 | MaC4H1 | Mba06_g09560 | MbC4H3 | 0.03574232 | 0.32611299 | 0.109601031 | 72.46955335 |
| Parent=Macma4_07_g20380 | MaC4H2 | Mba07_g18380 | MbC4H4 | 0.002614572 | 0.039980914 | 0.065395493 | 8.884647598 |
| Parent=Macma4_06_g10450 | MaC4H3 | Mba06_g09560 | MbC4H3 | 0.00437096 | 0.045587256 | 0.095881185 | 10.13050133 |
| Parent=Macma4_09_g04800 | MaC4H4 | Mba09_g04380 | MbC4H1 | 0.009588973 | 0.037598234 | 0.255037851 | 8.355163005 |
| Parent=Macma4_04_g18950 | MaCHI1 | Mba04_g18040 | MbCHI2 | 0.043092791 | 0.059758558 | 0.721114983 | 13.27967948 |
| Parent=Macma4_11_g22930 | MaCHI2 | Mba11_g21110 | MbCHI1 | 0.006200518 | 0.043754812 | 0.141710531 | 9.723291481 |
| Parent=Macma4_06_g12560 | MaCHS1 | Mba06_g11590 | MbCHS1 | 0.005637535 | 0.046932602 | 0.120119799 | 10.42946719 |
| Parent=Macma4_06_g18170 | MaCHS2 | Mba06_g16830 | MbCHS2 | 0.009610081 | 0.037664818 | 0.255147408 | 8.369959497 |
| Parent=Macma4_06_g18170 | MaCHS2 | Mba06_g09120 | MbCHS7 | 0.125578746 | 0.320627725 | 0.391665275 | 71.25060559 |

| | | | | | | | |
|---|---|---|---|---|---|---|---|
| Parent=Macma4_06_g18170 | MaCHS2 | Mba10_g10820 | MbCHS8 | 0.036427721 | 0.325308328 | 0.111979061 | 72.29073954 |
| Parent=Macma4_06_g10010 | MaCHS5 | Mba06_g09120 | MbCHS7 | 0.040649522 | 0.131898935 | 0.308186884 | 29.31087435 |
| Parent=Macma4_10_g16140 | MaCHS6 | Mba06_g16830 | MbCHS2 | 0.031517242 | 0.332252271 | 0.094859374 | 73.8338379 |
| Parent=Macma4_10_g16140 | MaCHS6 | Mba10_g10820 | MbCHS8 | 0.002246982 | 0.03968464 | 0.056620958 | 8.818808806 |
| Parent=Macma4_04_g33220 | MaDFR1 | Mba04_g31800 | MbDFR1 | 0.011463473 | 0.065311185 | 0.17552083 | 14.51359659 |
| Parent=Macma4_03_g33910 | MaDFR2 | Mba01_g01310 | MbDFR2 | 0.009222373 | 0.026488322 | 0.348167493 | 5.886293723 |
| Parent=Macma4_08_g08290 | MaF3'5'H1 | Mba08_g07950 | MbF3'5'H1 | 0.006184691 | 0.055710056 | 0.111015707 | 12.3800124 |
| Parent=Macma4_02_g03260 | MaF3'5'H10 | Mba02_g01940 | MbF3'5'H11 | 0.0320733 | 0.159387344 | 0.201228648 | 35.41940979 |
| Parent=Macma4_02_g03290 | MaF3'5'H11 | Mba02_g01920 | MbF3'5'H10 | 0.034419448 | 0.153695475 | 0.223945745 | 34.15454994 |
| Parent=Macma4_02_g03300 | MaF3'5'H12 | Mba02_g01950 | MbF3'5'H12 | 0.043827731 | 0.076625327 | 0.57197447 | 17.02785045 |
| Parent=Macma4_11_g15260 | MaF3'5'H13 | Mba11_g13650 | MbF3'5'H13 | 0.007953532 | 0.043638756 | 0.182258451 | 9.697501319 |
| Parent=Macma4_11_g05110 | MaF3'5'H2 | Mba11_g04840 | MbF3'5'H2 | 0.029145115 | 0.048170646 | 0.605038917 | 10.70458792 |
| Parent=Macma4_09_g17660 | MaF3'5'H4 | Mba09_g16030 | MbF3'5'H3 | 0.006156136 | 0.030065515 | 0.204757367 | 6.681225561 |
| Parent=Macma4_10_g01260 | MaF3'5'H6 | Mba10_g00740 | MbF3'5'H4 | 0.011354504 | 0.040981362 | 0.27706506 | 9.106969332 |
| Parent=Macma4_02_g03040 | MaF3'5'H7 | Mba02_g01920 | MbF3'5'H10 | 0.074227897 | 0.117392689 | 0.632304255 | 26.08726418 |
| Parent=Macma4_03_g33170 | MaF3'H | Mba01_g02030 | MbF3'H | 0.010826003 | 0.042282227 | 0.256041465 | 9.396050469 |
| Parent=Macma4_07_g17110 | MaF3H1 | Mba07_g15320 | MbF3H2 | 0.009437797 | 0.042537172 | 0.221871745 | 9.452704907 |
| Parent=Macma4_02_g06130 | MaF3H2 | Mba02_g04670 | MbF3H1 | 0.007050581 | 0.031295283 | 0.225292121 | 6.954507371 |
| Parent=Macma4_03_g07220 | MaFLS1 | Mba03_g06940 | MbFLS4 | 0.016535061 | 0.041616514 | 0.397319707 | 9.24811427 |
| Parent=Macma4_03_g07220 | MaFLS1 | Mba09_g14060 | MbFLS5 | 0.153355877 | 0.498751411 | 0.307479584 | 110.833647 |
| Parent=Macma4_08_g24690 | MaFLS2 | Mba08_g23630 | MbFLS1 | 0.00724906 | 0.080377973 | 0.090187142 | 17.86177179 |
| Parent=Macma4_05_g19560 | MaLAR | Mba05_g18490 | MbLAR | 0.018251851 | 0.01936216 | 0.942655765 | 4.302702115 |
| Parent=Macma4_05_g03920 | MaLDOX | Mba05_g13780 | MbLDOX | 0.010887288 | 0.041561879 | 0.261953697 | 9.23597319 |
| Parent=Macma4_01_g04800 | MaPAL1 | Mba01_g14280 | MbPAL8 | 0.005293224 | 0.040140084 | 0.131868777 | 8.920018758 |
| Parent=Macma4_05_g03790 | MaPAL2 | Mba05_g13940 | MbPAL7 | 0.005601395 | 0.039936234 | 0.140258473 | 8.874718755 |

| | | | | | | | |
|---|---|---|---|---|---|---|---|
| Parent=Macma4_11_g15930 | MaPAL3 | Mba11_g14220 | MbPAL6 | 0.020865022 | 0.047925308 | 0.435365414 | 10.65006852 |
| Parent=Macma4_02_g02760 | MaPAL4 | Mba02_g01510 | MbPAL5 | 0.050074297 | 0.088817771 | 0.563786908 | 19.73728255 |
| Parent=Macma4_11_g22170 | MaPAL6 | Mba11_g20430 | MbPAL2 | 0.003110102 | 0.021128158 | 0.147201755 | 4.695146161 |
| Parent=Macma4_08_g14800 | MaPAL7 | Mba08_g14430 | MbPAL3 | 0.00560925 | 0.015651041 | 0.35839471 | 3.478009049 |
| Parent=Macma4_09_g16190 | MaPAL8 | Mba09_g14610 | MbPAL1 | 0.005302718 | 0.039786335 | 0.13327987 | 8.841407869 |
| Parent=Macma4_06_g25800 | MaTT8 | Mba06_g23950 | MbTT8 | 0.072115694 | 0.098035442 | 0.735608393 | 21.78565374 |
| Parent=Macma4_03_g19880 | MaTT8.2 | Mba03_g18730 | MbTT8.2 | 0.093409853 | 0.128965766 | 0.724299601 | 28.65905911 |
| Parent=Macma4_03_g19880 | MaTT8.2 | Mba06_g23950 | MbTT8 | 0.113307434 | 0.421935181 | 0.268542277 | 93.76337363 |
| Parent=Macma4_05_g10420 | MaTTG1 | Mba05_g07670 | MbTTG1 | 0.00942575 | 0.036603787 | 0.257507506 | 8.134174944 |
| Parent=Macma4_04_g37870 | MaTTG1.2 | Mba04_g36060 | MbTTG1.2 | 0.00623186 | 0.051268312 | 0.121553831 | 11.3929582 |
| Parent=Macma4_10_g31950 | MaUGT75C1 | Mba10_g24560 | MbUGT75C1 | 0.0159793 | 0.014068408 | 1.135828569 | 3.126312845 |
| Parent=Macma4_06_g07210 | MaUGT75C1.2 | Mba06_g06490 | MbUGT75C1.3 | 0.011271346 | 0.03245344 | 0.347308185 | 7.211875657 |
| Parent=Macma4_06_g07200 | MaUGT75C1.3 | Mba06_g06480 | MbUGT75C1.2 | 0.019152014 | 0.053382209 | 0.358771482 | 11.86271308 |
| Parent=Macma4_10_g27580 | MaUGT78D2 | Mba10_g20800 | MbUGT78D2.2 | 0.030300388 | 0.059068052 | 0.51297422 | 13.12623384 |
| Parent=Macma4_07_g16200 | MaUGT78D2.2 | Mba07_g14580 | MbUGT78D2.3 | 0.010876921 | 0.040472246 | 0.26875012 | 8.993832482 |
| Parent=Macma4_03_g04880 | MaUGT78D2.3 | Mba03_g04830 | MbUGT78D2 | 0.014911028 | 0.051596271 | 0.288994294 | 11.465838 |

**Table S3e.** Divergence time, Ka, Ks and Ka/Ks values of orthologous flavonoid biosynthetic genes pairs between *M. acuminata* and *M. schizocharpa*

| **Seq_1** | **Name1** | **Seq_2** | **Name2** | **Ka** | **Ks** | **Ka_Ks** | **Diversity_year** |
|---|---|---|---|---|---|---|---|
| Parent=Macma4_08_g13040 | Ma4CL1 | Ms08t218760 | Ms4CL1 | 0.007492778 | 0.019208733 | 0.390071419 | 4.268607336 |

| | | | | | | | |
|---|---|---|---|---|---|---|---|
| Parent=Macma4_05_g22890 | Ma4CL1.2 | Ms05t136800 | Ms4CL1.2 | 0.004929836 | 0.022228423 | 0.221780753 | 4.939649472 |
| Parent=Macma4_01_g19800 | Ma4CL10 | Ms01t017810 | Ms4CL10 | 0.001631988 | 0.024362678 | 0.0669872 | 5.413928436 |
| Parent=Macma4_01_g19800 | Ma4CL10 | Ms03t058920 | Ms4CL11 | 0.038559243 | 0.333225729 | 0.115715083 | 74.05016192 |
| Parent=Macma4_01_g19800 | Ma4CL10 | Ms07t183930 | Ms4CL14 | 0.075310931 | 0.449422029 | 0.167572852 | 99.87156208 |
| Parent=Macma4_03_g13930 | Ma4CL11 | Ms01t017810 | Ms4CL10 | 0.039455271 | 0.347144062 | 0.113656764 | 77.14312494 |
| Parent=Macma4_03_g13930 | Ma4CL11 | Ms03t058920 | Ms4CL11 | 0.002445654 | 0.019513296 | 0.125332715 | 4.336287955 |
| Parent=Macma4_03_g13930 | Ma4CL11 | Ms07t183930 | Ms4CL14 | 0.071971516 | 0.480161248 | 0.149890305 | 106.7024995 |
| Parent=Macma4_04_g06110 | Ma4CL12 | Ms04t083350 | Ms4CL12 | 0.00578914 | 0.031985167 | 0.180994514 | 7.107814821 |
| Parent=Macma4_04_g06110 | Ma4CL12 | Ms04t089570 | Ms4CL13 | 0.044830481 | 0.37375165 | 0.119947245 | 83.05592232 |
| Parent=Macma4_04_g13130 | Ma4CL13 | Ms04t089570 | Ms4CL13 | 0.001637555 | 0.004938289 | 0.331603737 | 1.097397655 |
| Parent=Macma4_07_g05220 | Ma4CL14 | Ms01t017810 | Ms4CL10 | 0.076788935 | 0.427247549 | 0.179729374 | 94.94389985 |
| Parent=Macma4_07_g05220 | Ma4CL14 | Ms03t058920 | Ms4CL11 | 0.072459473 | 0.490835425 | 0.147624782 | 109.0745388 |
| Parent=Macma4_07_g05220 | Ma4CL14 | Ms07t183930 | Ms4CL14 | 0.007415096 | 0.027371895 | 0.270901827 | 6.082643403 |
| Parent=Macma4_07_g18890 | Ma4CL15 | Ms07t196560 | Ms4CL15 | 0.009149244 | 0.024471992 | 0.373865947 | 5.438220343 |
| Parent=Macma4_09_g24760 | Ma4CL16 | Ms09t262230 | Ms4CL16 | 0.004923095 | 0.022487631 | 0.218924556 | 4.9972513 |
| Parent=Macma4_11_g22750 | Ma4CL2.2 | Ms11t319670 | Ms4CL2.2 | 7.83E-04 | 0.01184391 | 0.066074021 | 2.63197995 |
| Parent=Macma4_11_g14090 | Ma4CL3 | Ms11t311430 | Ms4CL3 | 0.043415112 | 0.048132888 | 0.901984346 | 8.321008483 |
| Parent=Macma4_08_g13960 | Ma4CL4 | Ms08t219500 | Ms4CL4 | 8.34E-04 | 0.020370924 | 0.040959182 | 4.526872019 |
| Parent=Macma4_05_g28920 | Ma4CL5 | Ms05t142250 | Ms4CL5 | 0.010782602 | 0.020517997 | 0.525519237 | 4.559554869 |
| Parent=Macma4_01_g07350 | Ma4CL7 | Ms01t006680 | Ms4CL7 | 0.008184504 | 0.025633563 | 0.319288565 | 5.696347327 |
| Parent=Macma4_03_g14310 | Ma4CL8 | Ms03t059300 | Ms4CL8 | 0.010938719 | 0.015273682 | 0.716180888 | 2.34241607 |
| Parent=Macma4_06_g39580 | Ma4CL9 | Ms06t178240 | Ms4CL9 | 0.004899643 | 0.025594294 | 0.191434978 | 5.68762093 |
| Parent=Macma4_07_g10660 | MaANR2 | Ms07t188910 | MsANR3 | 0.028232302 | 0.058915661 | 0.479198586 | 13.09236919 |
| Parent=Macma4_08_g01440 | MaANR5 | Ms08t207990 | MsANR1 | 0.006338512 | 0.01941856 | 0.326415147 | 4.315235655 |
| Parent=Macma4_09_g29590 | MaC1.2 | Ms09t266370 | MsC1 | 0.010136716 | 0.010944107 | 0.926226019 | 2.432023698 |

| | | | | | | | |
|---|---|---|---|---|---|---|---|
| Parent=Macma4_10_g16840 | MaC4H1 | Ms10t281630 | MsC4H1 | 0.007019648 | 0.016667353 | 0.421161524 | 3.703856131 |
| Parent=Macma4_07_g20380 | MaC4H2 | Ms07t197940 | MsC4H2 | 0.003488885 | 0.037026978 | 0.0942255 | 8.228217264 |
| Parent=Macma4_06_g10450 | MaC4H3 | Ms10t281630 | MsC4H1 | 0.034363116 | 0.315819854 | 0.108806066 | 70.18218989 |
| Parent=Macma4_09_g04800 | MaC4H4 | Ms09t244050 | MsC4H3 | 0.001734606 | 0.025815168 | 0.067193293 | 5.736703981 |
| Parent=Macma4_11_g22930 | MaCHI2 | Ms11t319860 | MsCHI1 | 0.010952847 | 0.026182746 | 0.41832309 | 5.818388019 |
| Parent=Macma4_06_g12560 | MaCHS1 | MsSC66t327440 | MsCHS5 | 0.095019137 | 0.155167236 | 0.612365982 | 34.48160796 |
| Parent=Macma4_06_g18170 | MaCHS2 | Ms06t158650 | MsCHS2 | 0.0142117 | 0.041464448 | 0.342744208 | 9.214321842 |
| Parent=Macma4_06_g18170 | MaCHS2 | Ms10t280970 | MsCHS1 | 0.038777749 | 0.331098623 | 0.117118424 | 73.57747178 |
| Parent=Macma4_06_g10010 | MaCHS5 | MsSC51t327380 | MsCHS4 | 0.038761163 | 0.150595466 | 0.257385987 | 33.46565907 |
| Parent=Macma4_06_g10010 | MaCHS5 | Ms06t158650 | MsCHS2 | 0.117422023 | 0.388692181 | 0.302095151 | 86.37604011 |
| Parent=Macma4_10_g16140 | MaCHS6 | Ms06t158650 | MsCHS2 | 0.043030158 | 0.351872673 | 0.122289002 | 78.19392732 |
| Parent=Macma4_10_g16140 | MaCHS6 | Ms10t280970 | MsCHS1 | 0.002246351 | 0.014201608 | 0.158175849 | 3.155912834 |
| Parent=Macma4_04_g33220 | MaDFR1 | Ms04t107810 | MsDFR1 | 0.00490298 | 0.043333351 | 0.11314564 | 9.629633444 |
| Parent=Macma4_03_g33910 | MaDFR2 | Ms03t076720 | MsDFR3 | 0.009852358 | 0.020216857 | 0.487333822 | 4.492634998 |
| Parent=Macma4_04_g11180 | MaDFR3 | Ms04t087800 | MsDFR2 | 0.003678732 | 0.004204636 | 0.874922823 | 0.934363578 |
| Parent=Macma4_08_g08290 | MaF3'5'H1 | Ms08t214430 | MsF3'5'H1 | 0.007025225 | 0.030844748 | 0.227760816 | 6.854388376 |
| Parent=Macma4_02_g03290 | MaF3'5'H11 | Ms02t024400 | MsF3'5'H5 | 0.007037659 | 0.024530917 | 0.286889349 | 5.451314916 |
| Parent=Macma4_02_g03300 | MaF3'5'H12 | Ms02t024420 | MsF3'5'H7 | 0.189036887 | 0.221518094 | 0.853369957 | 49.2262432 |
| Parent=Macma4_11_g15260 | MaF3'5'H13 | Ms11t312500 | MsF3'5'H8 | 0.004407241 | 0.029747027 | 0.148157347 | 6.610450352 |
| Parent=Macma4_08_g05480 | MaF3'5'H3 | Ms08t211840 | MsF3'5'H3 | 0.009722461 | 0.016118821 | 0.603174447 | 3.58196012 |
| Parent=Macma4_09_g17660 | MaF3'5'H4 | Ms09t255750 | MsF3'5'H2 | 0.003748544 | 0.020315631 | 0.184515236 | 4.514584768 |
| Parent=Macma4_02_g03040 | MaF3'5'H7 | Ms02t024400 | MsF3'5'H5 | 0.093617041 | 0.195028562 | 0.48001708 | 43.33968054 |
| Parent=Macma4_03_g33170 | MaF3'H | Ms03t075990 | MsF3'H | 0.006935964 | 0.027927915 | 0.248352395 | 6.206203247 |
| Parent=Macma4_02_g06130 | MaF3H2 | Ms02t027200 | MsF3H | 0.00351838 | 0.027275733 | 0.128993056 | 6.061274028 |
| Parent=Macma4_03_g07220 | MaFLS1 | Ms03t052640 | MsFLS2 | 0.003931857 | 0.017291832 | 0.227382323 | 3.842629383 |

Musaceae flavonoid biosynthetic genes

| | | | | | | | |
|---|---|---|---|---|---|---|---|
| Parent=Macma4_08_g24690 | MaFLS2 | Ms08t229420 | MsFLS1 | 0.00524134 | 0.030638834 | 0.171068536 | 6.80862969 |
| Parent=Macma4_05_g19560 | MaLAR | Ms05t133790 | MsLAR | 0.017349837 | 0.035796267 | 0.484682847 | 7.9547261 |
| Parent=Macma4_05_g03920 | MaLDOX | Ms05t120140 | MsLDOX | 0 | 0.016360567 | 0 | 3.635681542 |
| Parent=Macma4_01_g04800 | MaPAL1 | Ms01t004140 | MsPAL1 | 6.20E-04 | 0.011549795 | 0.053716241 | 2.566621142 |
| Parent=Macma4_05_g03790 | MaPAL2 | Ms05t120010 | MsPAL2 | 6.20E-04 | 0.017717359 | 0.035006299 | 3.937190961 |
| Parent=Macma4_11_g15930 | MaPAL3 | Ms11t313180 | MsPAL3 | 0.002071348 | 0.01476841 | 0.140255298 | 3.281868815 |
| Parent=Macma4_02_g02760 | MaPAL4 | Ms02t024050 | MsPAL4 | 0.007531444 | 0.022917123 | 0.32863828 | 5.092694108 |
| Parent=Macma4_02_g02760 | MaPAL4 | Ms08t220210 | MsPAL5 | 0.06130516 | 0.38841449 | 0.157834378 | 86.31433114 |
| Parent=Macma4_05_g21700 | MaPAL5 | Ms05t135820 | MsPAL7 | 0.003742212 | 0.019268883 | 0.194210095 | 4.281973904 |
| Parent=Macma4_11_g22170 | MaPAL6 | Ms11t319150 | MsPAL6 | 0.00248808 | 0.01915813 | 0.129870727 | 4.257362175 |
| Parent=Macma4_08_g14800 | MaPAL7 | Ms02t024050 | MsPAL4 | 0.061075347 | 0.342676126 | 0.178230528 | 76.15025032 |
| Parent=Macma4_08_g14800 | MaPAL7 | Ms08t220210 | MsPAL5 | 0.006858972 | 0.019637843 | 0.349273205 | 4.363965011 |
| Parent=Macma4_09_g16190 | MaPAL8 | Ms09t254450 | MsPAL8 | 0.001175261 | 0.028715935 | 0.040927137 | 6.381318806 |
| Parent=Macma4_03_g19880 | MaTT8.2 | Ms03t064490 | MsTT8.2 | 0.01045853 | 0.024624242 | 0.42472494 | 5.472053777 |
| Parent=Macma4_05_g10420 | MaTTG1 | Ms05t125970 | MsTTG1.2 | 0.001252871 | 0.015180784 | 0.082530089 | 3.37350754 |
| Parent=Macma4_04_g37870 | MaTTG1.2 | Ms04t112120 | MsTTG1 | 0 | 0.019987858 | 0 | 4.441746332 |
| Parent=Macma4_10_g31950 | MaUGT75C1 | Ms10t295740 | MsUGT75C1.4 | 0.006647717 | 0.011439688 | 0.58110998 | 2.542152876 |
| Parent=Macma4_06_g07210 | MaUGT75C1.2 | Ms06t152730 | MsUGT75C1.3 | 0.003741531 | 0.017480762 | 0.214037056 | 3.88461382 |
| Parent=Macma4_06_g07200 | MaUGT75C1.3 | Ms06t152710 | MsUGT75C1 | 0.030358276 | 0.058194728 | 0.521667114 | 12.93216181 |
| Parent=Macma4_10_g27580 | MaUGT78D2 | Ms10t291590 | MsUGT78D2 | 0.015868361 | 0.035197074 | 0.450843171 | 7.821572037 |
| Parent=Macma4_07_g16200 | MaUGT78D2.2 | Ms07t193970 | MsUGT78D2.2 | 0.004971845 | 0.034788845 | 0.142914907 | 7.73085443 |
| Parent=Macma4_03_g04880 | MaUGT78D2.3 | Ms03t050760 | MsUGT78D2.3 | 0.008897781 | 0.014038781 | 0.6338001 | 3.119729209 |
| Parent=Macma4_09_g07100 | MaUGT79B1 | Ms09t246270 | MsUGT79B1 | 0.011530374 | 0.011442415 | 1.007687066 | 2.542758895 |

Musaceae flavonoid biosynthetic genes

**Table S3f.** Divergence time, Ka, Ks and Ka/Ks values of orthologous flavonoid biosynthetic genes pairs between *M. acuminata* and *M. schizocharpa*

| **Seq_1** | **Name1** | **Seq_2** | **Name2** | **Ka** | **Ks** | **Ka_Ks** | **Diversity_year** |
|---|---|---|---|---|---|---|---|
| Mba05_g21540 | Mb4CL1 | Ms05t136800 | Ms4CL1.2 | 0.011565698 | 0.055371049 | 0.208876268 | 12.30467747 |
| Mba08_g12720 | Mb4CL1.2 | Ms08t218760 | Ms4CL1 | 0.014261244 | 0.041427926 | 0.344242293 | 9.206205704 |
| Mba01_g28370 | Mb4CL10 | Ms01t017810 | Ms4CL10 | 0.005729514 | 0.054625997 | 0.104886218 | 12.13911052 |
| Mba01_g28370 | Mb4CL10 | Ms03t058920 | Ms4CL11 | 0.038572669 | 0.344127865 | 0.112088189 | 76.47285884 |
| Mba01_g28370 | Mb4CL10 | Ms07t183930 | Ms4CL14 | 0.077155151 | 0.4443799 | 0.173624304 | 98.75108895 |
| Mba04_g05720 | Mb4CL11 | Ms04t083350 | Ms4CL12 | 0.014136653 | 0.068042434 | 0.207762306 | 15.12054092 |
| Mba04_g05720 | Mb4CL11 | Ms04t089570 | Ms4CL13 | 0.048321794 | 0.361929109 | 0.133511763 | 80.42869088 |
| Mba04_g12340 | Mb4CL12 | Ms04t089570 | Ms4CL13 | 0.004922085 | 0.025044063 | 0.196536997 | 5.565347343 |
| Mba04_g12340 | Mb4CL12 | Ms04t083350 | Ms4CL12 | 0.046100056 | 0.393246213 | 0.117229498 | 87.38804732 |
| Mba07_g04890 | Mb4CL13 | Ms07t183930 | Ms4CL14 | 0.014559281 | 0.049176158 | 0.296063811 | 10.9280351 |
| Mba07_g17040 | Mb4CL14 | Ms07t196560 | Ms4CL15 | 0.016540987 | 0.05761229 | 0.287108663 | 12.80273102 |
| Mba11_g20940 | Mb4CL2.2 | Ms11t319670 | Ms4CL2.2 | 0.004709284 | 0.033612346 | 0.140105768 | 7.46941025 |
| Mba11_g12600 | Mb4CL3.2 | Ms11t311430 | Ms4CL3 | 0.117634338 | 0.102305839 | 1.149830142 | 22.73463092 |
| Mba08_g13620 | Mb4CL4 | Ms08t219500 | Ms4CL4 | 0.010561772 | 0.049382672 | 0.213876068 | 10.97392719 |
| Mba05_g27160 | Mb4CL5 | Ms05t142250 | Ms4CL5 | 0.021847225 | 0.039436993 | 0.553977951 | 8.763776221 |
| Mba01_g16650 | Mb4CL7 | Ms01t006680 | Ms4CL7 | 0.009448944 | 0.049935244 | 0.189223946 | 11.09672085 |
| Mba03_g13490 | Mb4CL8 | Ms03t059300 | Ms4CL8 | 0.042106403 | 0.067845131 | 0.620625272 | 15.07669578 |
| Mba06_g36630 | Mb4CL9 | Ms06t178240 | Ms4CL9 | 0.014781838 | 0.044178173 | 0.334595944 | 9.817371804 |
| Mba08_g01400 | MbANR | Ms08t207990 | MsANR1 | 0.016571739 | 0.039508805 | 0.419444189 | 8.779734391 |
| Mba06_g05630 | MbC1.2 | Ms06t151760 | MsC1.2 | 0.017586737 | 0.042830548 | 0.410612002 | 9.517899556 |
| Mba09_g23680 | MbC1.3 | Ms09t266370 | MsC1 | 0.182934985 | 0.250840554 | 0.729287914 | 55.74234541 |

| | | | | | | | |
|---|---|---|---|---|---|---|---|
| Mba09_g04380 | MbC4H1 | Ms09t244050 | MsC4H3 | 0.011347321 | 0.034621531 | 0.32775331 | 7.693673627 |
| Mba06_g09560 | MbC4H3 | Ms10t281630 | MsC4H1 | 0.033477738 | 0.331756574 | 0.10091055 | 73.72368313 |
| Mba07_g18380 | MbC4H4 | Ms07t197940 | MsC4H2 | 0.002614572 | 0.051806732 | 0.050467796 | 11.51260711 |
| Mba11_g21110 | MbCHI1 | Ms11t319860 | MsCHI1 | 0.006194116 | 0.043914982 | 0.141047911 | 9.758884792 |
| Mba04_g18040 | MbCHI2 | Ms04t093250 | MsCHI2 | 0.036279394 | 0.066520847 | 0.545383826 | 14.7824104 |
| Mba06_g11590 | MbCHS1 | MsSC66t327440 | MsCHS5 | 0.073701728 | 0.141407869 | 0.521199624 | 31.42397099 |
| Mba06_g16830 | MbCHS2 | Ms06t158650 | MsCHS2 | 0.013635448 | 0.062173869 | 0.219311555 | 13.81641543 |
| Mba06_g16830 | MbCHS2 | Ms10t280970 | MsCHS1 | 0.031517242 | 0.343261643 | 0.091816965 | 76.28036502 |
| Mba06_g09090 | MbCHS6 | MsSC51t327370 | MsCHS3 | 0.061634764 | 0.127288788 | 0.484212043 | 28.28639744 |
| Mba06_g09120 | MbCHS7 | MsSC51t327380 | MsCHS4 | 0.006520363 | 0.044819851 | 0.145479349 | 9.959966903 |
| Mba10_g10820 | MbCHS8 | Ms06t158650 | MsCHS2 | 0.040662452 | 0.35711559 | 0.11386356 | 79.35901995 |
| Mba10_g10820 | MbCHS8 | Ms10t280970 | MsCHS1 | 0.002246982 | 0.047131179 | 0.047675071 | 10.47359539 |
| Mba04_g31800 | MbDFR1 | Ms04t107810 | MsDFR1 | 0.0128093 | 0.045541144 | 0.281268737 | 10.12025429 |
| Mba01_g01310 | MbDFR2 | Ms03t076720 | MsDFR3 | 0.011696143 | 0.022382129 | 0.522566174 | 4.973806428 |
| Mba08_g07950 | MbF3'5'H1 | Ms08t214430 | MsF3'5'H1 | 0.00803019 | 0.050057667 | 0.160418774 | 11.1239261 |
| Mba02_g01950 | MbF3'5'H12 | Ms02t024400 | MsF3'5'H5 | 0.088357805 | 0.378038601 | 0.233726938 | 84.00857802 |
| Mba11_g13650 | MbF3'5'H13 | Ms11t312500 | MsF3'5'H8 | 0.005290638 | 0.049344122 | 0.107219224 | 10.9653604 |
| Mba09_g16030 | MbF3'5'H3 | Ms09t255750 | MsF3'5'H2 | 0.003755289 | 0.023134364 | 0.162325131 | 5.140969831 |
| Mba01_g02030 | MbF3'H | Ms03t075990 | MsF3'H | 0.012178593 | 0.03313344 | 0.367561983 | 7.362986769 |
| Mba02_g04670 | MbF3H1 | Ms02t027200 | MsF3H | 0.005872021 | 0.043347585 | 0.135463617 | 9.632796608 |
| Mba08_g23630 | MbFLS1 | Ms08t229420 | MsFLS1 | 0.001442135 | 0.075472086 | 0.019108188 | 16.77157461 |
| Mba10_g21980 | MbFLS3 | Ms10t292840 | MsFLS3 | 0.065206786 | 0.254366038 | 0.256350207 | 56.52578623 |
| Mba03_g06940 | MbFLS4 | Ms03t052640 | MsFLS2 | 0.019214832 | 0.041616514 | 0.461711708 | 9.24811427 |
| Mba09_g14060 | MbFLS5 | Ms03t052640 | MsFLS2 | 0.154976915 | 0.498751411 | 0.310729777 | 110.833647 |
| Mba05_g18490 | MbLAR | Ms05t133790 | MsLAR | 0.029105746 | 0.04528864 | 0.642672124 | 10.06414214 |

| | | | | | | | |
|---|---|---|---|---|---|---|---|
| Mba05_g13780 | MbLDOX | Ms05t120140 | MsLDOX | 0.010889483 | 0.033040442 | 0.329580439 | 7.342320523 |
| Mba09_g14610 | MbPAL1 | Ms09t254450 | MsPAL8 | 0.005303499 | 0.047231099 | 0.112288278 | 10.49579986 |
| Mba11_g20430 | MbPAL2 | Ms11t319150 | MsPAL6 | 0.00435912 | 0.025010333 | 0.17429275 | 5.557851737 |
| Mba08_g14430 | MbPAL3 | Ms02t024050 | MsPAL4 | 0.061421772 | 0.341024262 | 0.180109682 | 75.7831693 |
| Mba08_g14430 | MbPAL3 | Ms08t220210 | MsPAL5 | 0.006546516 | 0.024620625 | 0.265895616 | 5.47125008 |
| Mba02_g01510 | MbPAL5 | Ms02t024050 | MsPAL4 | 0.04281569 | 0.063804104 | 0.671049157 | 14.17868986 |
| Mba11_g14220 | MbPAL6 | Ms02t024050 | MsPAL4 | 0.072903193 | 0.425306377 | 0.171413356 | 94.51252818 |
| Mba11_g14220 | MbPAL6 | Ms11t313180 | MsPAL3 | 0.013682027 | 0.043727485 | 0.312893056 | 9.717218984 |
| Mba05_g13940 | MbPAL7 | Ms05t120010 | MsPAL2 | 0.006224425 | 0.035881282 | 0.17347276 | 7.973618164 |
| Mba01_g14280 | MbPAL8 | Ms01t004140 | MsPAL1 | 0.004667581 | 0.036151576 | 0.129111401 | 8.033683636 |
| Mba06_g23950 | MbTT8 | Ms03t064490 | MsTT8.2 | 0.11189735 | 0.429698066 | 0.260409248 | 95.48845915 |
| Mba03_g18730 | MbTT8.2 | Ms03t064490 | MsTT8.2 | 0.096555655 | 0.128765197 | 0.749858325 | 28.6144883 |
| Mba05_g07670 | MbTTG1 | Ms05t125970 | MsTTG1.2 | 0.007551191 | 0.034566117 | 0.218456449 | 7.681359387 |
| Mba04_g36060 | MbTTG1.2 | Ms04t112120 | MsTTG1 | 0.00655026 | 0.040550416 | 0.161533731 | 9.011203545 |
| Mba10_g24560 | MbUGT75C1 | Ms10t295740 | MsUGT75C1.4 | 0.015261635 | 0.014262374 | 1.070062701 | 3.169416482 |
| Mba06_g06480 | MbUGT75C1.2 | Ms06t152710 | MsUGT75C1 | 0.024956313 | 0.056729344 | 0.439918947 | 12.60652098 |
| Mba06_g06490 | MbUGT75C1.3 | Ms06t152730 | MsUGT75C1.3 | 0.00937732 | 0.026479828 | 0.354130696 | 5.884406236 |
| Mba03_g04830 | MbUGT78D2 | Ms03t050760 | MsUGT78D2.3 | 0.015889373 | 0.048867856 | 0.325149783 | 10.85952361 |
| Mba10_g20800 | MbUGT78D2.2 | Ms10t291590 | MsUGT78D2 | 0.029768135 | 0.057634703 | 0.516496715 | 12.80771168 |
| Mba07_g14580 | MbUGT78D2.3 | Ms07t193970 | MsUGT78D2.2 | 0.016013953 | 0.03487311 | 0.459206333 | 7.749579944 |

**Table S3g.** Divergence time, Ka, Ks and Ka/Ks values of orthologous flavonoid biosynthetic genes pairs between *M. beccarii* and *E. glaucum*

| **Seq_1** | **Name1** | **Seq_2** | **Name2** | **Ka** | **Ks** | **Ka_Ks** | **Diversity_year** |
|---|---|---|---|---|---|---|---|
| Mbe06_t016280 | Mbe4CL1 | Eg04_t006840 | Eg4CL9 | 0.028156771 | 0.127216837 | 0.221328962 | 28.27040831 |
| Mbe07_t013640 | Mbe4CL10 | Eg02_t002700 | Eg4CL1 | 0.03826276 | 0.123142796 | 0.310718622 | 27.36506571 |

| | | | | | | | |
|---|---|---|---|---|---|---|---|
| Mbe02_t027270 | Mbe4CL11 | Eg02_t037340 | Eg4CL2 | 0.032789868 | 0.1019178 | 0.321728569 | 22.64839991 |
| Mbe06_t040140 | Mbe4CL12 | Eg09_t018540 | Eg4CL2.2 | 0.014219371 | 0.108387028 | 0.131190711 | 24.08600627 |
| Mbe03_t056570 | Mbe4CL13 | Eg04_t025780 | Eg4CL12 | 0.050913726 | 0.424796606 | 0.119854361 | 94.39924582 |
| Mbe03_t056570 | Mbe4CL13 | Eg07_t014740 | Eg4CL15 | 0.015330967 | 0.092024666 | 0.166596278 | 20.44992568 |
| Mbe01_t005810 | Mbe4CL14 | Eg01_t025930 | Eg4CL11 | 0.015210168 | 0.101697954 | 0.149562181 | 22.59954532 |
| Mbe01_t005810 | Mbe4CL14 | Eg01_t014520 | Eg4CL10 | 0.052253646 | 0.376333647 | 0.138849253 | 83.62969941 |
| Mbe03_t000920 | Mbe4CL15 | Eg01_t014520 | Eg4CL10 | 0.021055463 | 0.082535626 | 0.255107569 | 18.34125019 |
| Mbe03_t000920 | Mbe4CL15 | Eg01_t025930 | Eg4CL11 | 0.047028135 | 0.356907932 | 0.131765453 | 79.31287385 |
| Mbe07_t022330 | Mbe4CL16 | Eg01_t025930 | Eg4CL11 | 0.075289271 | 0.445490417 | 0.169003122 | 98.99787054 |
| Mbe04_t006230 | Mbe4CL17 | Eg04_t032900 | Eg4CL13 | 0.021725513 | 0.073383394 | 0.296054893 | 16.30742093 |
| Mbe04_t006230 | Mbe4CL17 | Eg04_t025780 | Eg4CL12 | 0.04427457 | 0.348093526 | 0.127191594 | 77.35411695 |
| Mbe04_t013920 | Mbe4CL18 | Eg04_t025780 | Eg4CL12 | 0.014054838 | 0.105043564 | 0.133800084 | 23.34301431 |
| Mbe04_t013920 | Mbe4CL18 | Eg04_t032900 | Eg4CL13 | 0.047016097 | 0.36371296 | 0.129267037 | 80.82510232 |
| Mbe03_t000510 | Mbe4CL2 | Eg03_t041750 | Eg4CL8 | 0.04547516 | 0.094217396 | 0.482662032 | 20.93719918 |
| Mbe07_t014630 | Mbe4CL3 | Eg02_t001820 | Eg4CL4 | 0.016003976 | 0.100368254 | 0.159452573 | 22.30405655 |
| Mbe05_t019740 | Mbe4CL4 | Eg07_t001640 | Eg4CL14 | 0.030728815 | 0.108062268 | 0.284362117 | 24.01383736 |
| Mbe01_t019320 | Mbe4CL6 | Eg01_t038400 | Eg4CL7 | 0.031831154 | 0.107281695 | 0.29670629 | 23.8403766 |
| Mbe05_t004680 | Mbe4CL7 | Eg05_t033420 | Eg4CL5 | 0.02315161 | 0.10765073 | 0.215062268 | 23.9223845 |
| Mbe03_t020730 | Mbe4CL8 | Eg09_t002410 | Eg4CL3 | 0.044630582 | 0.073087895 | 0.610642596 | 16.24175436 |
| Mbe05_t011100 | Mbe4CL9 | Eg05_t023200 | Eg4CL1.2 | 0.031036019 | 0.084705308 | 0.366399927 | 18.82340186 |
| Mbe07_t028460 | MbeANR | Eg06_t013840 | EgANR1 | 0.045245013 | 0.124314848 | 0.363955022 | 27.62552174 |
| Mbe01_t033880 | MbeC1 | Eg08_t035970 | EgC1.2 | 0.038287124 | 0.14582585 | 0.262553749 | 32.40574448 |
| Mbe03_t051550 | MbeC1 | Eg07_t019620 | EgC1 | 0.049504291 | 0.119846005 | 0.413065848 | 26.63244545 |
| Mbe01_t038520 | MbeC4H1 | Eg08_t048930 | EgC4H3 | 0.009658434 | 0.096980096 | 0.099591925 | 21.55113255 |
| Mbe10_t000860 | MbeC4H2 | Eg07_t003130 | EgC4H2 | 0.00787696 | 0.075847915 | 0.103852025 | 16.85509226 |

Musaceae flavonoid biosynthetic genes

| | | | | | | | |
|---|---|---|---|---|---|---|---|
| Mbe02_t033650 | MbeC4H3 | Eg08_t019660 | EgC4H1 | 0.010536502 | 0.08469968 | 0.124398365 | 18.82215106 |
| Mbe02_t033650 | MbeC4H3 | Eg08_t048930 | EgC4H3 | 0.03866624 | 0.363923178 | 0.106248358 | 80.87181734 |
| Mbe08_t025090 | MbeC4H4 | Eg07_t034110 | EgC4H4 | 0.017547225 | 0.089435856 | 0.196198997 | 19.87463469 |
| Mbe04_t020690 | MbeCHI1 | Eg01_t015550 | EgCHI1 | 0.055726329 | 0.080869124 | 0.689092778 | 17.97091648 |
| Mbe06_t039860 | MbeCHI2 | Eg09_t018750 | EgCHI2 | 0.031561459 | 0.114143862 | 0.276505969 | 25.36530256 |
| Mbe01_t038030 | MbeCHS2 | Eg08_t049400 | EgCHS5 | 0.023030421 | 0.111966071 | 0.205691069 | 24.88134906 |
| Mbe01_t038030 | MbeCHS2 | Eg08_t040870 | EgCHS2 | 0.104952554 | 0.327177575 | 0.320781626 | 72.7061278 |
| Mbe01_t038030 | MbeCHS2 | Eg08_t020390 | EgCHS3 | 0.115305009 | 0.384043822 | 0.300239197 | 85.34307159 |
| Mbe01_t041060 | MbeCHS3 | Eg08_t046640 | EgCHS1 | 0.026331045 | 0.065476271 | 0.402146379 | 14.55028248 |
| Mbe01_t047580 | MbeCHS4 | Eg08_t040870 | EgCHS2 | 0.014734135 | 0.062154915 | 0.237055018 | 13.81220342 |
| Mbe01_t047580 | MbeCHS4 | Eg08_t020390 | EgCHS3 | 0.035020233 | 0.316611117 | 0.110609613 | 70.3580259 |
| Mbe01_t047580 | MbeCHS4 | Eg08_t049400 | EgCHS5 | 0.1014938 | 0.347533796 | 0.292040088 | 77.22973239 |
| Mbe11_t002790 | MbeCHS5 | Eg08_t049400 | EgCHS5 | 0.103976596 | 0.359949685 | 0.288864249 | 79.98881897 |
| Mbe11_t002790 | MbeCHS5 | Eg08_t020390 | EgCHS3 | 0.013572472 | 0.066260109 | 0.20483625 | 14.7244686 |
| Mbe11_t002790 | MbeCHS5 | Eg08_t040870 | EgCHS2 | 0.031490171 | 0.306411099 | 0.102770986 | 68.09135532 |
| Mbe04_t036630 | MbeDFR1 | Eg09_t036940 | EgDFR1 | 0.026225889 | 0.094631686 | 0.277136445 | 21.0292636 |
| Mbe04_t011700 | MbeDFR3 | Eg04_t027830 | EgDFR2 | 0.01954906 | 0.055070601 | 0.354981784 | 12.23791143 |
| Mbe07_t008510 | MbeF3'5'H1 | Eg02_t007770 | EgF3'5'H2 | 0.022786563 | 0.068771849 | 0.331335613 | 15.28263311 |
| Mbe07_t005300 | MbeF3'5'H10 | Eg02_t010650 | EgF3'5'H5 | 0.046562661 | 0.097605274 | 0.477050668 | 21.690061 |
| Mbe06_t049500 | MbeF3'5'H2 | Eg09_t010710 | EgF3'5'H10 | 0.01414094 | 0.076157433 | 0.18568037 | 16.92387401 |
| Mbe02_t005590 | MbeF3'5'H3 | Eg02_t019070 | EgF3'5'H6 | 0.095564557 | 0.474424643 | 0.201432531 | 105.4276985 |
| Mbe02_t005590 | MbeF3'5'H3 | Eg02_t019180 | EgF3'5'H9 | 0.016849659 | 0.09976735 | 0.168889514 | 22.17052223 |
| Mbe02_t005610 | MbeF3'5'H4 | Eg02_t019100 | EgF3'5'H7 | 0.155747086 | 0.316963641 | 0.491372089 | 70.43636456 |
| Mbe02_t006410 | MbeF3'5'H5 | Eg02_t019070 | EgF3'5'H6 | 0.026726902 | 0.128562607 | 0.207890169 | 28.56946812 |
| Mbe02_t006410 | MbeF3'5'H5 | Eg02_t019180 | EgF3'5'H9 | 0.096886161 | 0.431106961 | 0.224738104 | 95.80154696 |

| | | | | | | | |
|---|---|---|---|---|---|---|---|
| Mbe02_t005600 | MbeF3'5'H6 | Eg02_t019110 | EgF3'5'H8 | 0.0612697 | 0.132616347 | 0.462007147 | 29.47029925 |
| Mbe08_t039500 | MbeF3'5'H7 | Eg06_t004020 | EgF3'5'H3 | 0.036770421 | 0.128562607 | 0.28601179 | 28.56946812 |
| Mbe09_t008950 | MbeF3'5'H8 | Eg07_t042500 | EgF3'5'H4 | 0.020826979 | 0.103686455 | 0.200864992 | 23.04143442 |
| Mbe06_t019910 | MbeF3'H | Eg03_t018110 | EgF3'H1 | 0.04026328 | 0.114778749 | 0.350790367 | 25.50638878 |
| Mbe07_t035880 | MbeF3H1 | Eg07_t027880 | EgF3H1 | 0.020195241 | 0.149354476 | 0.135216845 | 33.1898836 |
| Mbe08_t015900 | MbeFLS1 | Eg03_t031640 | EgFLS4 | 0.025268347 | 0.066909126 | 0.377651734 | 14.8686946 |
| Mbe02_t047300 | MbeFLS2 | Eg08_t007020 | EgFLS3 | 0.031877308 | 0.104397848 | 0.305344494 | 23.19952169 |
| Mbe03_t008100 | MbeFLS4 | Eg01_t007540 | EgFLS2 | 0.007886162 | 0.100377354 | 0.078565156 | 22.3060787 |
| Mbe05_t014700 | MbeLAR | Eg05_t019870 | EgLAR | 0.030955495 | 0.117054756 | 0.264453119 | 26.012168 |
| Mbe03_t031300 | MbeLDOX | Eg05_t014690 | EgLDOX | 0.025669648 | 0.076391062 | 0.336029461 | 16.97579166 |
| Mbe05_t009880 | MbePAL1 | Eg02_t001010 | EgPAL6 | 0.06394699 | 0.412667672 | 0.154960019 | 91.70392722 |
| Mbe05_t009880 | MbePAL1 | Eg05_t024420 | EgPAL8 | 0.00815609 | 0.067006623 | 0.121720658 | 14.89036062 |
| Mbe05_t009880 | MbePAL1 | Eg09_t017980 | EgPAL7 | 0.037499799 | 0.396747313 | 0.094518093 | 88.16606957 |
| Mbe06_t040740 | MbePAL2 | Eg05_t024420 | EgPAL8 | 0.034974975 | 0.362222832 | 0.096556518 | 80.49396268 |
| Mbe06_t040740 | MbePAL2 | Eg09_t017980 | EgPAL7 | 0.013826724 | 0.091656191 | 0.150854234 | 20.36804247 |
| Mbe07_t015550 | MbePAL3 | Eg02_t001010 | EgPAL6 | 0.01826862 | 0.072612932 | 0.251589067 | 16.13620707 |
| Mbe07_t015550 | MbePAL3 | Eg02_t018790 | EgPAL4 | 0.058359751 | 0.370536095 | 0.157500852 | 82.34135434 |
| Mbe06_t048840 | MbePAL5 | Eg02_t001010 | EgPAL6 | 0.085861938 | 0.386316199 | 0.222258186 | 85.84804432 |
| Mbe06_t048840 | MbePAL5 | Eg02_t018790 | EgPAL4 | 0.06818482 | 0.427384155 | 0.159539888 | 94.97425675 |
| Mbe06_t048840 | MbePAL5 | Eg09_t011330 | EgPAL5 | 0.025059889 | 0.059962 | 0.417929497 | 13.32488898 |
| Mbe01_t022110 | MbePAL7 | Eg01_t040920 | EgPAL1 | 0.011582637 | 0.07695211 | 0.150517476 | 17.10046898 |
| Mbe08_t037020 | MbePAL8 | Eg06_t005550 | EgPAL9 | 0.007673456 | 0.058548459 | 0.131061621 | 13.01076877 |
| Mbe06_t034420 | MbeTT8 | Eg03_t004200 | EgTT8.2 | 0.029540166 | 0.137254456 | 0.215221911 | 30.50099024 |
| Mbe06_t001720 | MbeTT8 | Eg04_t020700 | EgTT8 | 0.035710727 | 0.128057456 | 0.278864877 | 28.45721249 |
| Mbe03_t038300 | MbeTTG1 | Eg05_t009200 | EgTTG1 | 0.014790462 | 0.130086288 | 0.113697314 | 28.90806402 |

| | | | | | | | |
|---|---|---|---|---|---|---|---|
| Mbe04_t041380 | MbeTTG1 | Eg09_t041330 | EgTTG1.2 | 0.015819547 | 0.110329134 | 0.143385039 | 24.51758544 |
| Mbe01_t034870 | MbeUGT75C1.3 | Eg08_t037030 | EgUGT75C1.2 | 0.071319721 | 0.1107188 | 0.644151862 | 24.60417784 |
| Mbe02_t045890 | MbeUGT78D2 | Eg08_t008340 | EgUGT78D2 | 0.037937959 | 0.115138777 | 0.329497676 | 25.58639478 |
| Mbe07_t034770 | MbeUGT78D2 | Eg06_t008220 | EgUGT78D2.2 | 0.024917166 | 0.067976597 | 0.366555062 | 15.10591048 |
| Mbe03_t010580 | MbeUGT78D2.3 | Eg01_t005290 | EgUGT78D2.3 | 0.030374287 | 0.104516641 | 0.290616755 | 23.22592015 |
| Mbe08_t027530 | MbeUGT79B1 | Eg07_t037900 | EgUGT79B1 | 0.036338449 | 0.081561768 | 0.445532874 | 18.12483734 |

**Table S3h.** Divergence time, Ka, Ks and Ka/Ks values of orthologous flavonoid biosynthetic genes pairs between *M. beccarii* and *M. acuminata*

| **Seq_1** | **Name1** | **Seq_2** | **Name2** | **Ka** | **Ks** | **Ka_Ks** | **Diversity_year** |
|---|---|---|---|---|---|---|---|
| Mbe06_t016280 | Mbe4CL1 | Parent=Macma4_06_g39580 | Ma4CL9 | 0.022346481 | 0.095960332 | 0.232872065 | 21.32451819 |
| Mbe07_t013640 | Mbe4CL10 | Parent=Macma4_08_g13040 | Ma4CL1 | 0.028703559 | 0.077375288 | 0.370965457 | 17.19450855 |
| Mbe06_t040140 | Mbe4CL12 | Parent=Macma4_11_g22750 | Ma4CL2.2 | 0.011031059 | 0.087126445 | 0.126609768 | 19.36143223 |
| Mbe03_t056570 | Mbe4CL13 | Parent=Macma4_09_g24760 | Ma4CL16 | 0.018246805 | 0.058516634 | 0.311822531 | 13.00369652 |
| Mbe01_t005810 | Mbe4CL14 | Parent=Macma4_01_g19800 | Ma4CL10 | 0.009017118 | 0.073028919 | 0.12347325 | 16.22864863 |
| Mbe01_t005810 | Mbe4CL14 | Parent=Macma4_03_g13930 | Ma4CL11 | 0.040157357 | 0.352052587 | 0.11406636 | 78.2339082 |
| Mbe01_t005810 | Mbe4CL14 | Parent=Macma4_07_g05220 | Ma4CL14 | 0.079952596 | 0.474385963 | 0.168539126 | 105.4191029 |
| Mbe03_t000920 | Mbe4CL15 | Parent=Macma4_01_g19800 | Ma4CL10 | 0.040174166 | 0.328863615 | 0.122160569 | 73.08080324 |
| Mbe03_t000920 | Mbe4CL15 | Parent=Macma4_03_g13930 | Ma4CL11 | 0.012321449 | 0.04961511 | 0.248340655 | 11.02558009 |
| Mbe03_t000920 | Mbe4CL15 | Parent=Macma4_07_g05220 | Ma4CL14 | 0.076772329 | 0.485633162 | 0.158087082 | 107.9184804 |
| Mbe07_t022330 | Mbe4CL16 | Parent=Macma4_01_g19800 | Ma4CL10 | 0.066804516 | 0.415380401 | 0.160827318 | 92.30675578 |
| Mbe07_t022330 | Mbe4CL16 | Parent=Macma4_03_g13930 | Ma4CL11 | 0.067533619 | 0.48300948 | 0.139818413 | 107.33544 |
| Mbe07_t022330 | Mbe4CL16 | Parent=Macma4_07_g05220 | Ma4CL14 | 0.023335216 | 0.063543488 | 0.367232214 | 14.12077501 |
| Mbe04_t006230 | Mbe4CL17 | Parent=Macma4_04_g06110 | Ma4CL12 | 0.014132735 | 0.084110961 | 0.168024896 | 18.69132468 |
| Mbe04_t006230 | Mbe4CL17 | Parent=Macma4_04_g13130 | Ma4CL13 | 0.044644302 | 0.336495894 | 0.132674136 | 74.77686538 |
| Mbe04_t013920 | Mbe4CL18 | Parent=Macma4_04_g13130 | Ma4CL13 | 0.010708584 | 0.061520381 | 0.17406563 | 13.67119579 |

| | | | | | | | |
|---|---|---|---|---|---|---|---|
| Mbe04_t013920 | Mbe4CL18 | Parent=Macma4_04_g06110 | Ma4CL12 | 0.040918279 | 0.396103457 | 0.103301999 | 88.02299046 |
| Mbe03_t000510 | Mbe4CL2 | Parent=Macma4_03_g14310 | Ma4CL8 | 0.036871087 | 0.080043579 | 0.460637661 | 17.78746206 |
| Mbe07_t014630 | Mbe4CL3 | Parent=Macma4_08_g13960 | Ma4CL4 | 0.010075718 | 0.067697559 | 0.148834292 | 15.04390206 |
| Mbe05_t019740 | Mbe4CL4 | Parent=Macma4_07_g18890 | Ma4CL15 | 0.020592961 | 0.08111312 | 0.253879531 | 18.02513779 |
| Mbe01_t019320 | Mbe4CL6 | Parent=Macma4_01_g07350 | Ma4CL7 | 0.027588841 | 0.104692896 | 0.26352161 | 23.26508808 |
| Mbe05_t004680 | Mbe4CL7 | Parent=Macma4_05_g28920 | Ma4CL5 | 0.016328884 | 0.064835163 | 0.251852294 | 14.40781393 |
| Mbe03_t020730 | Mbe4CL8 | Parent=Macma4_11_g06130 | Ma4CL6 | 0.034417316 | 0.075575831 | 0.455401085 | 16.79462915 |
| Mbe05_t011100 | Mbe4CL9 | Parent=Macma4_05_g22890 | Ma4CL1.2 | 0.018244283 | 0.055475815 | 0.328869127 | 12.32795893 |
| Mbe07_t028460 | MbeANR | Parent=Macma4_07_g10660 | MaANR2 | 0.039643374 | 0.093468426 | 0.424136532 | 20.77076125 |
| Mbe03_t051550 | MbeC1 | Parent=Macma4_09_g29590 | MaC1.2 | 0.029100784 | 0.080037796 | 0.363588017 | 17.78617687 |
| Mbe02_t039140 | MbeC1.3 | Parent=Macma4_10_g21630 | MaC1 | 0.018993365 | 0.135069213 | 0.140619501 | 30.01538064 |
| Mbe01_t038520 | MbeC4H1 | Parent=Macma4_06_g10450 | MaC4H3 | 0.00788386 | 0.048576841 | 0.162296673 | 10.79485364 |
| Mbe10_t000860 | MbeC4H2 | Parent=Macma4_07_g20380 | MaC4H2 | 0.005243249 | 0.06362194 | 0.082412591 | 14.13820879 |
| Mbe02_t033650 | MbeC4H3 | Parent=Macma4_06_g10450 | MaC4H3 | 0.036633501 | 0.310074958 | 0.118144015 | 68.90554629 |
| Mbe02_t033650 | MbeC4H3 | Parent=Macma4_10_g16840 | MaC4H1 | 0.011435495 | 0.045361012 | 0.252099644 | 10.08022486 |
| Mbe08_t025090 | MbeC4H4 | Parent=Macma4_09_g04800 | MaC4H4 | 0.008713429 | 0.043532527 | 0.200159052 | 9.673894942 |
| Mbe04_t020690 | MbeCHI1 | Parent=Macma4_04_g18950 | MaCHI1 | 0.041691182 | 0.066778156 | 0.62432365 | 14.83959027 |
| Mbe06_t039860 | MbeCHI2 | Parent=Macma4_11_g22930 | MaCHI2 | 0.023022387 | 0.097760208 | 0.235498545 | 21.72449057 |
| Mbe01_t038030 | MbeCHS2 | Parent=Macma4_06_g10010 | MaCHS5 | 0.037056423 | 0.132735585 | 0.279174746 | 29.49679669 |
| Mbe01_t038030 | MbeCHS2 | Parent=Macma4_06_g18170 | MaCHS2 | 0.112781267 | 0.343201107 | 0.328615686 | 76.26691266 |
| Mbe01_t038030 | MbeCHS2 | Parent=Macma4_10_g16140 | MaCHS6 | 0.111797989 | 0.382389646 | 0.292366674 | 84.97547684 |
| Mbe01_t041060 | MbeCHS3 | Parent=Macma4_06_g12560 | MaCHS1 | 0.012454291 | 0.065875907 | 0.189056847 | 14.63909042 |
| Mbe01_t047580 | MbeCHS4 | Parent=Macma4_06_g18170 | MaCHS2 | 0.014166065 | 0.04143954 | 0.34184899 | 9.20878656 |
| Mbe01_t047580 | MbeCHS4 | Parent=Macma4_06_g10010 | MaCHS5 | 0.10983443 | 0.373831173 | 0.293807574 | 83.07359409 |
| Mbe01_t047580 | MbeCHS4 | Parent=Macma4_10_g16140 | MaCHS6 | 0.0326668 | 0.338612226 | 0.096472595 | 75.24716144 |

Musaceae flavonoid biosynthetic genes

| | | | | | | | |
|---|---|---|---|---|---|---|---|
| Mbe11_t002790 | MbeCHS5 | Parent=Macma4_06_g18170 | MaCHS2 | 0.039338811 | 0.292112523 | 0.134670061 | 64.91389408 |
| Mbe11_t002790 | MbeCHS5 | Parent=Macma4_10_g16140 | MaCHS6 | 0.009019274 | 0.036108056 | 0.249785656 | 8.02401241 |
| Mbe04_t036630 | MbeDFR1 | Parent=Macma4_04_g33220 | MaDFR1 | 0.017789906 | 0.082779185 | 0.214907968 | 18.39537452 |
| Mbe06_t019010 | MbeDFR2 | Parent=Macma4_03_g33910 | MaDFR2 | 0.018615227 | 0.10633488 | 0.175062281 | 23.62997325 |
| Mbe04_t011700 | MbeDFR3 | Parent=Macma4_04_g11180 | MaDFR3 | 0.013374215 | 0.073071347 | 0.183029534 | 16.23807712 |
| Mbe07_t008510 | MbeF3'5'H1 | Parent=Macma4_08_g08290 | MaF3'5'H1 | 0.016903377 | 0.070358562 | 0.240246198 | 15.63523591 |
| Mbe07_t005300 | MbeF3'5'H10 | Parent=Macma4_08_g05480 | MaF3'5'H3 | 0.051141161 | 0.104782026 | 0.488071891 | 23.28489459 |
| Mbe06_t049500 | MbeF3'5'H2 | Parent=Macma4_11_g15260 | MaF3'5'H13 | 0.013726307 | 0.065364483 | 0.209996419 | 14.52544069 |
| Mbe02_t005590 | MbeF3'5'H3 | Parent=Macma4_02_g03290 | MaF3'5'H11 | 0.084989209 | 0.344417286 | 0.246762323 | 76.53717464 |
| Mbe02_t005610 | MbeF3'5'H4 | Parent=Macma4_02_g03300 | MaF3'5'H12 | 0.26455997 | 0.587838341 | 0.450055655 | 130.6307424 |
| Mbe02_t006410 | MbeF3'5'H5 | Parent=Macma4_02_g03040 | MaF3'5'H7 | 0.077005053 | 0.138666714 | 0.555324712 | 30.81482538 |
| Mbe02_t005600 | MbeF3'5'H6 | Parent=Macma4_02_g03260 | MaF3'5'H10 | 0.048718422 | 0.162695314 | 0.299445761 | 36.15451433 |
| Mbe08_t039500 | MbeF3'5'H7 | Parent=Macma4_09_g17660 | MaF3'5'H4 | 0.024037666 | 0.049787861 | 0.482801737 | 11.06396911 |
| Mbe09_t008950 | MbeF3'5'H8 | Parent=Macma4_09_g17660 | MaF3'5'H4 | 0.117104106 | 0.440303758 | 0.265962087 | 97.84527949 |
| Mbe09_t008950 | MbeF3'5'H8 | Parent=Macma4_10_g01260 | MaF3'5'H6 | 0.011843325 | 0.058476157 | 0.20253255 | 12.99470165 |
| Mbe06_t019910 | MbeF3'H | Parent=Macma4_03_g33170 | MaF3'H | 0.03306928 | 0.099925827 | 0.330938267 | 22.2057394 |
| Mbe07_t035880 | MbeF3H1 | Parent=Macma4_07_g17110 | MaF3H1 | 0.017801469 | 0.066710615 | 0.266846119 | 14.824581 |
| Mbe08_t015900 | MbeFLS1 | Parent=Macma4_08_g24690 | MaFLS2 | 0.021202825 | 0.057806378 | 0.366790418 | 12.84586172 |
| Mbe03_t008100 | MbeFLS4 | Parent=Macma4_03_g07220 | MaFLS1 | 0.014523504 | 0.076327492 | 0.190278801 | 16.96166487 |
| Mbe05_t014700 | MbeLAR | Parent=Macma4_05_g19560 | MaLAR | 0.020608788 | 0.083446878 | 0.246968955 | 18.5437507 |
| Mbe03_t031300 | MbeLDOX | Parent=Macma4_05_g03920 | MaLDOX | 0.01209216 | 0.033200441 | 0.364216857 | 7.377875842 |
| Mbe05_t009880 | MbePAL1 | Parent=Macma4_05_g21700 | MaPAL5 | 0.010975423 | 0.043972214 | 0.249599059 | 9.77160309 |
| Mbe05_t009880 | MbePAL1 | Parent=Macma4_11_g22170 | MaPAL6 | 0.035791444 | 0.34818076 | 0.102795583 | 77.37350225 |
| Mbe06_t040740 | MbePAL2 | Parent=Macma4_05_g21700 | MaPAL5 | 0.041128139 | 0.342568802 | 0.12005804 | 76.12640042 |
| Mbe06_t040740 | MbePAL2 | Parent=Macma4_11_g22170 | MaPAL6 | 0.007496548 | 0.058824846 | 0.12743847 | 13.07218806 |

| | | | | | | | |
|---|---|---|---|---|---|---|---|
| Mbe07_t015550 | MbePAL3 | Parent=Macma4_02_g02760 | MaPAL4 | 0.059828397 | 0.352447054 | 0.169751447 | 78.32156746 |
| Mbe07_t015550 | MbePAL3 | Parent=Macma4_08_g14800 | MaPAL7 | 0.010014753 | 0.064382671 | 0.155550449 | 14.30726031 |
| Mbe06_t048840 | MbePAL5 | Parent=Macma4_02_g02760 | MaPAL4 | 0.063233637 | 0.392271547 | 0.161198633 | 87.17145493 |
| Mbe06_t048840 | MbePAL5 | Parent=Macma4_11_g15930 | MaPAL3 | 0.019412277 | 0.056611572 | 0.342902991 | 12.58034939 |
| Mbe01_t022110 | MbePAL7 | Parent=Macma4_01_g04800 | MaPAL1 | 0.013464509 | 0.066710615 | 0.201834588 | 14.824581 |
| Mbe08_t037020 | MbePAL8 | Parent=Macma4_09_g16190 | MaPAL8 | 0.004121901 | 0.043483875 | 0.094791473 | 9.663083325 |
| Mbe06_t034420 | MbeTT8 | Parent=Macma4_03_g19880 | MaTT8.2 | 0.021164221 | 0.092312077 | 0.229268177 | 20.51379479 |
| Mbe06_t001720 | MbeTT8 | Parent=Macma4_06_g25800 | MaTT8 | 0.047226502 | 0.058162737 | 0.811971791 | 12.92505272 |
| Mbe03_t038300 | MbeTTG1 | Parent=Macma4_05_g10420 | MaTTG1 | 0.010604006 | 0.067397986 | 0.157334165 | 14.97733015 |
| Mbe04_t041380 | MbeTTG1 | Parent=Macma4_04_g37870 | MaTTG1.2 | 0.014488432 | 0.061657434 | 0.234982728 | 13.70165207 |
| Mbe01_t034880 | MbeUGT75C1 | Parent=Macma4_06_g07210 | MaUGT75C1.2 | 0.031445654 | 0.063021309 | 0.498968589 | 14.00473542 |
| Mbe02_t051150 | MbeUGT75C1 | Parent=Macma4_10_g31950 | MaUGT75C1 | 0.027471695 | 0.048926478 | 0.561489326 | 10.87255073 |
| Mbe01_t034870 | MbeUGT75C1.3 | Parent=Macma4_06_g07200 | MaUGT75C1.3 | 0.049253963 | 0.077184987 | 0.638128807 | 17.15221924 |
| Mbe02_t045890 | MbeUGT78D2 | Parent=Macma4_10_g27580 | MaUGT78D2 | 0.042160138 | 0.125638289 | 0.335567588 | 27.91961973 |
| Mbe07_t034770 | MbeUGT78D2 | Parent=Macma4_07_g16200 | MaUGT78D2.2 | 0.016863002 | 0.077181239 | 0.218485759 | 17.15138653 |
| Mbe03_t010580 | MbeUGT78D2.3 | Parent=Macma4_03_g04880 | MaUGT78D2.3 | 0.020112235 | 0.072002237 | 0.279327917 | 16.00049704 |
| Mbe08_t027530 | MbeUGT79B1 | Parent=Macma4_09_g07100 | MaUGT79B1 | 0.02523285 | 0.074355047 | 0.339356259 | 16.52334371 |

**Table S3i.** Divergence time, Ka, Ks and Ka/Ks values of orthologous flavonoid biosynthetic genes pairs between *M. beccarii* and *M. balbisiana*

| Seq_1 | Name1 | Seq_2 | Name2 | Ka | Ks | Ka_Ks | Diversity_year |
|---|---|---|---|---|---|---|---|
| Mbe06_t016280 | Mbe4CL1 | Mba06_g36630 | Mb4CL9 | 0.02400536 | 0.090575557 | 0.265031326 | 20.12790153 |
| Mbe07_t013640 | Mbe4CL10 | Mba08_g12720 | Mb4CL1.2 | 0.026878576 | 0.073057245 | 0.367911168 | 16.23494336 |
| Mbe06_t040140 | Mbe4CL12 | Mba11_g20940 | Mb4CL2.2 | 0.015015319 | 0.073966689 | 0.203001093 | 16.43704205 |
| Mbe01_t005810 | Mbe4CL14 | Mba01_g28370 | Mb4CL10 | 0.008195683 | 0.083595506 | 0.098039759 | 18.57677914 |
| Mbe03_t000920 | Mbe4CL15 | Mba01_g28370 | Mb4CL10 | 0.040331547 | 0.335289628 | 0.12028868 | 74.50880628 |

| | | | | | | | |
|---|---|---|---|---|---|---|---|
| Mbe07_t022330 | Mbe4CL16 | Mba01_g28370 | Mb4CL10 | 0.068626634 | 0.410607407 | 0.167134427 | 91.24609038 |
| Mbe07_t022330 | Mbe4CL16 | Mba07_g04890 | Mb4CL13 | 0.025348889 | 0.071306185 | 0.355493549 | 15.84581889 |
| Mbe04_t006230 | Mbe4CL17 | Mba04_g05720 | Mb4CL11 | 0.018339341 | 0.070759511 | 0.25917846 | 15.72433576 |
| Mbe04_t006230 | Mbe4CL17 | Mba04_g12340 | Mb4CL12 | 0.044631727 | 0.360149382 | 0.123925597 | 80.03319599 |
| Mbe04_t013920 | Mbe4CL18 | Mba04_g12340 | Mb4CL12 | 0.012366315 | 0.075062594 | 0.164746702 | 16.68057643 |
| Mbe04_t013920 | Mbe4CL18 | Mba04_g05720 | Mb4CL11 | 0.044391654 | 0.375855239 | 0.118108381 | 83.52338634 |
| Mbe03_t000510 | Mbe4CL2 | Mba03_g13490 | Mb4CL8 | 0.091801832 | 0.158323822 | 0.579835871 | 35.18307164 |
| Mbe07_t014630 | Mbe4CL3 | Mba08_g13620 | Mb4CL4 | 0.017738111 | 0.084032105 | 0.211087316 | 18.67380104 |
| Mbe05_t019740 | Mbe4CL4 | Mba07_g17040 | Mb4CL14 | 0.01975965 | 0.095924944 | 0.205990739 | 21.31665417 |
| Mbe01_t019320 | Mbe4CL6 | Mba01_g16650 | Mb4CL7 | 0.028358011 | 0.077087997 | 0.367865448 | 17.13066601 |
| Mbe05_t004680 | Mbe4CL7 | Mba05_g27160 | Mb4CL5 | 0.02356931 | 0.075909095 | 0.310493886 | 16.86868781 |
| Mbe05_t011100 | Mbe4CL9 | Mba05_g21540 | Mb4CL1 | 0.016584424 | 0.055301424 | 0.29989144 | 12.28920531 |
| Mbe01_t033880 | MbeC1 | Mba06_g05630 | MbC1.2 | 0.025815939 | 0.082728407 | 0.312056528 | 18.38409034 |
| Mbe03_t051550 | MbeC1 | Mba09_g23680 | MbC1.3 | 0.147611107 | 0.271515804 | 0.543655672 | 60.33684525 |
| Mbe02_t039140 | MbeC1.3 | Mba10_g15520 | MbC1 | 0.019020964 | 0.110239313 | 0.172542476 | 24.49762501 |
| Mbe01_t038520 | MbeC4H1 | Mba06_g09560 | MbC4H3 | 0.007009397 | 0.066325612 | 0.105681599 | 14.73902499 |
| Mbe10_t000860 | MbeC4H2 | Mba07_g18380 | MbC4H4 | 0.002616472 | 0.069733232 | 0.037521162 | 15.4962737 |
| Mbe02_t033650 | MbeC4H3 | Mba06_g09560 | MbC4H3 | 0.033006664 | 0.334478731 | 0.098680906 | 74.32860678 |
| Mbe08_t025090 | MbeC4H4 | Mba09_g04380 | MbC4H1 | 0.016643019 | 0.046526543 | 0.357710192 | 10.33923182 |
| Mbe04_t020690 | MbeCHI1 | Mba04_g18040 | MbCHI2 | 0.057765709 | 0.102431651 | 0.563943942 | 22.76258917 |
| Mbe06_t039860 | MbeCHI2 | Mba11_g21110 | MbCHI1 | 0.020885443 | 0.114289091 | 0.182742226 | 25.39757572 |
| Mbe01_t038030 | MbeCHS2 | Mba06_g09120 | MbCHS7 | 0.012214802 | 0.104002719 | 0.117446945 | 23.11171528 |
| Mbe01_t038030 | MbeCHS2 | Mba06_g16830 | MbCHS2 | 0.104317132 | 0.363048407 | 0.287336701 | 80.6774238 |
| Mbe01_t041060 | MbeCHS3 | Mba06_g11590 | MbCHS1 | 0.006769508 | 0.050682729 | 0.133566371 | 11.26282859 |
| Mbe01_t047580 | MbeCHS4 | Mba06_g16830 | MbCHS2 | 0.006765056 | 0.05455297 | 0.124008936 | 12.1228822 |

| | | | | | | | |
|---|---|---|---|---|---|---|---|
| Mbe01_t047580 | MbeCHS4 | Mba10_g10820 | MbCHS8 | 0.03033607 | 0.33823646 | 0.08968894 | 75.16365782 |
| Mbe11_t002790 | MbeCHS5 | Mba10_g10820 | MbCHS8 | 0.006756168 | 0.047245408 | 0.143001588 | 10.49897962 |
| Mbe04_t036630 | MbeDFR1 | Mba04_g31800 | MbDFR1 | 0.0128093 | 0.086160207 | 0.1486684 | 19.14671267 |
| Mbe06_t019010 | MbeDFR2 | Mba01_g01310 | MbDFR2 | 0.017332867 | 0.093241603 | 0.185891986 | 20.72035621 |
| Mbe07_t008510 | MbeF3'5'H1 | Mba08_g07950 | MbF3'5'H1 | 0.012412548 | 0.067508649 | 0.183866037 | 15.00192205 |
| Mbe06_t049500 | MbeF3'5'H2 | Mba11_g13650 | MbF3'5'H13 | 0.012384182 | 0.072716435 | 0.170307887 | 16.15920768 |
| Mbe02_t005590 | MbeF3'5'H3 | Mba02_g01950 | MbF3'5'H12 | 0.012376883 | 0.105940449 | 0.116828684 | 23.54232193 |
| Mbe02_t005610 | MbeF3'5'H4 | Mba02_g01880 | MbF3'5'H6 | 0.157999701 | 0.255337944 | 0.618786609 | 56.74176537 |
| Mbe02_t006410 | MbeF3'5'H5 | Mba02_g01950 | MbF3'5'H12 | 0.110202853 | 0.509436164 | 0.216323185 | 113.2080365 |
| Mbe02_t005600 | MbeF3'5'H6 | Mba02_g01940 | MbF3'5'H11 | 0.032142037 | 0.063755179 | 0.504147861 | 14.16781747 |
| Mbe08_t039500 | MbeF3'5'H7 | Mba09_g16030 | MbF3'5'H3 | 0.019561011 | 0.052374891 | 0.373480702 | 11.63886461 |
| Mbe09_t008950 | MbeF3'5'H8 | Mba10_g00740 | MbF3'5'H4 | 0.016218176 | 0.056558492 | 0.286750511 | 12.56855372 |
| Mbe06_t019910 | MbeF3'H | Mba01_g02030 | MbF3'H | 0.038946803 | 0.092874685 | 0.419347886 | 20.63881888 |
| Mbe07_t035880 | MbeF3H1 | Mba07_g15320 | MbF3H2 | 0.017797948 | 0.087574515 | 0.203232048 | 19.46100335 |
| Mbe08_t015900 | MbeFLS1 | Mba08_g23630 | MbFLS1 | 0.016029754 | 0.091290257 | 0.175591071 | 20.28672382 |
| Mbe02_t047300 | MbeFLS2 | Mba10_g21980 | MbFLS3 | 0.025850344 | 0.068253888 | 0.378738045 | 15.16753065 |
| Mbe08_t036240 | MbeFLS3 | Mba09_g14060 | MbFLS5 | 0.083368337 | 0.145369652 | 0.573492028 | 32.30436722 |
| Mbe03_t008100 | MbeFLS4 | Mba03_g06940 | MbFLS4 | 0.014536299 | 0.080825798 | 0.17984727 | 17.96128843 |
| Mbe03_t008100 | MbeFLS4 | Mba09_g14060 | MbFLS5 | 0.149641669 | 0.517019962 | 0.289431124 | 114.8933249 |
| Mbe05_t014700 | MbeLAR | Mba05_g18490 | MbLAR | 0.028868641 | 0.068092097 | 0.42396464 | 15.13157713 |
| Mbe03_t031300 | MbeLDOX | Mba05_g13780 | MbLDOX | 0.062273669 | 0.126456267 | 0.492452218 | 28.10139275 |
| Mbe06_t040740 | MbePAL2 | Mba11_g20430 | MbPAL2 | 0.009381474 | 0.056806388 | 0.165148227 | 12.62364178 |
| Mbe07_t015550 | MbePAL3 | Mba08_g14430 | MbPAL3 | 0.009071115 | 0.065411864 | 0.138676901 | 14.53596987 |
| Mbe06_t048840 | MbePAL5 | Mba11_g14220 | MbPAL6 | 0.030104148 | 0.073469868 | 0.409748221 | 16.32663741 |
| Mbe01_t022110 | MbePAL7 | Mba01_g14280 | MbPAL8 | 0.011891352 | 0.067662693 | 0.17574458 | 15.03615408 |

| | | | | | | | |
|---|---|---|---|---|---|---|---|
| Mbe08_t037020 | MbePAL8 | Mba09_g14610 | MbPAL1 | 0.005895677 | 0.047216799 | 0.124863979 | 10.49262203 |
| Mbe06_t034420 | MbeTT8 | Mba03_g18730 | MbTT8.2 | 0.104261625 | 0.155083736 | 0.672292451 | 34.4630524 |
| Mbe06_t034420 | MbeTT8 | Mba06_g23950 | MbTT8 | 0.110492037 | 0.46406338 | 0.238096868 | 103.1251956 |
| Mbe03_t038300 | MbeTTG1 | Mba05_g07670 | MbTTG1 | 0.012984782 | 0.075203837 | 0.172661168 | 16.71196375 |
| Mbe04_t041380 | MbeTTG1 | Mba04_g36060 | MbTTG1.2 | 0.011830676 | 0.065999274 | 0.179254637 | 14.66650523 |
| Mbe02_t051150 | MbeUGT75C1 | Mba10_g24560 | MbUGT75C1 | 0.024611768 | 0.039914406 | 0.616613664 | 8.869868045 |
| Mbe01_t034870 | MbeUGT75C1.3 | Mba06_g06490 | MbUGT75C1.3 | 0.133288396 | 0.328032643 | 0.406326623 | 72.89614281 |
| Mbe02_t045890 | MbeUGT78D2 | Mba10_g20800 | MbUGT78D2.2 | 0.030631872 | 0.096217156 | 0.318361857 | 21.3815902 |
| Mbe07_t034770 | MbeUGT78D2 | Mba07_g14580 | MbUGT78D2.3 | 0.015847404 | 0.068026895 | 0.232957916 | 15.11708779 |
| Mbe03_t010580 | MbeUGT78D2.3 | Mba03_g04830 | MbUGT78D2 | 0.015036527 | 0.096193127 | 0.156316026 | 21.37625054 |

**Table S3j.** Divergence time, Ka, Ks and Ka/Ks values of orthologous flavonoid biosynthetic genes pairs between *M. beccarii* and *M. schizocarpa*

| Seq_1 | Name1 | Seq_2 | Name2 | Ka | Ks | Ka_Ks | Diversity_year |
|---|---|---|---|---|---|---|---|
| Mbe06_t016280 | Mbe4CL1 | Ms06t178240 | Ms4CL9 | 0.023185452 | 0.110339622 | 0.210128074 | 24.51991593 |
| Mbe07_t013640 | Mbe4CL10 | Ms08t218760 | Ms4CL1 | 0.029572966 | 0.072075286 | 0.410306613 | 16.01673022 |
| Mbe02_t027270 | Mbe4CL11 | Ms02t041000 | Ms4CL2 | 0.023896204 | 0.078202793 | 0.305567146 | 17.37839844 |
| Mbe06_t040140 | Mbe4CL12 | Ms11t319670 | Ms4CL2.2 | 0.010237723 | 0.089770784 | 0.114042926 | 19.94906322 |
| Mbe03_t056570 | Mbe4CL13 | Ms04t089570 | Ms4CL13 | 0.052154378 | 0.45144524 | 0.115527584 | 100.3211643 |
| Mbe03_t056570 | Mbe4CL13 | Ms09t262230 | Ms4CL16 | 0.021602158 | 0.055958148 | 0.386041332 | 12.43514397 |
| Mbe01_t005810 | Mbe4CL14 | Ms01t017810 | Ms4CL10 | 0.009022049 | 0.08622531 | 0.104633413 | 19.16118001 |
| Mbe01_t005810 | Mbe4CL14 | Ms03t058920 | Ms4CL11 | 0.039711163 | 0.354519054 | 0.112014184 | 78.7820121 |
| Mbe01_t005810 | Mbe4CL14 | Ms07t183930 | Ms4CL14 | 0.080334157 | 0.487716512 | 0.16471486 | 108.3814471 |
| Mbe03_t000920 | Mbe4CL15 | Ms01t017810 | Ms4CL10 | 0.0401966 | 0.34698525 | 0.115845271 | 77.10783333 |
| Mbe03_t000920 | Mbe4CL15 | Ms03t058920 | Ms4CL11 | 0.011488975 | 0.054830303 | 0.209536963 | 12.18451188 |

| | | | | | | | |
|---|---|---|---|---|---|---|---|
| Mbe03_t000920 | Mbe4CL15 | Ms07t183930 | Ms4CL14 | 0.075338023 | 0.485108771 | 0.155301302 | 107.801949 |
| Mbe07_t022330 | Mbe4CL16 | Ms01t017810 | Ms4CL10 | 0.068641396 | 0.406092075 | 0.169029143 | 90.24268324 |
| Mbe07_t022330 | Mbe4CL16 | Ms03t058920 | Ms4CL11 | 0.066607248 | 0.493183151 | 0.135055807 | 109.5962557 |
| Mbe07_t022330 | Mbe4CL16 | Ms07t183930 | Ms4CL14 | 0.024160367 | 0.077139221 | 0.313204711 | 17.14204914 |
| Mbe04_t006230 | Mbe4CL17 | Ms04t083350 | Ms4CL12 | 0.016645388 | 0.08967803 | 0.185612777 | 19.92845104 |
| Mbe04_t006230 | Mbe4CL17 | Ms04t089570 | Ms4CL13 | 0.046387644 | 0.344175142 | 0.134779183 | 76.48336493 |
| Mbe04_t013920 | Mbe4CL18 | Ms04t089570 | Ms4CL13 | 0.012369714 | 0.066882334 | 0.184947405 | 14.86274093 |
| Mbe04_t013920 | Mbe4CL18 | Ms04t083350 | Ms4CL12 | 0.04393457 | 0.403182143 | 0.108969533 | 89.59603168 |
| Mbe03_t000510 | Mbe4CL2 | Ms03t059300 | Ms4CL8 | 0.043272474 | 0.07875672 | 0.549444844 | 17.50149334 |
| Mbe07_t014630 | Mbe4CL3 | Ms08t219500 | Ms4CL4 | 0.010921504 | 0.075062594 | 0.145498616 | 16.68057643 |
| Mbe05_t019740 | Mbe4CL4 | Ms07t196560 | Ms4CL15 | 0.025948534 | 0.083595506 | 0.310405849 | 18.57677914 |
| Mbe01_t019320 | Mbe4CL6 | Ms01t006680 | Ms4CL7 | 0.027170122 | 0.087766283 | 0.30957357 | 19.50361849 |
| Mbe05_t004680 | Mbe4CL7 | Ms05t142250 | Ms4CL5 | 0.019725555 | 0.072230891 | 0.273090285 | 16.05130919 |
| Mbe05_t011100 | Mbe4CL9 | Ms05t136800 | Ms4CL1.2 | 0.01657755 | 0.057987711 | 0.285880404 | 12.88615793 |
| Mbe07_t028460 | MbeANR | Ms07t188910 | MsANR3 | 0.039327929 | 0.105907211 | 0.371343263 | 23.53493571 |
| Mbe01_t033880 | MbeC1 | Ms06t151760 | MsC1.2 | 0.02747412 | 0.076779313 | 0.357832318 | 17.0620696 |
| Mbe03_t051550 | MbeC1 | Ms09t266370 | MsC1 | 0.039664415 | 0.067971013 | 0.58354897 | 15.10466958 |
| Mbe01_t038520 | MbeC4H1 | Ms10t281630 | MsC4H1 | 0.033011573 | 0.321501004 | 0.102679533 | 71.44466753 |
| Mbe10_t000860 | MbeC4H2 | Ms07t197940 | MsC4H2 | 0.005243249 | 0.078866367 | 0.066482698 | 17.52585929 |
| Mbe02_t033650 | MbeC4H3 | Ms10t281630 | MsC4H1 | 0.011442205 | 0.045275413 | 0.252724478 | 10.06120298 |
| Mbe08_t025090 | MbeC4H4 | Ms09t244050 | MsC4H3 | 0.010469852 | 0.046503998 | 0.225138743 | 10.33422172 |
| Mbe04_t020690 | MbeCHI1 | Ms04t093250 | MsCHI2 | 0.035650275 | 0.072784342 | 0.489806934 | 16.17429819 |
| Mbe06_t039860 | MbeCHI2 | Ms11t319860 | MsCHI1 | 0.022998318 | 0.114434691 | 0.200973302 | 25.42993127 |
| Mbe01_t038030 | MbeCHS2 | MsSC51t327380 | MsCHS4 | 0.011532872 | 0.091712473 | 0.125750306 | 20.38054955 |
| Mbe01_t038030 | MbeCHS2 | Ms06t158650 | MsCHS2 | 0.116787417 | 0.345525927 | 0.337998998 | 76.78353943 |

| | | | | | | | |
|---|---|---|---|---|---|---|---|
| Mbe01_t038030 | MbeCHS2 | Ms10t280970 | MsCHS1 | 0.110495764 | 0.394148828 | 0.280340207 | 87.58862853 |
| Mbe01_t041060 | MbeCHS3 | MsSC66t327440 | MsCHS5 | 0.081540411 | 0.168281506 | 0.484547667 | 37.39589029 |
| Mbe01_t047580 | MbeCHS4 | Ms06t158650 | MsCHS2 | 0.018224131 | 0.062306872 | 0.292489907 | 13.84597162 |
| Mbe01_t047580 | MbeCHS4 | Ms10t280970 | MsCHS1 | 0.0326668 | 0.349738995 | 0.093403369 | 77.71977672 |
| Mbe11_t002790 | MbeCHS5 | Ms06t158650 | MsCHS2 | 0.043592535 | 0.322505347 | 0.13516841 | 71.66785494 |
| Mbe11_t002790 | MbeCHS5 | Ms10t280970 | MsCHS1 | 0.009019274 | 0.043543061 | 0.207134598 | 9.676235771 |
| Mbe04_t036630 | MbeDFR1 | Ms04t107810 | MsDFR1 | 0.017771087 | 0.073396948 | 0.242122976 | 16.31043281 |
| Mbe06_t019010 | MbeDFR2 | Ms03t076720 | MsDFR3 | 0.0198646 | 0.101923357 | 0.194897427 | 22.64963493 |
| Mbe04_t011700 | MbeDFR3 | Ms04t087800 | MsDFR2 | 0.012358673 | 0.069767041 | 0.177141997 | 15.50378695 |
| Mbe07_t008510 | MbeF3'5'H1 | Ms08t214430 | MsF3'5'H1 | 0.020746811 | 0.068120671 | 0.3045597 | 15.13792688 |
| Mbe07_t005300 | MbeF3'5'H10 | Ms08t211840 | MsF3'5'H3 | 0.04936988 | 0.10118761 | 0.487904394 | 22.48613562 |
| Mbe06_t049500 | MbeF3'5'H2 | Ms11t312500 | MsF3'5'H8 | 0.012827441 | 0.074238302 | 0.172787374 | 16.49740053 |
| Mbe02_t005590 | MbeF3'5'H3 | Ms02t024400 | MsF3'5'H5 | 0.084531185 | 0.363179019 | 0.232753493 | 80.70644865 |
| Mbe02_t005610 | MbeF3'5'H4 | Ms02t024420 | MsF3'5'H7 | 0.233654738 | 0.56302065 | 0.415002074 | 125.1157001 |
| Mbe02_t006410 | MbeF3'5'H5 | Ms02t024400 | MsF3'5'H5 | 0.038613944 | 0.134931273 | 0.286174905 | 29.98472725 |
| Mbe08_t039500 | MbeF3'5'H7 | Ms09t255750 | MsF3'5'H2 | 0.021869864 | 0.043882854 | 0.498369234 | 9.751745292 |
| Mbe06_t019910 | MbeF3'H | Ms03t075990 | MsF3'H | 0.039275796 | 0.088940851 | 0.441594556 | 19.76463367 |
| Mbe08_t015900 | MbeFLS1 | Ms08t229420 | MsFLS1 | 0.015824763 | 0.044260629 | 0.357535882 | 9.83569524 |
| Mbe02_t047300 | MbeFLS2 | Ms10t292840 | MsFLS3 | 0.07091356 | 0.208223802 | 0.340564139 | 46.2719561 |
| Mbe03_t008100 | MbeFLS4 | Ms03t052640 | MsFLS2 | 0.017194625 | 0.081068717 | 0.212099386 | 18.01527038 |
| Mbe05_t014700 | MbeLAR | Ms05t133790 | MsLAR | 0.035800456 | 0.113254989 | 0.316104893 | 25.16777539 |
| Mbe03_t031300 | MbeLDOX | Ms05t120140 | MsLDOX | 0.012094598 | 0.033177489 | 0.364542288 | 7.372775439 |
| Mbe05_t009880 | MbePAL1 | Ms05t135820 | MsPAL7 | 0.008147996 | 0.05694589 | 0.14308313 | 12.65464217 |
| Mbe05_t009880 | MbePAL1 | Ms11t319150 | MsPAL6 | 0.034821266 | 0.349267872 | 0.099697878 | 77.61508264 |
| Mbe06_t040740 | MbePAL2 | Ms05t135820 | MsPAL7 | 0.039533435 | 0.354459876 | 0.111531482 | 78.76886136 |

Musaceae flavonoid biosynthetic genes

| | | | | | | | |
|---|---|---|---|---|---|---|---|
| Mbe06_t040740 | MbePAL2 | Ms11t319150 | MsPAL6 | 0.007498891 | 0.058767201 | 0.127603335 | 13.0593779 |
| Mbe07_t015550 | MbePAL3 | Ms02t024050 | MsPAL4 | 0.061168463 | 0.331768271 | 0.184371046 | 73.7262824 |
| Mbe07_t015550 | MbePAL3 | Ms08t220210 | MsPAL5 | 0.01127311 | 0.077160124 | 0.14610021 | 17.14669423 |
| Mbe06_t048840 | MbePAL5 | Ms02t024050 | MsPAL4 | 0.067020691 | 0.391149273 | 0.171343002 | 86.92206069 |
| Mbe06_t048840 | MbePAL5 | Ms11t313180 | MsPAL3 | 0.017656183 | 0.054529823 | 0.323789474 | 12.11773837 |
| Mbe01_t022110 | MbePAL7 | Ms01t004140 | MsPAL1 | 0.012830187 | 0.064673788 | 0.198383102 | 14.3719529 |
| Mbe08_t037020 | MbePAL8 | Ms09t254450 | MsPAL8 | 0.004122507 | 0.045331016 | 0.090942314 | 10.07355903 |
| Mbe06_t034420 | MbeTT8 | Ms03t064490 | MsTT8.2 | 0.019450037 | 0.092697813 | 0.209821958 | 20.5995139 |
| Mbe03_t038300 | MbeTTG1 | Ms05t125970 | MsTTG1.2 | 0.006282759 | 0.070965572 | 0.088532497 | 15.77012706 |
| Mbe04_t041380 | MbeTTG1 | Ms04t112120 | MsTTG1 | 0.014488432 | 0.048913765 | 0.296203577 | 10.86972553 |
| Mbe02_t051150 | MbeUGT75C1 | Ms10t295740 | MsUGT75C1.4 | 0.026007903 | 0.049665257 | 0.52366392 | 11.03672382 |
| Mbe01_t034870 | MbeUGT75C1.3 | Ms06t152730 | MsUGT75C1.3 | 0.132854542 | 0.349272971 | 0.380374529 | 77.61621581 |
| Mbe02_t045890 | MbeUGT78D2 | Ms10t291590 | MsUGT78D2 | 0.041611839 | 0.124202882 | 0.335031188 | 27.60064037 |
| Mbe07_t034770 | MbeUGT78D2 | Ms07t193970 | MsUGT78D2.2 | 0.02208995 | 0.074763043 | 0.295466172 | 16.61400961 |
| Mbe03_t010580 | MbeUGT78D2.3 | Ms03t050760 | MsUGT78D2.3 | 0.020070185 | 0.072460248 | 0.276982003 | 16.10227742 |
| Mbe08_t027530 | MbeUGT79B1 | Ms09t246270 | MsUGT79B1 | 0.024228869 | 0.074521414 | 0.325126269 | 16.56031433 |

**Table S3k.** Ka/Ks values of orthologous flavonoid biosynthetic genes pairs between *M. lasiocarpa* and *E. glaucum*

| **Seq_1** | **Name1** | **Seq_2** | **Name2** | **Ka** | **Ks** | **Ka/Ks** | **Divergence (Mya)** |
|---|---|---|---|---|---|---|---|
| ML1h07G00167 | Ml4CL1 | Eg02_t001820 | Eg4CL4 | 0.010912336 | 0.062974035 | 0.173283096 | 13.99423002 |
| ML1h07G00248 | Ml4CL10 | Eg02_t002700 | Eg4CL1 | 0.028479506 | 0.062549797 | 0.455309333 | 13.8999548 |
| ML1h04G02613 | Ml4CL11 | Eg09_t018540 | Eg4CL2.2 | 0.007076915 | 0.048452768 | 0.146058014 | 10.76728184 |
| ML1h07G03382 | Ml4CL12 | Eg02_t037340 | Eg4CL2 | 0.021146424 | 0.061016831 | 0.346567061 | 13.55929576 |
| ML1h02G02377 | Ml4CL13 | Eg01_t025930 | Eg4CL11 | 0.009006044 | 0.049779826 | 0.180917545 | 11.06218358 |
| ML1h02G02377 | Ml4CL13 | Eg01_t014520 | Eg4CL10 | 0.053560305 | 0.354882672 | 0.150923978 | 78.86281603 |

| | | | | | | | |
|---|---|---|---|---|---|---|---|
| ML1h09G01309 | Ml4CL14 | Eg01_t014520 | Eg4CL10 | 0.017696294 | 0.061540782 | 0.287553938 | 13.67572934 |
| ML1h09G01309 | Ml4CL14 | Eg01_t025930 | Eg4CL11 | 0.050453313 | 0.369960986 | 0.136374687 | 82.21355235 |
| ML1h08G00557 | Ml4CL15 | Eg07_t014740 | Eg4CL15 | 0.010733636 | 0.058429874 | 0.183701164 | 12.98441635 |
| ML1h06G03426 | Ml4CL16 | Eg04_t025780 | Eg4CL12 | 0.009074147 | 0.053130744 | 0.170789 | 11.80683192 |
| ML1h06G03426 | Ml4CL16 | Eg04_t032900 | Eg4CL13 | 0.04663177 | 0.344725008 | 0.135272375 | 76.60555743 |
| ML1h06G02761 | Ml4CL17 | Eg04_t032900 | Eg4CL13 | 0.018480001 | 0.044288728 | 0.41726196 | 9.841939445 |
| ML1h06G02761 | Ml4CL17 | Eg04_t025780 | Eg4CL12 | 0.054798151 | 0.348272923 | 0.157342553 | 77.39398278 |
| ML1h09G02144 | Ml4CL18 | Eg04_t006840 | Eg4CL9 | 0.013819657 | 0.046109268 | 0.299715378 | 10.246504 |
| ML1h08G00140 | Ml4CL2 | Eg07_t001640 | Eg4CL14 | 0.024419318 | 0.079353915 | 0.307726693 | 17.63420344 |
| ML1h04G04085 | Ml4CL3 | Eg09_t002410 | Eg4CL3 | 0.02720826 | 0.055816111 | 0.487462478 | 12.40358021 |
| ML1h03G00142 | Ml4CL4 | Eg03_t041750 | Eg4CL8 | 0.019625693 | 0.065769815 | 0.298399702 | 14.61551446 |
| ML1h02G00664 | Ml4CL5 | Eg01_t038400 | Eg4CL7 | 0.011481124 | 0.059614287 | 0.192590136 | 13.24761924 |
| ML1h01G00405 | Ml4CL8 | Eg05_t033420 | Eg4CL5 | 0.018452147 | 0.068497612 | 0.269383796 | 15.22169163 |
| ML1h01G00972 | Ml4CL9 | Eg05_t023200 | Eg4CL1.2 | 0.011529978 | 0.052466756 | 0.219757784 | 11.65927905 |
| ML1h06G01342 | MlANR2 | Eg06_t013840 | EgANR1 | 0.037270463 | 0.079574011 | 0.468374818 | 17.68311346 |
| ML1h07G01339 | MlANR4 | Eg02_t015210 | EgANR2 | 0.019034056 | 0.038289039 | 0.497115004 | 8.508675389 |
| ML1h08G01785 | MlC1 | Eg07_t019620 | EgC1 | 0.029138198 | 0.067778924 | 0.429900564 | 15.06198303 |
| ML1h05G03234 | MlC1 | Eg08_t035970 | EgC1.2 | 0.019109314 | 0.086896354 | 0.219909272 | 19.31030091 |
| ML1h05G01754 | MlC4H1 | Eg08_t019660 | EgC4H1 | 0.004370323 | 0.045608921 | 0.095821679 | 10.13531575 |
| ML1h05G01754 | MlC4H1 | Eg08_t048930 | EgC4H3 | 0.036355405 | 0.353455818 | 0.102857 | 78.54573723 |
| ML1h08G00252 | MlC4H2 | Eg07_t003130 | EgC4H2 | 0.008209264 | 0.057955651 | 0.141647344 | 12.87903362 |
| ML1h05G04363 | MlC4H3 | Eg08_t048930 | EgC4H3 | 0.008780374 | 0.060132358 | 0.146017455 | 13.36274633 |
| ML1h05G04363 | MlC4H3 | Eg08_t019660 | EgC4H1 | 0.033425509 | 0.295940509 | 0.112946718 | 65.76455764 |
| ML1h08G03392 | MlC4H4 | Eg07_t034110 | EgC4H4 | 0.013124651 | 0.061422859 | 0.213676987 | 13.64952412 |
| ML1h09G01419 | MlCHI1 | Eg01_t015550 | EgCHI1 | 0.020913903 | 0.053382209 | 0.391776644 | 11.86271308 |

Musaceae flavonoid biosynthetic genes

| | | | | | | | |
|---|---|---|---|---|---|---|---|
| ML1h04G02595 | MlCHI2 | Eg09_t018750 | EgCHI2 | 0.010254817 | 0.043464182 | 0.235937192 | 9.658707121 |
| ML1h05G01818 | MlCHS1 | Eg08_t049400 | EgCHS5 | 0.106062297 | 0.401538611 | 0.264139722 | 89.23080248 |
| ML1h05G01818 | MlCHS1 | Eg08_t020390 | EgCHS3 | 0.011308167 | 0.058352626 | 0.193790208 | 12.96725019 |
| ML1h05G01818 | MlCHS1 | Eg08_t040870 | EgCHS2 | 0.033290974 | 0.323508417 | 0.102906053 | 71.89075935 |
| ML1h05G03634 | MlCHS2 | Eg08_t040870 | EgCHS2 | 0.011310299 | 0.035913503 | 0.314931657 | 7.980778545 |
| ML1h05G03634 | MlCHS2 | Eg08_t049400 | EgCHS5 | 0.102801707 | 0.358470458 | 0.286778742 | 79.66010169 |
| ML1h05G03634 | MlCHS2 | Eg08_t020390 | EgCHS3 | 0.033265512 | 0.313707975 | 0.106039739 | 69.71288336 |
| ML1h05G04408 | MlCHS3 | Eg08_t049400 | EgCHS5 | 0.030391162 | 0.102804521 | 0.295620873 | 22.84544902 |
| ML1h05G04408 | MlCHS3 | Eg08_t020390 | EgCHS3 | 0.128648066 | 0.366730391 | 0.350797395 | 81.49564242 |
| ML1h05G04408 | MlCHS3 | Eg08_t040870 | EgCHS2 | 0.120098339 | 0.324407833 | 0.370207891 | 72.09062946 |
| ML1h04G00849 | MlDFR1 | Eg09_t036940 | EgDFR1 | 0.010144929 | 0.06288755 | 0.161318562 | 13.97501103 |
| ML1h06G03238 | MlDFR2 | Eg04_t027830 | EgDFR2 | 0.015187317 | 0.04458334 | 0.340650043 | 9.907408801 |
| ML1h07G00690 | MlF3'5'H1 | Eg02_t007770 | EgF3'5'H2 | 0.01291102 | 0.045162796 | 0.285877339 | 10.03617679 |
| ML1h04G03337 | MlF3'5'H12 | Eg09_t010710 | EgF3'5'H10 | 0.022583935 | 0.04069856 | 0.554907474 | 9.044124427 |
| ML1h04G02425 | MlF3'5'H2 | Eg09_t020640 | EgF3'5'H1 | 0.024096666 | 0.068966544 | 0.34939646 | 15.32589862 |
| ML1h07G00951 | MlF3'5'H3 | Eg02_t010650 | EgF3'5'H5 | 0.008861423 | 0.032277505 | 0.274538678 | 7.172778783 |
| ML1h08G03854 | MlF3'5'H4 | Eg07_t042500 | EgF3'5'H4 | 0.017689499 | 0.069429744 | 0.254782723 | 15.42883202 |
| ML1h06G02237 | MlF3'5'H6 | Eg06_t004020 | EgF3'5'H3 | 0.022611436 | 0.099521676 | 0.22720112 | 22.11592797 |
| ML1h07G01736 | MlF3'5'H7 | Eg02_t019070 | EgF3'5'H6 | 0.023118078 | 0.097209379 | 0.237817364 | 21.60208421 |
| ML1h07G01736 | MlF3'5'H7 | Eg02_t019180 | EgF3'5'H9 | 0.095935569 | 0.430326585 | 0.222936653 | 95.62813004 |
| ML1h07G02004 | MlF3H1 | Eg02_t022060 | EgF3H2 | 0.010571827 | 0.063783435 | 0.165745656 | 14.17409675 |
| ML1h08G02519 | MlF3H2 | Eg07_t027880 | EgF3H1 | 0.011200837 | 0.103192792 | 0.108542824 | 22.93173148 |
| ML1h03G02252 | MlF3H3 | Eg03_t018110 | EgF3'H1 | 0.013015511 | 0.046707697 | 0.278658804 | 10.37948825 |
| ML1h03G01030 | MlFLS1 | Eg03_t031640 | EgFLS4 | 0.015877109 | 0.057423179 | 0.276493034 | 12.76070637 |
| ML1h06G02043 | MlFLS4 | Eg01_t007540 | EgFLS2 | 0.085980299 | 0.354362068 | 0.242634037 | 78.74712612 |

Musaceae flavonoid biosynthetic genes

| | | | | | | | |
|---|---|---|---|---|---|---|---|
| ML1h06G02043 | MlFLS4 | Eg06_t006150 | EgFLS1 | 0.072917831 | 0.116560578 | 0.625578836 | 25.90235065 |
| ML1h09G00678 | MlFLS5 | Eg01_t007540 | EgFLS2 | 0.00854055 | 0.046355318 | 0.184240998 | 10.3011817 |
| ML1h01G01259 | MlLAR | Eg05_t019870 | EgLAR | 0.019209246 | 0.054129765 | 0.35487399 | 12.02883656 |
| ML1h01G01735 | MlLDOX | Eg05_t014690 | EgLDOX | 0.019529175 | 0.045343477 | 0.430694255 | 10.07632817 |
| ML1h04G02672 | MlPAL1 | Eg05_t024420 | EgPAL8 | 0.036274381 | 0.368810564 | 0.098355049 | 81.95790315 |
| ML1h04G02672 | MlPAL1 | Eg09_t017980 | EgPAL7 | 0.00437046 | 0.052482546 | 0.08327454 | 11.6627879 |
| ML1h01G00856 | MlPAL2 | Eg05_t024420 | EgPAL8 | 0.006260797 | 0.032839625 | 0.190647639 | 7.297694506 |
| ML1h01G00856 | MlPAL2 | Eg09_t017980 | EgPAL7 | 0.03735832 | 0.395393758 | 0.094483838 | 87.86527959 |
| ML1h07G00086 | MlPAL3 | Eg02_t001010 | EgPAL6 | 0.010654033 | 0.051744654 | 0.205896312 | 11.49881191 |
| ML1h07G00086 | MlPAL3 | Eg02_t018790 | EgPAL4 | 0.060040726 | 0.452258101 | 0.132757657 | 100.5018002 |
| ML1h07G00086 | MlPAL3 | Eg09_t011330 | EgPAL5 | 0.08367939 | 0.43435677 | 0.192651285 | 96.52372669 |
| ML1h04G03282 | MlPAL4 | Eg09_t011330 | EgPAL5 | 0.016562941 | 0.057739944 | 0.286854119 | 12.83109873 |
| ML1h07G01701 | MlPAL5 | Eg02_t018790 | EgPAL4 | 0.015103557 | 0.052971838 | 0.285124281 | 11.77151959 |
| ML1h07G01701 | MlPAL5 | Eg02_t001010 | EgPAL6 | 0.067895942 | 0.405963074 | 0.167246595 | 90.2140164 |
| ML1h07G01701 | MlPAL5 | Eg09_t011330 | EgPAL5 | 0.067114369 | 0.420507428 | 0.159603289 | 93.44609515 |
| ML1h02G00424 | MlPAL6 | Eg01_t040920 | EgPAL1 | 0.011894453 | 0.057274075 | 0.207676047 | 12.72757223 |
| ML1h01G01719 | MlPAL8 | Eg05_t014880 | EgPAL2 | 0.009368291 | 0.048032416 | 0.195041009 | 10.67387018 |
| ML1h06G02093 | MlPAL9 | Eg06_t005550 | EgPAL9 | 0.002943056 | 0.045296783 | 0.064972732 | 10.06595171 |
| ML1h02G03012 | MlTT8 | Eg04_t020700 | EgTT8 | 0.018037727 | 0.078340838 | 0.230246802 | 17.40907505 |
| ML1h03G03510 | MlTT8 | Eg03_t004200 | EgTT8.2 | 0.023274927 | 0.09188372 | 0.253308497 | 20.41860437 |
| ML1h01G02333 | MlTTG1 | Eg05_t009200 | EgTTG1 | 0.004714217 | 0.074272141 | 0.063472211 | 16.50492028 |
| ML1h04G00438 | MlTTG1 | Eg09_t041330 | EgTTG1.2 | 0.009913988 | 0.079986938 | 0.123945091 | 17.77487519 |
| ML1h05G03318 | MlUGT75C1 | Eg08_t037040 | EgUGT75C1.3 | 0.038625976 | 0.068466729 | 0.564156881 | 15.21482861 |
| ML1h05G03317 | MlUGT75C1.3 | Eg08_t037030 | EgUGT75C1.2 | 0.036043461 | 0.075969715 | 0.474445129 | 16.88215887 |
| ML1h09G00476 | MlUGT78D2 | Eg01_t005290 | EgUGT78D2.3 | 0.021946703 | 0.063775726 | 0.344123137 | 14.17238367 |

Musaceae flavonoid biosynthetic genes

| | | | | | | | |
|---|---|---|---|---|---|---|---|
| ML1h06G01843 | MlUGT78D2 | Eg06_t008220 | EgUGT78D2.2 | 0.008881683 | 0.049532559 | 0.179309991 | 11.00723539 |
| ML1h05G00763 | MlUGT78D2.3 | Eg08_t008340 | EgUGT78D2 | 0.028141648 | 0.068516976 | 0.410725194 | 15.22599461 |
| ML1h08G03185 | MlUGT79B1 | Eg07_t037900 | EgUGT79B1 | 0.018758493 | 0.048063033 | 0.390289415 | 10.68067389 |

**Table S3l.** Divergence time, Ka, Ks and Ka/Ks values of orthologous flavonoid biosynthetic genes pairs between *M. lasiocarpa* and *M. acuminata*

| **Seq_1** | **Name1** | **Seq_2** | **Name2** | **Ka** | **Ks** | **Ka_Ks** | **Diversity_year** |
|---|---|---|---|---|---|---|---|
| ML1h07G00167 | Ml4CL1 | Parent=Macma4_08_g13960 | Ma4CL4 | 0.008386418 | 0.082516897 | 0.101632732 | 18.33708813 |
| ML1h07G00248 | Ml4CL10 | Parent=Macma4_08_g13040 | Ma4CL1 | 0.029365358 | 0.079211822 | 0.37071939 | 17.60262712 |
| ML1h04G02613 | Ml4CL11 | Parent=Macma4_11_g22750 | Ma4CL2.2 | 0.010239067 | 0.066273671 | 0.154496752 | 14.72748243 |
| ML1h02G02377 | Ml4CL13 | Parent=Macma4_01_g19800 | Ma4CL10 | 0.008192326 | 0.06777183 | 0.120880993 | 15.06040668 |
| ML1h02G02377 | Ml4CL13 | Parent=Macma4_03_g13930 | Ma4CL11 | 0.041444695 | 0.338817297 | 0.122321662 | 75.29273258 |
| ML1h02G02377 | Ml4CL13 | Parent=Macma4_07_g05220 | Ma4CL14 | 0.078577548 | 0.415155625 | 0.189272513 | 92.25680564 |
| ML1h09G01309 | Ml4CL14 | Parent=Macma4_01_g19800 | Ma4CL10 | 0.043578549 | 0.34511291 | 0.126273307 | 76.69175779 |
| ML1h09G01309 | Ml4CL14 | Parent=Macma4_03_g13930 | Ma4CL11 | 0.018958355 | 0.075934134 | 0.249668421 | 16.87425206 |
| ML1h09G01309 | Ml4CL14 | Parent=Macma4_07_g05220 | Ma4CL14 | 0.07836337 | 0.460836246 | 0.170046021 | 102.4080547 |
| ML1h08G00557 | Ml4CL15 | Parent=Macma4_09_g24760 | Ma4CL16 | 0.019915428 | 0.055981891 | 0.355747686 | 12.44042022 |
| ML1h06G03426 | Ml4CL16 | Parent=Macma4_04_g13130 | Ma4CL13 | 0.010720358 | 0.091178917 | 0.117574971 | 20.26198145 |
| ML1h06G03426 | Ml4CL16 | Parent=Macma4_04_g06110 | Ma4CL12 | 0.040096755 | 0.37818499 | 0.106024184 | 84.04110878 |
| ML1h06G02761 | Ml4CL17 | Parent=Macma4_04_g06110 | Ma4CL12 | 0.024268207 | 0.108834688 | 0.222982286 | 24.18548613 |
| ML1h06G02761 | Ml4CL17 | Parent=Macma4_04_g13130 | Ma4CL13 | 0.056483251 | 0.338582284 | 0.16682282 | 75.24050748 |
| ML1h09G02144 | Ml4CL18 | Parent=Macma4_06_g39580 | Ma4CL9 | 0.023578393 | 0.053338465 | 0.442052337 | 11.85299212 |
| ML1h08G00140 | Ml4CL2 | Parent=Macma4_07_g18890 | Ma4CL15 | 0.025643524 | 0.103207052 | 0.248466776 | 22.93490053 |
| ML1h04G04085 | Ml4CL3 | Parent=Macma4_11_g06130 | Ma4CL6 | 0.040373156 | 0.105896704 | 0.381250354 | 23.53260087 |
| ML1h03G00142 | Ml4CL4 | Parent=Macma4_03_g14310 | Ma4CL8 | 0.027443393 | 0.08570551 | 0.320205703 | 19.04566898 |

| | | | | | | | |
|---|---|---|---|---|---|---|---|
| ML1h02G00664 | Ml4CL5 | Parent=Macma4_01_g07350 | Ma4CL7 | 0.021879777 | 0.112347627 | 0.194750678 | 24.96613939 |
| ML1h01G00405 | Ml4CL8 | Parent=Macma4_05_g28920 | Ma4CL5 | 0.018436683 | 0.099018175 | 0.186194934 | 22.00403889 |
| ML1h01G00972 | Ml4CL9 | Parent=Macma4_05_g22890 | Ma4CL1.2 | 0.025044063 | 0.082203667 | 0.304658709 | 18.26748154 |
| ML1h06G01342 | MlANR2 | Parent=Macma4_07_g10660 | MaANR2 | 0.043325898 | 0.104110832 | 0.416151686 | 23.1357404 |
| ML1h07G01339 | MlANR4 | Parent=Macma4_08_g01440 | MaANR5 | 0.027053169 | 0.072735285 | 0.371940098 | 16.16339666 |
| ML1h08G01785 | MlC1 | Parent=Macma4_09_g29590 | MaC1.2 | 0.03876832 | 0.07100853 | 0.545967082 | 15.77967333 |
| ML1h05G01322 | MlC1.3 | Parent=Macma4_10_g21630 | MaC1 | 0.035234662 | 0.12844674 | 0.274313397 | 28.54372005 |
| ML1h05G01754 | MlC4H1 | Parent=Macma4_06_g10450 | MaC4H3 | 0.034332467 | 0.312700142 | 0.109793577 | 69.48892037 |
| ML1h05G01754 | MlC4H1 | Parent=Macma4_10_g16840 | MaC4H1 | 0.007013494 | 0.072240593 | 0.097085215 | 16.05346522 |
| ML1h08G00252 | MlC4H2 | Parent=Macma4_07_g20380 | MaC4H2 | 0.009129059 | 0.08042723 | 0.113507069 | 17.87271771 |
| ML1h05G04363 | MlC4H3 | Parent=Macma4_06_g10450 | MaC4H3 | 0.008770107 | 0.063362198 | 0.138412283 | 14.08048844 |
| ML1h05G04363 | MlC4H3 | Parent=Macma4_10_g16840 | MaC4H1 | 0.036210847 | 0.319391298 | 0.113374556 | 70.97584397 |
| ML1h08G03392 | MlC4H4 | Parent=Macma4_09_g04800 | MaC4H4 | 0.014876882 | 0.089681225 | 0.165886254 | 19.9291611 |
| ML1h09G01419 | MlCHI1 | Parent=Macma4_04_g18950 | MaCHI1 | 0.04247556 | 0.071803674 | 0.591551356 | 15.95637191 |
| ML1h04G02595 | MlCHI2 | Parent=Macma4_11_g22930 | MaCHI2 | 0.010258324 | 0.105502785 | 0.097232729 | 23.44506344 |
| ML1h05G01818 | MlCHS1 | Parent=Macma4_06_g18170 | MaCHS2 | 0.041169457 | 0.319572413 | 0.128826694 | 71.0160917 |
| ML1h05G01818 | MlCHS1 | Parent=Macma4_10_g16140 | MaCHS6 | 0.013587841 | 0.077607636 | 0.175083815 | 17.2461413 |
| ML1h05G03634 | MlCHS2 | Parent=Macma4_06_g18170 | MaCHS2 | 0.017609443 | 0.063965686 | 0.275295145 | 14.21459693 |
| ML1h05G03634 | MlCHS2 | Parent=Macma4_06_g10010 | MaCHS5 | 0.111157805 | 0.373548322 | 0.29757276 | 83.01073828 |
| ML1h05G03634 | MlCHS2 | Parent=Macma4_10_g16140 | MaCHS6 | 0.030917458 | 0.335608062 | 0.092123706 | 74.57956924 |
| ML1h05G04408 | MlCHS3 | Parent=Macma4_06_g10010 | MaCHS5 | 0.043947439 | 0.146382931 | 0.300222427 | 32.52954024 |
| ML1h05G04408 | MlCHS3 | Parent=Macma4_06_g18170 | MaCHS2 | 0.126755524 | 0.329464642 | 0.384731798 | 73.21436497 |
| ML1h05G04408 | MlCHS3 | Parent=Macma4_10_g16140 | MaCHS6 | 0.127290951 | 0.361301944 | 0.352311834 | 80.28932087 |
| ML1h04G00849 | MlDFR1 | Parent=Macma4_04_g33220 | MaDFR1 | 0.015251532 | 0.096834218 | 0.15750147 | 21.51871514 |
| ML1h06G03238 | MlDFR2 | Parent=Macma4_04_g11180 | MaDFR3 | 0.017016835 | 0.082963447 | 0.205112433 | 18.43632156 |

Musaceae flavonoid biosynthetic genes

| | | | | | | | |
|---|---|---|---|---|---|---|---|
| ML1h03G02164 | MlDFR3 | Parent=Macma4_03_g33910 | MaDFR2 | 0.016017036 | 0.071987515 | 0.222497411 | 15.99722564 |
| ML1h07G00690 | MlF3'5'H1 | Parent=Macma4_08_g08290 | MaF3'5'H1 | 0.016920349 | 0.058677754 | 0.28836055 | 13.03950085 |
| ML1h04G03337 | MlF3'5'H12 | Parent=Macma4_11_g15260 | MaF3'5'H13 | 0.025649849 | 0.079629213 | 0.322116066 | 17.69538059 |
| ML1h04G02425 | MlF3'5'H2 | Parent=Macma4_11_g05110 | MaF3'5'H2 | 0.032383984 | 0.090292082 | 0.358658074 | 20.06490704 |
| ML1h07G00951 | MlF3'5'H3 | Parent=Macma4_08_g05480 | MaF3'5'H3 | 0.021407727 | 0.054894178 | 0.389981735 | 12.19870612 |
| ML1h08G03854 | MlF3'5'H4 | Parent=Macma4_10_g01260 | MaF3'5'H6 | 0.016837947 | 0.104599596 | 0.16097526 | 23.24435469 |
| ML1h06G02237 | MlF3'5'H6 | Parent=Macma4_09_g17660 | MaF3'5'H4 | 0.029881321 | 0.057330485 | 0.521211724 | 12.74010784 |
| ML1h07G01736 | MlF3'5'H7 | Parent=Macma4_02_g03290 | MaF3'5'H11 | 0.037235725 | 0.139733036 | 0.266477603 | 31.05178586 |
| ML1h07G01736 | MlF3'5'H7 | Parent=Macma4_02_g03040 | MaF3'5'H7 | 0.073799377 | 0.145464074 | 0.507337478 | 32.32534988 |
| ML1h07G02004 | MlF3H1 | Parent=Macma4_02_g06130 | MaF3H2 | 0.017761067 | 0.08069583 | 0.220098941 | 17.93240672 |
| ML1h08G02519 | MlF3H2 | Parent=Macma4_07_g17110 | MaF3H1 | 0.01657526 | 0.083752264 | 0.197908206 | 18.61161414 |
| ML1h03G02252 | MlF3H3 | Parent=Macma4_03_g33170 | MaF3'H | 0.020962063 | 0.079229271 | 0.264574731 | 17.60650473 |
| ML1h03G01030 | MlFLS1 | Parent=Macma4_08_g24690 | MaFLS2 | 0.01988188 | 0.095366382 | 0.208478917 | 21.19252925 |
| ML1h06G02043 | MlFLS4 | Parent=Macma4_03_g07220 | MaFLS1 | 0.084518182 | 0.354203423 | 0.238614809 | 78.71187174 |
| ML1h09G00678 | MlFLS5 | Parent=Macma4_03_g07220 | MaFLS1 | 0.023221669 | 0.064772372 | 0.358511947 | 14.39386047 |
| ML1h01G01259 | MlLAR | Parent=Macma4_05_g19560 | MaLAR | 0.02062207 | 0.087487352 | 0.235714876 | 19.44163375 |
| ML1h01G01735 | MlLDOX | Parent=Macma4_05_g03920 | MaLDOX | 0.020728814 | 0.054227921 | 0.382253524 | 12.05064907 |
| ML1h04G02672 | MlPAL1 | Parent=Macma4_05_g21700 | MaPAL5 | 0.043754161 | 0.339985024 | 0.128694378 | 75.55222761 |
| ML1h04G02672 | MlPAL1 | Parent=Macma4_11_g22170 | MaPAL6 | 0.010639031 | 0.054809967 | 0.1941076 | 12.1799926 |
| ML1h01G00856 | MlPAL2 | Parent=Macma4_05_g21700 | MaPAL5 | 0.016351135 | 0.045096889 | 0.362577896 | 10.0215308 |
| ML1h07G00086 | MlPAL3 | Parent=Macma4_02_g02760 | MaPAL4 | 0.061181789 | 0.413009578 | 0.148136489 | 91.77990625 |
| ML1h07G00086 | MlPAL3 | Parent=Macma4_08_g14800 | MaPAL7 | 0.00938392 | 0.074986467 | 0.12514151 | 16.66365939 |
| ML1h04G03282 | MlPAL4 | Parent=Macma4_02_g02760 | MaPAL4 | 0.073162606 | 0.389712726 | 0.187734712 | 86.6028279 |
| ML1h04G03282 | MlPAL4 | Parent=Macma4_11_g15930 | MaPAL3 | 0.026905808 | 0.074487676 | 0.361211537 | 16.55281681 |
| ML1h07G01701 | MlPAL5 | Parent=Macma4_02_g02760 | MaPAL4 | 0.020881918 | 0.09426511 | 0.221523302 | 20.94780214 |

Musaceae flavonoid biosynthetic genes

| | | | | | | | |
|---|---|---|---|---|---|---|---|
| ML1h07G01701 | MlPAL5 | Parent=Macma4_08_g14800 | MaPAL7 | 0.059995105 | 0.389509319 | 0.154027392 | 86.55762639 |
| ML1h02G00424 | MlPAL6 | Parent=Macma4_01_g04800 | MaPAL1 | 0.014724784 | 0.068864831 | 0.213821533 | 15.30329581 |
| ML1h01G01719 | MlPAL8 | Parent=Macma4_05_g03790 | MaPAL2 | 0.008736448 | 0.092870115 | 0.094071685 | 20.63780344 |
| ML1h06G02093 | MlPAL9 | Parent=Macma4_09_g16190 | MaPAL8 | 0.008863605 | 0.056596185 | 0.156611356 | 12.57692995 |
| ML1h02G03012 | MlTT8 | Parent=Macma4_06_g25800 | MaTT8 | 0.015649846 | 0.053509215 | 0.292470115 | 11.89093677 |
| ML1h03G03510 | MlTT8 | Parent=Macma4_03_g19880 | MaTT8.2 | 0.029538475 | 0.137283085 | 0.215164708 | 30.50735232 |
| ML1h01G02333 | MlTTG1 | Parent=Macma4_05_g10420 | MaTTG1 | 0.015397304 | 0.043968633 | 0.350188364 | 9.770807225 |
| ML1h04G00438 | MlTTG1 | Parent=Macma4_04_g37870 | MaTTG1.2 | 0.012807142 | 0.08982685 | 0.142575872 | 19.96152227 |
| ML1h05G03318 | MlUGT75C1 | Parent=Macma4_06_g07210 | MaUGT75C1.2 | 0.056531459 | 0.103420116 | 0.54661956 | 22.98224806 |
| ML1h05G00327 | MlUGT75C1 | Parent=Macma4_10_g31950 | MaUGT75C1 | 0.029492504 | 0.057873964 | 0.509598817 | 12.86088095 |
| ML1h05G03317 | MlUGT75C1.3 | Parent=Macma4_06_g07200 | MaUGT75C1.3 | 0.04952077 | 0.103683777 | 0.477613481 | 23.04083943 |
| ML1h09G00476 | MlUGT78D2 | Parent=Macma4_03_g04880 | MaUGT78D2.3 | 0.026516018 | 0.080604581 | 0.328964158 | 17.91212921 |
| ML1h06G01843 | MlUGT78D2 | Parent=Macma4_07_g16200 | MaUGT78D2.2 | 0.020902074 | 0.077009636 | 0.271421546 | 17.11325234 |
| ML1h05G00763 | MlUGT78D2.3 | Parent=Macma4_10_g27580 | MaUGT78D2 | 0.03869547 | 0.11667959 | 0.331638721 | 25.92879775 |
| ML1h08G03185 | MlUGT79B1 | Parent=Macma4_09_g07100 | MaUGT79B1 | 0.027207258 | 0.083815005 | 0.32461083 | 18.6255566 |

**Table S3m.** Divergence time, Ka, Ks and Ka/Ks values of orthologous flavonoid biosynthetic genes pairs between *M. lasiocarpa* and *M. balbisiana*

| **Seq_1** | **Name1** | **Seq_2** | **Name2** | **Ka** | **Ks** | **Ka_Ks** | **Diversity_year** |
|---|---|---|---|---|---|---|---|
| ML1h07G00167 | Ml4CL1 | Mba08_g13620 | Mb4CL4 | 0.01553557 | 0.100872186 | 0.154012422 | 22.41604138 |
| ML1h07G00248 | Ml4CL10 | Mba08_g12720 | Mb4CL1.2 | 0.02800495 | 0.092752812 | 0.301931014 | 20.61173596 |
| ML1h04G02613 | Ml4CL11 | Mba11_g20940 | Mb4CL2.2 | 0.014219371 | 0.056021163 | 0.253821423 | 12.44914734 |
| ML1h02G02377 | Ml4CL13 | Mba01_g28370 | Mb4CL10 | 0.00737157 | 0.07827151 | 0.094179476 | 17.39366889 |
| ML1h09G01309 | Ml4CL14 | Mba01_g28370 | Mb4CL10 | 0.043737653 | 0.355532462 | 0.12302014 | 79.00721369 |
| ML1h08G00557 | Ml4CL15 | Mba04_g12340 | Mb4CL12 | 0.052984387 | 0.485221546 | 0.109196278 | 107.8270102 |

| | | | | | | | |
|---|---|---|---|---|---|---|---|
| ML1h06G03426 | Ml4CL16 | Mba04_g12340 | Mb4CL12 | 0.012379923 | 0.105251203 | 0.117622631 | 23.38915632 |
| ML1h06G03426 | Ml4CL16 | Mba04_g05720 | Mb4CL11 | 0.043570186 | 0.366331946 | 0.118936353 | 81.40709917 |
| ML1h06G02761 | Ml4CL17 | Mba04_g05720 | Mb4CL11 | 0.028530919 | 0.09230584 | 0.309091157 | 20.51240887 |
| ML1h06G02761 | Ml4CL17 | Mba04_g12340 | Mb4CL12 | 0.058237564 | 0.358366037 | 0.162508603 | 79.63689714 |
| ML1h09G02144 | Ml4CL18 | Mba06_g36630 | Mb4CL9 | 0.02481616 | 0.052149585 | 0.475864952 | 11.58879662 |
| ML1h08G00140 | Ml4CL2 | Mba07_g17040 | Mb4CL14 | 0.026487338 | 0.120995616 | 0.218911549 | 26.88791474 |
| ML1h03G00142 | Ml4CL4 | Mba03_g13490 | Mb4CL8 | 0.047463818 | 0.109910569 | 0.431840345 | 24.42457085 |
| ML1h02G00664 | Ml4CL5 | Mba01_g16650 | Mb4CL7 | 0.011057196 | 0.081057297 | 0.136412104 | 18.01273272 |
| ML1h01G00405 | Ml4CL8 | Mba05_g27160 | Mb4CL5 | 0.022700346 | 0.098898578 | 0.229531569 | 21.97746173 |
| ML1h01G00972 | Ml4CL9 | Mba05_g21540 | Mb4CL1 | 0.026771751 | 0.081940287 | 0.326722693 | 18.20895275 |
| ML1h07G01339 | MlANR4 | Mba08_g01400 | MbANR | 0.025709458 | 0.06878132 | 0.373785472 | 15.28473775 |
| ML1h08G01785 | MlC1 | Mba09_g23680 | MbC1.3 | 0.173514535 | 0.216601891 | 0.801075812 | 48.13375346 |
| ML1h05G03234 | MlC1 | Mba06_g05630 | MbC1.2 | 0.029110243 | 0.100381601 | 0.289995806 | 22.3070225 |
| ML1h05G01322 | MlC1.3 | Mba10_g15520 | MbC1 | 0.035286476 | 0.140004964 | 0.252037318 | 31.11221419 |
| ML1h05G01754 | MlC4H1 | Mba06_g09560 | MbC4H3 | 0.031625826 | 0.332907993 | 0.094998697 | 73.97955411 |
| ML1h08G00252 | MlC4H2 | Mba07_g18380 | MbC4H4 | 0.006377202 | 0.083764649 | 0.07613238 | 18.61436655 |
| ML1h05G04363 | MlC4H3 | Mba06_g09560 | MbC4H3 | 0.00789481 | 0.075280111 | 0.10487245 | 16.72891363 |
| ML1h08G03392 | MlC4H4 | Mba09_g04380 | MbC4H1 | 0.022873996 | 0.086512485 | 0.264401101 | 19.22499668 |
| ML1h09G01419 | MlCHI1 | Mba04_g18040 | MbCHI2 | 0.066382369 | 0.119540262 | 0.555313901 | 26.56450256 |
| ML1h04G02595 | MlCHI2 | Mba11_g21110 | MbCHI1 | 0.012446267 | 0.106575428 | 0.116783642 | 23.68342846 |
| ML1h05G01818 | MlCHS1 | Mba06_g16830 | MbCHS2 | 0.033881359 | 0.331520521 | 0.102199885 | 73.67122693 |
| ML1h05G01818 | MlCHS1 | Mba10_g10820 | MbCHS8 | 0.011309233 | 0.093258753 | 0.121267258 | 20.72416727 |
| ML1h05G03634 | MlCHS2 | Mba06_g16830 | MbCHS2 | 0.010172521 | 0.077488211 | 0.131278306 | 17.21960234 |
| ML1h05G03634 | MlCHS2 | Mba10_g10820 | MbCHS8 | 0.028591226 | 0.335236646 | 0.085286698 | 74.49703249 |
| ML1h05G04408 | MlCHS3 | Mba06_g09120 | MbCHS7 | 0.038595901 | 0.098214943 | 0.39297382 | 21.82554278 |

| | | | | | | | |
|---|---|---|---|---|---|---|---|
| ML1h05G04408 | MlCHS3 | Mba06_g16830 | MbCHS2 | 0.119451815 | 0.337905479 | 0.353506593 | 75.09010649 |
| ML1h05G04408 | MlCHS3 | Mba10_g10820 | MbCHS8 | 0.123350065 | 0.366592145 | 0.336477655 | 81.46492119 |
| ML1h04G00849 | MlDFR1 | Mba04_g31800 | MbDFR1 | 0.013507794 | 0.093563779 | 0.144369909 | 20.79195083 |
| ML1h03G02164 | MlDFR3 | Mba01_g01310 | MbDFR2 | 0.01350973 | 0.054703524 | 0.246962696 | 12.15633861 |
| ML1h07G00690 | MlF3'5'H1 | Mba08_g07950 | MbF3'5'H1 | 0.013901929 | 0.065561783 | 0.212043179 | 14.56928508 |
| ML1h04G03337 | MlF3'5'H12 | Mba11_g13650 | MbF3'5'H13 | 0.022643229 | 0.096648269 | 0.234284892 | 21.47739308 |
| ML1h04G02425 | MlF3'5'H2 | Mba11_g04840 | MbF3'5'H2 | 0.030395221 | 0.12074783 | 0.25172478 | 26.83285111 |
| ML1h08G03854 | MlF3'5'H4 | Mba10_g00740 | MbF3'5'H4 | 0.018212544 | 0.088935235 | 0.204784346 | 19.76338545 |
| ML1h06G02237 | MlF3'5'H6 | Mba09_g16030 | MbF3'5'H3 | 0.029021478 | 0.062827412 | 0.461923813 | 13.96164718 |
| ML1h07G01736 | MlF3'5'H7 | Mba02_g01950 | MbF3'5'H12 | 0.092941402 | 0.445007983 | 0.208853336 | 98.89066294 |
| ML1h07G02004 | MlF3H1 | Mba02_g04670 | MbF3H1 | 0.015365454 | 0.063889622 | 0.240500002 | 14.19769374 |
| ML1h08G02519 | MlF3H2 | Mba07_g15320 | MbF3H2 | 0.017769834 | 0.10522276 | 0.168878237 | 23.38283552 |
| ML1h03G02252 | MlF3H3 | Mba01_g02030 | MbF3'H | 0.021845144 | 0.076482549 | 0.285622596 | 16.9961219 |
| ML1h03G01030 | MlFLS1 | Mba08_g23630 | MbFLS1 | 0.017525441 | 0.129463865 | 0.135369367 | 28.76974782 |
| ML1h06G02043 | MlFLS4 | Mba03_g06940 | MbFLS4 | 0.086070159 | 0.346259135 | 0.248571518 | 76.94647449 |
| ML1h06G02043 | MlFLS4 | Mba09_g14060 | MbFLS5 | 0.070790543 | 0.151310838 | 0.46784846 | 33.62463072 |
| ML1h09G00678 | MlFLS5 | Mba03_g06940 | MbFLS4 | 0.019877489 | 0.048475018 | 0.410056342 | 10.77222622 |
| ML1h09G00678 | MlFLS5 | Mba09_g14060 | MbFLS5 | 0.160871935 | 0.47849146 | 0.336206492 | 106.3314356 |
| ML1h01G01259 | MlLAR | Mba05_g18490 | MbLAR | 0.028887597 | 0.072151723 | 0.400372934 | 16.03371623 |
| ML1h01G01735 | MlLDOX | Mba05_g13780 | MbLDOX | 0.026947173 | 0.071633353 | 0.376181927 | 15.91852295 |
| ML1h04G02672 | MlPAL1 | Mba11_g20430 | MbPAL2 | 0.01189942 | 0.052799121 | 0.225371553 | 11.73313789 |
| ML1h07G00086 | MlPAL3 | Mba08_g14430 | MbPAL3 | 0.009384898 | 0.074961126 | 0.125196869 | 16.658028 |
| ML1h04G03282 | MlPAL4 | Mba11_g14220 | MbPAL6 | 0.036644975 | 0.09750796 | 0.375815215 | 21.66843551 |
| ML1h07G01701 | MlPAL5 | Mba02_g01510 | MbPAL5 | 0.054050785 | 0.110011401 | 0.491319847 | 24.44697801 |
| ML1h07G01701 | MlPAL5 | Mba08_g14430 | MbPAL3 | 0.060001618 | 0.389345547 | 0.15410891 | 86.5212327 |

| | | | | | | | |
|---|---|---|---|---|---|---|---|
| ML1h07G01701 | MlPAL5 | Mba11_g14220 | MbPAL6 | 0.068781133 | 0.460907277 | 0.149229869 | 102.4238393 |
| ML1h02G00424 | MlPAL6 | Mba01_g14280 | MbPAL8 | 0.012518546 | 0.061501724 | 0.203547895 | 13.66704973 |
| ML1h01G01719 | MlPAL8 | Mba05_g13940 | MbPAL7 | 0.014404701 | 0.09511955 | 0.151437864 | 21.13767777 |
| ML1h06G02093 | MlPAL9 | Mba09_g14610 | MbPAL1 | 0.009458719 | 0.064210315 | 0.147308411 | 14.26895879 |
| ML1h02G03012 | MlTT8 | Mba06_g23950 | MbTT8 | 0.079639792 | 0.131859868 | 0.603972939 | 29.30219288 |
| ML1h03G03510 | MlTT8 | Mba03_g18730 | MbTT8.2 | 0.10088637 | 0.167042755 | 0.60395538 | 37.12061218 |
| ML1h03G03510 | MlTT8 | Mba06_g23950 | MbTT8 | 0.110357306 | 0.439747898 | 0.250955846 | 97.72175511 |
| ML1h01G02333 | MlTTG1 | Mba05_g07670 | MbTTG1 | 0.017799708 | 0.051506596 | 0.345581139 | 11.44591026 |
| ML1h04G00438 | MlTTG1 | Mba04_g36060 | MbTTG1.2 | 0.012767775 | 0.0942591 | 0.13545403 | 20.9464667 |
| ML1h05G03318 | MlUGT75C1 | Mba06_g06490 | MbUGT75C1.3 | 0.0581719 | 0.090908254 | 0.639896793 | 20.20183416 |
| ML1h05G00327 | MlUGT75C1 | Mba10_g24560 | MbUGT75C1 | 0.029042029 | 0.050209949 | 0.578411838 | 11.15776644 |
| ML1h09G00476 | MlUGT78D2 | Mba03_g04830 | MbUGT78D2 | 0.022449298 | 0.08998993 | 0.249464558 | 19.9977623 |
| ML1h06G01843 | MlUGT78D2 | Mba07_g14580 | MbUGT78D2.3 | 0.01987694 | 0.064793764 | 0.306772418 | 14.39861429 |
| ML1h05G00763 | MlUGT78D2.3 | Mba10_g20800 | MbUGT78D2.2 | 0.028071107 | 0.084917665 | 0.330568523 | 18.87059224 |

**Table S3n.** Divergence time, Ka, Ks and Ka/Ks values of orthologous flavonoid biosynthetic genes pairs between *M. lasiocarpa* and *M. beccarii*

| **Seq_1** | **Name1** | **Seq_2** | **Name2** | **Ka** | **Ks** | **Ka_Ks** | **Diversity_year** |
|---|---|---|---|---|---|---|---|
| ML1h07G00167 | Ml4CL1 | Mbe07_t014630 | Mbe4CL3 | 0.010076423 | 0.09251595 | 0.108915526 | 20.55910003 |
| ML1h07G00248 | Ml4CL10 | Mbe07_t013640 | Mbe4CL10 | 0.03940911 | 0.104750957 | 0.376217182 | 23.27799037 |
| ML1h04G02613 | Ml4CL11 | Mbe06_t040140 | Mbe4CL12 | 0.015012353 | 0.097719325 | 0.153627266 | 21.71540549 |
| ML1h07G03382 | Ml4CL12 | Mbe02_t027270 | Mbe4CL11 | 0.031972054 | 0.098930023 | 0.323178477 | 21.98444948 |
| ML1h02G02377 | Ml4CL13 | Mbe01_t005810 | Mbe4CL14 | 0.010669765 | 0.070359151 | 0.15164715 | 15.63536697 |
| ML1h02G02377 | Ml4CL13 | Mbe03_t000920 | Mbe4CL15 | 0.04276394 | 0.336168407 | 0.127209871 | 74.70409045 |
| ML1h02G02377 | Ml4CL13 | Mbe07_t022330 | Mbe4CL16 | 0.070413997 | 0.402578179 | 0.174907634 | 89.46181758 |
| ML1h09G01309 | Ml4CL14 | Mbe01_t005810 | Mbe4CL14 | 0.044883393 | 0.370013245 | 0.121302126 | 82.22516554 |

| | | | | | | | |
|---|---|---|---|---|---|---|---|
| ML1h09G01309 | Ml4CL14 | Mbe03_t000920 | Mbe4CL15 | 0.013145619 | 0.081125862 | 0.16203981 | 18.02796941 |
| ML1h09G01309 | Ml4CL14 | Mbe07_t022330 | Mbe4CL16 | 0.070658987 | 0.463091322 | 0.152581107 | 102.9091826 |
| ML1h08G00557 | Ml4CL15 | Mbe03_t056570 | Mbe4CL13 | 0.014908352 | 0.066327453 | 0.224768951 | 14.73943396 |
| ML1h08G00557 | Ml4CL15 | Mbe04_t013920 | Mbe4CL18 | 0.049930312 | 0.48232639 | 0.103519759 | 107.1836422 |
| ML1h06G03426 | Ml4CL16 | Mbe04_t013920 | Mbe4CL18 | 0.009890253 | 0.113673574 | 0.087005738 | 25.26079414 |
| ML1h06G03426 | Ml4CL16 | Mbe04_t006230 | Mbe4CL17 | 0.041650014 | 0.356321178 | 0.116888966 | 79.18248398 |
| ML1h06G02761 | Ml4CL17 | Mbe04_t006230 | Mbe4CL17 | 0.030218103 | 0.092543426 | 0.326528899 | 20.56520588 |
| ML1h06G02761 | Ml4CL17 | Mbe04_t013920 | Mbe4CL18 | 0.052074284 | 0.369884371 | 0.140785304 | 82.19652681 |
| ML1h09G02144 | Ml4CL18 | Mbe06_t016280 | Mbe4CL1 | 0.028985114 | 0.109919794 | 0.263693309 | 24.42662081 |
| ML1h08G00140 | Ml4CL2 | Mbe05_t019740 | Mbe4CL4 | 0.023156687 | 0.105292316 | 0.219927611 | 23.39829241 |
| ML1h04G04085 | Ml4CL3 | Mbe03_t020730 | Mbe4CL8 | 0.040761295 | 0.077205453 | 0.527958761 | 17.15676723 |
| ML1h03G00142 | Ml4CL4 | Mbe03_t000510 | Mbe4CL2 | 0.034655288 | 0.083689314 | 0.414094534 | 18.59762536 |
| ML1h02G00664 | Ml4CL5 | Mbe01_t019320 | Mbe4CL6 | 0.03139534 | 0.09560869 | 0.328373286 | 21.24637554 |
| ML1h01G00405 | Ml4CL8 | Mbe05_t004680 | Mbe4CL7 | 0.021428529 | 0.110480097 | 0.19395827 | 24.5511327 |
| ML1h01G00972 | Ml4CL9 | Mbe05_t011100 | Mbe4CL9 | 0.027623964 | 0.057975524 | 0.476476309 | 12.88344976 |
| ML1h06G01342 | MlANR2 | Mbe07_t028460 | MbeANR | 0.03809047 | 0.133636416 | 0.285030618 | 29.69698136 |
| ML1h08G01785 | MlC1 | Mbe03_t051550 | MbeC1 | 0.044136742 | 0.07077383 | 0.623630828 | 15.72751775 |
| ML1h05G03234 | MlC1 | Mbe01_t033880 | MbeC1 | 0.023385848 | 0.115017234 | 0.203324732 | 25.55938543 |
| ML1h05G01322 | MlC1.3 | Mbe02_t039140 | MbeC1.3 | 0.022176906 | 0.12341204 | 0.179698075 | 27.42489784 |
| ML1h05G01754 | MlC4H1 | Mbe01_t038520 | MbeC4H1 | 0.032982141 | 0.326864996 | 0.100904477 | 72.63666589 |
| ML1h05G01754 | MlC4H1 | Mbe02_t033650 | MbeC4H3 | 0.007895964 | 0.078294106 | 0.100850046 | 17.39869011 |
| ML1h08G00252 | MlC4H2 | Mbe10_t000860 | MbeC4H2 | 0.007299883 | 0.060931741 | 0.11980427 | 13.54038692 |
| ML1h05G04363 | MlC4H3 | Mbe01_t038520 | MbeC4H1 | 0.006131421 | 0.075352898 | 0.081369412 | 16.74508839 |
| ML1h05G04363 | MlC4H3 | Mbe02_t033650 | MbeC4H3 | 0.036216244 | 0.327677441 | 0.11052407 | 72.81720905 |
| ML1h08G03392 | MlC4H4 | Mbe08_t025090 | MbeC4H4 | 0.016649095 | 0.09281701 | 0.179375478 | 20.62600221 |

| | | | | | | | |
|---|---|---|---|---|---|---|---|
| ML1h09G01419 | MlCHI1 | Mbe04_t020690 | MbeCHI1 | 0.045762577 | 0.086204466 | 0.530860854 | 19.15654808 |
| ML1h04G02595 | MlCHI2 | Mbe06_t039860 | MbeCHI2 | 0.02513763 | 0.10603639 | 0.237066065 | 23.56364215 |
| ML1h05G01818 | MlCHS1 | Mbe01_t047580 | MbeCHS4 | 0.035033644 | 0.337861541 | 0.103692311 | 75.08034245 |
| ML1h05G01818 | MlCHS1 | Mbe01_t038030 | MbeCHS2 | 0.112735691 | 0.407157206 | 0.276884922 | 90.47937908 |
| ML1h05G01818 | MlCHS1 | Mbe11_t002790 | MbeCHS5 | 0.013577591 | 0.0739059 | 0.183714578 | 16.4235334 |
| ML1h05G03634 | MlCHS2 | Mbe01_t047580 | MbeCHS4 | 0.012443724 | 0.06605734 | 0.188377618 | 14.67940887 |
| ML1h05G03634 | MlCHS2 | Mbe01_t038030 | MbeCHS2 | 0.109445378 | 0.352444381 | 0.310532339 | 78.32097359 |
| ML1h05G03634 | MlCHS2 | Mbe11_t002790 | MbeCHS5 | 0.031478154 | 0.312177109 | 0.100834279 | 69.37269091 |
| ML1h05G04408 | MlCHS3 | Mbe01_t038030 | MbeCHS2 | 0.035270178 | 0.139237302 | 0.253309834 | 30.94162258 |
| ML1h05G04408 | MlCHS3 | Mbe01_t047580 | MbeCHS4 | 0.123324989 | 0.344314185 | 0.358175743 | 76.51426331 |
| ML1h05G04408 | MlCHS3 | Mbe11_t002790 | MbeCHS5 | 0.123873307 | 0.348262531 | 0.355689447 | 77.39167362 |
| ML1h04G00849 | MlDFR1 | Mbe04_t036630 | MbeDFR1 | 0.019729606 | 0.075405922 | 0.261645313 | 16.75687154 |
| ML1h06G03238 | MlDFR2 | Mbe04_t011700 | MbeDFR3 | 0.017047918 | 0.077851607 | 0.218979656 | 17.30035715 |
| ML1h03G02164 | MlDFR3 | Mbe06_t019010 | MbeDFR2 | 0.021045326 | 0.126661846 | 0.16615363 | 28.14707693 |
| ML1h07G00690 | MlF3'5'H1 | Mbe07_t008510 | MbeF3'5'H1 | 0.018928449 | 0.045296783 | 0.417876244 | 10.06595171 |
| ML1h04G03337 | MlF3'5'H12 | Mbe06_t049500 | MbeF3'5'H2 | 0.026599406 | 0.076785167 | 0.346413334 | 17.06337046 |
| ML1h07G00951 | MlF3'5'H3 | Mbe07_t005300 | MbeF3'5'H10 | 0.048966095 | 0.088483743 | 0.553390869 | 19.66305394 |
| ML1h08G03854 | MlF3'5'H4 | Mbe09_t008950 | MbeF3'5'H8 | 0.019460073 | 0.102504679 | 0.189845704 | 22.77881766 |
| ML1h06G02237 | MlF3'5'H6 | Mbe08_t039500 | MbeF3'5'H7 | 0.040042495 | 0.068595224 | 0.583750485 | 15.243383 |
| ML1h06G02237 | MlF3'5'H6 | Mbe09_t008950 | MbeF3'5'H8 | 0.12945665 | 0.450429109 | 0.287407379 | 100.0953575 |
| ML1h07G01736 | MlF3'5'H7 | Mbe02_t005590 | MbeF3'5'H3 | 0.093553434 | 0.402936374 | 0.232179173 | 89.54141643 |
| ML1h07G01736 | MlF3'5'H7 | Mbe02_t006410 | MbeF3'5'H5 | 0.023102671 | 0.091324072 | 0.252974602 | 20.29423811 |
| ML1h08G02519 | MlF3H2 | Mbe07_t035880 | MbeF3H1 | 0.016578532 | 0.083696573 | 0.19807898 | 18.59923837 |
| ML1h03G02252 | MlF3H3 | Mbe06_t019910 | MbeF3'H | 0.039803002 | 0.116291814 | 0.342268304 | 25.8426253 |
| ML1h03G01030 | MlFLS1 | Mbe08_t015900 | MbeFLS1 | 0.02525715 | 0.081160188 | 0.31120122 | 18.03559744 |

Musaceae flavonoid biosynthetic genes

| | | | | | | | |
|---|---|---|---|---|---|---|---|
| ML1h06G02043 | MlFLS4 | Mbe03_t008100 | MbeFLS4 | 0.083812345 | 0.370742318 | 0.226066303 | 82.38718168 |
| ML1h06G02043 | MlFLS4 | Mbe08_t036240 | MbeFLS3 | 0.071015216 | 0.151376519 | 0.469129667 | 33.63922653 |
| ML1h09G00678 | MlFLS5 | Mbe03_t008100 | MbeFLS4 | 0.010529953 | 0.076441995 | 0.137750885 | 16.98711006 |
| ML1h01G01259 | MlLAR | Mbe05_t014700 | MbeLAR | 0.028341715 | 0.099615685 | 0.284510566 | 22.13681884 |
| ML1h01G01735 | MlLDOX | Mbe03_t031300 | MbeLDOX | 0.025669648 | 0.063106582 | 0.406766562 | 14.02368496 |
| ML1h04G02672 | MlPAL1 | Mbe05_t009880 | MbePAL1 | 0.03823436 | 0.353889393 | 0.108040425 | 78.64208744 |
| ML1h04G02672 | MlPAL1 | Mbe06_t040740 | MbePAL2 | 0.013181634 | 0.060678659 | 0.217236747 | 13.48414634 |
| ML1h01G00856 | MlPAL2 | Mbe05_t009880 | MbePAL1 | 0.010671869 | 0.052893976 | 0.201759626 | 11.75421678 |
| ML1h01G00856 | MlPAL2 | Mbe06_t040740 | MbePAL2 | 0.035481209 | 0.346049955 | 0.102532042 | 76.89999001 |
| ML1h07G00086 | MlPAL3 | Mbe05_t009880 | MbePAL1 | 0.058552059 | 0.438930994 | 0.133396956 | 97.54022078 |
| ML1h07G00086 | MlPAL3 | Mbe06_t048840 | MbePAL5 | 0.084132775 | 0.41693586 | 0.201788292 | 92.65241337 |
| ML1h07G00086 | MlPAL3 | Mbe07_t015550 | MbePAL3 | 0.013189914 | 0.089628387 | 0.147162236 | 19.91741943 |
| ML1h04G03282 | MlPAL4 | Mbe06_t048840 | MbePAL5 | 0.028331068 | 0.049645911 | 0.570662669 | 11.03242478 |
| ML1h07G01701 | MlPAL5 | Mbe07_t015550 | MbePAL3 | 0.061111894 | 0.363372944 | 0.16817954 | 80.749543 |
| ML1h02G00424 | MlPAL6 | Mbe01_t022110 | MbePAL7 | 0.013468727 | 0.068727549 | 0.195972752 | 15.27278869 |
| ML1h06G02093 | MlPAL9 | Mbe08_t037020 | MbePAL8 | 0.008271464 | 0.056553144 | 0.146260017 | 12.56736541 |
| ML1h02G03012 | MlTT8 | Mbe06_t001720 | MbeTT8 | 0.04835441 | 0.097771363 | 0.494566181 | 21.72696964 |
| ML1h03G03510 | MlTT8 | Mbe06_t034420 | MbeTT8 | 0.025297219 | 0.143805131 | 0.175913191 | 31.95669587 |
| ML1h01G02333 | MlTTG1 | Mbe03_t038300 | MbeTTG1 | 0.013597771 | 0.080829318 | 0.168228206 | 17.96207073 |
| ML1h04G00438 | MlTTG1 | Mbe04_t041380 | MbeTTG1 | 0.011371931 | 0.085102379 | 0.133626473 | 18.91163975 |
| ML1h05G03318 | MlUGT75C1 | Mbe01_t034870 | MbeUGT75C1.3 | 0.155528597 | 0.406098334 | 0.382982603 | 90.24407426 |
| ML1h05G00327 | MlUGT75C1 | Mbe02_t051150 | MbeUGT75C1 | 0.028561281 | 0.072766674 | 0.392504966 | 16.17037195 |
| ML1h09G00476 | MlUGT78D2 | Mbe03_t010580 | MbeUGT78D2.3 | 0.01817701 | 0.086440298 | 0.210283982 | 19.20895504 |
| ML1h06G01843 | MlUGT78D2 | Mbe07_t034770 | MbeUGT78D2 | 0.020363804 | 0.054236077 | 0.375466014 | 12.05246159 |
| ML1h05G00763 | MlUGT78D2.3 | Mbe02_t045890 | MbeUGT78D2 | 0.030101858 | 0.09475826 | 0.317670019 | 21.057391 |

Musaceae flavonoid biosynthetic genes

| | | | | | | | |
|---|---|---|---|---|---|---|---|
| ML1h08G03185 | MlUGT79B1 | Mbe08_t027530 | MbeUGT79B1 | 0.034016681 | 0.086598694 | 0.392808247 | 19.24415425 |

**Table S3h.** Divergence time, Ka, Ks and Ka/Ks values of orthologous flavonoid biosynthetic genes pairs between *M. lasiocarpa* and *M. schizocarpa*

| **Seq_1** | **Name1** | **Seq_2** | **Name2** | **Ka** | **Ks** | **Ka_Ks** | **Diversity_year** |
|---|---|---|---|---|---|---|---|
| ML1h07G00167 | Ml4CL1 | Ms08t219500 | Ms4CL4 | 0.00923024 | 0.090031048 | 0.102522857 | 20.00689963 |
| ML1h07G00248 | Ml4CL10 | Ms08t218760 | Ms4CL1 | 0.030282406 | 0.079142102 | 0.382633336 | 17.58713375 |
| ML1h04G02613 | Ml4CL11 | Ms11t319670 | Ms4CL2.2 | 0.011032508 | 0.063711756 | 0.173162829 | 14.1581679 |
| ML1h07G03382 | Ml4CL12 | Ms02t041000 | Ms4CL2 | 0.025954871 | 0.084858624 | 0.305860146 | 18.8574721 |
| ML1h02G02377 | Ml4CL13 | Ms01t017810 | Ms4CL10 | 0.008196803 | 0.080886391 | 0.101337231 | 17.97475353 |
| ML1h02G02377 | Ml4CL13 | Ms03t058920 | Ms4CL11 | 0.040567379 | 0.343122745 | 0.118229933 | 76.24949899 |
| ML1h02G02377 | Ml4CL13 | Ms07t183930 | Ms4CL14 | 0.078959606 | 0.42731524 | 0.184780693 | 94.95894222 |
| ML1h09G01309 | Ml4CL14 | Ms01t017810 | Ms4CL10 | 0.043602916 | 0.359772179 | 0.121195909 | 79.94937311 |
| ML1h09G01309 | Ml4CL14 | Ms03t058920 | Ms4CL11 | 0.018116574 | 0.081384019 | 0.222606041 | 18.08533755 |
| ML1h09G01309 | Ml4CL14 | Ms07t183930 | Ms4CL14 | 0.077380379 | 0.458010428 | 0.168948945 | 101.7800951 |
| ML1h08G00557 | Ml4CL15 | Ms04t089570 | Ms4CL13 | 0.052121214 | 0.457085002 | 0.114029587 | 101.5744449 |
| ML1h08G00557 | Ml4CL15 | Ms09t262230 | Ms4CL16 | 0.023275401 | 0.053423722 | 0.435675395 | 11.87193821 |
| ML1h06G03426 | Ml4CL16 | Ms04t089570 | Ms4CL13 | 0.01238333 | 0.096739646 | 0.128006777 | 21.49769919 |
| ML1h06G03426 | Ml4CL16 | Ms04t083350 | Ms4CL12 | 0.043113495 | 0.38911356 | 0.110799262 | 86.46968005 |
| ML1h06G02761 | Ml4CL17 | Ms04t083350 | Ms4CL12 | 0.027662561 | 0.111822524 | 0.247379154 | 24.84944978 |
| ML1h06G02761 | Ml4CL17 | Ms04t089570 | Ms4CL13 | 0.058254122 | 0.346286182 | 0.168225371 | 76.95248487 |
| ML1h09G02144 | Ml4CL18 | Ms06t178240 | Ms4CL9 | 0.025259443 | 0.064217331 | 0.393343088 | 14.2705181 |
| ML1h08G00140 | Ml4CL2 | Ms07t196560 | Ms4CL15 | 0.033545964 | 0.116311659 | 0.288414458 | 25.84703538 |
| ML1h03G00142 | Ml4CL4 | Ms03t059300 | Ms4CL8 | 0.031069531 | 0.080762548 | 0.384702218 | 17.94723285 |
| ML1h02G00664 | Ml4CL5 | Ms01t006680 | Ms4CL7 | 0.018955751 | 0.097798566 | 0.193824427 | 21.73301475 |

| | | | | | | | |
|---|---|---|---|---|---|---|---|
| ML1h01G00405 | Ml4CL8 | Ms05t142250 | Ms4CL5 | 0.020989197 | 0.096402062 | 0.217725601 | 21.42268035 |
| ML1h01G00972 | Ml4CL9 | Ms05t136800 | Ms4CL1.2 | 0.022515762 | 0.084759757 | 0.265642127 | 18.8355016 |
| ML1h06G01342 | MlANR2 | Ms07t188910 | MsANR3 | 0.041563768 | 0.102780639 | 0.404392968 | 22.84014204 |
| ML1h07G01339 | MlANR4 | Ms08t207990 | MsANR1 | 0.030992143 | 0.068550033 | 0.452109826 | 15.23334062 |
| ML1h08G01785 | MlC1 | Ms09t266370 | MsC1 | 0.045895142 | 0.070907754 | 0.647251383 | 15.75727861 |
| ML1h05G03234 | MlC1 | Ms06t151760 | MsC1.2 | 0.030777702 | 0.088377986 | 0.348250778 | 19.63955236 |
| ML1h05G01754 | MlC4H1 | Ms10t281630 | MsC4H1 | 0.008782302 | 0.078180649 | 0.112333446 | 17.37347761 |
| ML1h08G00252 | MlC4H2 | Ms07t197940 | MsC4H2 | 0.009129059 | 0.083699665 | 0.109069243 | 18.59992549 |
| ML1h05G04363 | MlC4H3 | Ms10t281630 | MsC4H1 | 0.033031223 | 0.324991547 | 0.101637177 | 72.22034377 |
| ML1h08G03392 | MlC4H4 | Ms09t244050 | MsC4H3 | 0.016649095 | 0.086469445 | 0.192543106 | 19.21543221 |
| ML1h09G01419 | MlCHI1 | Ms04t093250 | MsCHI2 | 0.040556944 | 0.071447141 | 0.567649637 | 15.87714247 |
| ML1h04G02595 | MlCHI2 | Ms11t319860 | MsCHI1 | 0.021357434 | 0.119111687 | 0.179305946 | 26.46926375 |
| ML1h05G01818 | MlCHS1 | Ms06t158650 | MsCHS2 | 0.045443547 | 0.35108469 | 0.129437563 | 78.01881991 |
| ML1h05G01818 | MlCHS1 | Ms10t280970 | MsCHS1 | 0.013587841 | 0.085435496 | 0.159042104 | 18.98566568 |
| ML1h05G03634 | MlCHS2 | Ms06t158650 | MsCHS2 | 0.021695138 | 0.089491504 | 0.242426788 | 19.88700081 |
| ML1h05G03634 | MlCHS2 | Ms10t280970 | MsCHS1 | 0.030917458 | 0.346683482 | 0.08918065 | 77.04077379 |
| ML1h05G04408 | MlCHS3 | MsSC51t327380 | MsCHS4 | 0.030420239 | 0.122722068 | 0.247879124 | 27.27157063 |
| ML1h05G04408 | MlCHS3 | Ms06t158650 | MsCHS2 | 0.130421122 | 0.344569991 | 0.378504006 | 76.57110904 |
| ML1h05G04408 | MlCHS3 | Ms10t280970 | MsCHS1 | 0.125962439 | 0.372756191 | 0.337921789 | 82.8347092 |
| ML1h04G00849 | MlDFR1 | Ms04t107810 | MsDFR1 | 0.016519092 | 0.082525913 | 0.200168547 | 18.33909185 |
| ML1h06G03238 | MlDFR2 | Ms04t087800 | MsDFR2 | 0.014813772 | 0.079750456 | 0.18575157 | 17.72232354 |
| ML1h03G02164 | MlDFR3 | Ms03t076720 | MsDFR3 | 0.018504228 | 0.067651466 | 0.273522941 | 15.0336592 |
| ML1h07G00690 | MlF3'5'H1 | Ms08t214430 | MsF3'5'H1 | 0.019552574 | 0.060069614 | 0.325498572 | 13.3488032 |
| ML1h04G03337 | MlF3'5'H12 | Ms11t312500 | MsF3'5'H8 | 0.022633944 | 0.09334291 | 0.242481663 | 20.74286885 |
| ML1h07G00951 | MlF3'5'H3 | Ms08t211840 | MsF3'5'H3 | 0.018696717 | 0.049231614 | 0.379770549 | 10.9403587 |

| | | | | | | | |
|---|---|---|---|---|---|---|---|
| ML1h06G02237 | MlF3'5'H6 | Ms09t255750 | MsF3'5'H2 | 0.029069712 | 0.054920671 | 0.529303666 | 12.20459354 |
| ML1h07G01736 | MlF3'5'H7 | Ms02t024400 | MsF3'5'H5 | 0.039092406 | 0.129935677 | 0.300859681 | 28.87459479 |
| ML1h07G01736 | MlF3'5'H7 | Ms11t312500 | MsF3'5'H8 | 0.148718793 | 0.639189586 | 0.232667735 | 142.0421301 |
| ML1h07G02004 | MlF3H1 | Ms02t027200 | MsF3H | 0.015969061 | 0.095690424 | 0.166882536 | 21.26453865 |
| ML1h03G02252 | MlF3H3 | Ms03t075990 | MsF3'H | 0.024525943 | 0.070810361 | 0.346360936 | 15.7356357 |
| ML1h03G01030 | MlFLS1 | Ms08t229420 | MsFLS1 | 0.017177584 | 0.090951571 | 0.188865173 | 20.21146019 |
| ML1h06G02043 | MlFLS4 | Ms03t052640 | MsFLS2 | 0.085990274 | 0.354203423 | 0.242770873 | 78.71187174 |
| ML1h09G00678 | MlFLS5 | Ms03t052640 | MsFLS2 | 0.025921942 | 0.064772372 | 0.4002006 | 14.39386047 |
| ML1h01G01259 | MlLAR | Ms05t133790 | MsLAR | 0.035825609 | 0.094112368 | 0.380668452 | 20.9138596 |
| ML1h01G01735 | MlLDOX | Ms05t120140 | MsLDOX | 0.020733028 | 0.045594147 | 0.454730023 | 10.1320327 |
| ML1h04G02672 | MlPAL1 | Ms05t135820 | MsPAL7 | 0.040838047 | 0.345826936 | 0.118088103 | 76.85043015 |
| ML1h04G02672 | MlPAL1 | Ms11t319150 | MsPAL6 | 0.009382452 | 0.05475635 | 0.171349118 | 12.1680778 |
| ML1h01G00856 | MlPAL2 | Ms05t135820 | MsPAL7 | 0.013829622 | 0.057095449 | 0.242219336 | 12.6878776 |
| ML1h01G00856 | MlPAL2 | Ms11t319150 | MsPAL6 | 0.034682727 | 0.350899187 | 0.098839575 | 77.97759704 |
| ML1h07G00086 | MlPAL3 | Ms02t024050 | MsPAL4 | 0.062183002 | 0.392279352 | 0.158517143 | 87.17318925 |
| ML1h07G00086 | MlPAL3 | Ms05t135820 | MsPAL7 | 0.062454791 | 0.417082356 | 0.149742108 | 92.68496804 |
| ML1h07G00086 | MlPAL3 | Ms08t220210 | MsPAL5 | 0.011903147 | 0.085793141 | 0.138742412 | 19.06514241 |
| ML1h04G03282 | MlPAL4 | Ms02t024050 | MsPAL4 | 0.074883672 | 0.382188697 | 0.195933769 | 84.93082147 |
| ML1h04G03282 | MlPAL4 | Ms11t313180 | MsPAL3 | 0.027277344 | 0.078423409 | 0.347821446 | 17.42742432 |
| ML1h07G01701 | MlPAL5 | Ms02t024050 | MsPAL4 | 0.021517059 | 0.077436879 | 0.277865779 | 17.20819535 |
| ML1h07G01701 | MlPAL5 | Ms08t220210 | MsPAL5 | 0.062240548 | 0.415180286 | 0.1499121 | 92.26228588 |
| ML1h02G00424 | MlPAL6 | Ms01t004140 | MsPAL1 | 0.014089331 | 0.064738241 | 0.217635369 | 14.38627586 |
| ML1h01G01719 | MlPAL8 | Ms05t120010 | MsPAL2 | 0.00936147 | 0.088580192 | 0.105683555 | 19.68448705 |
| ML1h06G02093 | MlPAL9 | Ms09t254450 | MsPAL8 | 0.008864915 | 0.060370836 | 0.146841015 | 13.4157413 |
| ML1h03G03510 | MlTT8 | Ms03t064490 | MsTT8.2 | 0.02817784 | 0.14049816 | 0.200556645 | 31.22181324 |

| | | | | | | | |
|---|---|---|---|---|---|---|---|
| ML1h01G02333 | MlTTG1 | Ms05t125970 | MsTTG1.2 | 0.011363854 | 0.046346789 | 0.245191822 | 10.29928654 |
| ML1h04G00438 | MlTTG1 | Ms04t112120 | MsTTG1 | 0.012807142 | 0.075647617 | 0.169299999 | 16.81058163 |
| ML1h05G03318 | MlUGT75C1 | Ms06t152730 | MsUGT75C1.3 | 0.054437732 | 0.103530107 | 0.525815472 | 23.00669043 |
| ML1h05G00327 | MlUGT75C1 | Ms10t295740 | MsUGT75C1.4 | 0.028494373 | 0.057058946 | 0.499384843 | 12.67976571 |
| ML1h09G00476 | MlUGT78D2 | Ms03t050760 | MsUGT78D2.3 | 0.029528418 | 0.081000663 | 0.364545382 | 18.00014729 |
| ML1h06G01843 | MlUGT78D2 | Ms07t193970 | MsUGT78D2.2 | 0.026196937 | 0.068347186 | 0.383292112 | 15.1882636 |
| ML1h05G00763 | MlUGT78D2.3 | Ms10t291590 | MsUGT78D2 | 0.039014596 | 0.115802655 | 0.336905886 | 25.73392325 |
| ML1h08G03185 | MlUGT79B1 | Ms09t246270 | MsUGT79B1 | 0.025212458 | 0.08083589 | 0.311896831 | 17.96353121 |

**Table S4a.** Tandem duplication FBG pairs in Musaceae

| Gene1 | Name1 | Gene2 | Name2 | Gene3 | Name3 | Gene4 | Name4 |
|---|---|---|---|---|---|---|---|
| ***M. beccarii*** | | | | | | | |
| Mbe01_t034870 | ***MbeUGT75C1.3*** | Mbe01_t034880 | ***MbeUGT75C1.2*** | | | | |
| Mbe02_t005590 | ***MbeF3'5'H3*** | Mbe02_t005600 | ***MbeF3'5'H6*** | | | | |
| Mbe09_t008950 | ***MbeF3'5'H8*** | Mbe09_t008960 | ***MbeF3'5'H9*** | | | | |
| ***M. lasiocarpa*** | | | | | | | |
| ML1h01G01720 | ***MlPAL7*** | ML1h01G01719 | ***MlPAL8*** | | | | |
| ML1h05G00635 | ***MlFLS3*** | ML1h05G00636 | ***MlFLS2*** | | | | |
| ML1h05G03317 | ***MlUGT75C1.3*** | ML1h05G03318 | ***MlUGT75C1.2*** | | | | |
| ML1h06G01340 | ***MlANR1*** | ML1h06G01342 | ***MlANR2*** | ML1h06G01341 | ***MlANR3*** | | |
| ML1h07G01736 | ***MlF3'5'H7*** | ML1h07G01737 | ***MlF3'5'H8*** | | | | |
| ***E. glaucum*** | | | | | | | |
| Eg05_t014870 | ***EgPAL3*** | Eg05_t014880 | ***EgPAL2*** | | | | |
| Eg08_t037030 | ***EgUGT75C1.2*** | Eg08_t037040 | ***EgUGT75C1.3*** | | | | |
| Eg08_t049390 | ***EgCHS4*** | Eg08_t049400 | ***EgCHS5*** | Eg08_t049410 | ***EgCHS6*** | | |
| ***M. acuminata*** | | | | | | | |
| Macma4_06_g07190.1 | ***MaUGT75C1.4*** | Macma4_06_g07200.1 | ***MaUGT75C1.3*** | Macma4_06_g07210.1 | ***MaUGT75C1.2*** | | |
| Macma4_06_g10010.1 | ***MaCHS5*** | Macma4_06_g10020.1 | ***MaCHS4*** | | | | |
| Macma4_07_g10660.1 | ***MaANR2*** | Macma4_07_g10670.1 | ***MaANR3*** | Macma4_07_g10680.1 | ***MaANR4*** | | |
| Macma4_02_g03240.1 | ***MaF3'5'H8*** | Macma4_02_g03250.1 | ***MaF3'5'H9*** | Macma4_02_g03260.1 | ***MaF3'5'H10*** | | |
| Macma4_02_g03290.1 | ***MaF3'5'H11*** | Macma4_02_g03300.1 | ***MaF3'5'H12*** | | | | |
| ***M. balbisiana*** | | | | | | | |
| Mba06_g06480.1 | ***MbUGT75C1.2*** | Mba06_g06490.1 | ***MbUGT75C1.3*** | | | | |

| | | | | | | | |
|---|---|---|---|---|---|---|---|
| Mba06_g09080.1 | ***MbCHS5*** | Mba06_g09090.1 | ***MbCHS6*** | | | | |
| Mba06_g09110.1 | ***MbCHS4*** | Mba06_g09120.1 | ***MbCHS7*** | | | | |
| Mba02_g01890.1 | ***MbF3'5'H7*** | Mba02_g01900.1 | ***MbF3'5'H8*** | Mba02_g01910.1 | ***MbF3'5'H9*** | Mba02_g01920.1 | ***MbF3'5'H10*** |
| ***M. schizocarpa*** | | | | | | | |
| Ms02t024400 | ***MsF3'5'H5*** | Ms02t024410 | ***MsF3'5'H6*** | | | | |
| Ms06t152710 | ***MsUGT75C1***.1 | Ms06t152720 | ***MsUGT75C1.2*** | Ms06t152730 | ***MsUGT75C1.3*** | | |
| Ms07t188910 | ***MsANR3*** | Ms07t188920 | ***MsANR2*** | | | | |
| MsSC51t327370 | ***MsCHS3*** | MsSC51t327380 | ***MsCHS4*** | | | | |

**Table S4b.** Segmental duplication FBG pairs in Musaceae.

| Gene1 | Name1 | Gene2 | Name2 | |
|---|---|---|---|---|
| Mbe02_t027270 | ***Mbe4CL11*** | Mbe06_t040140 | ***Mbe4CL12*** | |
| Mbe03_t056570 | ***Mbe4CL13*** | Mbe04_t006230 | ***Mbe4CL17*** | |
| Mbe01_t005810 | ***Mbe4CL14*** | Mbe03_t000920 | ***Mbe4CL15*** | |
| Mbe01_t005810 | ***Mbe4CL14*** | Mbe07_t022330 | ***Mbe4CL16*** | * |
| Mbe03_t000920 | ***Mbe4CL15*** | Mbe07_t022330 | ***Mbe4CL16*** | |
| Mbe05_t011100 | ***Mbe4CL9*** | Mbe07_t013640 | ***Mbe4CL10*** | |
| Mbe01_t038520 | ***MbeC4H1*** | Mbe02_t033650 | ***MbeC4H3*** | |
| Mbe01_t038520 | ***MbeC4H1*** | Mbe10_t000860 | ***MbeC4H2*** | |
| Mbe02_t033650 | ***MbeC4H3*** | Mbe08_t025090 | ***MbeC4H4*** | * |
| Mbe01_t038010 | ***MbeCHS1*** | Mbe01_t047580 | ***MbeCHS4*** | |
| Mbe01_t038010 | ***MbeCHS1*** | Mbe11_t002790 | ***MbeCHS5*** | |
| Mbe01_t047580 | ***MbeCHS4*** | Mbe11_t002790 | ***MbeCHS5*** | |
| Mbe04_t011700 | ***MbeDFR3*** | Mbe06_t019010 | ***MbeDFR2*** | |
| Mbe02_t005590 | ***MbeF3'5'H3*** | Mbe06_t049500 | ***MbeF3'5'H2*** | |

| | | | | |
|---|---|---|---|---|
| Mbe08_t039500 | ***MbeF3'5'H7*** | Mbe09_t008950 | ***MbeF3'5'H8*** | |
| Mbe02_t047300 | ***MbeFLS2*** | Mbe03_t008100 | ***MbeFLS4*** | * |
| Mbe02_t047300 | ***MbeFLS2*** | Mbe08_t036240 | ***MbeFLS3*** | |
| Mbe05_t009880 | ***MbePAL1*** | Mbe06_t040740 | ***MbePAL2*** | |
| Mbe05_t009880 | ***MbePAL1*** | Mbe07_t015550 | ***MbePAL3*** | * |
| Mbe06_t040740 | ***MbePAL2*** | Mbe07_t015550 | ***MbePAL3*** | |
| Mbe06_t048840 | ***MbePAL5*** | Mbe07_t015550 | ***MbePAL3*** | |
| Mbe06_t001720 | ***MbeTT8.2*** | Mbe06_t034420 | ***MbeTT8.1*** | |
| Mbe02_t045890 | ***MbeUGT78D2.2*** | Mbe03_t010580 | ***MbeUGT78D2.3*** | |
| Mbe02_t045890 | ***MbeUGT78D2.2*** | Mbe07_t034770 | ***MbeUGT78D2.1*** | * |
| Mbe03_t010580 | ***MbeUGT78D2.3*** | Mbe07_t034770 | ***MbeUGT78D2.1*** | |
| Eg02_t018790 | ***EgPAL4*** | Eg09_t011330 | ***EgPAL5*** | |
| Eg02_t001010 | ***EgPAL6*** | Eg09_t011330 | ***EgPAL5*** | |
| Eg02_t001010 | ***EgPAL6*** | Eg09_t017980 | ***EgPAL7*** | * |
| Eg02_t001010 | ***EgPAL6*** | Eg05_t024420 | ***EgPAL8*** | |
| Eg05_t024420 | ***EgPAL8*** | Eg09_t017980 | ***EgPAL7*** | |
| Eg08_t019660 | ***EgC4H1*** | Eg08_t048930 | ***EgC4H3*** | |
| Eg07_t034110 | ***EgC4H4*** | Eg08_t019660 | ***EgC4H1*** | * |
| Eg07_t003130 | ***EgC4H2*** | Eg08_t048930 | ***EgC4H3*** | |
| Eg02_t002700 | ***Eg4CL1.1*** | Eg05_t023200 | ***Eg4CL1.2*** | |
| Eg02_t037340 | ***Eg4CL2.1*** | Eg09_t018540 | ***Eg4CL2.2*** | |
| Eg01_t014520 | ***Eg4CL10*** | Eg01_t025930 | ***Eg4CL11*** | |
| Eg04_t032900 | ***Eg4CL13*** | Eg07_t014740 | ***Eg4CL15*** | |
| Eg08_t020390 | ***EgCHS3*** | Eg08_t040870 | ***EgCHS2*** | * |
| Eg08_t040870 | ***EgCHS2*** | Eg08_t049390 | ***EgCHS4*** | |

| | | | | |
|---|---|---|---|---|
| Eg08_t020390 | ***EgCHS3*** | Eg08_t049390 | ***EgCHS4*** | |
| Eg01_t007540 | ***EgFLS2*** | Eg08_t007020 | ***EgFLS3*** | |
| Eg06_t008220 | ***EgUGT78D2.2*** | Eg08_t008340 | ***EgUGT78D2.1*** | |
| Eg01_t005290 | ***EgUGT78D2.3*** | Eg08_t008340 | ***EgUGT78D2.1*** | * |
| Eg01_t005290 | ***EgUGT78D2.3*** | Eg06_t008220 | ***EgUGT78D2.2*** | |
| Eg03_t004200 | ***EgTT8.2*** | Eg04_t020700 | ***EgTT8.1*** | |
| Macma4_08_g14800.1 | ***MaPAL7*** | Macma4_11_g15930.1 | ***MaPAL3*** | |
| Macma4_02_g02760.1 | ***MaPAL4*** | Macma4_08_g14800.1 | ***MaPAL7*** | |
| Macma4_05_g21700.1 | ***MaPAL5*** | Macma4_11_g22170.1 | ***MaPAL6*** | * |
| Macma4_05_g21700.1 | ***MaPAL5*** | Macma4_08_g14800.1 | ***MaPAL7*** | |
| Macma4_08_g14800.1 | ***MaPAL7*** | Macma4_11_g22170.1 | ***MaPAL6*** | |
| Macma4_06_g10450.1 | ***MaC4H3*** | Macma4_10_g16840.1 | ***MaC4H1*** | |
| Macma4_09_g04800.1 | ***MaC4H4*** | Macma4_10_g16840.1 | ***MaC4H1*** | * |
| Macma4_06_g10450.1 | ***MaC4H3*** | Macma4_07_g20380.1 | ***MaC4H2*** | |
| Macma4_05_g22890.1 | ***Ma4CL1.2*** | Macma4_08_g13040.1 | ***Ma4CL1.1*** | |
| Macma4_02_g21360.1 | ***Ma4CL2.1*** | Macma4_11_g22750.1 | ***Ma4CL2.2*** | |
| Macma4_01_g19800 | ***Ma4CL10*** | Macma4_03_g13930 | ***Ma4CL11*** | |
| Macma4_01_g19800 | ***Ma4CL10*** | Macma4_07_g05220 | ***Ma4CL14*** | * |
| Macma4_03_g13930 | ***Ma4CL11*** | Macma4_07_g05220 | ***Ma4CL14*** | |
| Macma4_04_g06110 | ***Ma4CL12*** | Macma4_09_g24760 | ***Ma4CL16*** | |
| Macma4_06_g10010.1 | ***MaCHS5*** | Macma4_06_g18170.1 | ***MaCHS2*** | |
| Macma4_06_g18170.1 | ***MaCHS2*** | Macma4_10_g16140.1 | ***MaCHS6*** | * |
| Macma4_06_g10010.1 | ***MaCHS5*** | Macma4_10_g16140.1 | ***MaCHS6*** | |
| Macma4_07_g16200.1 | ***MaUGT78D2.2*** | Macma4_10_g27580.1 | ***MaUGT78D2.1*** | * |
| Macma4_03_g04880.1 | ***MaUGT78D2.3*** | Macma4_10_g27580.1 | ***MaUGT78D2.1*** | |

Musaceae flavonoid biosynthetic genes

| | | | | |
|---|---|---|---|---|
| Macma4_03_g04880.1 | ***MaUGT78D2.3*** | Macma4_07_g16200.1 | ***MaUGT78D2.2*** | |
| Macma4_03_g19880.1 | ***MaTT8.2*** | Macma4_06_g25800.1 | ***MaTT8.1*** | |
| Macma4_09_g17660.1 | ***MaF3'5'H4*** | Macma4_10_g01240.1 | ***MaF3'5'H5*** | * |
| Macma4_02_g03240.1 | ***MaF3'5'H8*** | Macma4_10_g01240.1 | ***MaF3'5'H5*** | |
| Mba08_g14430 | ***MbPAL3*** | Mba11_g20430 | ***MbPAL2*** | * |
| Mba02_g01510 | ***MbPAL5*** | Mba08_g14430 | ***MbPAL3*** | |
| Mba09_g04380 | ***MbC4H1*** | Mba10_g11450 | ***MbC4H2*** | * |
| Mba06_g09560 | ***MbC4H3*** | Mba10_g11450 | ***MbC4H2*** | |
| Mba02_g18560 | ***Mb4CL2.1*** | Mba11_g20940 | ***Mb4CL2.2*** | |
| Mba01_g28370 | ***Mb4CL10*** | Mba07_g04890 | ***Mb4CL13*** | |
| Mba04_g05720 | ***Mb4CL11*** | Mba04_g12340 | ***Mb4CL12*** | |
| Mba06_g09080 | ***MbCHS5*** | Mba06_g16830 | ***MbCHS2*** | |
| Mba06_g16830 | ***MbCHS2*** | Mba10_g10820 | ***MbCHS8*** | * |
| Mba06_g09080 | ***MbCHS5*** | Mba10_g10820 | ***MbCHS8*** | |
| Mba03_g06940 | ***MbFLS4*** | Mba10_g21980 | ***MbFLS3*** | |
| Mba03_g04830 | ***MbUGT78D2.1*** | Mba10_g20800 | ***MbUGT78D2.2*** | |
| Mba03_g04830 | ***MbUGT78D2.1*** | Mba07_g14580 | ***MbUGT78D2.3*** | * |
| Mba07_g14580 | ***MbUGT78D2.3*** | Mba10_g20800 | ***MbUGT78D2.2*** | |
| Mba03_g18730 | ***MbTT8.2*** | Mba06_g23950 | ***MbTT8.1*** | |
| Mba09_g16030 | ***MbF3'5'H3*** | Mba10_g00740 | ***MbF3'5'H4*** | |
| Mba02_g01840 | ***MbF3'5'H5*** | Mba11_g13650 | ***MbF3'5'H13*** | |
| Ms02t024050 | ***MsPAL4*** | Ms11t313180 | ***MsPAL3*** | |
| Ms08t220210 | ***MsPAL5*** | Ms11t313180 | ***MsPAL3*** | * |
| Ms05t135820 | ***MsPAL7*** | Ms11t313180 | ***MsPAL3*** | |
| Ms02t024050 | ***MsPAL4*** | Ms08t220210 | ***MsPAL5*** | |

Musaceae flavonoid biosynthetic genes

| | | | | |
|---|---|---|---|---|
| Ms08t220210 | ***MsPAL5*** | Ms11t319150 | ***MsPAL6*** | |
| Ms05t135820 | ***MsPAL7*** | Ms08t220210 | ***MsPAL5*** | |
| Ms05t135820 | ***MsPAL7*** | Ms11t319150 | ***MsPAL6*** | |
| Ms09t244050 | ***MsC4H3*** | Ms10t281630 | ***MsC4H1*** | |
| Ms05t136800 | ***Ms4CL1.2*** | Ms08t218760 | ***Ms4CL1.1*** | |
| Ms02t041000 | ***Ms4CL2.1*** | Ms11t319670 | ***Ms4CL2.2*** | |
| Ms01t017810 | ***Ms4CL10*** | Ms03t058920 | ***Ms4CL11*** | |
| Ms01t017810 | ***Ms4CL10*** | Ms07t183930 | ***Ms4CL14*** | * |
| Ms03t058920 | ***Ms4CL11*** | Ms07t183930 | ***Ms4CL14*** | |
| Ms04t083350 | ***Ms4CL12*** | Ms04t089570 | ***Ms4CL13*** | * |
| Ms04t083350 | ***Ms4CL12*** | Ms09t262230 | ***Ms4CL16*** | |
| Ms06t158650 | ***MsCHS2*** | Ms10t280970 | ***MsCHS1*** | |
| Ms10t280970 | ***MsCHS1*** | MsSC51t327370 | ***MsCHS3*** | * |
| Ms06t158650 | ***MsCHS2*** | MsSC51t327370 | ***MsCHS3*** | |
| Ms03t076720 | ***MsDFR3*** | Ms04t087800 | ***MsDFR2*** | |
| Ms07t193970 | ***MsUGT78D2.2*** | Ms10t291590 | ***MsUGT78D2.1*** | |
| Ms03t050760 | ***MsUGT78D2.3*** | Ms10t291590 | ***MsUGT78D2.1*** | * |
| Ms03t050760 | ***MsUGT78D2.3*** | Ms07t193970 | ***MsUGT78D2.2*** | |
